# Decoding the molecular complexity governing corneal wound closure *in vivo*

**DOI:** 10.1101/2025.03.20.642766

**Authors:** Nadège Feret, Alicia Caballero Megido, Solja Kalha, Alison Kuony, Laura Fichter, Sonia Llorens Garcia, Aurore Attina, Naima Nhiri, Eric Jacquet, Jerome Viaralet, Alexandre David, Christophe Hirtz, Karine Loulier, Frederic Michon

**Affiliations:** Institute for Neurosciences of Montpellier, Univ Montpellier, INSERM, Montpellier, France; Institute of Biotechnology, HiLIFE, University of Helsinki, Helsinki, Finland; IRMB-PPC, Univ Montpellier, CHU Montpellier, INSERM CNRS, Montpellier, France; Institut de Chimie des Substances Naturelles, Paris-Saclay University, CNRS, Gif-sur-Yvette, France; Institut de Recherche en Cancérologie de Montpellier, Univ Montpellier, ICM, INSERM, Montpellier, France; Department of Ophthalmology, Gui de Chauliac Hospital, Montpellier, France

**Keywords:** cornea, epithelium, lacrimal gland, tears, proteomics, transcriptomics, epitranscriptomics

## Abstract

The cornea, the transparent outermost layer of the eye, possesses exceptional wound healing capabilities essential for vision preservation. The complexity of the corneal microenvironment is central to its rapid healing; however, the molecular mechanisms orchestrating this process remain poorly defined, limiting therapeutic advancements. Here, we elucidate the extensive remodeling of the corneal molecular landscape following physical injury. Multi-omics analyses—including transcriptomic, epitranscriptomic, and proteomic profiling—uncover significant induction of epithelial cell plasticity driving wound closure. Moreover, lacrimal gland ablation further suppresses Pax6 expression, highlighting its regulatory role. Our multi-omic approach uniquely reveals bilateral remodeling of the molecular environment, a phenomenon constrained by an intact tear film. Collectively, our findings identify novel molecular factors critical to corneal healing, significantly advancing the understanding of epithelial plasticity. These insights will facilitate the translation of cell plasticity research into innovative strategies for tissue and organ regeneration.

## Introduction

The cornea is a transparent, multilayered structure at the anterior surface of the eye, composed primarily of an inner endothelium, a mesenchymal stromal layer, and a stratified squamous epithelium. By virtue of its anatomical position and optical properties, the cornea is elemental to focusing and transmitting light to the retina, while simultaneously protecting underlying ocular structures from environmental and internal perturbations^1^. Owing to its exposed location, the cornea is vulnerable to damage from external insults^2^, as well as age-related changes^3^, and various systemic^4^ and local pathologies^5,6^. Under these conditions, the homeostasis of the corneal microenvironment—which encompasses the corneal epithelium, its innervation, and the tear film— is disrupted, often resulting in progressive thickening, leading to opacification, or thinning, leading to its disintegration. These structural changes lead to corneal blindness, a condition that affects an estimated 28 million people worldwide and is the fourth leading cause of blindness globally^7,8^, underscoring its significant clinical and public health implications.

Among the myriad causes of corneal dysfunction, corneal abrasions are the most frequently encountered ocular injuries in clinical practice^9,10^. A corneal abrasion involves a nonpenetrating disruption of the epithelial barrier, manifesting as ocular pain, photophobia, and excessive tearing^11,12^. Clinical confirmation is typically achieved using fluorescein staining under cobalt-blue illumination, which reveals epithelial defects through characteristic dye uptake^13^. In otherwise healthy eyes with proper care, minimal intervention, and the corneal intrinsic regenerative capacity^14^, these superficial injuries usually heal rapidly (within 24 to 72 hours), restoring corneal clarity. Standard treatments include artificial tears to maintain lubrication, topical NSAIDs for pain management, and prophylactic topical antibiotics to prevent secondary infections^11,15,16^.

However, when not promptly and properly managed, corneal abrasions can lead to persistent epithelial defects^17^ and elevated infection risk. Delayed wound closure compromises the corneal protective function, permitting the ingress of various pathogens^18,19^ and increasing the likelihood of ulcers, perforations, severe pain, and vision loss. Multiple factors contribute to corneal abrasions, including direct mechanical trauma, chemical injuries, foreign bodies, contact lens wear^20^, and inadequate tear film quality^21^. Corneal abrasions are also a common perioperative complication in patients undergoing general anesthesia for non-ocular surgeries^22,23^, and they are an intentional step in certain refractive procedures. For example, photorefractive keratectomy (PRK) and laser-assisted in situ keratomileusis (LASIK) both involve controlled disruptions of the corneal epithelium to correct refractive errors^24,25^. Despite their high success rates^26,27^, these surgeries can be complicated by delayed re-epithelialization^28^, recurrent erosions^29^, or dry eye syndrome^30^, all of which highlight the delicate balance required for effective corneal wound healing.

This duality of corneal abrasion—ranging from rapid, complete healing to severe complications culminating in scarring and persistent vision impairment—demonstrates the complexity of the corneal wound healing process. The need to better understand these mechanisms has driven the development of mouse models that replicate human corneal abrasion. Such models utilize controlled epithelial debridement techniques^31–35^, and allow precise regulation of wound size. The resulting lesions, evaluated by fluorescein staining, typically heal within 72 hours and serve as robust systems to dissect the cellular and molecular mechanisms underlying corneal epithelial homeostasis and repair. To date, however, the exact impact of corneal abrasions on the epithelial interface and its interplay with the tear film remains poorly understood, warranting further investigation.

## Results

### Corneal abrasion triggers profound molecular changes of the cornea and lacrimal gland

To gain deeper insights into the molecular landscape underlying corneal epithelial wound healing, we performed mechanical debridement^34^ and analyzed the transcriptomic signatures of the cornea and lacrimal gland, along with the proteomic profile of the tear film (Fig. 1a, Fig. S1a). Consistent with previous reports^36^, the corneal transcriptome exhibited rapid and significant changes within the first 48 hours of the wound healing process (Fig. 1b, Fig. S1b). Specifically, at 18 hours post-abrasion, 1288 differentially expressed genes (DEGs) showed at least a twofold increase in expression, while 1127 DEGs displayed a reduction of at least 50%. The extensive modulation of the corneal transcriptome led to a reduction in RNA and protein processing (Fig. S2) and an upregulation of immune response effectors, as revealed by Gene Ontology (GO) analysis (Fig. S3). Additionally, KEGG enrichment analysis identified key pathways that were significantly altered during the wound healing process (Table S1, Table S2). While NF-kB, HIF-1, VEGF, and TGF-β signaling appeared to be among the earliest pathways activated, the most pronounced changes occurred 48 hours post-abrasion.

**Figure 1.**
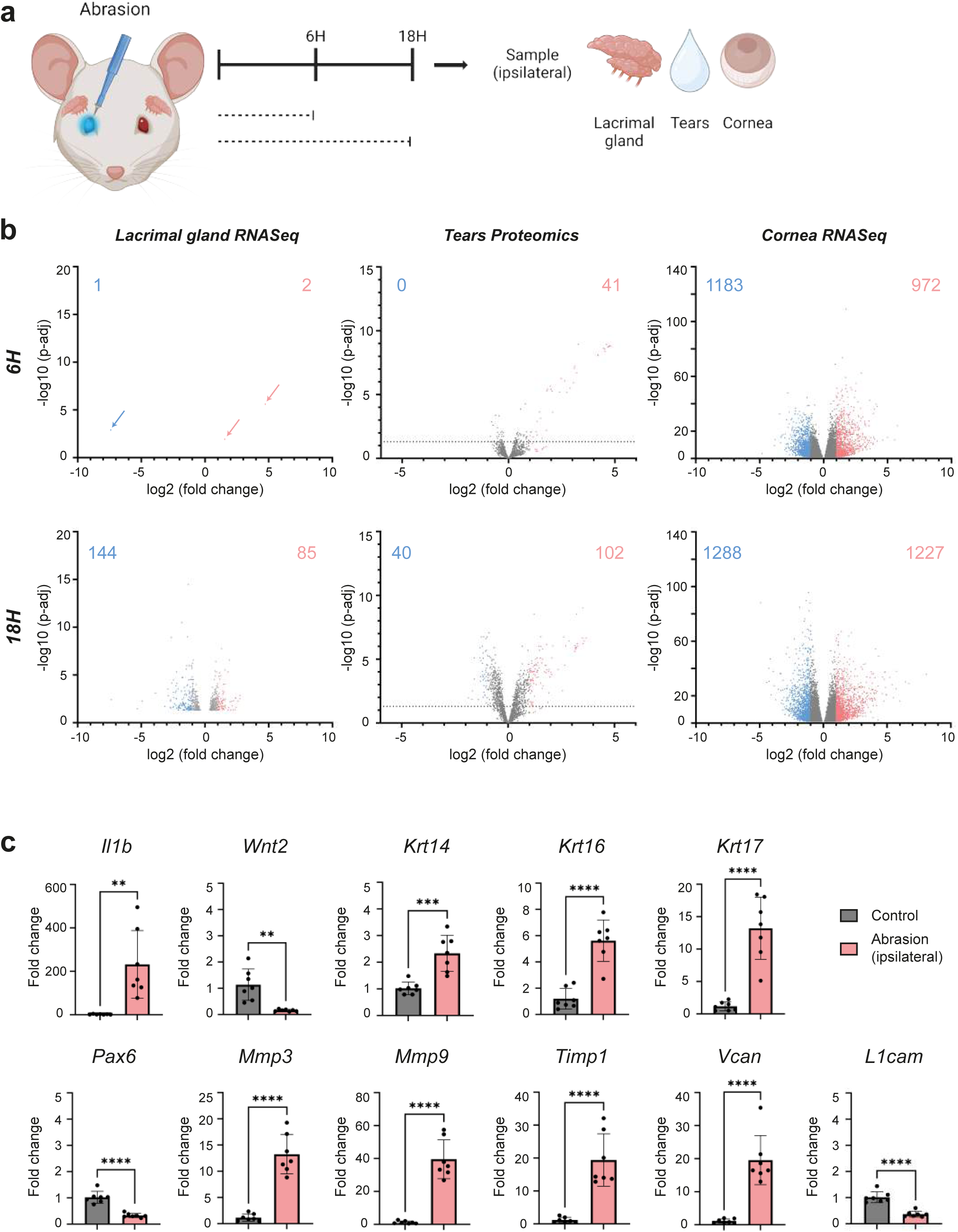
Corneal abrasion induces molecular changes in lacrimal gland, tears and cornea. (a) Schematic overview of the experimental design. Abrasion was performed unilaterally. Samples from the abraded side were taken before the abrasion as a control and at 6H and 18H post-abrasion. (b) Volcano plot representations of lacrimal gland RNA-Seq analysis (n=3 per group), tears proteomics analysis by mass spectrometry (n=7 per group) and cornea RNA-Seq analysis (n=5 per group) at 6H and 18H post-abrasion vs control. For RNA-Seq volcano plots, statistically significant genes are represented only. For proteomics volcano plots, all identified proteins are represented and the limit of statistical significance is shown with the horizontal dotted line. Genes and proteins with a fold change lower than 0.5 are colored in blue and their number is indicated in the top left corner of each plot. Genes and proteins with a fold change higher than 2 are colored in red and their number is indicated in the top right corner of each plot. The genes and proteins with an intermediate fold change are colored in grey. (c) Histograms of RT-qPCR analysis on cornea samples at 18H post-abrasion vs control (n=7 per group). Data are represented as mean ± SD and normalized to a reference gene (*Ppia*). Statistical significance was assessed by unpaired two-tailed t-test (*, p < 0.05; **, p < 0.01; ***, p < 0.001, ****, p < 0.0001).

To further explore post-transcriptional regulation, we analyzed total RNA epitranscriptomic modifications (Table S3), as these reflect alterations in RNA biology and dynamics^37,38^. We observed epitranscriptomic modifications affecting all nucleosides at 18 and 72 hours post-injury. Notably, these modified nucleosides were significantly reduced following abrasion, with most modifications corresponding to methylation events, including m¹A, m³C, m⁷G, and m⁶,⁶A.

The lacrimal gland exhibited subtle yet significant changes in its transcriptomic profile, displaying only 3 DEGs at 6 hours post-injury but more than 600 DEGs at 18 hours post-injury (Fig. 1b). In contrast, tear proteomic analysis showed a slightly stronger response at 6 hours post-injury, with 41 proteins exhibiting at least a twofold change in expression. This modulation intensified by 12 hours post-injury, and the expression profile subsequently remained stable up to 24 hours post-injury (Fig. S1b). KEGG enrichment analysis of proteomic data, specifically focusing on pathway ligands, identified the activation of the MAPK, PI3K-Akt, TGF-β, JAK-STAT, HIF-1, focal adhesion, and calcium signaling pathways starting from 12 hours post-injury (data not shown). Interestingly, EGF was identified as the activating ligand for all these pathways except TGF-β, which was activated by GDF5. Thus, these transcriptomic and proteomic profiles highlight dynamic modulation of tear film composition, as well as molecular alterations in the lacrimal gland and cornea throughout the wound healing process.

To further investigate these molecular alterations, we conducted qPCR analyses on eleven selected DEGs with known relevance to corneal biology at 18 hours post-injury (Fig. 1c). Specifically, we examined two secreted ligands: Il-1β, a known marker of corneal inflammation^39,40^, and Wnt2, expressed by limbal stem cells^41,42^; one transcription factor, Pax6, critical for corneal epithelial cell identity^43,44^; three intracellular markers, Krt14, Krt16, and Krt17, indicative of epithelial differentiation states and stress^45–47^; and five extracellular matrix (ECM)-associated factors: Mmp3 (essential for directional epithelial cell migration^48,49^), Mmp9 (involved in collagen remodeling^50,51^), Timp1 (an inhibitor of MMP9 activity^52,53^), Versican (Vcan), which participates in ECM remodeling^54,55^, and L1cam, an ECM component implicated in cell migration and cohesiveness^56,57^. At this critical time point, wound healing is actively underway, and significant molecular changes have already commenced, as indicated by our global molecular data. Our results demonstrated substantial modulation in the expression of all examined genes. Notably, Wnt2, Pax6, and L1cam were downregulated, whereas the other genes showed increased expression. In particular, the elevated expression of Krt14, a marker of undifferentiated corneal epithelial cells^58^, and decreased Pax6, crucial for maintaining corneal epithelial cell identity, strongly support a dedifferentiation process during the initial phase of wound healing.

To confirm cellular-level dedifferentiation, immunostainings for KRT14 and KRT12, a marker for terminally differentiated corneal epithelial cells^59^, were performed on whole-mounted corneas (Fig. 2a, Fig. 2b). At 18 hours post-injury, we observed elevated KRT14 presence primarily localized at the limbus and wound margins, in contrast to its restriction to the limbus in unwounded corneas. Remarkably, *krt14*+ cells persisted and extended across the entire corneal epithelium at 72 hours post-injury, despite the complete closure of the epithelial wound. KRT12 presence was observed in the whole cornea except for the limbus and some specific regions of the wound border, where KRT14 was overexpressed. This indicates the coexistence of both undifferentiated and differentiated epithelial cell populations during wound healing.

**Figure 2.**
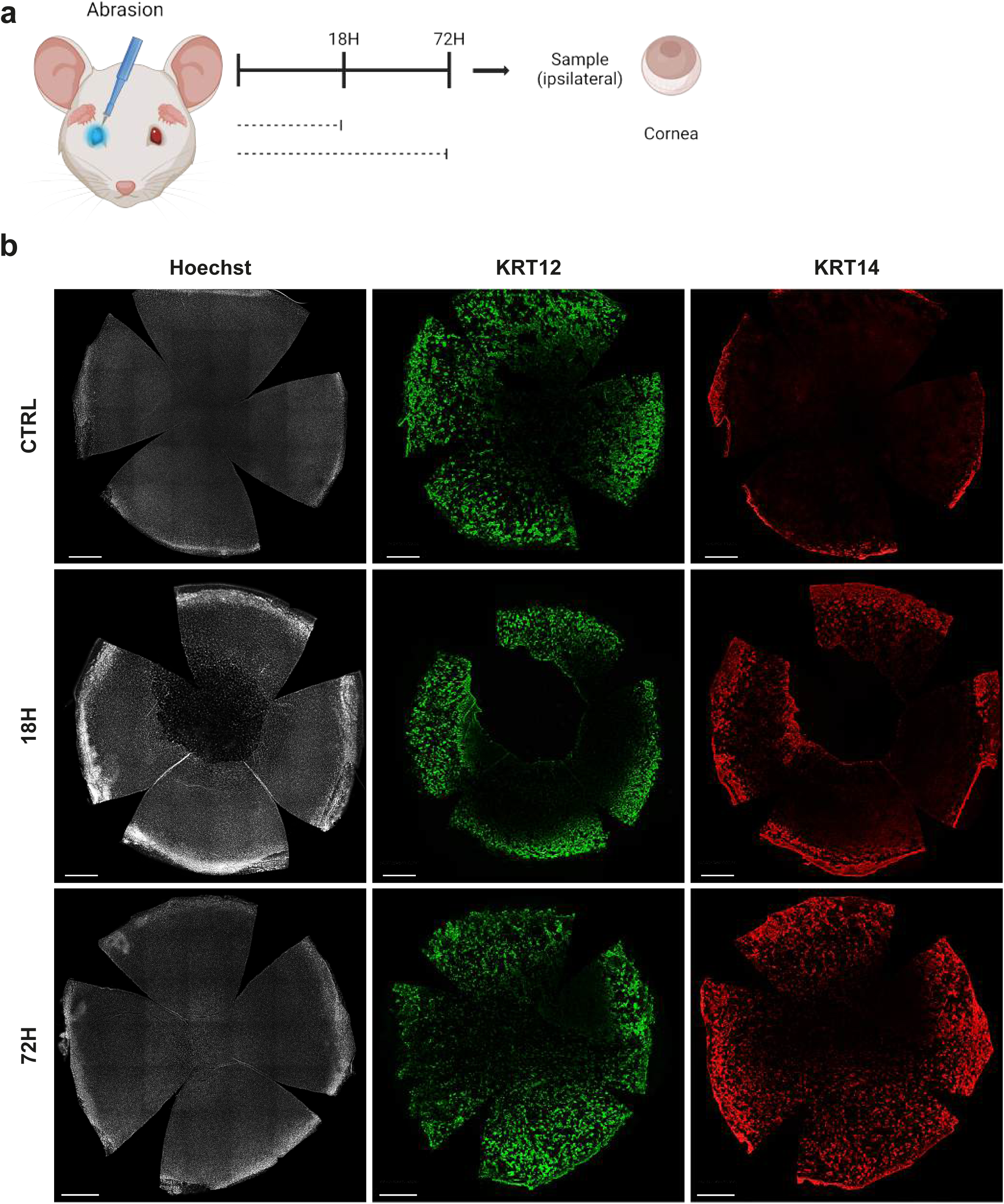
Corneal abrasion induces a strong KRT14 overexpression in the regenerating epithelium. (a) Schematic overview of the experimental design. Abrasion was performed unilaterally. Samples from the abraded side were taken before the abrasion as a control and at 18H and 72H post-abrasion. (b) Wholemount immunofluorescence staining for differentiated (KRT12) and undifferentiated (KRT14) corneal markers. Data are representative of 3 biological replicates. Nuclei were counterstained with Hoechst. Scale bars are 500µm.

Collectively, our findings demonstrate that corneal abrasion triggers significant changes in corneal intrinsic transcriptomic signature, lacrimal gland transcriptomic, and tear film proteomic profiles, supporting a dedifferentiation process in the early wound-healing phase. Additionally, the lacrimal gland and tear film also undergo considerable, though comparatively less pronounced, molecular remodeling.

### Moderate impact of tear film on corneal molecular dynamics during healing

Our next objective was to elucidate the specific role of the tear film by performing extraorbital lacrimal gland ablation alone or combined with corneal abrasion (Fig. 3a). We validated our surgery by analysing the wound closure rate, which was significantly impaired in the absence of tears, as expected (Fig. 3b). Therefore, to limit confounding effects from dry eye conditions caused by gland ablation, molecular analyses were conducted at 18 and 72 hours post-ablation. We validated our approach by analysing our candidate gene expression in lacrimal gland-ablated conditions alone. Our findings showed only Krt17 and L1CAM expressions where mildly modified solely due to lacrimal gland ablation at 18 hours post-surgery (Fig. S4a, Fig. S4b), no other of our candidate list were modulated.

**Figure 3.**
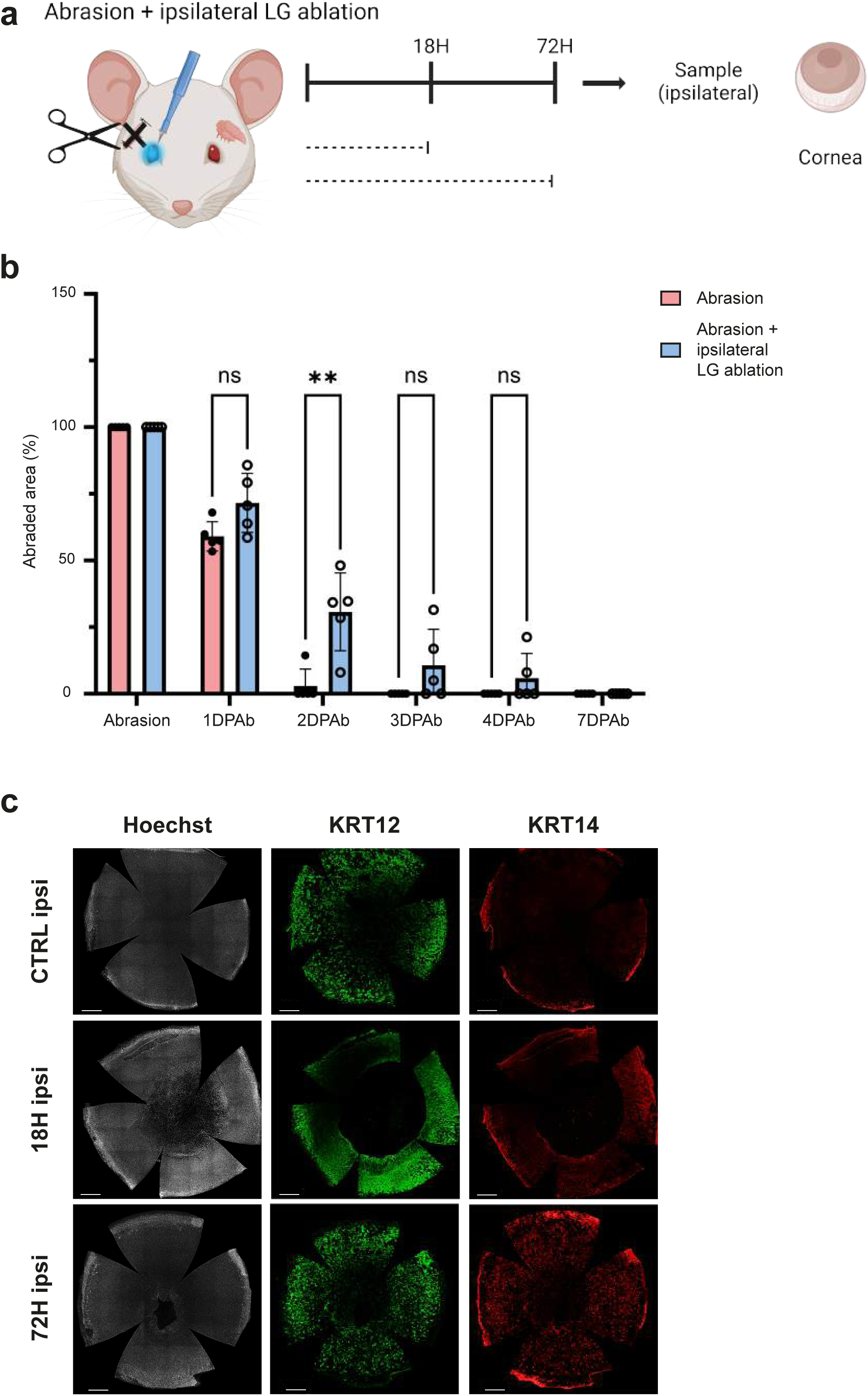
The absence of tears after a corneal abrasion delays the epithelial wound closure. (a) Schematic overview of the experimental design. Abrasion was performed unilaterally and the extraorbital lacrimal gland was removed on the ipsilateral side at the same time. Samples from the wounded side were taken before the surgery as a control and at 18H and 72H post-surgery. (b) Wound closure quantification of the abraded epithelial surface from the abrasion up to 7 days after the abrasion (DPAb) (n=5 per group). Data are represented as mean ± SD. Statistical significance was assessed by two-way repeated measures ANOVA with Geisser-Greenhouse correction and Tukey’s multiple comparisons test (**, p < 0.01). (c) Wholemount immunofluorescence staining for differentiated (KRT12) and undifferentiated (KRT14) corneal markers. Data are representative of 3 biological replicates. Nuclei were counterstained with Hoechst. Scale bars are 500µm.

When combining the lacrimal gland ablation and corneal abrasion, our 11 selected genes were significantly impacted by the abrasion, even in the absence of tear production (Fig 4a, Fig. 4b). Notably, we observed significant modulation in Wnt2, Pax6, and Mmp9. While the downregulation of Pax6 and upregulation of Mmp9 were enhanced in the absence of tears, Wnt2 expression was unexpectedly higher during wound healing without tears compared to the tear-intact condition (Fig. 4c).

**Figure 4.**
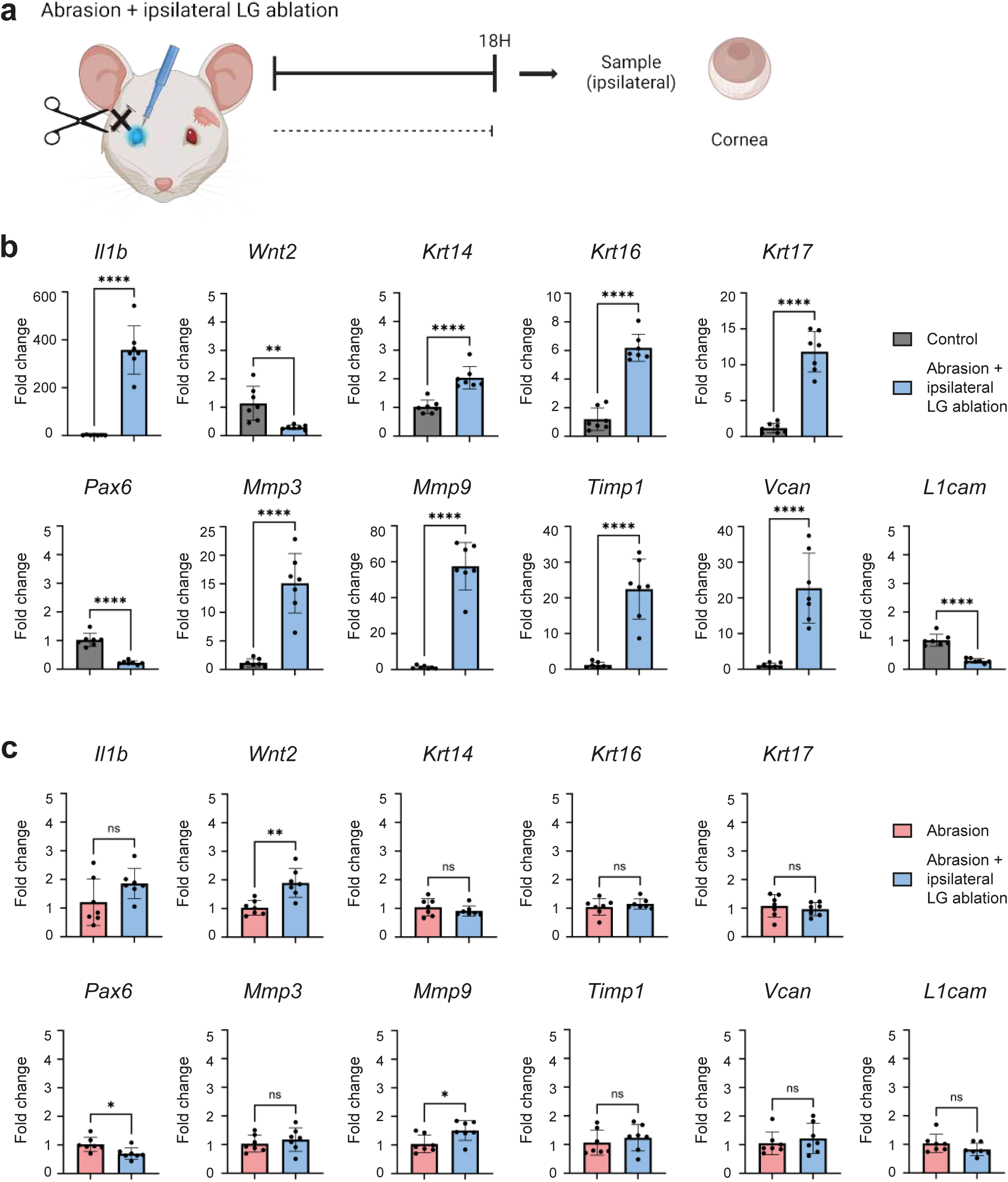
The absence of tears after a corneal abrasion mildly alters the molecular pattern of the corneal wound healing process. (a) Schematic overview of the experimental design. Abrasion was performed unilaterally and the extraorbital lacrimal gland was removed on the ipsilateral side at the same time. Samples from the wounded side were taken before the surgery as a control and at 18H post-surgery. (b) Histograms of RT-qPCR analysis on cornea samples at 18H post-surgery vs control (n=7 per group). Data are represented as mean ± SD and normalized to a reference gene (*Ppia*). Statistical significance was assessed by unpaired two-tailed t-test (*, p < 0.05; **, p < 0.01; ***, p < 0.001, ****, p < 0.0001). (c) Histograms of RT-qPCR analysis on cornea samples at 18H post-surgery vs 18H post-abrasion only (n=7 per group). Data are represented as mean ± SD and normalized to a reference gene (*Ppia*). Statistical significance was assessed by unpaired two-tailed t-test (*, p < 0.05; **, p < 0.01; ***, p < 0.001, ****, p < 0.0001).

Furthermore, we examined the epitranscriptomic profile following lacrimal gland ablation alone (Table S3). At 18 hours post-surgery, the absence of tears did not significantly affect the studied nucleoside modifications. However, by 72 hours post-surgery, four nucleoside modifications were significantly reduced. Importantly, following abrasion combined with lacrimal gland ablation, significant alterations were observed in the epitranscriptomic profile at 18 hours post-injury compared to physiological controls, affecting multiple nucleosides. When directly comparing conditions of corneal abrasion with and without tears (Table S4), only two nucleoside modifications differed significantly between the groups. Specifically, m^6^A levels were notably decreased in the absence of tears.

To confirm these molecular observations at the cellular level, immunostaining for KRT14 was performed on whole-mounted corneas (Fig. 3c). Our results indicated increased expression of KRT14 at 18 hours post-surgery, comparable to conditions with intact tear production. Furthermore, this overexpression of KRT14 expanded across the entire cornea, paralleling observations made after abrasion with an intact lacrimal gland.

Collectively, these findings underscore the critical role of the tear film and a functional corneal microenvironment for efficient wound closure. Nevertheless, our data also reveal distinct molecular changes between wound healing conditions with or without tear presence, emphasizing the nuanced role tears play in the healing process.

### Bilateral molecular response after unilateral corneal injury

Previous studies have provided evidence of a bilateral response to unilateral corneal injury. For instance, in zebrafish, a second abrasion inflicted on the contralateral eye one hour after injuring the ipsilateral eye demonstrated an accelerated wound closure^60^. However, such a bilateral molecular response has not yet been thoroughly characterized in mammalian corneal and tear film contexts. To address this gap, we investigated molecular changes occurring in the corneal microenvironment of the contralateral eye following unilateral corneal abrasion.

We first analyzed the transcriptomic profile of the cornea and the proteomic profile of tears from the contralateral eye (Fig. 5a, Fig. S5a). Transcriptomic analysis revealed early, moderate changes, with significant modulations most pronounced at 24 hours post-abrasion—occurring later than the ipsilateral response peak. Proteomic data similarly indicated early alterations with intermediate modulation of significant proteins (Fig. 5b, Fig. S5b). GO enrichment analysis showed broad similarity with ipsilateral changes (Fig. S6, Fig. S7). KEGG enrichment analysis highlighted slight but specific pathway activation at 6 hours post-injury, excluding the Notch, Hedgehog, and cell cycle pathways (Table S1, Table S2). However, activation of the Notch pathway emerged at 12 hours post-abrasion and persisted until 48 hours. Notably, significant suppression of pathways, including MAPK, TGF-β, Wnt, NF-kB and the apoptosis pathway, occurred initially at 18 hours, with JAK-STAT pathway activation most prominent between 12 and 24 hours post-abrasion.

**Figure 5.**
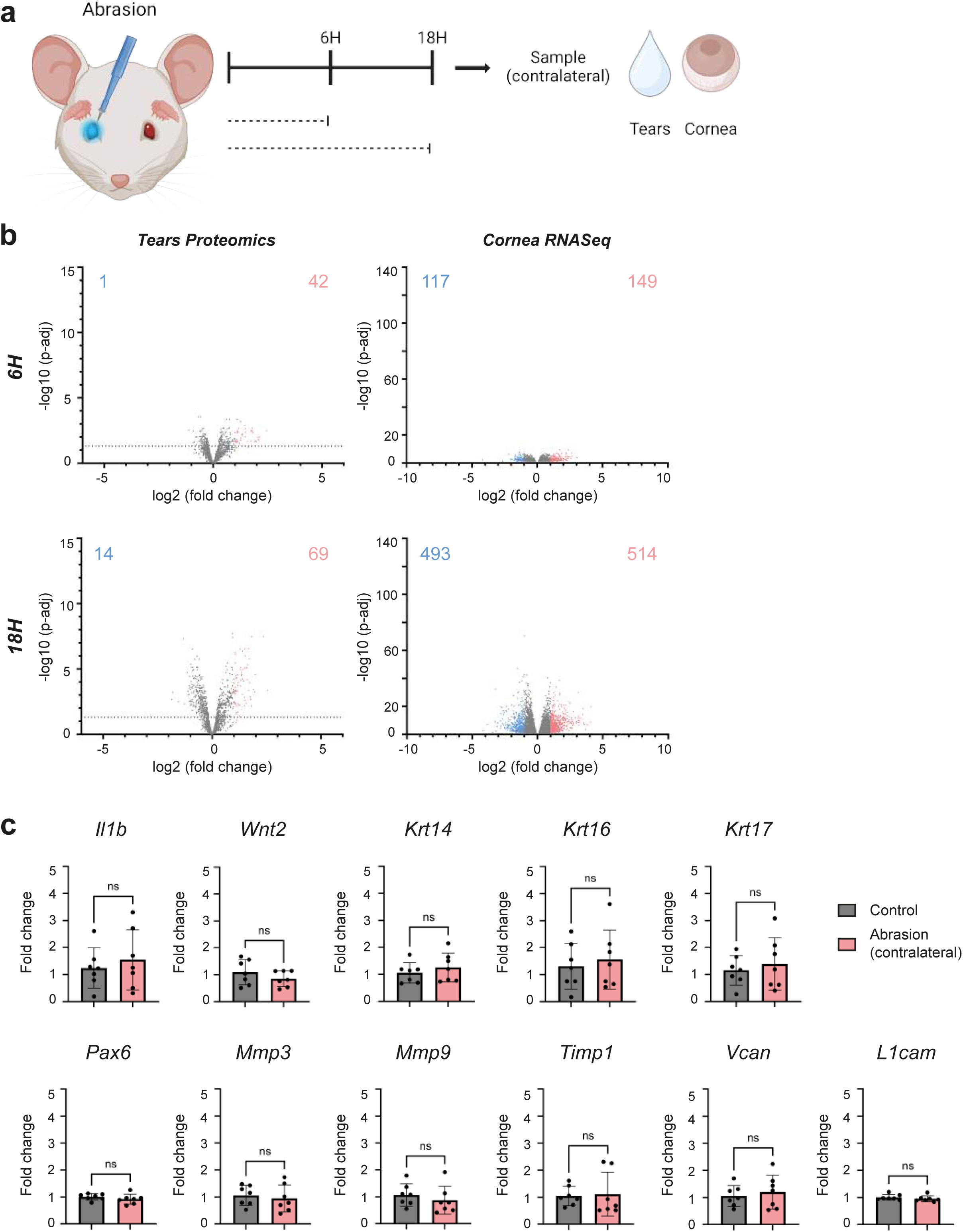
Corneal abrasion induces contralateral molecular changes in tears and cornea. (a) Schematic overview of the experimental design. Abrasion was performed unilaterally. Samples from the contralateral side of the abrasion were taken before the abrasion as a control and at 6H and 18H post-abrasion. (b) Volcano plot representations of contralateral tears proteomics analysis by mass spectrometry (n=7 per group) and contralateral cornea RNA-Seq analysis (n=5 per group) at 6H and 18H post-abrasion vs control. For RNA-Seq volcano plots, statistically significant genes are represented only. For proteomics volcano plots, all identified proteins are represented and the limit of statistical significance is shown with the horizontal dotted line. Genes and proteins with a fold change lower than 0.5 are colored in blue and their number is indicated in the top left corner of each plot. Genes and proteins with a fold change higher than 2 are colored in red and their number is indicated in the top right corner of each plot. The genes and proteins with an intermediate fold change are colored in grey. (c) Histograms of RT-qPCR analysis on contralateral cornea samples at 18H post-abrasion vs control (n=7 per group). Data are represented as mean ± SD and normalized to a reference gene (*Ppia*). Statistical significance was assessed by unpaired two-tailed t-test.

We expanded these observations through epitranscriptomic analyses of contralateral corneas, identifying no significant modifications at 18 hours post-injury (Table S3). However, at this time point, qPCR analyses demonstrated no significant bilateral alterations in our selected genes compared to controls (Fig. 5c). Cellular-level validation through immunostaining for KRT14 and KRT12 further supported this result, showing no clear bilateral increase in KRT14 or change in KRT12 domains at 18 and 72 hours post-injury, despite slight KRT14 enhancement restricted to specific regions (Fig. 6a, Fig. 6b).

**Figure 6.**
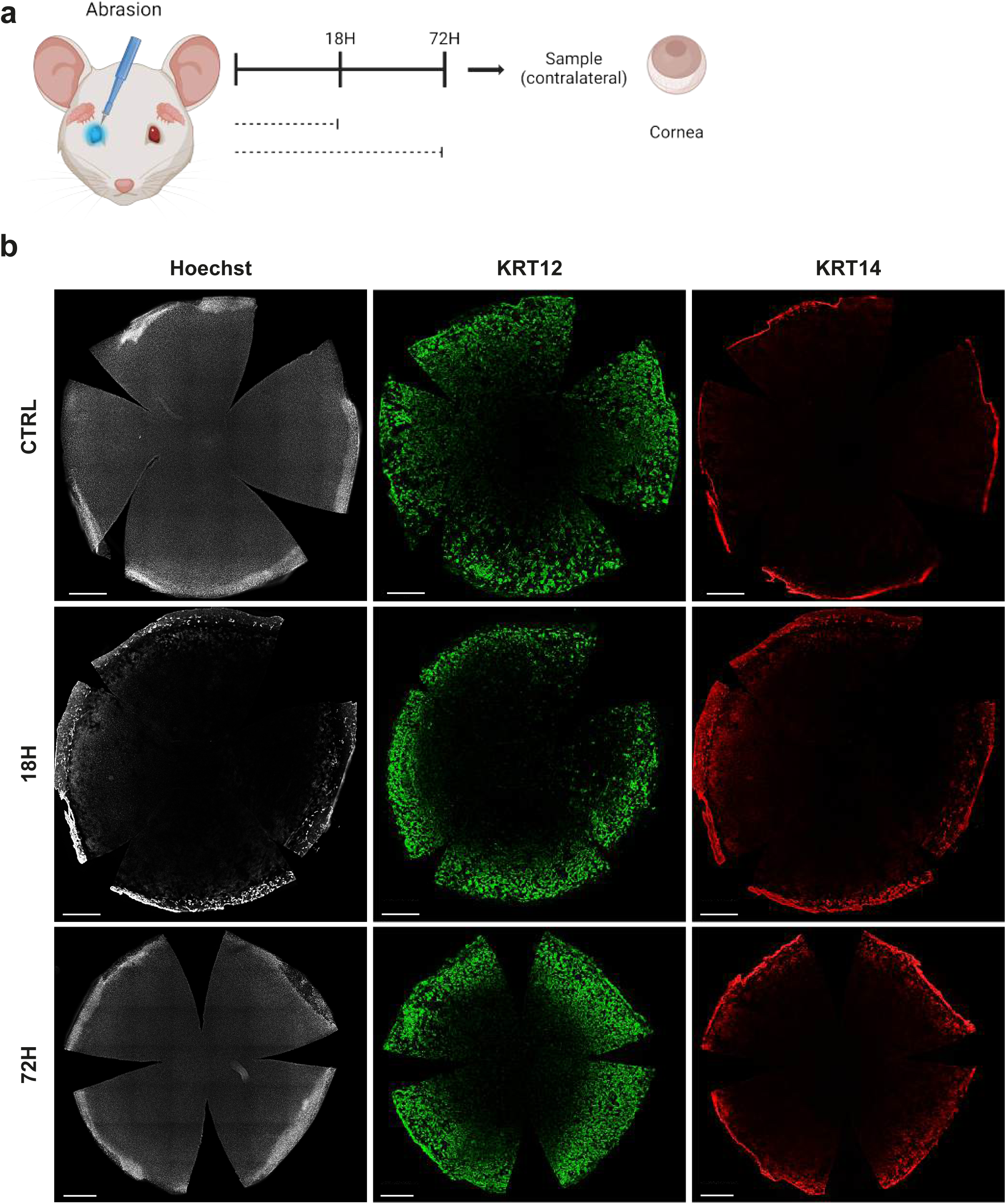
Corneal abrasion induces a mild KRT14 overexpression in the contralateral peripheral epithelium. (a) Schematic overview of the experimental design. Abrasion was performed unilaterally. Samples from the contralateral side of the abrasion were taken before the abrasion as a control and at 18H and 72H post-abrasion. (b) Wholemount immunofluorescence staining for differentiated (KRT12) and undifferentiated (KRT14) corneal markers. Data are representative of 3 biological replicates. Nuclei were counterstained with Hoechst. Scale bars are 500µm.

Finally, direct comparisons between ipsilateral and contralateral sides revealed substantial transcriptomic discrepancies, characterized by significant and sustained differences in gene expression persisting up to 48 hours post-injury (Fig. S8a, Fig. S8b), as further validated by qPCR analyses (Fig. S8c). At 18 hours post-abrasion, 26% of all DEGs showing at least a twofold increase in the wounded eye were also upregulated in the contralateral eye; however, 74% were specifically upregulated only in the wounded eye. Conversely, among DEGs exhibiting a reduction of 50% or more, 20% were commonly downregulated in both eyes, while the remaining 80% were specifically downregulated in the wounded eye (Table S5, Table S6). These commonly regulated genes represented 62% of the upregulated DEGs and 53% of the downregulated DEGs detected in the contralateral eye.

Biological processes commonly downregulated in both eyes differed notably from those exclusively downregulated in the wounded eye. However, the biological processes upregulated in both eyes were consistently associated with immune responses, as indicated by GO analysis (Fig. S9). Despite transcriptomic differences, proteomic profiles of tear fluids remained largely comparable between eyes (Fig. S10a, Fig. S10b). This observation was supported by analyzing commonly significant proteins, where 65% of the proteins upregulated and 63% of the proteins downregulated in the wounded eye showed significant changes also in the contralateral eye, representing respectively 84% and 79% of the contralaterally regulated proteins (Table S7, Table S8). In contrast, immunostaining analyses revealed pronounced and widespread KRT14 overexpression exclusively on the ipsilateral side at the cellular level (Fig. S11a, Fig. S11b).

Taken together, these results indicate a significant bilateral molecular response to unilateral corneal injury at transcriptomic and proteomic levels. However, this bilateral effect is not evident at the cellular scale, suggesting a unique and localized molecular signature associated with direct injury exposure.

### Tears constrain bilateral injury response

Finally, we sought to clarify the specific contribution of the tear film to the bilateral molecular response observed during corneal wound healing. To address this, we performed unilateral corneal abrasion combined with or without contralateral extraorbital lacrimal gland ablation and subsequently examined the expression of the previously selected genes by qPCR in the contralateral cornea at 18 hours post-surgery (Fig. 7a). Notably, Il-1β, Wnt2, Pax6, and L1cam expression levels remained unchanged in the contralateral cornea lacking tears after abrasion, compared to physiological controls (Fig. 7b). However, further comparisons between contralateral corneas with and without tear production revealed significant modulations specifically in genes associated with extracellular matrix remodeling (Mmp3, Mmp9, Timp1, Vcan) and cytoskeletal markers (Krt14 and Krt16), all showing increased expression in the absence of tears (Fig. 7c).

**Figure 7.**
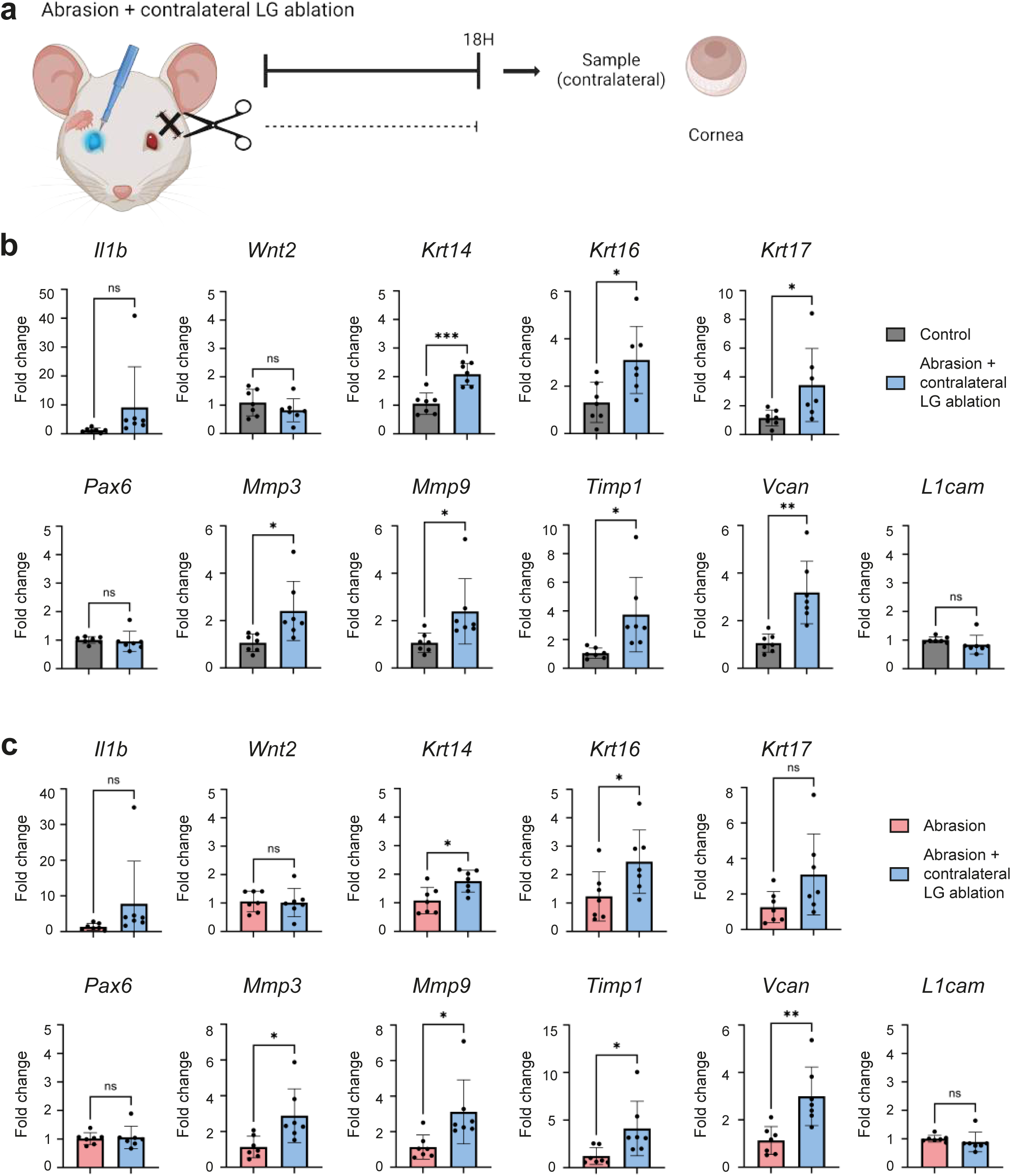
The lack of tears on the contralateral side after a corneal abrasion widely alters the molecular state of the unwounded cornea. (a) Schematic overview of the experimental design. Abrasion was performed unilaterally and the extraorbital lacrimal gland was removed on the contralateral side at the same time. Samples from the contralateral side of the abrasion were taken before the surgery as a control and at 18H post-surgery. (b) Histograms of RT-qPCR analysis on contralateral cornea samples at 18H post-surgery vs control (n=7 per group). Data are represented as mean ± SD and normalized to a reference gene (*Ppia*). Statistical significance was assessed by unpaired two-tailed t-test (*, p < 0.05; **, p < 0.01; ***, p < 0.001). (c) Histograms of RT-qPCR analysis on contralateral cornea samples at 18H post-surgery vs 18H post-abrasion only (n=7 per group). Data are represented as mean ± SD and normalized to a reference gene (*Ppia*). Statistical significance was assessed by unpaired two-tailed t-test (*, p < 0.05; **, p < 0.01).

Expanding our molecular analysis, we evaluated the epitranscriptomic profile of the contralateral corneas (Table S3). At 18 hours post-surgery, lacrimal gland ablation alone did not significantly affect the selected nucleoside modifications, except for a minor but notable reduction in specific nucleosides at later time points. When combining abrasion with gland ablation, the epitranscriptomic profile at 18 hours post-surgery indicated only modest changes, with a significant reduction limited to the m^3^C nucleoside.

We complemented these molecular findings with immunostainings for KRT14 and KRT12 on whole-mounted contralateral corneas (Fig. 8a, Fig. 8b). Despite qPCR results indicating elevated Krt14 expression, immunostaining showed only mild increases localized to peripheral cornea, without widespread bilateral alterations in KRT12 or KRT14 expression at either 18 or 72 hours post-abrasion.

**Figure 8.**
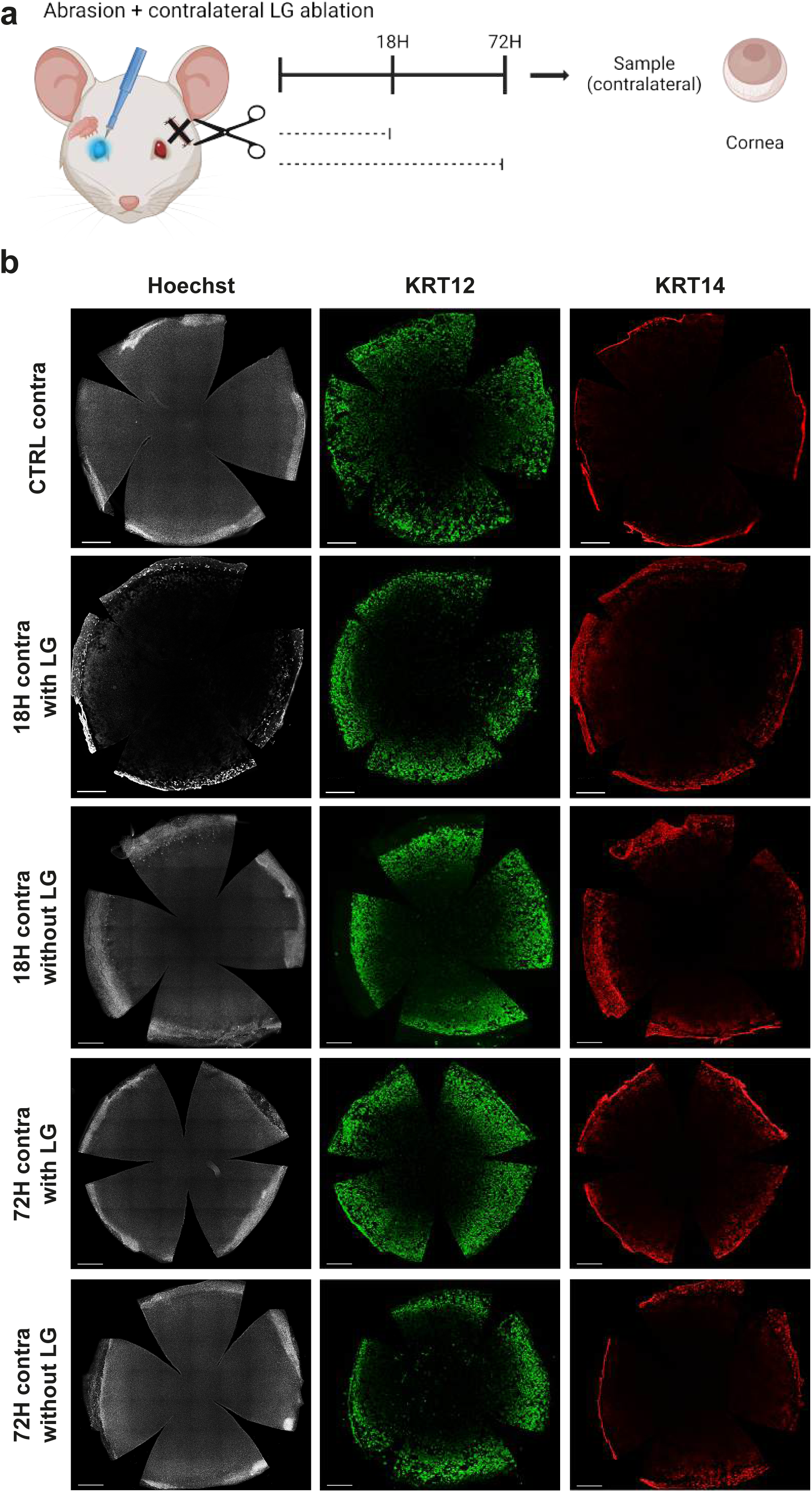
The lack of tears on the contralateral side after a corneal abrasion slightly affects KRT14 overexpression in the peripheral unwounded cornea. (a) Schematic overview of the experimental design. Abrasion was performed unilaterally and the extraorbital lacrimal gland was removed on the contralateral side at the same time. Samples from the contralateral side of the abrasion were taken before the surgery as a control and at 18H and 72H post-surgery. (b) Wholemount immunofluorescence staining for differentiated (KRT12) and undifferentiated (KRT14) corneal markers. Data are representative of 3 biological replicates. Nuclei were counterstained with Hoechst. Scale bars are 500µm.

Together, our data suggest that tear film presence exerts a selective negative regulatory role on gene expression changes in the contralateral cornea, moderating extracellular matrix remodeling and cytoskeletal rearrangements during the bilateral response to unilateral corneal injury. This regulatory mechanism likely helps prevent excessive molecular remodeling in the contralateral cornea, maintaining tissue homeostasis.

## Discussion

Corneal abrasion, characterized by a transient rupture in cellular cohesion and the integrity of the epithelial barrier, represents one of the most common threats to corneal homeostasis. Typically, this type of injury heals rapidly, within a few days, thanks to a robust wound healing process that relies heavily on the corneal microenvironment. Corneal epithelial cells contribute critical cellular factors essential for intra-tissular crosstalk, corneal innervation provides neurotrophic factors, and the tear film supplies essential growth factors. Under these conditions, corneal transparency is generally maintained. However, complications can arise, leading to clinical issues such as persistent epithelial defects or ulcerations, which are challenging to heal. This duality—from efficient healing to severe complications—highlights the need to thoroughly understand the underlying mechanisms of the wound healing process and the involvement of the entire microenvironment, an aspect frequently overlooked in prior research. This study uniquely provides a dynamic and comprehensive molecular analysis of the cornea and its microenvironment during wound healing following corneal abrasion and extraorbital lacrimal gland ablation.

Our findings demonstrate rapid and extensive changes in both corneal transcriptome and tear proteome, detectable from 6 hours and persisting up to 48 hours post-abrasion. These molecular shifts impacted numerous pathways, including TGF-β, HIF-1, and VEGF. Activation of the TGF-β pathway has previously been associated with modulating corneal epithelial migration and proliferation, keratocyte proliferation, and their transdifferentiation into myofibroblasts^61,62^. Several studies have underscored the critical roles of VEGF in nerve regeneration^63,64^ and implicated HIF-1 in corneal pathophysiology^65,66^, though its precise role remains to be clearly defined. Additionally, the downregulation of stromal components such as Mmp3, Mmp9, Timp1, and Vcan at 18 hours post-abrasion signifies significant stromal remodeling, which is essential for keratocyte activation, migration, and transdifferentiation^67,68^. Consistent with prior research, the cornea exhibited increased expression of Krt14, a marker of undifferentiated epithelial cells^58^, along with decreased expression of Pax6, which is crucial for maintaining epithelial identity. These observations strongly suggest a dedifferentiation process occurring in the early phases of wound healing.

The lacrimal gland, as part of the corneal microenvironment, produces proteins secreted into the tear film, crucial for maintaining corneal homeostasis. Although tear proteomics studies have expanded recently, particularly in dry eye syndrome^69,70^, very few have addressed tear proteomic changes during corneal wound healing *in vivo*^71^. Our study reveals significant molecular differences between healing processes in the presence or absence of tears. Notably, Pax6 downregulation was more pronounced without tears, suggesting a greater loss of epithelial identity. Additionally, Wnt2 downregulation was less marked without tears, indicating tears might enhance inhibition of the Wnt signaling pathway during healing. Wnt/β-catenin signaling contributes to corneal epithelial homeostasis by regulating proliferation and stratification^72^. Among others, Wnt2 ligand is found preferentially in the limbus, where the limbal epithelial stem cells are located, and act as a regulator of cell proliferation and cell fate^73,74^. The observed Wnt2 downregulation after abrasion suggests an activation of limbal stem cell niches, complementing epithelial dedifferentiation at wound margins, thus collectively facilitating wound closure. Indeed, we observed a significantly reduced wound closure rate in the absence of tears, emphasizing tear essential role in activating stem cell niches and regulating epithelial identity for efficient healing.

Evidence of a bilateral ocular response following unilateral injury has previously been documented in a zebrafish model^60^, yet such phenomena have not been extensively explored in mammalian systems. To address this gap, we analyzed molecular profiles in the contralateral eye following corneal abrasion. Our findings uniquely demonstrate a bilateral molecular response commencing as early as 6 hours post-abrasion and persisting up to 48 hours, detectable through corneal transcriptomic and tear proteomic analyses. However, this bilateral modulation involved distinct molecular candidates that remain to be fully characterized, as none of the specifically selected candidate genes showed contralateral expression variations. Notably, the Notch signaling pathway was activated exclusively in the contralateral eye from 12 hours post-injury and maintained through 48 hours. Prior research has associated the Notch pathway with maintaining corneal epithelial cell fate^75^, and controlling proliferation and differentiation^76,77^. Our data support a robust maintenance of epithelial cell identity in the contralateral eye during the wound healing process.

Moreover, significant differences emerged when comparing corneal transcriptomic profiles between wounded and contralateral eyes. At 18 hours post-injury, only 26% of differentially expressed genes (DEGs) showing at least a twofold increase in the wounded eye were also upregulated contralaterally. Interestingly, such transcriptomic discrepancies were less evident in tear proteomic profiles, with both eyes maintaining relatively similar molecular profiles—65% of upregulated proteins in the wounded eye were similarly modulated in the contralateral eye. Notably, when tears were absent, the contralateral side demonstrated pronounced molecular alterations, underscoring the regulatory role of tears in limiting stromal bilateral responses. Specifically, increased expression of Mmp3, Mmp9, Timp1, and Vcan was observed without tears. Consistent with our hypothesis regarding the preservation of cell fate in the contralateral eye during healing, tear absence led to increased molecular expression of Krt14; however, this was not reflected at the protein level, as KRT14 cellular localization remained unchanged.

Collectively, our findings establish a molecular bilateral response occurring early in the wound healing process, sustained for at least 48 hours post-injury. Importantly, we provide evidence that bilateralization is not driven primarily by tear film factors. Further investigation is required to clarify the underlying mechanisms. We hypothesize two potential mechanisms: firstly, bilateral responses may predominantly arise via corneal innervation and be regulated by the tear film; alternatively, systemic signals may reach the limbal niche, subsequently signaling to the broader corneal tissue. Future research exploring the specific role of corneal innervation and potential involvement of systemic factors (such as hormones or immune mediators) will be crucial to fully elucidate the nature and pathways of this bilateral phenomenon.

Our study underscores the essential role of the microenvironment in corneal wound healing, providing a detailed molecular characterization of changes occurring in both injured and contralateral corneas and tear fluid. This comprehensive approach highlights the necessity of an intact microenvironment for optimal wound repair, offering critical insights into the pathophysiological processes and potential therapeutic targets for improved wound healing outcomes.

Additionally, this research provides valuable insights relevant to wound healing processes occurring in other epithelial tissues, such as skin. The skin microenvironment, characterized by systemic influences, innervation, resident immune cells, and sweat glands, similarly plays a pivotal role in determining wound responses. Exploring how these diverse factors and their interactions influence skin injury responses would significantly enhance our understanding of the microenvironment’s broader role in complex and multifactorial wound healing scenarios.

Such investigations not only deepen our comprehension of fundamental pathophysiological processes but also pave the way for therapeutic strategies specifically designed to target the microenvironment, promoting quicker and more effective wound healing in various pathological conditions where the microenvironment is compromised.

In conclusion, this study emphasizes the essential role of the microenvironment in wound healing, elucidating profound transcriptomic and proteomic changes occurring within the cornea and tear fluid. For the first time, we have extensively characterized molecular events following in vivo corneal abrasion, encompassing both injured and contralateral sides, thereby establishing the crucial requirement of an intact microenvironment for efficient wound closure.

## Material and Methods

### Animals included in this study

All the experiments conducted on mice were approved by the local ethical committee and the Ministère de la Recherche et de l’enseignement Supérieur (authorization 2016080510211993 version2). All of the procedures were carried out in accordance with the French regulation for the animal procedure (French decree 2013-118) and with specific European Union guidelines for the protection of animal welfare (Directive 2010/63/EU). Mice were housed in plastic boxes, on a standard light cycle (12h light, 12h dark), with food and water ad libitum, in a 40-60% relative humidity environment and a 21°C-22°C ambient temperature. Swiss/CD1 female mice (RjOrl:SWISS, Janvier Labs, France) of 11 to 12 weeks of age were used for all experiments.

### Corneal abrasions and wound healing monitoring

Corneal abrasions were performed as previously described^34,78^. Briefly, mice were anesthetized by intraperitoneal injection with a mix of ketamine (90mg/kg, Imalgene® 1000, Centravet) and medetomidine (1mg/kg, Domitor®, Centravet). The epithelium was abraded unilaterally on one eye with an ocular burr (Algerbrush II, reference BR2-5 0.5 mm, Alger company, USA). The abrasion was immediately checked by using a fluorescein solution (1% in PBS, Sigma-Aldrich) under cobalt blue light and a picture was taken. After abrasion, a drop of Ocrygel (TVM) was applied on both eyes, analgesia was given using a buprenorphine (0.1mg/kg, Burprecare®, Centravet) analgesic solution and mice were woken up with atipamezole (1mg/kg, Antisedan®, Centravet). The wellbeing of the animals was monitored in the following days. For wound healing monitoring, the eye surface was specifically checked at 1, 2, 3, 4 and 7 days after the abrasion using fluorescein solution on conscious hand-restrained mice and a picture was taken each time until complete wound closure.

### Lacrimal gland ablation

Mice were anesthetized by intraperitoneal injection with a mix of ketamine (90mg/kg) and medetomidine (1mg/kg). A drop of Ocrygel was applied on both eyes. The mouse was put on its side, the fur under the ear was trimmed carefully, disinfected with 70% ethanol/PBS solution followed by vetedine solution (Vétoquinol). A skin incision was made above the lacrimal gland location, the extraobital lacrimal gland was removed by carefully cutting the duct and the surrounding connective tissues. The incision was sutured with two individual stitches using 6-0 absorbable suture thread (Vicryl® Polyglactin 910, reference W9981, Ethicon). For specific cohorts, a corneal abrasion was performed, as described above, after lacrimal gland ablation. The sutured skin was disinfected with vetedine solution, analgesia was given using a buprenorphine (0.1mg/kg) analgesic solution and mice were woken up with atipamezole (1mg/kg). The wellbeing of the animals was monitored in the following days.

### Sample collection and processing

Mice were euthanazied by cervical disclocation. The eyes were collected by enucleation, using curved scissors to cut the optic nerve, and washed briefly in PBS. For RNA-Seq, qPCR and epitranscriptomics analyses, the corneas were immediately dissected in an RNAse free environment, snap frozen in liquid nitrogen and then stored at −80°C before the analyses. For RNA-Seq, two corneas were pooled to form each sample, except for three samples at 12H post-abrasion that needed five corneas per sample. For qPCR, five corneas were pooled to form each sample. For epitranscriptomics, three corneas were pooled to form each sample. For each mouse, both eyes were dissected individually, to ensure ipsilateral and contralateral analyses later on. Otherwise, for immunofluorescence labeling, the eyes were fixed in a 4% paraformaldehyde solution (Antigenfix) for 20 min and washed 3 times with PBS for 15 min each. Eyes were then dehydrated for 2H in 50% ethanol/PBS and stored in 70% ethanol/PBS at 4°C. The extraorbital lacrimal glands were collected by cutting the skin above the lacrimal gland location and carefully cutting the duct and the surrounding connective tissues. The glands were snap frozen in liquid nitrogen and stored at −80°C before the analyses. Lacrimal gland and cornea samples were taken from different mice for the different time points of the experiments.

Tear samples were collected from conscious hand-restrained mice by using 1µL disposable microcapillary glass tubes (reference 022.7726, CAMAG). Briefly, the mouse was restrained by hand and a microcapillary glass tube was gently tapped on the outer corner of the eye, avoiding contact with the eye itself. Both eyes were sampled for 1 min each (with approximatively 1 tap per second) and stored separately for each mouse in 12µL of ABC (Ammonium Bicarbonate) buffer, provided by the proteomics platform. The samples were temporarily kept on ice while the tear collection was performed and then stored at −80°C before the analyses. For each experiment, tears were sampled from the same mice for all the time points as a longitudinal follow-up.

### RNA-Seq on cornea and lacrimal glands

The same protocol was performed for both cornea and lacrimal gland samples. TRIzol reagent was used to extract total RNA, which was then purified with Qiagen RNeasy kit following the manufacturer’s procedure. Total RNA concentration was quantified with Qubit. RNA ScreenTape Assay (Agilent TapeStation 4200) was used to determine RNA quality. The ribodepletion of 500ng of good quality total RNA (RIN > 8.8) was achieved using Illumina’s mammalian Ribo-Zero Magnetic Gold Kit HMN/Mouse/Rat (reference MRZG12324, Illumina). Ribodepleted RNA was then used to complete the library preparation with the NEBNext Ultra Directional RNA Library prep kit (reference E7420L, New England Biolabs) using 8 cycles of PCR amplification, followed by single (i7) indexing. Indexed library preparations from each sample were pooled and the NextSeq 500 using a NextSeq High Output 75 cycle flow cell (Illumina) was used for sequencing with 75SE reads at a pool concentration of 1.4 pM.

### RT-qPCR on cornea

Samples composed of five pooled corneas were used for the extraction of total RNA. CKMix lysing kit and Precellys homogenizer (Bertin Technologies) were used to perform tissue grinding with 1mL of TRIzol reagent (Life Technologies). RNA purification and DNase treatment were then achieved with the Nucleospin RNA Kit (Macherey Nagel). Quantification of the RNA samples was performed by using the NanodropOne spectrophotometer (ThermoFisher Scientific). The integrity of the RNAs was then checked using the Agilent 2100 bioanalyzer with the RNA 6000 Nano kit (Agilent Technologies). The High-Capacity Reverse Transcription kit (Applied Biosystems) was used following the manufacturer’s instructions to reverse transcribe 500ng of total RNA in a 20µL final reaction volume, using random primers and the presence of RNase inhibitor. The quantitative PCR experiments were conducted using a TaqMan detection protocol with the QuantStudio 12 K Flex Real-Time PCR System (Life Technologies). The TaqMan assays (ThermoFisher Scientific) selected for each target and reference gene are presented in ***Table 1***. For qPCR reactions, after mixing 3ng of cDNA with TaqMan FastAdvance Master Mix and TaqMan assay in a final volume of 10µL, samples were loaded on 384-well microplates and 40 cycles of PCR were performed (50°C/2min; (95°C/1s; 60°C/20s) X40). To check the absence of genomic DNA contaminants, negative controls without the reverse transcriptase were implemented. Normalization of the data was achieved by adding three reference genes in the experiments (*Hprt1*, *Ppia*, *Ubc*) and using the GenEx software (MultiD) to select the most stable one (*Ppia*). Duplicates were made for each measurement, and the determined Ct values were used for analysis. The relative gene expression ratio was determined using the ΔΔCt method.

**Table 1.** List of the genes and their identification used for RT-qPCR analysis.

| **Gene** | **Probe ID** |
| --- | --- |
| *Hprt1* | Mm00446968_m1 |
| *Ppia* | Mm02342430_g1 |
| *Ubc* | Mm01198158_m1 |
| *Il1b* | Mm00434228_m1 |
| *Wnt2* | Mm00470018_m1 |
| *Krt14* | Mm00516876_m1 |
| *Krt16* | Mm01306668_g1 |
| *Krt17* | Mm00495207_m1 |
| *Pax6* | Mm00443081_m1 |
| *Mmp3* | Mm00440295_m1 |
| *Mmp9* | Mm00442991_m1 |
| *Timp1* | Mm01341361_m1 |
| *Vcan* | Mm01283063_m1 |
| *L1cam* | Mm00493049_m1 |

### Proteomics study

The proteomic profile of tear samples was studied. The total protein concentration was quantified using the NanodropOne Microvolume UV-Vis Spectrophotometer (reference ND-ONE-W, ThermoFisher Scientific). 3µg of total protein were reduced, alkylated and digested with trypsin using automated SP3 protocol on the Bravo 96LT AssayMap Protein Sample Prep Platform (Agilent Technologies). Peptides were desalted and injected in triplicate on the system Evosep one/timsTOF HT (Bruker Daltonics). Protein identification was performed with DIA-NN software (version 1.8.1). The parameters used were the following: the digestion enzyme is trypsin, the number of missed cleavages was 1, the minimum peptide size was 7 amino acids, the precursor charge range was set from 1 to 4, the precursor m/z set from 300 to 1800 and a protein identification FDR was set at 1%. The Uniprot database of the mouse proteome was used as reference (Release_2024_03). Some modifications, induced by the sample preparation protocol, were studied: asparagine deamidation and methionine oxidation as variable modifications and cysteine carbamidomethylation as fixed modification. The maximum number of variable modifications was set to 2. LFQ intensities obtained were processed using the Perseus software (version 1.6.15.0) and DIA-Analyst platform (version 0.8.6). The mean intensity of the three technical replicates was calculated and used for statistical analysis. The LFQ data for each protein were transformed by applying the log2(x) formula and grouped according to condition (CTRL, T6, T12, T18 and T24) for each eye (left and right). For biological replicates, proteins with 100% of valid values were selected. A cutoff of the adjusted p-value of 0.05 (t-statistic correction) along with a log2(fold change) of 1 has been applied to determine significantly regulated proteins in each pairwise comparison for paired samples.

### ImmunoHistoChemistry (IHC) studies

The following antibodies were used for IHC studies: rabbit anti-KRT12 (1:200, reference MA5-42701, Invitrogen) and chicken anti-KRT14 (1:500, reference 906004, BioLegend) as primary antibodies, goat anti-Rabbit IgG H&L Alexa Fluor 488 (1:500, reference A11008, Fisher Scientific) and goat anti-chicken IgY H&L Alexa Fluor 568 (1:500, reference ab175711, Abcam) as secondary antibodies respectively. Nuclei were counterstained with Hoechst 33342 (1:5000, reference H3570, Thermofisher Scientific).

### Immunofluorescence labeling on whole cornea

To study the expression of KRT12 and KRT14 in the corneal epithelium, eyes were rehydrated for 2H in 50% ethanol/PBS and washed two times in PBS for 10 min each at room temperature. After dissection, corneas were blocked and permeabilized with a 5% Goat Serum (GS) (reference 16210064, Thermo Fisher Scientific) 5% Fish Skin Gelatin (FSG) (reference G7765, Sigma-Aldrich) in 0.5% Triton X-100/PBS solution, with agitation for 1H at room temperature. Corneas were incubated in primary antibody diluted in blocking solution (5% GS 5% FSG in 0,1% Triton X-100/PBS) overnight at 4°C with agitation and rinsed in 0,1% Triton X-100/PBS at room temperature 3 times for 1H each. Next, samples were incubated in secondary antibody diluted in blocking solution overnight at 4°C on agitation and rinsed 3 times for 1H each in 0,1% Triton X-100/PBS at room temperature. After the washes, nuclei were stained for 10 min with Hoechst 33342 and washed 5 min in PBS. Corneas were then cut at four cardinal points with a carbon steel surgical blade (15C, reference 0221, Swann-Morton) and flat mounted in Vectashield medium (Vector laboratories, H-1000), with the epithelium facing the coverslip.

### Images acquisition and processing

The acquisition of the images was performed as previously described^79^. A Leica Thunder Imager Tissue microscope was used to acquire the whole-cornea images, using the navigator module with the large volume computational clearing (LVCC) process. The LAS X software (version 3.7.4) was used to obtain the images using a 20X/0.55 objective. Imaris Bitplane software (version 9.8.0) was used to process the images. All of the images from a single panel were acquired and processed with the same parameters. For wound healing monitoring, the Fiji measurement plug-in (FIJI (RRID:SCR_002285)) was used to determine the size of the wounded area, highlighted with the fluorescein staining.

### Gene Ontology and Kyoto Encyclopedia of Genes and Genomes enrichment analyses

To analyse the Gene Ontology (GO) biological processes from the RNA-Seq data on corneas, statistical analyses were performed using R and RStudio (version 4.4.1). The *BiocManager* package from the Bioconductor open source software project was used (*clusterProfiler* and *AnnotationDbi* packages). Genes with a p-adj <0.05, baseMean >50 and log2(FoldChange) >1 (for upregulated genes) or log2(FoldChange) <1 (for downregulated genes) were chosen. Genes were then associated with biological processes (BP) using gene ontology (GO) enrichment analyses, with the *Mus musculus* reference genome as the background dataset (org.Mm.eg.db). Results are presented as dot plots and show the top 15 upregulated or downregulated pathways.

To analyse the activation or inhibition of various pathways after abrasion, 17 pathways were taken from the Kyoto Encyclopedia of Genes and Genomes (KEGG) pathway database and analysed for the Mus musculus (house mouse) organism. The chosen process pathways were the following: focal adhesion (mmu04510), chemokine (mmu04062), calcium (mmu04020), neurotrophin (mmu04722), apoptosis (mmu04210), cell cycle (mmu04110) and stem cells pluripotency (mmu04550). The studied molecular pathways were the following: MAPK (mmu04010), PI3K-Akt (mmu04151), TGF-β (mmu04350), Notch (mmu04330), Wnt (mmu04310), Hedgehog (mmu04340), JAK-STAT (mmu04630), NF-kB (mmu04064), HIF-1 (mmu04066) and VEGF (mmu04370). The list of all genes included in each pathway was extracted (“background genes”). From the RNA-Seq data on corneas, differentially expressed genes with a statistical significance (p-adj <0.05) and log2(FoldChange) >1 (upregulated genes) or <1 (dowregulated genes) were isolated. From the RNA-Seq data on lacrimal gland, differentially expressed genes with a statistical significance (p-adj <0.05) were isolated, without a log2(FoldChange) restriction. Next, the presence of each gene was verified in the list of background genes for each pathway. For all present genes in the pathways, the log2(FoldChange) was taken and genes were sorted out between the upregulated ones, with a positive log2(FoldChange), and the downregulated ones, with a negative log2(FoldChange). The upregulated genes were searched by name for each pathway with the “Search” option. The total number of actors in the pathway was counted (green boxes), and the number of actors found after searching was noted (pink boxes). The ratio between pink boxes and green boxes was calculated for each pathway and converted in a percentage. The same process was done for the downregulated genes.

### Statistical analysis

Data were analysed using GraphPad Prism software (version 10.1.2, Prism, CA, USA) and expressed as the mean ± SD as indicated in the Figure legend. Statistical differences were tested using unpaired two-tailed t-test, or two-way repeated measures ANOVA with Geisser-Greenhouse correction followed by Tukey’s multiple comparisons test, as indicated in the Figure legend. Significant p-values were represented as *p < 0.05, ** p < 0.01, *** p < 0.001, **** p < 0.0001.

## Data availability

The raw data is available upon request and the datasets produced for the current study are publicly available. The RNA-Sequencing data is available in the GEO repository. The mass spectrometry proteomics data is available in the PRIDE repository.

## Author contributions

Conceptualization: N.F., F.M.; Methodology: N.F., F.M.; Validation: N.F., F.M.; Formal analysis: N.F., F.M.; Data analysis: N.F., F.M.; Writing: N.F., F.M.; Supervision: F.M.; Project administration: F.M.; Funding acquisition: F.M.

## Acknowledgments

This research was supported by ATIP-Avenir program, Inserm, the Région Occitanie, ANR (ANR-21-CE17-0061, TeFiCoPa), FRM (REP202110014140), Support for research: I-SITE 2024 - program of excellence of the University of Montpellier, CBS2 Doctoral School, and the Fondation Groupama. The RNA-Sequencing service was provided by the Biomedicum Functional Genomics Unit at the Helsinki Institute of Life Science and Biocenter Finland at the University of Helsinki. The RT-qPCR service was provided by the Dr Eric Jacquet at the QPCR plateform from Réseau des plateformes de Génomique Paris-Saclay at the Institut de Chimie des Substances Naturelles at Gif-sur-Yvette. Mass spectrometry experiments were carried out using the facilities of the Montpellier Proteomics Platform (PPM-PPC, BioCampus Montpellier), member of the Proteomics French Infrastructure (ProFI). We thank the MRI-DBS imaging facility, member of the France-BioImaging national infrastructure supported by the French National Research Agency (ANR-10-INBS-04, «Investments for the future»). We thank the personel of the INM animal core facility, member of the Animal Facility Network in Montpellier (RAM).

## Supplementary Material and Methods

### Epitranscriptomics study

Samples containing three pooled corneas were used for the extraction of total RNA. For each sample, 250µL of PBS was added in 2mL LoBind® tubes (Eppendorf) and tissue grinding was performed manually with a mechanical grinder (FastPrep-24™ Classic, MP Biomedicals) (20sec X7). 100µL of material was then transferred in 1.5mL LoBind® tubes. 1mL of TRIzol reagent (reference T9424, Sigma-Aldrich) and 200µL of chloroform (reference C2432, Sigma-Aldrich) were added in each tube and a manual agitation was performed vigorously for 15sec. Samples were incubated for 5 min at room temperature and then centrifugated for 15 min at 12000g at 4°C. The aqueous phase of each sample was collected (400µL) and transferred in new LoBind® tubes. 500µL of isopropanol (reference 0016264101BS, Biosolve) was added and samples were mixed by hand before being incubated for 10 min at room temperature. A centrifugation was performed for 30 min at 12000g at 4°C and the supernatant was discarded, keeping the pellet containing the RNAs. 1mL of 75% ethanol diluted in ULC/MS grade water (reference 0023214102BS, Biosolve) was added, samples were vortexed to detach the pellet and then centrifugated for 10 min at 7500g at 4°C. The supernatant was discarded and the tubes were left open on clean paper towel at room temperature to let the RNAs dry completely (1H minimum). All samples were resuspended in 20µL of Milli-Q® water (Merck Millipore) and incubated for 10 min at 55°C. Quantification of the RNA samples was then performed (ND-ONE-W, ThermoFisher Scientific). 200ng of total RNA was transferred in new LoBind® tubes in a 20µL final volume completed with Milli-Q® water. 3µL of 100mM pH 5.3 ammonium acetate and 1µL of nuclease P1 (1U/mL) were added. Samples were incubated for 2H at 42°C before adding 3µL of 1M pH 7 ammonium acetate and 1µL of alkaline phosphatase from *E. coli* (1U/mL). Samples were incubated for 2H at 37°C and 60µL of phase A (5mM pH 5.3 ammonium acetate) was added. Filtration of the samples was achieved using 0.22µm filters (reference SLGVR04NL, Millipore) and the filtrate was transferred in vials (reference 5188-2788, Agilent). For nucleoside analysis, 5µL of sample were injected into a Nexera LC-40 system coupled to a Triple Quadrupole NX-8060 (Shimadzu).

### LC method

The chromatographic separation of nucleosides was performed on a polar end-capped C18 column (Synergi Hydro-RP, 4µm particle size, 250mm x 2.0mm, 80Å, reference 00G-4375-B0, Phenomenex). The flow rate and temperature were set at 0.4 mL/min and 35 °C, respectively, and a gradient elution was used, with 0.5% Acetic Acid (reference 0001074131BS, Biosolve) in ULC/MS water as mobile phase A and pure acetonitrile (reference 0001204101BS, Biosolve) as phase B. The total run time was 35 min and the gradient elution program started with 100% phase A for 3 min, followed by an increase in phase B up to 8% for 10 min, and continued for an additional 10 min up to 40% solvent B. At 23 min, the column was rinsed with 80% phase B for 7 min, and the initial conditions were regenerated by flushing the column with 100% phase A for another 5 min.

### MRM method

For the detection of nucleosides, a triple quadrupole mass spectrometry (Shimadzu NX-8060 Triple Quadrupole) equipped with an electrospray ionization (ESI) source was used for MRM analysis in the positive ion mode. ESI source parameters were set as described in Table S9, and multiple reaction monitoring (MRM) mode was used for the detection of 51 targets based on their fragmentation pattern. Instrument settings for the analysis were set individually for each nucleoside (Table S10) and, for each transition, the retention time window was set to 3 min and the dwell time to 20ms.

The total area under curve (AUC) for each unmodified and modified nucleoside was extracted using the LabSolution Insight software (version 3.8 SP3, Shimadzu). The ratio of the total AUC of each modified nucleoside relative to the total AUC of Uridine was calculated. The list of all modified nucleosides detected by LCMS in this experiment is shown in Table S11. The Modomics database was used to gather information regarding the RNA type in which is found each modified nucleoside.

## Supplementary material

**Figure S1.**
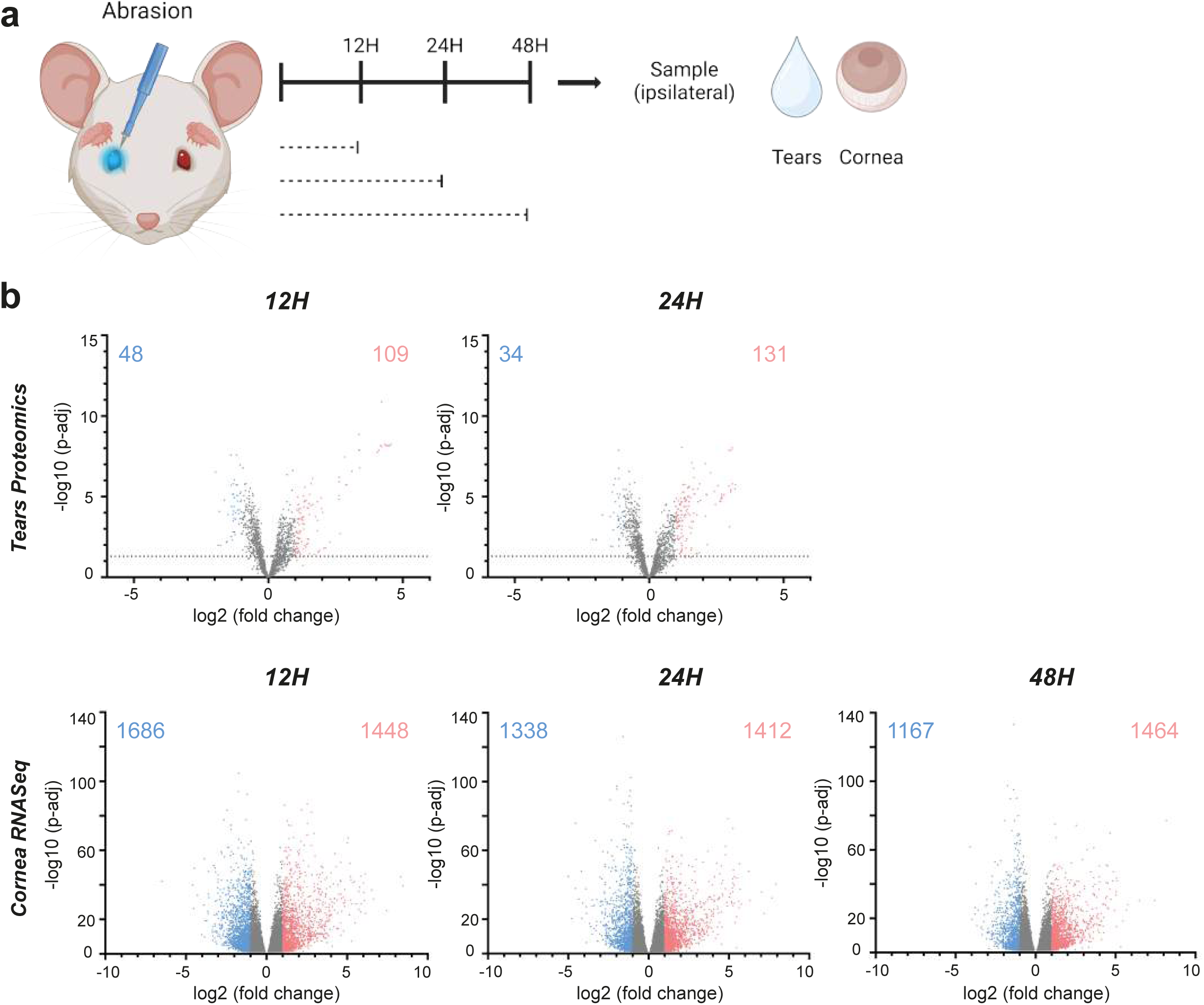
Corneal abrasion induces molecular changes in tears and cornea. (a) Schematic overview of the experimental design. Abrasion was performed unilaterally. Samples from the abraded side were taken before the abrasion as a control and at 12H, 24H and 48H post-abrasion. (b) Volcano plot representations of tears proteomics analysis by mass spectrometry (n=7 per group) and cornea RNA-Seq analysis (n=5 per group) at 12H, 24H and 48H post-abrasion vs control. For RNA-Seq volcano plots, statistically significant genes are represented only. For proteomics volcano plots, all identified proteins are represented and the limit of statistical significance is shown with the horizontal dotted line. Genes and proteins with a fold change lower than 0.5 are colored in blue and their number is indicated in the top left corner of each plot. Genes and proteins with a fold change higher than 2 are colored in red and their number is indicated in the top right corner of each plot. The genes and proteins with an intermediate fold change are colored in grey.

**Figure S2.**
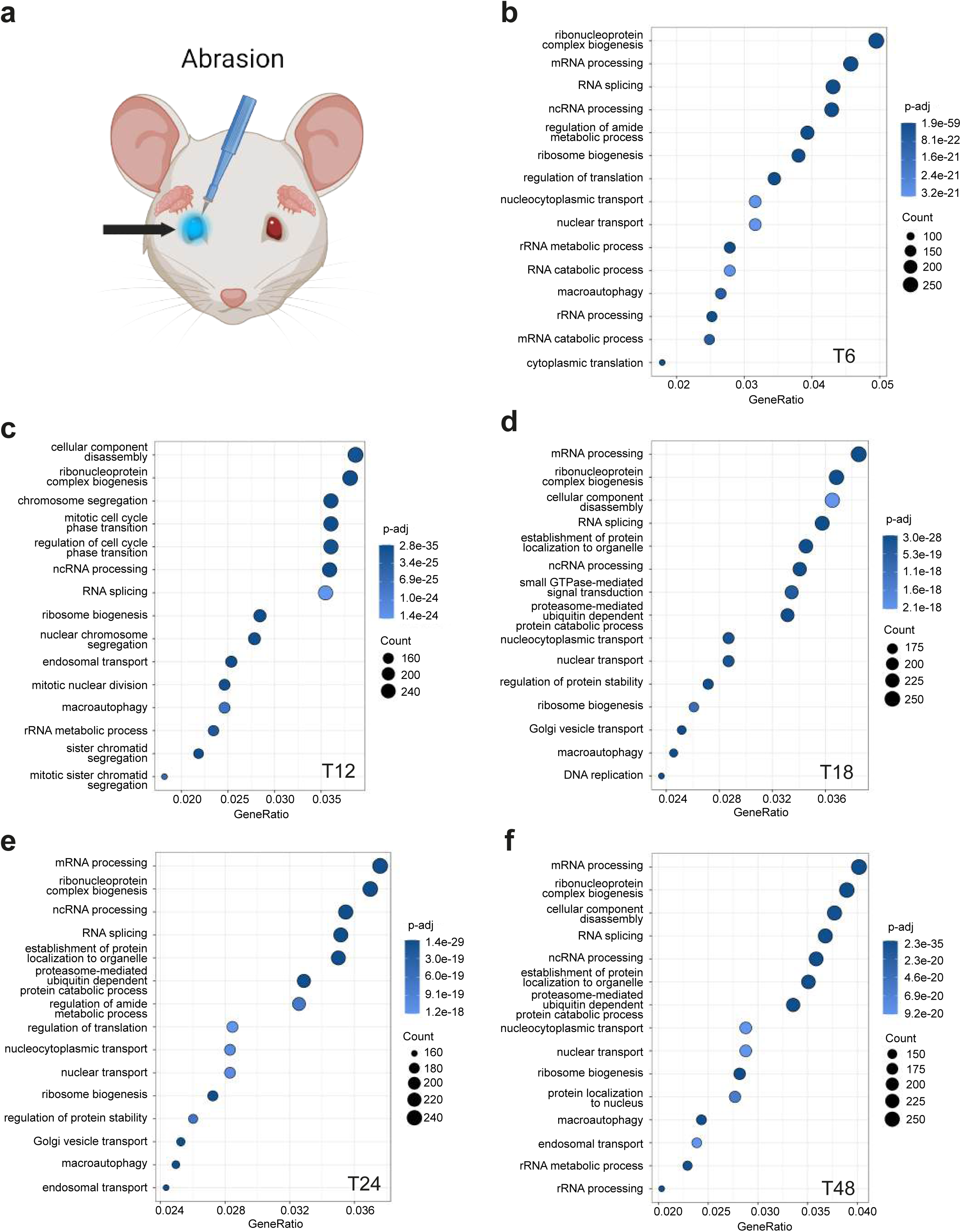
Corneal abrasion induces the downregulation of various biological processes revolving around RNA and protein processing. (a) Schematic overview of the experimental design. Dot plot representations of Gene Ontology (GO) enrichment analysis performed in R from RNA-Seq data on the wounded cornea. (b, c, d, e, f) The top 15 downregulated biological processes are shown for each plot, from 6H to 48H post-abrasion. GeneRatio represents the number of differentially expressed genes in each GO term (Count), divided by the number of genes in that GO term.

**Figure S3.**
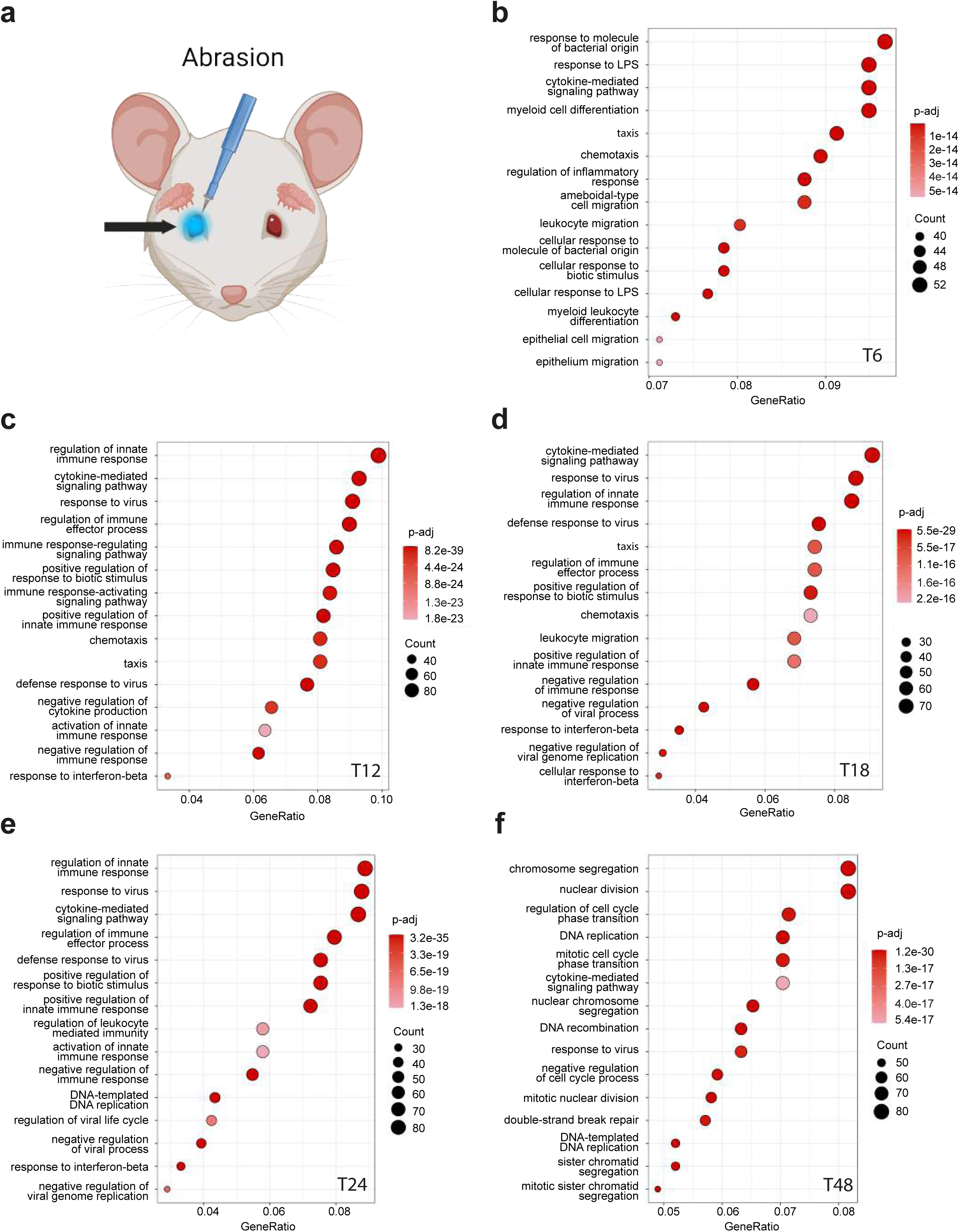
Corneal abrasion induces the upregulation of various biological processes involved in the inflammatory response. (a) Schematic overview of the experimental design. Dot plot representations of Gene Ontology (GO) enrichment analysis performed in R from RNA-Seq data on the wounded cornea. (b, c, d, e, f) The top 15 upregulated biological processes are shown for each plot, from 6H to 48H post-abrasion. GeneRatio represents the number of differentially expressed genes in each GO term (Count), divided by the number of genes in that GO term.

**Figure S4.**
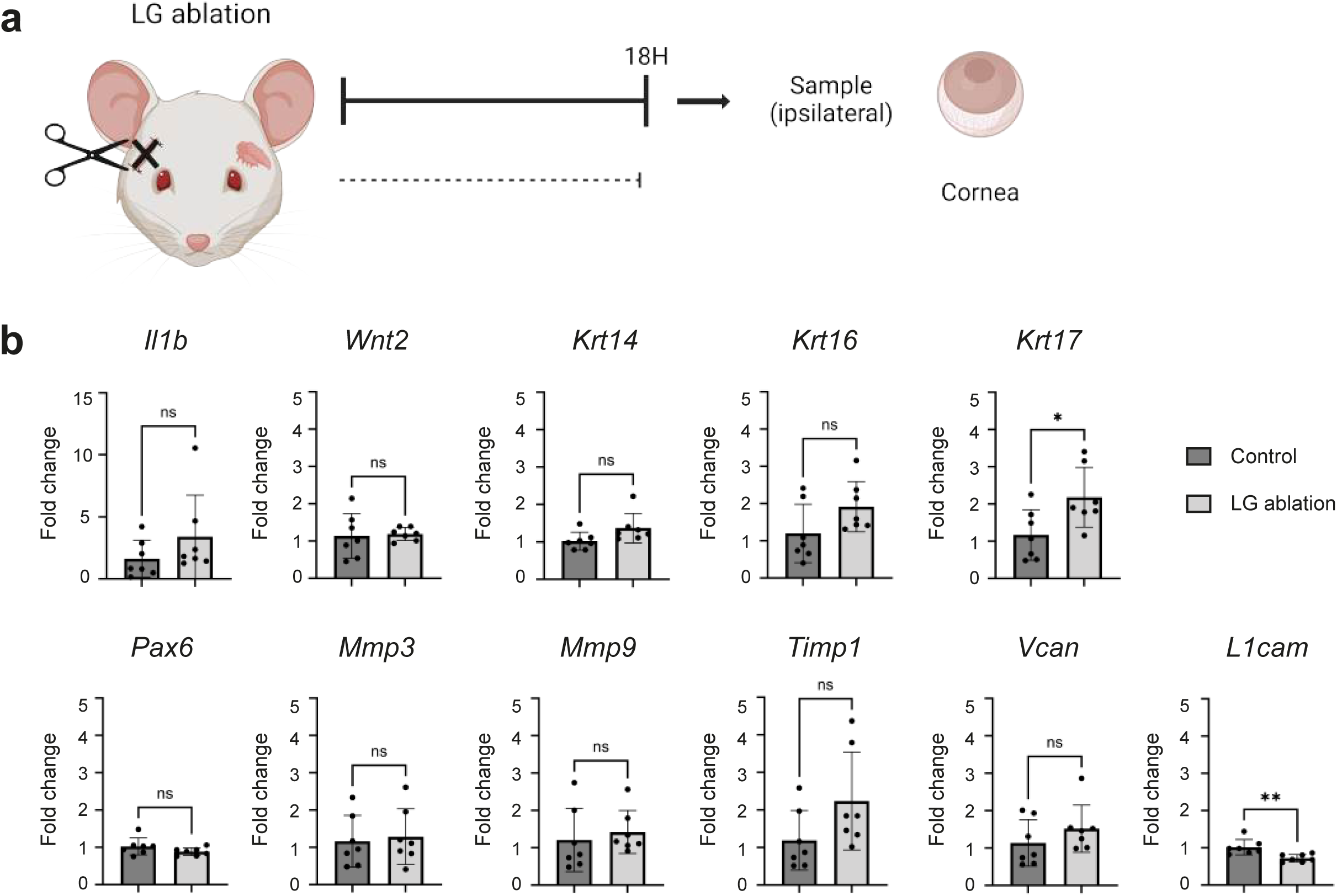
The absence of tears in an unwounded condition has a restricted impact on the molecular state of the cornea. (a) Schematic overview of the experimental design. Extraorbital lacrimal gland ablation was performed unilaterally. Samples from the ipsilateral side of the ablation were taken before the surgery as a control and at 18H post-surgery. (b) Histograms of RT-qPCR analysis on cornea samples at 18H post-surgery vs control (n=7 per group). Data are represented as mean ± SD and normalized to a reference gene (*Ppia*). Statistical significance was assessed by unpaired two-tailed t-test (*, p < 0.05; **, p < 0.01).

**Figure S5.**
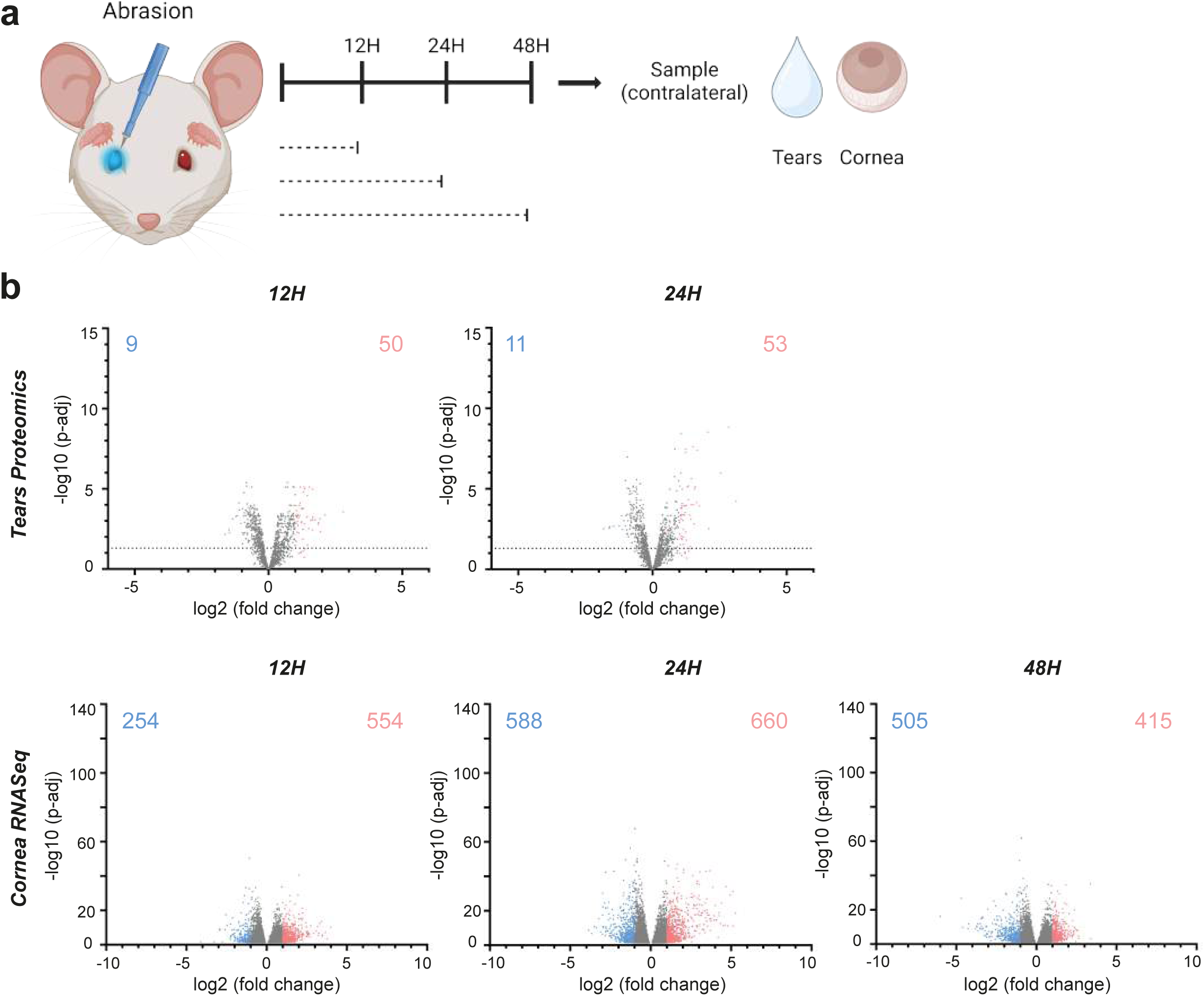
Corneal abrasion induces contralateral molecular changes in tears and cornea. (a) Schematic overview of the experimental design. Abrasion was performed unilaterally. Samples from the contralateral side of the abrasion were taken before the abrasion as a control and at 12H, 24H and 48H post-abrasion. (b) Volcano plot representations of contralateral tears proteomics analysis by mass spectrometry (n=7 per group) and contralateral cornea RNA-Seq analysis (n=5 per group) at 12H, 24H and 48H post-abrasion vs control. For RNA-Seq volcano plots, statistically significant genes are represented only. For proteomics volcano plots, all identified proteins are represented and the limit of statistical significance is shown with the horizontal dotted line. Genes and proteins with a fold change lower than 0.5 are colored in blue and their number is indicated in the top left corner of each plot. Genes and proteins with a fold change higher than 2 are colored in red and their number is indicated in the top right corner of each plot. The genes and proteins with an intermediate fold change are colored in grey.

**Figure S6.**
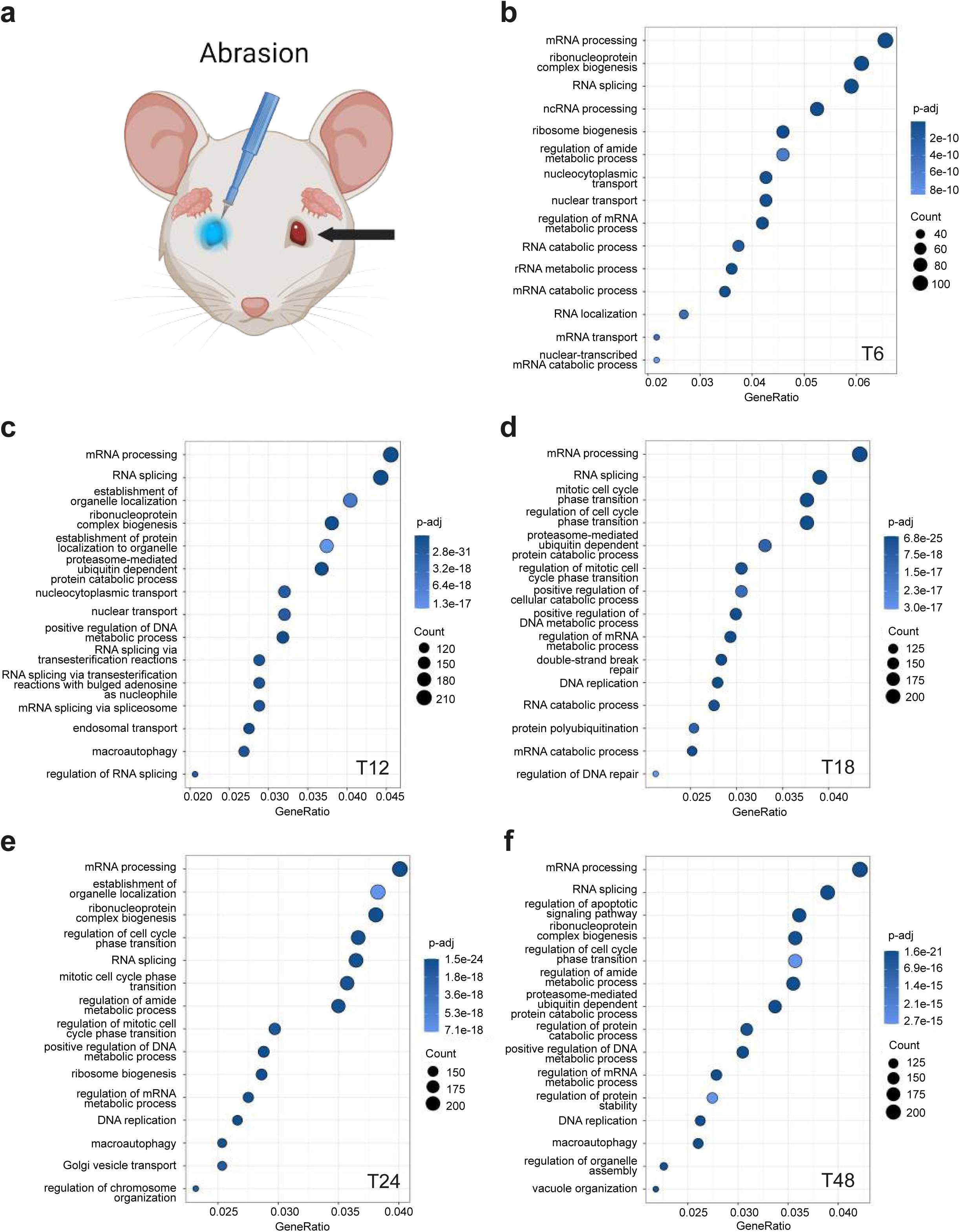
Corneal abrasion induces the downregulation of various biological processes related to cell cycle regulation as well as RNA and protein processing in the contralateral side. (a) Schematic overview of the experimental design. Dot plot representations of Gene Ontology (GO) enrichment analysis performed in R from RNA-Seq data on the contralateral side of the abrasion. (b, c, d, e, f) The top 15 downregulated biological processes are shown for each plot, from 6H to 48H post-abrasion. GeneRatio represents the number of differentially expressed genes in each GO term (Count), divided by the number of genes in that GO term.

**Figure S7.**
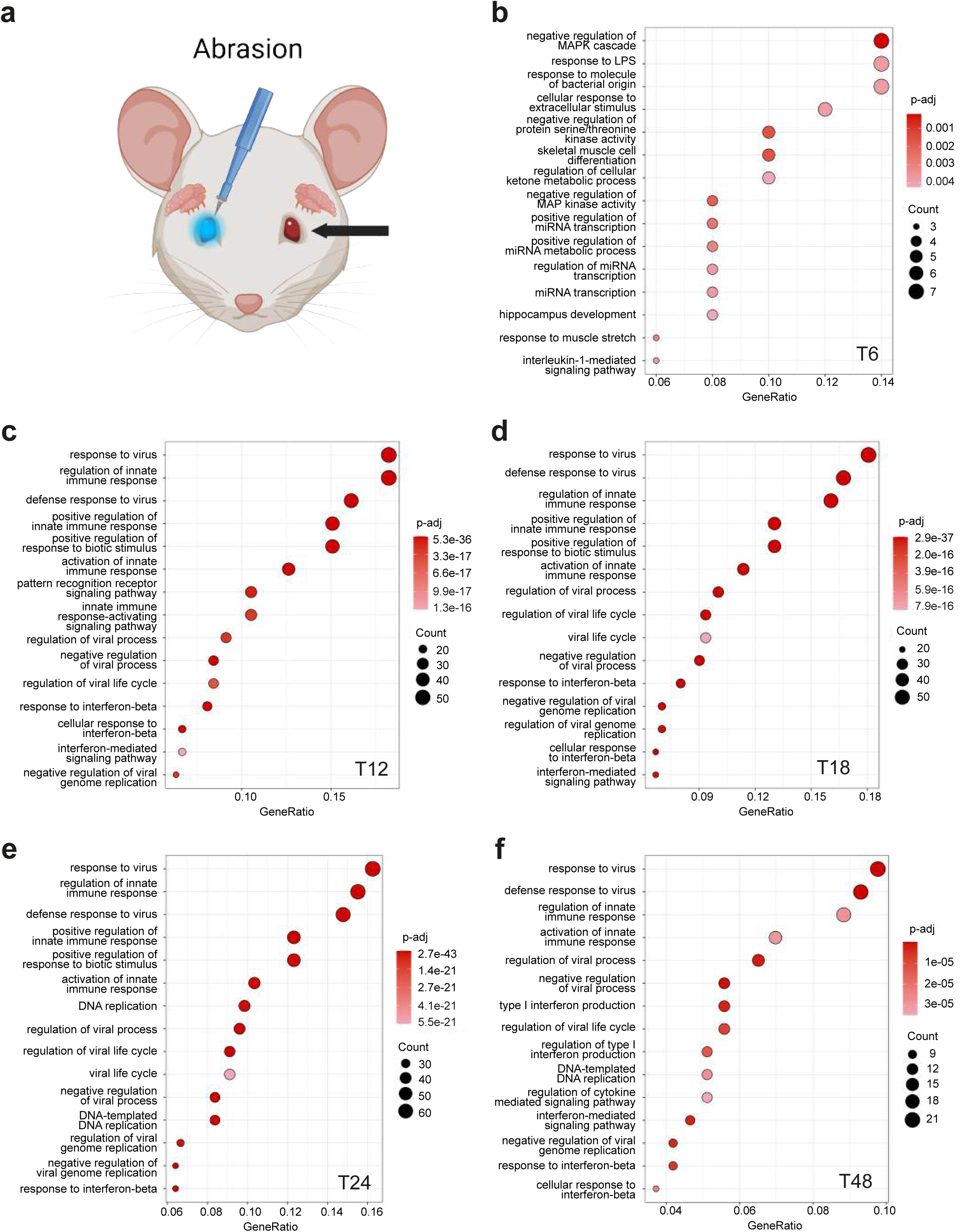
Corneal abrasion induces the upregulation of various biological processes linked to the inflammatory response in the contralateral side. (a) Schematic overview of the experimental design. Dot plot representations of Gene Ontology (GO) enrichment analysis performed in R from RNA-Seq data on the contralateral side of the abrasion. (b, c, d, e, f) The top 15 upregulated biological processes are shown for each plot, from 6H to 48H post-abrasion. GeneRatio represents the number of differentially expressed genes in each GO term (Count), divided by the number of genes in that GO term.

**Figure S8.**
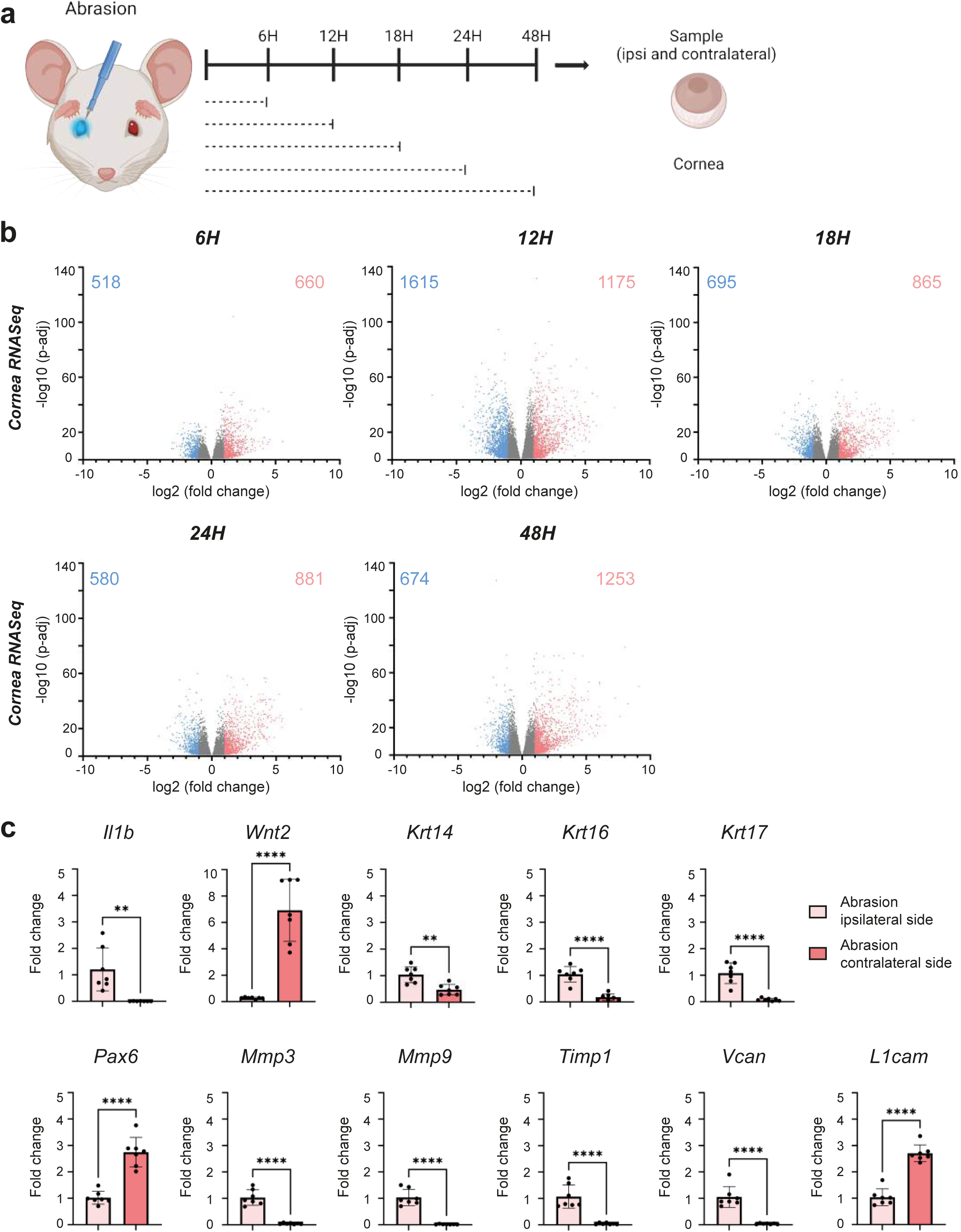
Corneal abrasion induces bilateral molecular changes in the cornea. (a) Schematic overview of the experimental design. Abrasion was performed unilaterally. Samples from the ipsilateral and contralateral side of the abrasion were taken before the abrasion as a control and at 6H, 12H, 18H, 24H and 48H post-abrasion. (b) Volcano plot representations of cornea RNA-Seq analysis (n=5 per group) at 6H, 12H, 18H, 24H and 48H post-abrasion vs control. Statistically significant genes are represented only. Genes with a fold change lower than 0.5 are colored in blue and their number is indicated in the top left corner of each plot. Genes with a fold change higher than 2 are colored in red and their number is indicated in the top right corner of each plot. The genes with an intermediate fold change are colored in grey. (c) Histograms of RT-qPCR analysis on cornea samples at 18H post-abrasion ipsilateral vs contralateral (n=7 per group). Data are represented as mean ± SD and normalized to a reference gene (*Ppia*). Statistical significance was assessed by unpaired two-tailed t-test (*, p < 0.05; **, p < 0.01; ***, p < 0.001, ****, p < 0.0001).

**Figure S9.**
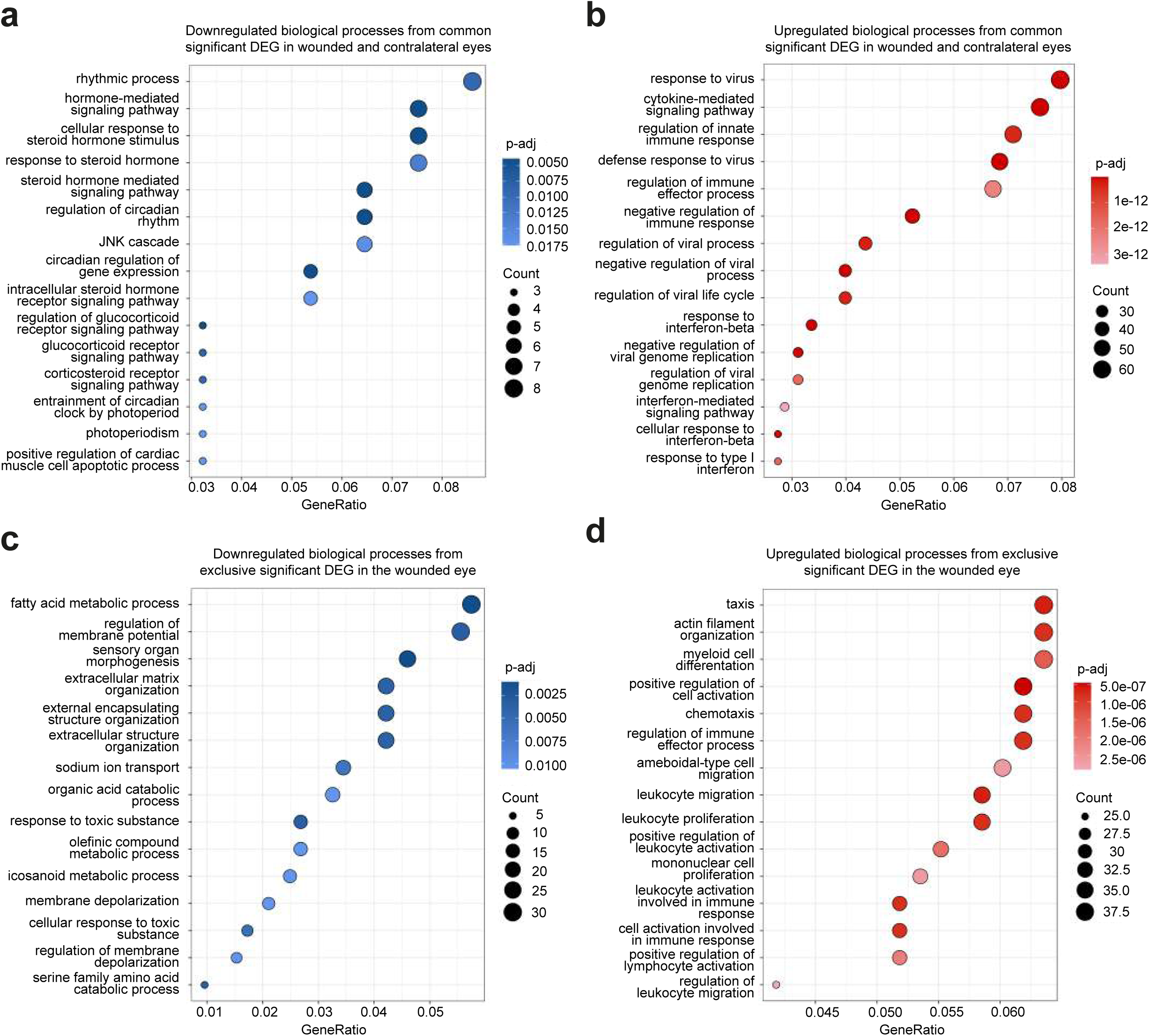
Corneal abrasion induces the downregulation of specific biological processes in each eye, but a similar upregulation of processes related to immune response. Dot plot representations of Gene Ontology (GO) enrichment analysis performed in R from RNA-Seq data on the wounded cornea and the contralateral cornea. (a, b) The top 15 modified biological processes, from common significant DEG in both eyes, are shown for each plot, at 18H post-abrasion. (c, d) The top 15 modified biological processes, from significant DEG in the wounded eye solely, are shown for each plot, at 18H post-abrasion. The GeneRatio represents the number of differentially expressed genes in each GO term (Count), divided by the number of genes in that GO term.

**Figure S10.**
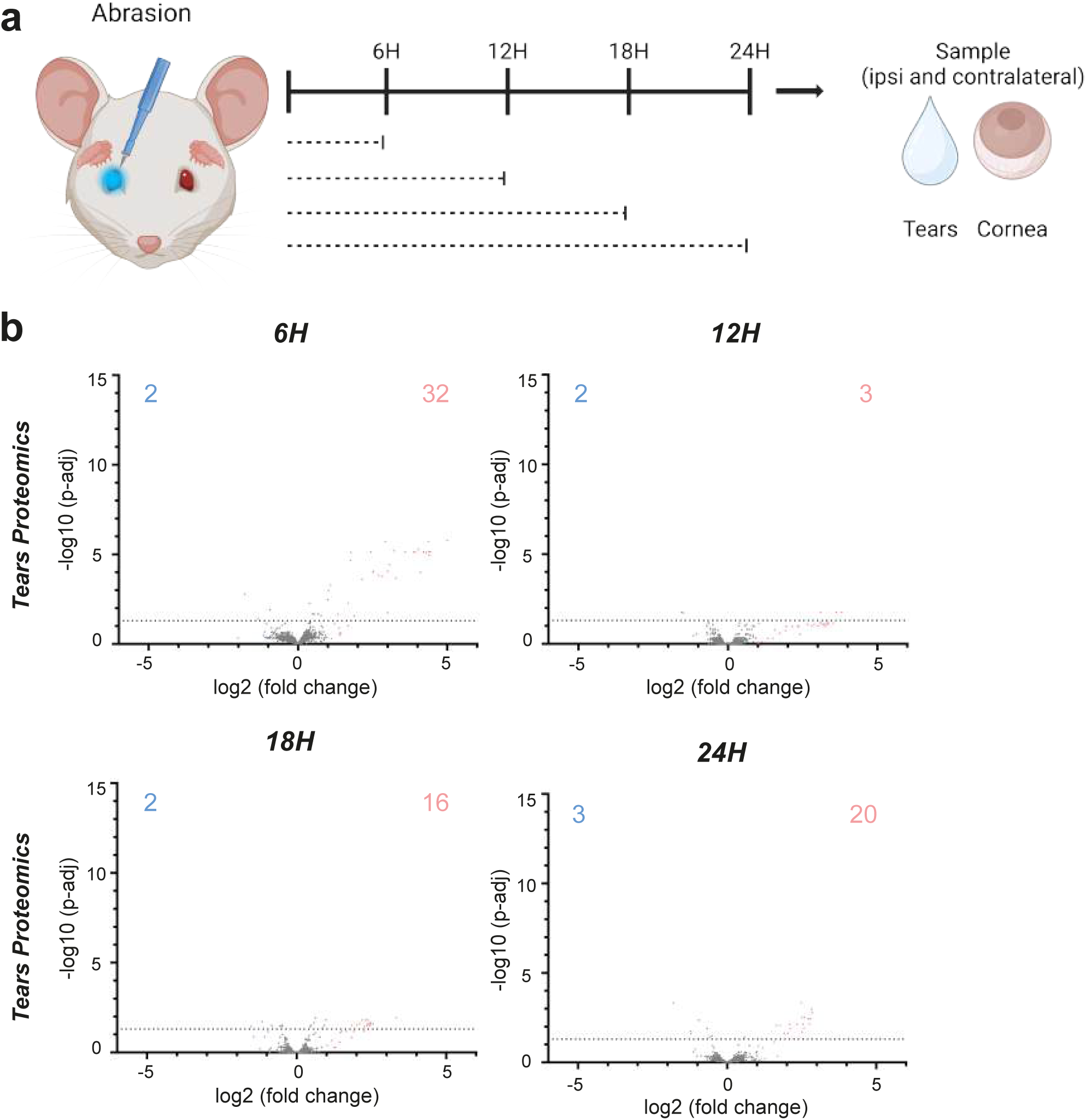
Corneal abrasion induces bilateral molecular changes in tears. (a) Schematic overview of the experimental design. Abrasion was performed unilaterally. Samples from the ipsilateral and contralateral side of the abrasion were taken before the abrasion as a control and at 6H, 12H, 18H and 24H post-abrasion. (b) Volcano plot representations of tears proteomics analysis by mass spectrometry (n=7 per group) at 6H, 12H, 18H and 24H post-abrasion vs control. All identified proteins are represented and the limit of statistical significance is shown with the horizontal dotted line. Proteins with a fold change lower than 0.5 are colored in blue and their number is indicated in the top left corner of each plot. Proteins with a fold change higher than 2 are colored in red and their number is indicated in the top right corner of each plot. The proteins with an intermediate fold change are colored in grey.

**Figure S11.**
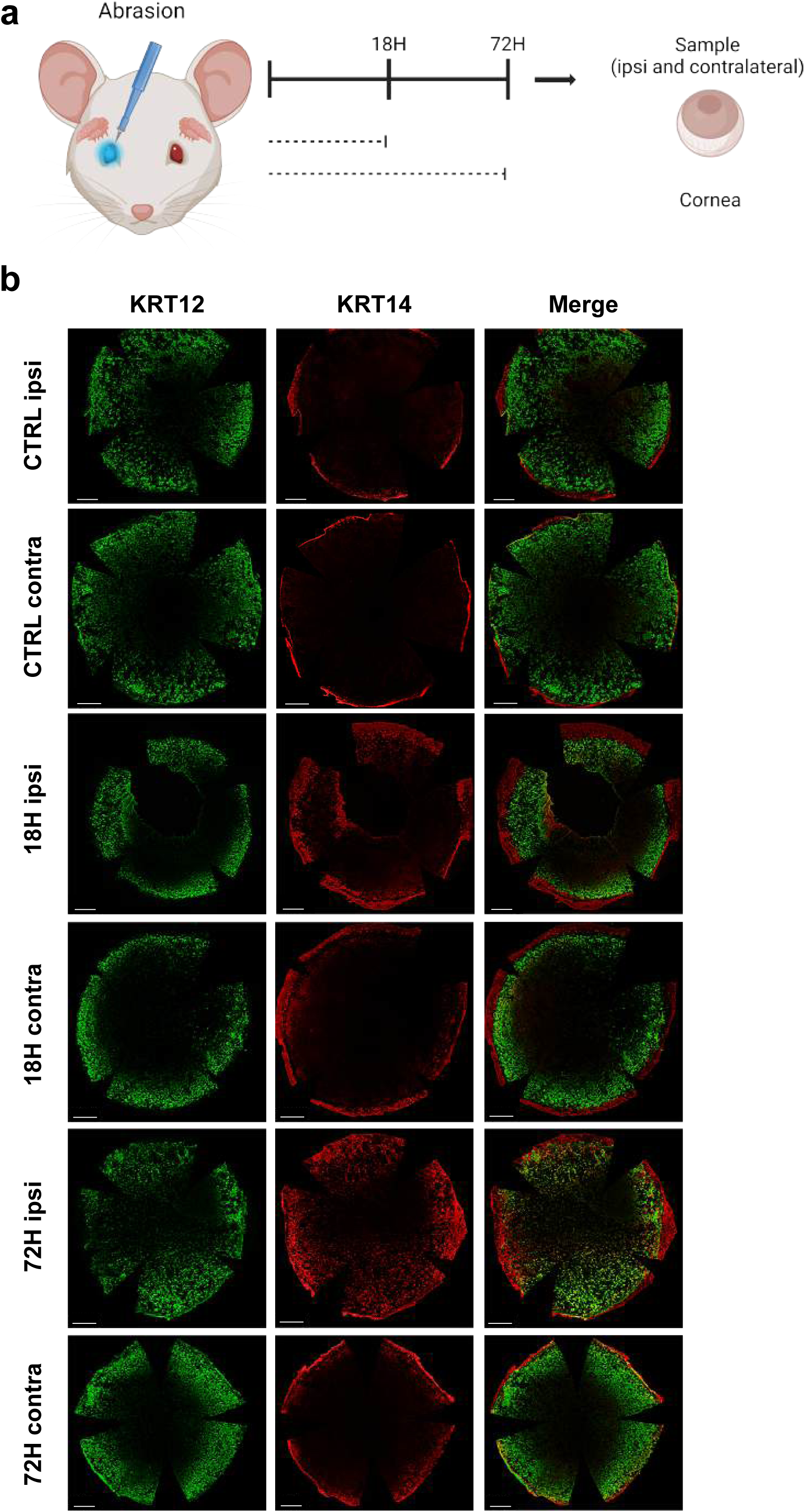
The corneal abrasion induces a limited bilateral cellular response throughout the wound healing process. (a) Schematic overview of the experimental design. Abrasion was performed unilaterally. Samples from the ipsilateral and contralateral side of the abrasion were taken before the surgery as a control and at 18H and 72H post-surgery. (b) Wholemount immunofluorescence staining for differentiated (KRT12) and undifferentiated (KRT14) corneal markers. Data are representative of 3 biological replicates. Nuclei were counterstained with Hoechst (not shown). Scale bars are 500µm.

**Table S1.** Overview of the evolution of KEGG molecular pathways, based on cornea RNA-Seq results after abrasion. Data are shown as the percentage of upregulated (red) or downregulated (blue) genes among each pathway, from 6H to 48H post-abrasion.

|  |  |  | Molecular pathways (KEGG) |  |  |  |  |  |  |  |  |  |
| --- | --- | --- | --- | --- | --- | --- | --- | --- | --- | --- | --- | --- |
| | | | MAPK | PI3K-Akt | TGF-β | Notch | Wnt | Hedgehog | JAK-STAT | NF-kB | HIF-1 | VEGF |
| Abrasion (ipsilateral)<br>vs control | 6H | % up | 19.8 | 21.6 | 16.7 | 9.7 | 8.8 | 0.0 | 23.5 | 29.5 | 28.8 | 25.0 |
|  |  | % down | 3.3 | 5.7 | 8.3 | 6.5 | 4.4 | 5.1 | 7.1 | 1.4 | 4.5 | 0.0 |
|  | 12H | % up | 21.5 | 18.2 | 22.6 | 16.1 | 4.4 | 5.1 | 38.8 | 17.3 | 25.8 | 21.4 |
|  |  | % down | 10.7 | 17.0 | 8.3 | 6.5 | 18.7 | 11.9 | 10.6 | 5.8 | 15.2 | 10.7 |
|  | 18H | % up | 12.4 | 18.2 | 13.1 | 9.7 | 8.8 | 1.7 | 30.6 | 7.2 | 21.2 | 17.9 |
|  |  | % down | 10.7 | 8.0 | 6.0 | 6.5 | 17.6 | 6.8 | 7.1 | 2.2 | 13.6 | 3.6 |
|  | 24H | % up | 13.2 | 18.2 | 15.5 | 12.9 | 11.0 | 5.1 | 34.1 | 6.5 | 15.2 | 14.3 |
|  |  | % down | 13.2 | 10.2 | 7.1 | 3.2 | 18.7 | 5.1 | 7.1 | 1.4 | 12.1 | 7.1 |
|  | 48H | % up | 40.5 | 38.6 | 27.4 | 38.7 | 36.3 | 42.4 | 54.1 | 27.3 | 50.0 | 42.9 |
|  |  | % down | 53.7 | 44.3 | 50.0 | 48.4 | 54.9 | 33.9 | 56.5 | 41.7 | 33.3 | 50.0 |
| Abrasion (contralateral)<br>vs control | 6H | % up | 7.4 | 3.4 | 1.2 | 0.0 | 3.3 | 0.0 | 2.4 | 7.2 | 1.5 | 3.6 |
|  |  | % down | 0.0 | 0.0 | 0.0 | 0.0 | 0.0 | 0.0 | 0.0 | 0.0 | 0.0 | 0.0 |
|  | 12H | % up | 5.0 | 10.2 | 3.6 | 19.4 | 8.8 | 1.7 | 22.4 | 2.9 | 7.6 | 7.1 |
|  |  | % down | 5.8 | 5.7 | 1.2 | 0.0 | 6.6 | 0.0 | 2.4 | 1.4 | 1.5 | 0.0 |
|  | 18H | % up | 1.7 | 8.0 | 1.2 | 19.4 | 6.6 | 1.7 | 20.0 | 0.7 | 3.0 | 3.6 |
|  |  | % down | 13.2 | 8.0 | 4.8 | 0.0 | 11.0 | 5.1 | 4.7 | 13.7 | 4.5 | 3.6 |
|  | 24H | % up | 2.5 | 6.8 | 2.4 | 16.1 | 4.4 | 1.7 | 20.0 | 1.4 | 6.1 | 3.6 |
|  |  | % down | 9.9 | 9.1 | 6.0 | 0.0 | 9.9 | 5.1 | 4.7 | 12.9 | 7.6 | 10.7 |
|  | 48H | % up | 2.5 | 10.2 | 1.2 | 12.9 | 6.6 | 5.1 | 3.5 | 0.7 | 3.0 | 3.6 |
|  |  | % down | 11.6 | 10.2 | 8.3 | 0.0 | 6.6 | 0.0 | 4.7 | 15.1 | 6.1 | 7.1 |
| Abrasion (ipsilateral)<br>vs abrasion (contralateral) | 6H | % up | 17.4 | 15.9 | 14.3 | 9.7 | 5.5 | 0.0 | 20.0 | 25.9 | 21.2 | 25.0 |
|  |  | % down | 1.7 | 4.5 | 3.6 | 3.2 | 3.3 | 0.0 | 4.7 | 0.0 | 4.5 | 0.0 |
|  | 12H | % up | 24.0 | 21.6 | 23.8 | 6.5 | 7.7 | 5.1 | 23.5 | 12.2 | 28.8 | 32.1 |
|  |  | % down | 8.3 | 17.0 | 6.0 | 9.7 | 13.2 | 11.9 | 11.8 | 5.0 | 13.6 | 7.1 |
|  | 18H | % up | 13.2 | 18.2 | 13.1 | 3.2 | 5.5 | 0.0 | 12.9 | 9.4 | 25.8 | 21.4 |
|  |  | % down | 3.3 | 4.5 | 3.6 | 6.5 | 6.6 | 3.4 | 8.2 | 0.7 | 12.1 | 3.6 |
|  | 24H | % up | 17.4 | 21.6 | 16.7 | 6.5 | 11.0 | 1.7 | 14.1 | 18.7 | 18.2 | 21.4 |
|  |  | % down | 3.3 | 5.7 | 0.0 | 3.2 | 2.2 | 0.0 | 7.1 | 1.4 | 9.1 | 3.6 |
|  | 48H | % up | 16.5 | 15.9 | 15.5 | 6.5 | 9.9 | 10.2 | 14.1 | 7.2 | 24.2 | 17.9 |
|  |  | % down | 4.1 | 8.0 | 1.2 | 0.0 | 6.6 | 5.1 | 5.9 | 7.2 | 9.1 | 0.0 |

**Table S2.** Overview of the evolution of KEGG process pathways, based on cornea RNA-Seq results after abrasion. Data are shown as the percentage of upregulated (red) or downregulated (blue) genes among each pathway, from 6H to 48H post-abrasion.

|  |  |  | Process pathways (KEGG) |  |  |  |  |  |  |
| --- | --- | --- | --- | --- | --- | --- | --- | --- | --- |
|  |  |  | Focal adhesion | Chemokine | Calcium | Neurotrophin | Apoptosis | Cell cycle | Stem cells pluripotency |
| Abrasion (ipsilateral)<br>vs control | 6H | % up | 17.7 | 19.0 | 25.0 | 11.7 | 22.6 | 8.9 | 13.3 |
|  |  | % down | 6.5 | 1.7 | 5.4 | 2.6 | 0.0 | 3.6 | 13.3 |
|  | 12H | % up | 21.0 | 27.6 | 19.6 | 14.3 | 26.4 | 4.5 | 17.3 |
|  |  | % down | 16.1 | 13.8 | 26.8 | 6.5 | 11.3 | 17.9 | 15.3 |
|  | 18H | % up | 17.7 | 20.7 | 17.9 | 7.8 | 14.2 | 11.6 | 11.2 |
|  |  | % down | 12.9 | 6.9 | 26.8 | 6.5 | 5.7 | 8.0 | 10.2 |
|  | 24H | % up | 17.7 | 22.4 | 19.6 | 11.7 | 19.8 | 20.5 | 13.3 |
|  |  | % down | 12.9 | 10.3 | 30.4 | 9.1 | 4.7 | 8.9 | 10.2 |
|  | 48H | % up | 45.2 | 41.4 | 48.2 | 32.5 | 43.4 | 62.5 | 37.8 |
|  |  | % down | 54.8 | 55.2 | 51.8 | 53.2 | 33.0 | 39.3 | 46.9 |
| Abrasion (contralateral)<br>vs control | 6H | % up | 3.2 | 3.4 | 3.6 | 5.2 | 4.7 | 0.0 | 2.0 |
|  |  | % down | 0.0 | 0.0 | 0.0 | 0.0 | 0.0 | 0.0 | 0.0 |
|  | 12H | % up | 12.9 | 13.8 | 10.7 | 6.5 | 6.6 | 0.0 | 10.2 |
|  |  | % down | 1.6 | 0.0 | 3.6 | 2.6 | 3.8 | 0.9 | 3.1 |
|  | 18H | % up | 9.7 | 10.3 | 7.1 | 1.3 | 1.9 | 5.4 | 6.1 |
|  |  | % down | 6.5 | 5.2 | 8.9 | 7.8 | 17.9 | 1.8 | 8.2 |
|  | 24H | % up | 8.1 | 12.1 | 8.9 | 2.6 | 3.8 | 13.4 | 6.1 |
|  |  | % down | 6.5 | 3.4 | 14.3 | 6.5 | 12.3 | 4.5 | 5.1 |
|  | 48H | % up | 6.5 | 6.9 | 5.4 | 2.6 | 1.9 | 6.3 | 6.1 |
|  |  | % down | 4.8 | 5.2 | 10.7 | 6.5 | 17.0 | 0.9 | 4.1 |
| Abrasion (ipsilateral)<br>vs abrasion (contralateral) | 6H | % up | 16.1 | 17.2 | 19.6 | 9.1 | 19.8 | 7.1 | 10.2 |
|  |  | % down | 3.2 | 0.0 | 3.6 | 1.3 | 0.0 | 2.7 | 8.2 |
|  | 12H | % up | 27.4 | 22.4 | 26.8 | 16.9 | 23.6 | 5.4 | 16.3 |
|  |  | % down | 12.9 | 10.3 | 23.2 | 7.8 | 11.3 | 11.6 | 13.3 |
|  | 18H | % up | 22.6 | 20.7 | 17.9 | 11.7 | 16.0 | 1.8 | 9.2 |
|  |  | % down | 8.1 | 5.2 | 16.1 | 2.6 | 1.9 | 0.9 | 5.1 |
|  | 24H | % up | 24.2 | 27.6 | 23.2 | 13.0 | 23.6 | 1.8 | 12.2 |
|  |  | % down | 9.7 | 5.2 | 19.6 | 1.3 | 1.9 | 0.9 | 4.1 |
|  | 48H | % up | 22.6 | 22.4 | 23.2 | 11.7 | 16.0 | 17.9 | 9.2 |
|  |  | % down | 8.1 | 3.4 | 16.1 | 1.3 | 0.9 | 1.8 | 7.1 |

**Table S3.** Overview of the epitranscriptomics modifications compared to control. Data are shown as the percentage of variation of the area under curve ratio (modified nucleoside/uridine) in the various experimental conditions compared to control, at 18H and 72H post-surgery. Statistically significant negative variations are shown in blue. All non-significant variations are shown in grey. Statistical significance was assessed by unpaired two-tailed t-test (*, p < 0.05; **, p < 0.01; ***, p < 0.001, ****, p < 0.0001).

| Condition<br>vs control | Abrasion (ipsilateral) |  | Abrasion + ipsilateral LG<br>ablation (ipsilateral) |  | Abrasion (contralateral) |  | Abrasion + contralateral<br>LG ablation (contralateral) |  | Ablation (ipsilateral) |  |
| --- | --- | --- | --- | --- | --- | --- | --- | --- | --- | --- |
|  | 18H | 72H | 18H | 72H | 18H | 72H | 18H | 72H | 18H | 72H |
| ac<sup>4</sup>C | -12.6 * | -12.6 * | -24.1 ** | -12.6 <sup>ns</sup> | 3.4 <sup>ns</sup> | 1.1 <sup>ns</sup> | -3.4 <sup>ns</sup> | 3.4 <sup>ns</sup> | 3.4 <sup>ns</sup> | -5.7 <sup>ns</sup> |
| G<sub>m</sub> | -5.2 <sup>ns</sup> | -17.2 * | -15.5 * | -20.7 ** | -6.9 <sup>ns</sup> | -6.9 <sup>ns</sup> | -10.3 <sup>ns</sup> | -6.9 <sup>ns</sup> | -5.2 <sup>ns</sup> | -5.2 <sup>ns</sup> |
| i<sup>6</sup>A | -27.2 *** | -20.8 ** | -33.3 *** | -14.3 * | -4.0 <sup>ns</sup> | 6.3 <sup>ns</sup> | -7.9 <sup>ns</sup> | -0.2 <sup>ns</sup> | -2.7 <sup>ns</sup> | -17.9 <sup>ns</sup> |
| m<sup>1</sup>A | -21.4 *** | -16.3 ** | -19.7 * | -13.4 * | -2.8 <sup>ns</sup> | 6.6 <sup>ns</sup> | -4.0 <sup>ns</sup> | 4.0 <sup>ns</sup> | -0.6 <sup>ns</sup> | -9.6 <sup>ns</sup> |
| m<sup>3</sup>C | -24.6 *** | -23.0 *** | -32.4 *** | -26.2 *** | -7.4 <sup>ns</sup> | -0.4 <sup>ns</sup> | -9.4 * | -4.9 <sup>ns</sup> | -7.8 <sup>ns</sup> | -21.7 ** |
| m<sup>5</sup>U | -19.0 <sup>ns</sup> | -16.5 <sup>ns</sup> | -18.3 <sup>ns</sup> | -6.9 <sup>ns</sup> | -4.7 <sup>ns</sup> | 4.3 <sup>ns</sup> | -9.5 <sup>ns</sup> | -1.6 <sup>ns</sup> | -1.2 <sup>ns</sup> | -4.8 <sup>ns</sup> |
| m<sup>6</sup>A | -2.5 <sup>ns</sup> | -4.6 <sup>ns</sup> | -32.0 * | -45.2 *** | 5.6 <sup>ns</sup> | -2.5 <sup>ns</sup> | 4.6 <sup>ns</sup> | -2.5 <sup>ns</sup> | 8.6 <sup>ns</sup> | -12.7 <sup>ns</sup> |
| m<sup>7</sup>G | -26.2 **** | -11.5 ** | -23.0 ** | -16.4 ** | -1.6 <sup>ns</sup> | 8.2 <sup>ns</sup> | -4.9 <sup>ns</sup> | 1.6 <sup>ns</sup> | -1.6 <sup>ns</sup> | -14.8 * |
| m<sup>2,2</sup>G | -17.0 * | -17.5 * | -26.2 ** | -9.7 <sup>ns</sup> | -4.4 <sup>ns</sup> | 1.0 <sup>ns</sup> | -9.2 <sup>ns</sup> | -3.4 <sup>ns</sup> | -4.4 <sup>ns</sup> | -15.0 * |
| m<sup>6,6</sup>A | -29.7 *** | -13.5 ** | -36.5 *** | -7.4 <sup>ns</sup> | -3.4 <sup>ns</sup> | 6.8 <sup>ns</sup> | -8.1 <sup>ns</sup> | 3.4 <sup>ns</sup> | -2.7 <sup>ns</sup> | -18.9 * |
| mcm<sup>5</sup>s<sup>2</sup>U | -24.2 ** | -14.5 * | -22.6 ** | -8.1 <sup>ns</sup> | -8.1 <sup>ns</sup> | 3.2 <sup>ns</sup> | -9.7 <sup>ns</sup> | -1.6 <sup>ns</sup> | -3.2 <sup>ns</sup> | -6.5 <sup>ns</sup> |
| ncm<sup>5</sup>U | -11.9 <sup>ns</sup> | -4.4 <sup>ns</sup> | -11.9 <sup>ns</sup> | -1.9 <sup>ns</sup> | -0.6 <sup>ns</sup> | 1.9 <sup>ns</sup> | -5.7 <sup>ns</sup> | -5.7 <sup>ns</sup> | -4.4 <sup>ns</sup> | -5.7 <sup>ns</sup> |
| Queuosine | -19.0 <sup>ns</sup> | -26.1 <sup>ns</sup> | -21.0 <sup>ns</sup> | -19.5 <sup>ns</sup> | -3.8 <sup>ns</sup> | 1.3 <sup>ns</sup> | -11.4 <sup>ns</sup> | -3.8 <sup>ns</sup> | -13.9 <sup>ns</sup> | -18.5 <sup>ns</sup> |
| U<sub>m</sub> | -14.3 * | -16.2 ** | -20.0 ** | -27.6 *** | -2.9 <sup>ns</sup> | 1.0 <sup>ns</sup> | -6.7 <sup>ns</sup> | -6.7 <sup>ns</sup> | -4.8 <sup>ns</sup> | -8.6 <sup>ns</sup> |

**Table S4.**
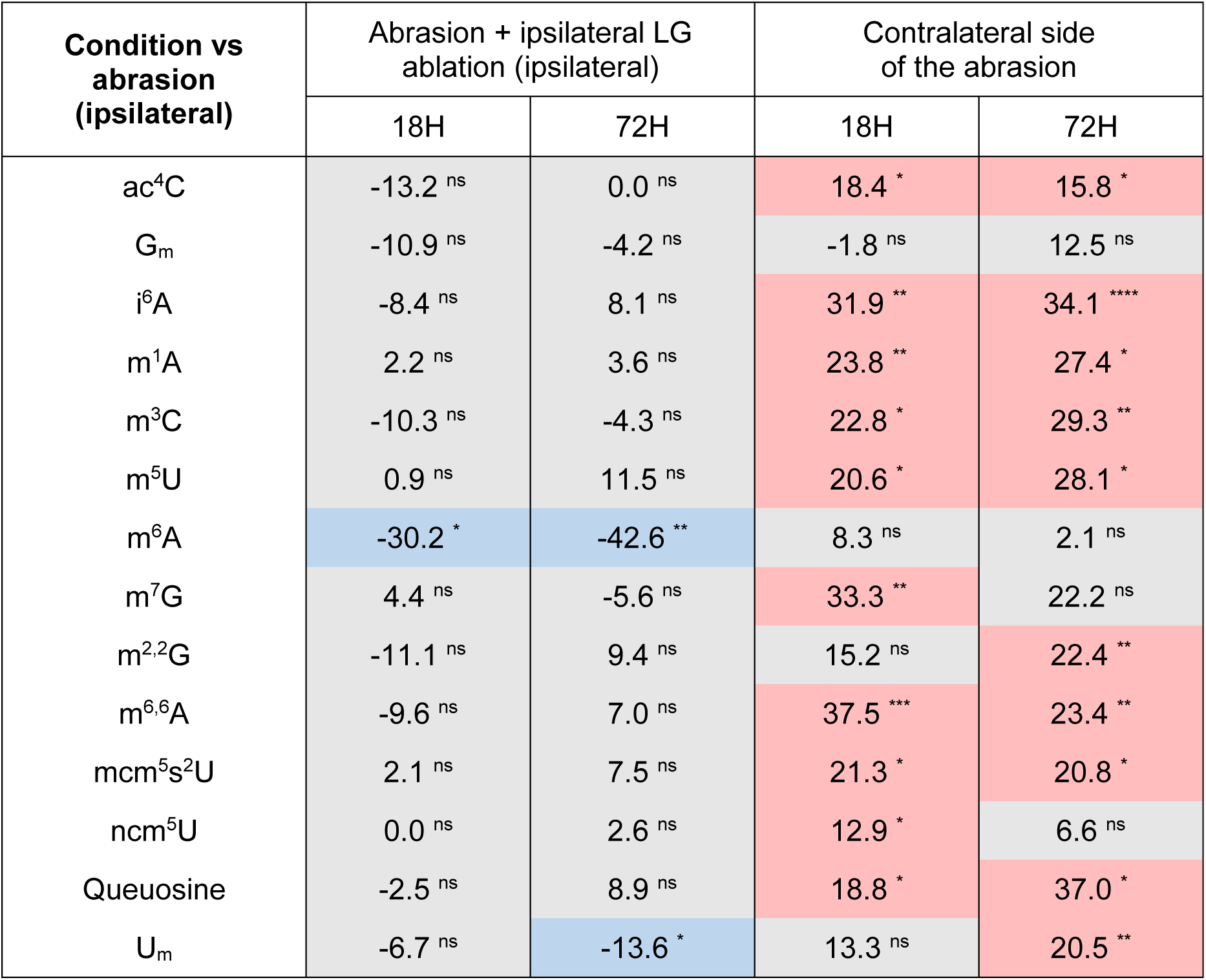
Overview of the epitranscriptomics modifications compared to the abrasion. Data are shown as the percentage of variation of the area under curve ratio (modified nucleoside/uridine) in the various experimental conditions compared to the ipsilateral side of the abrasion, at 18H and 72H post-surgery. Statistically significant negative variations are shown in blue. Statistically significant positive variations are shown in red. All non-significant variations are shown in grey. Statistical significance was assessed by unpaired two-tailed t-test (*, p < 0.05; **, p < 0.01; ***, p < 0.001, ****, p < 0.0001).

**Table S5.**
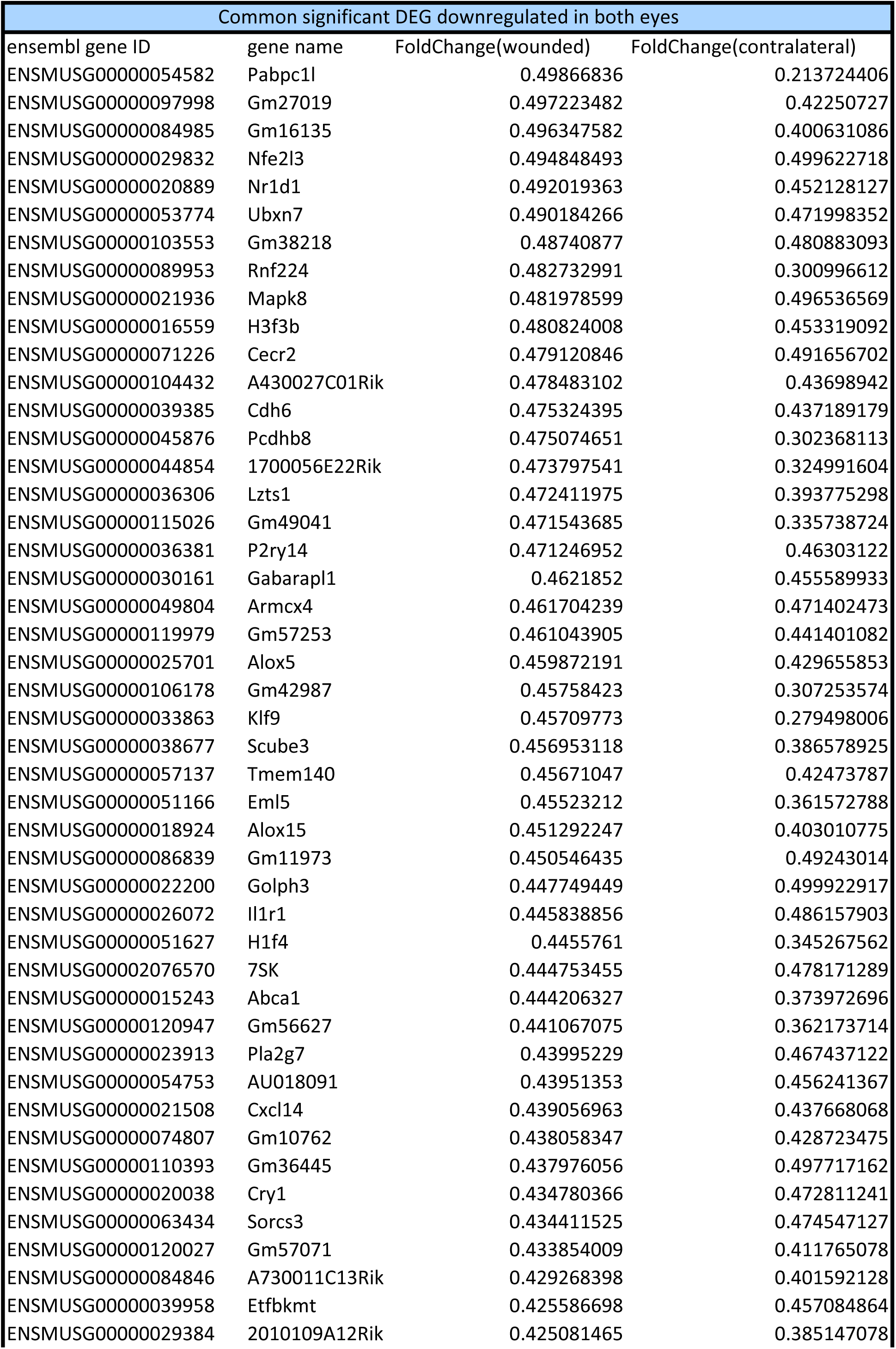
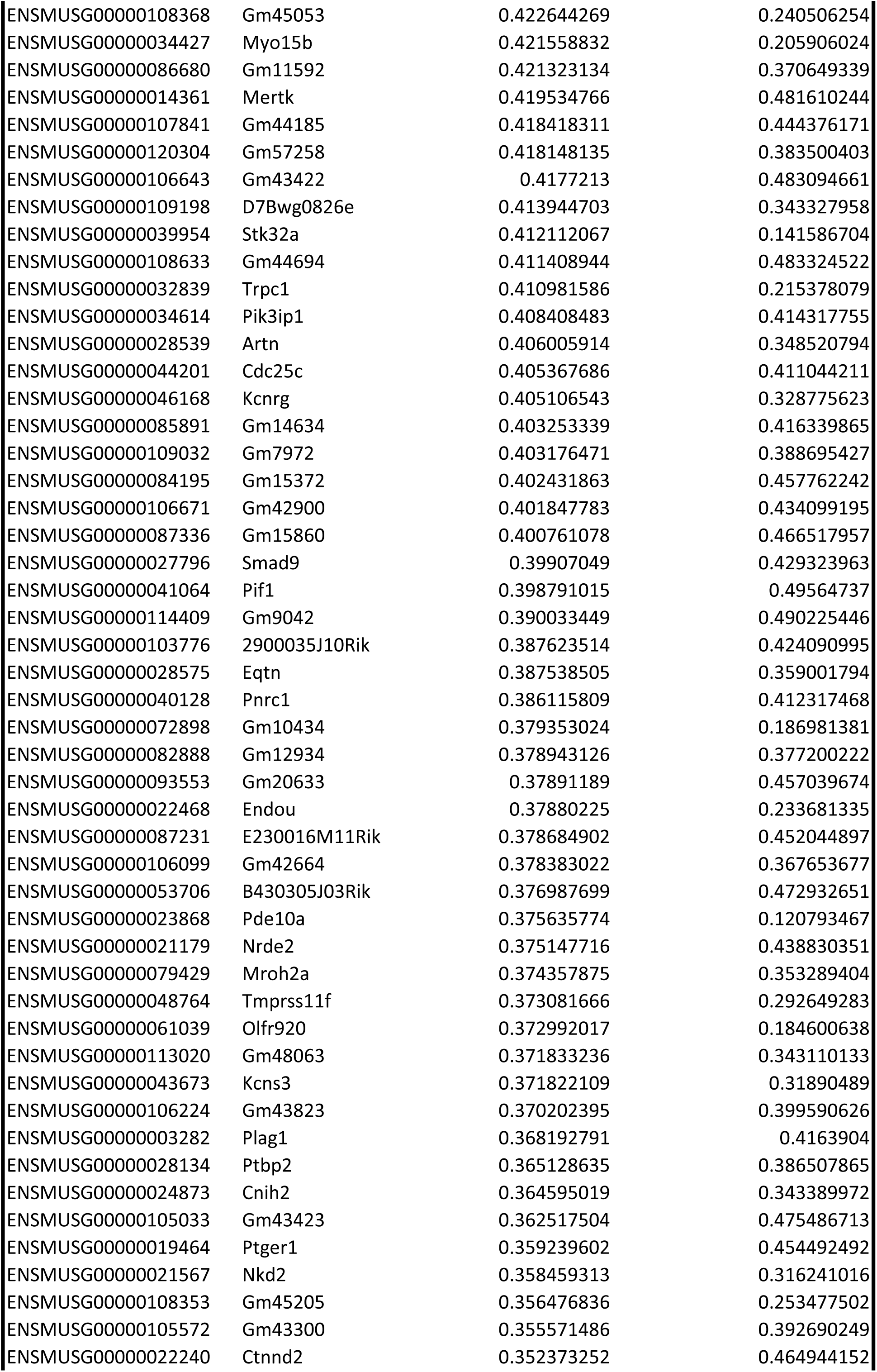
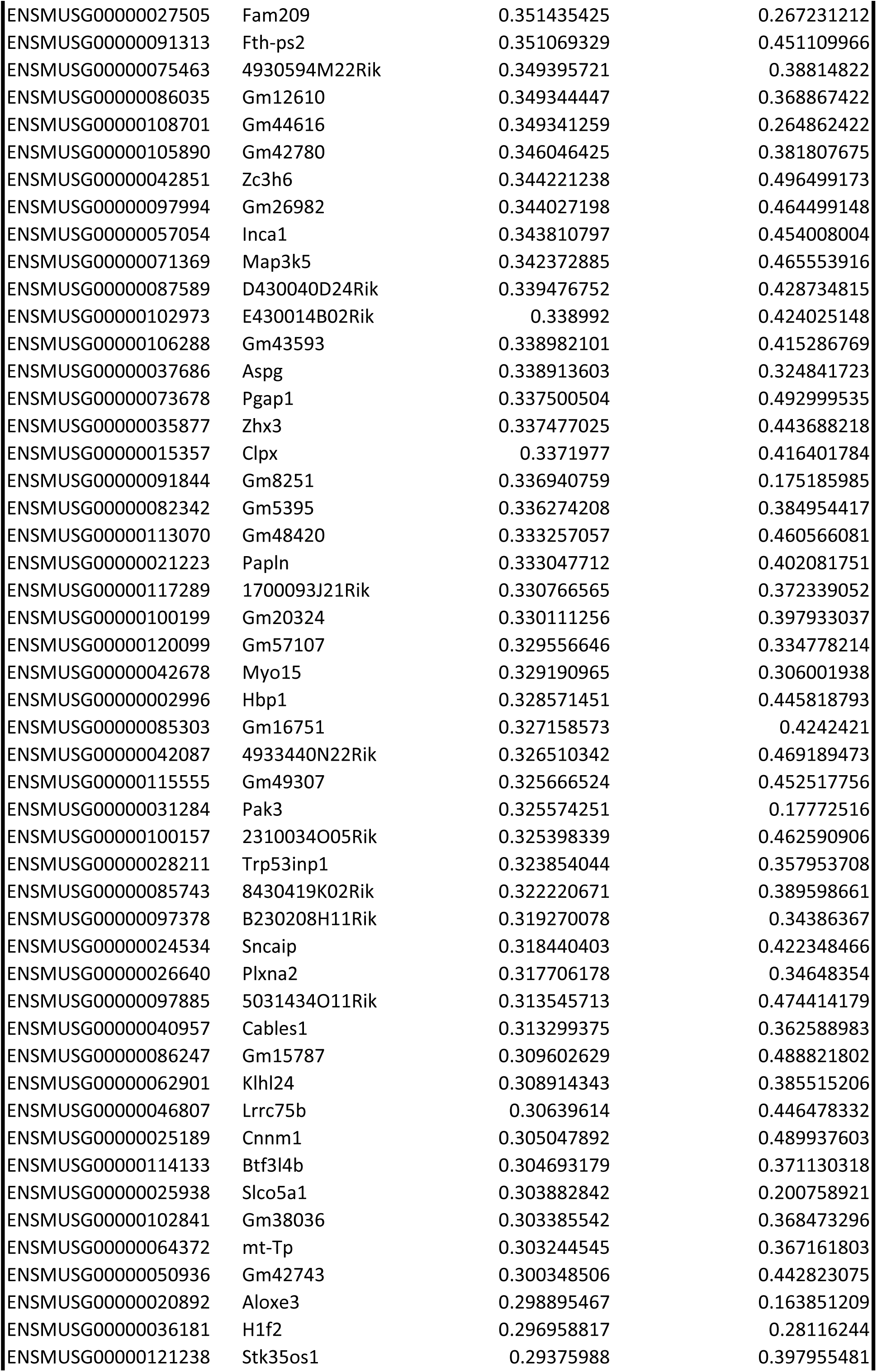
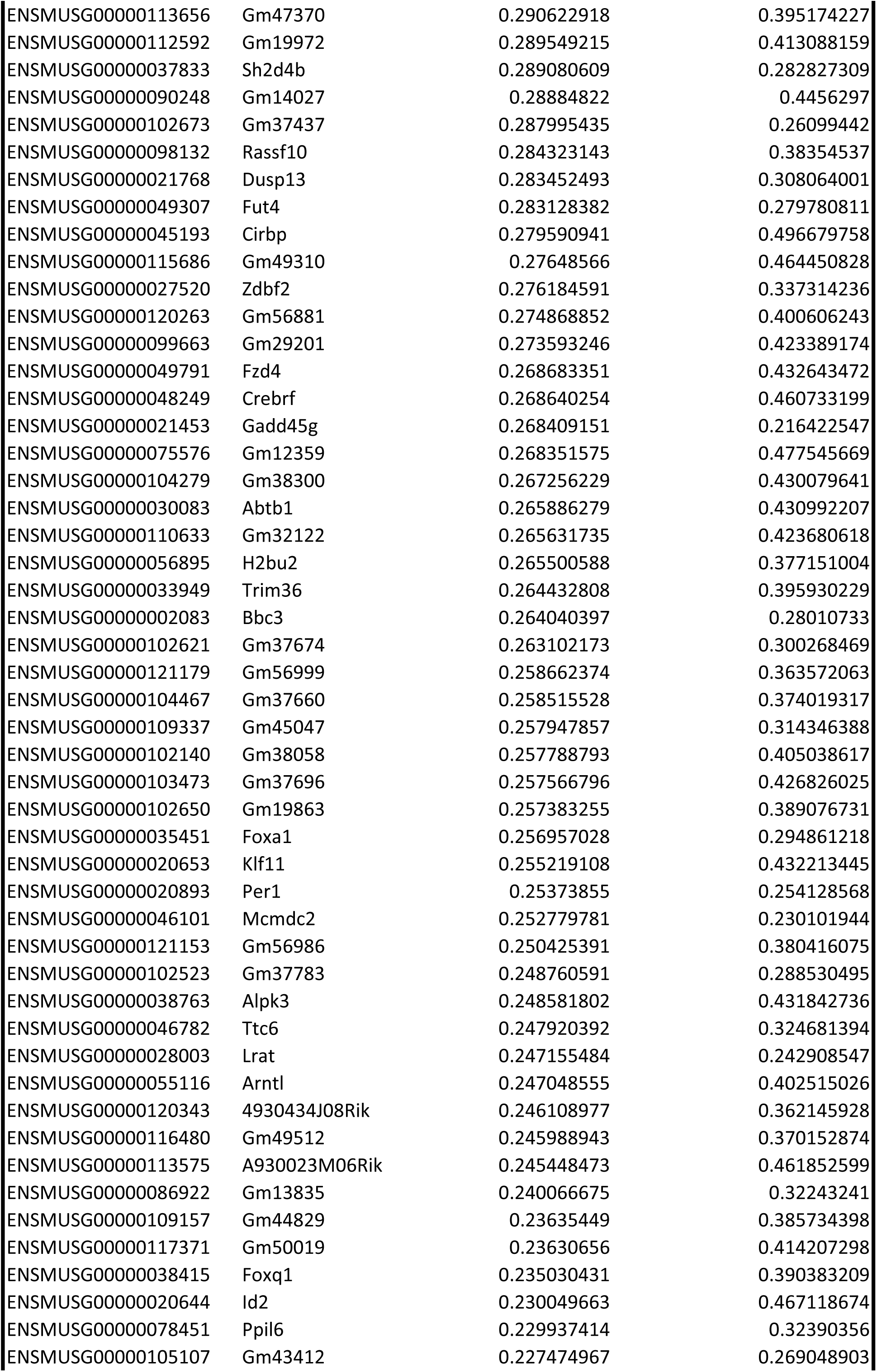
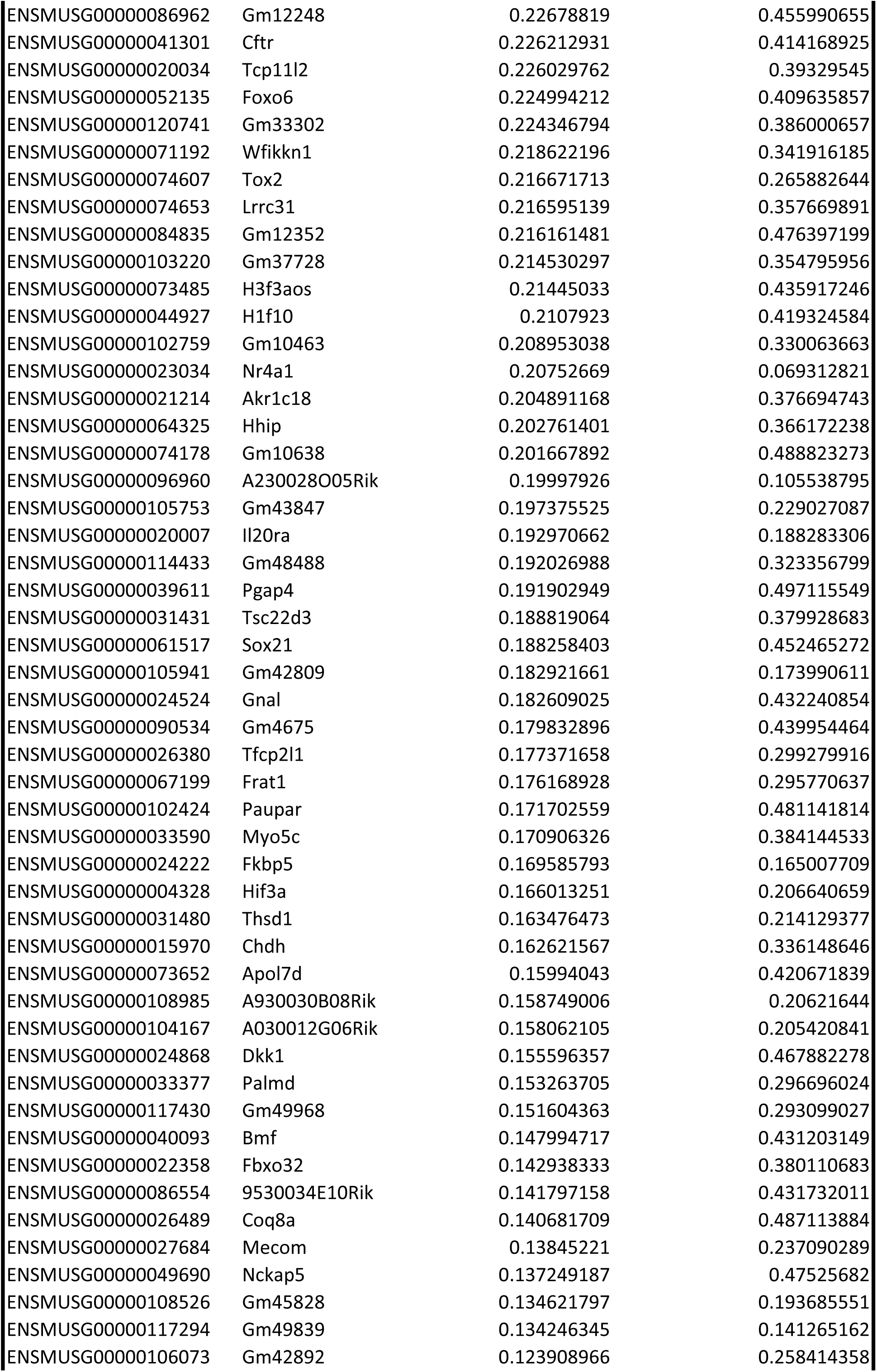
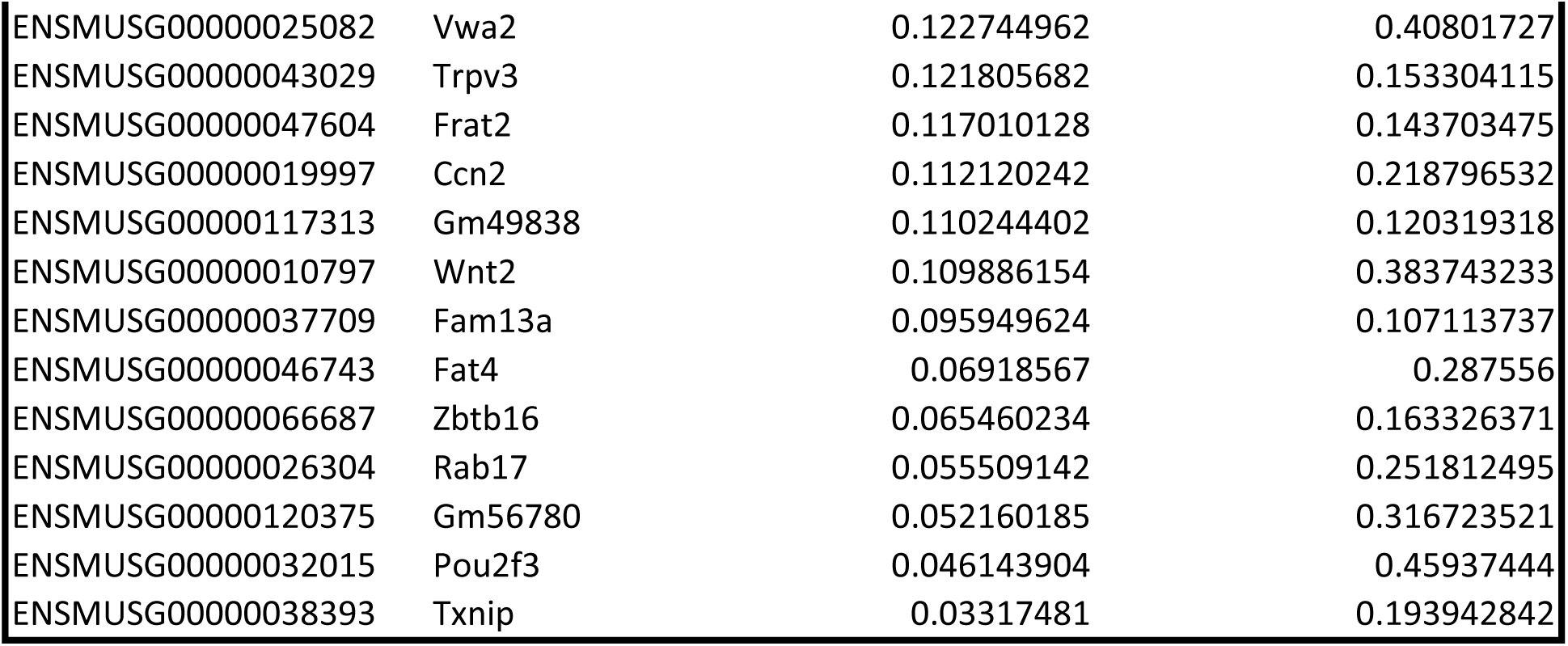
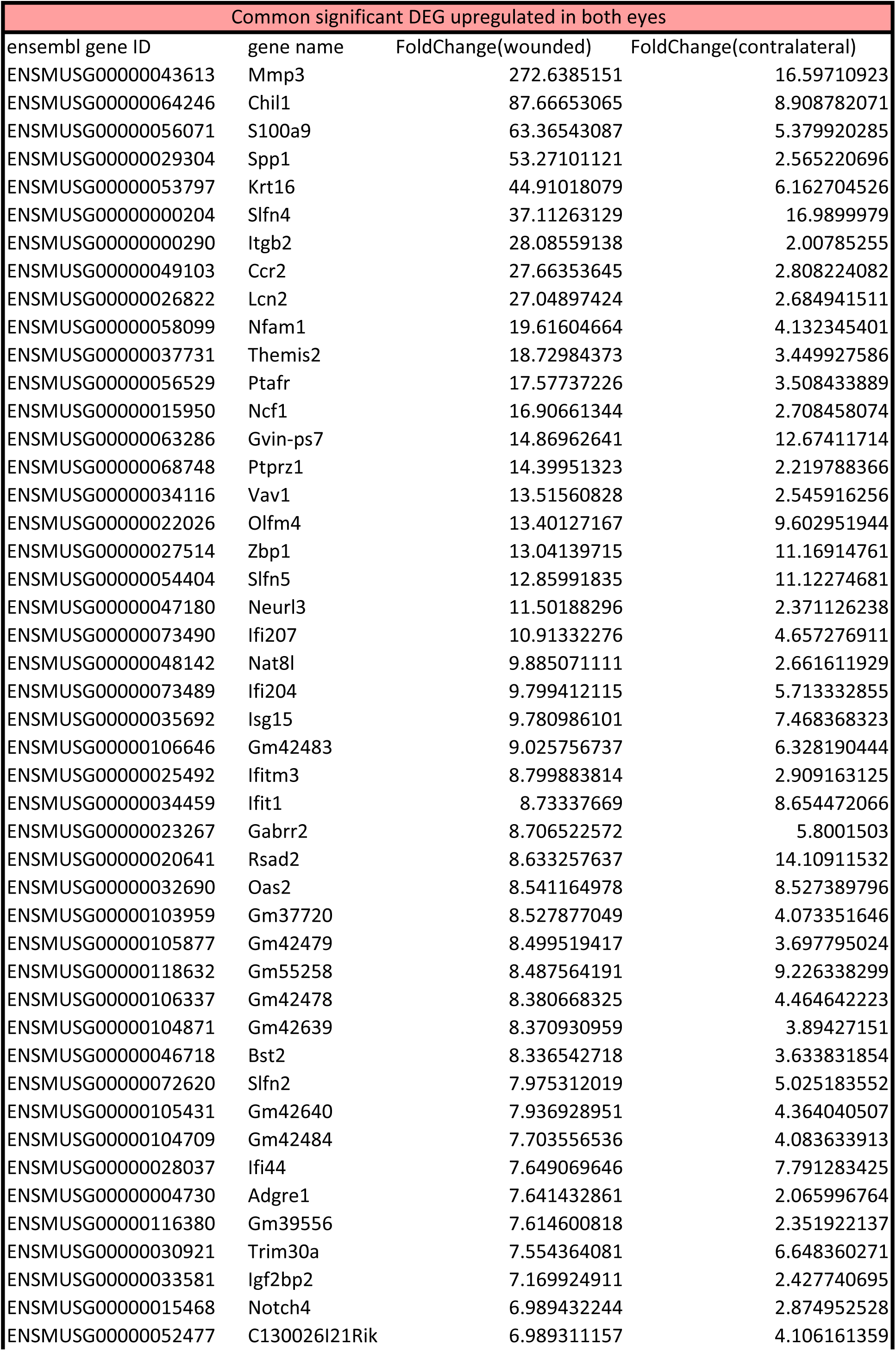
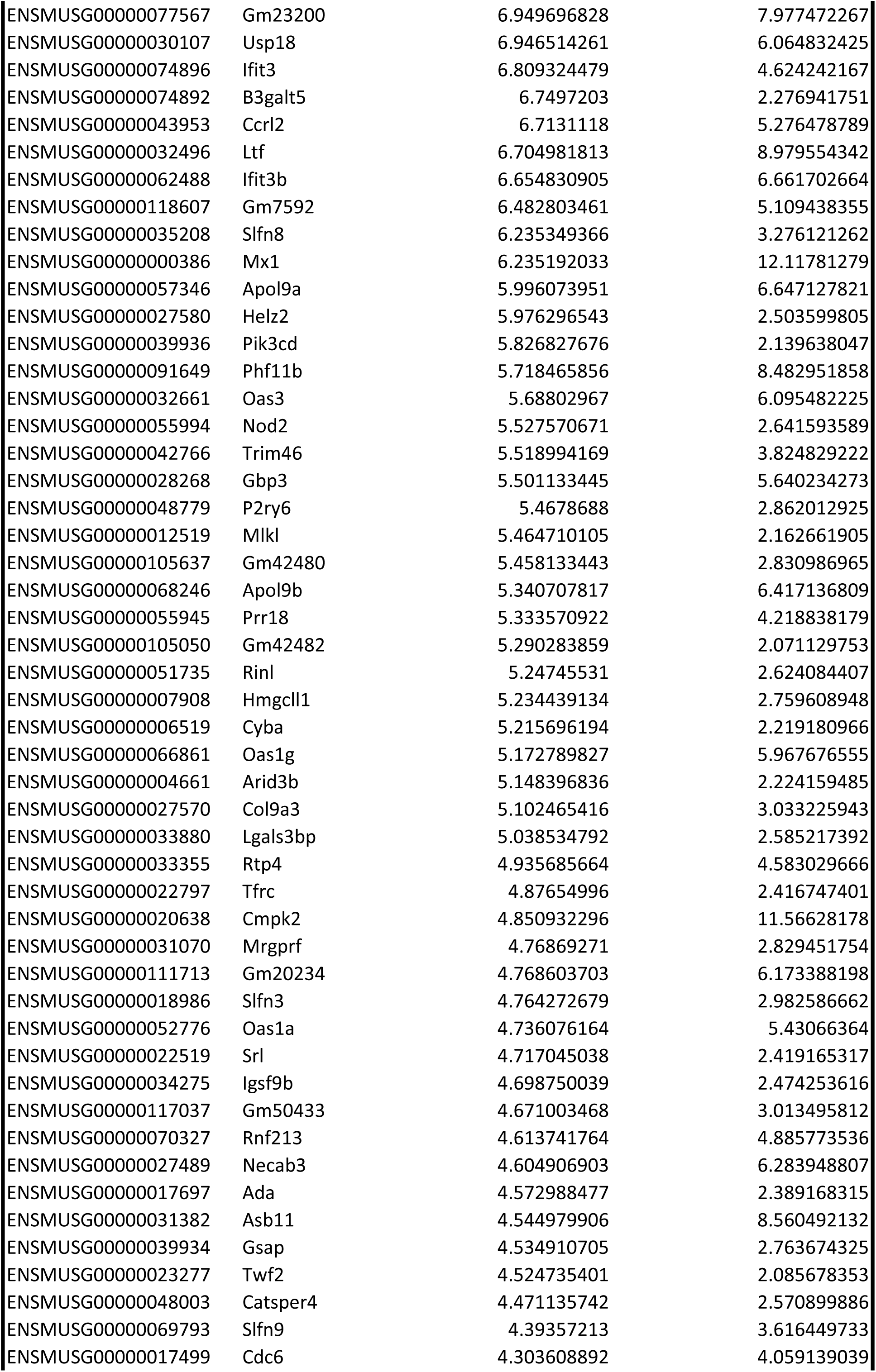
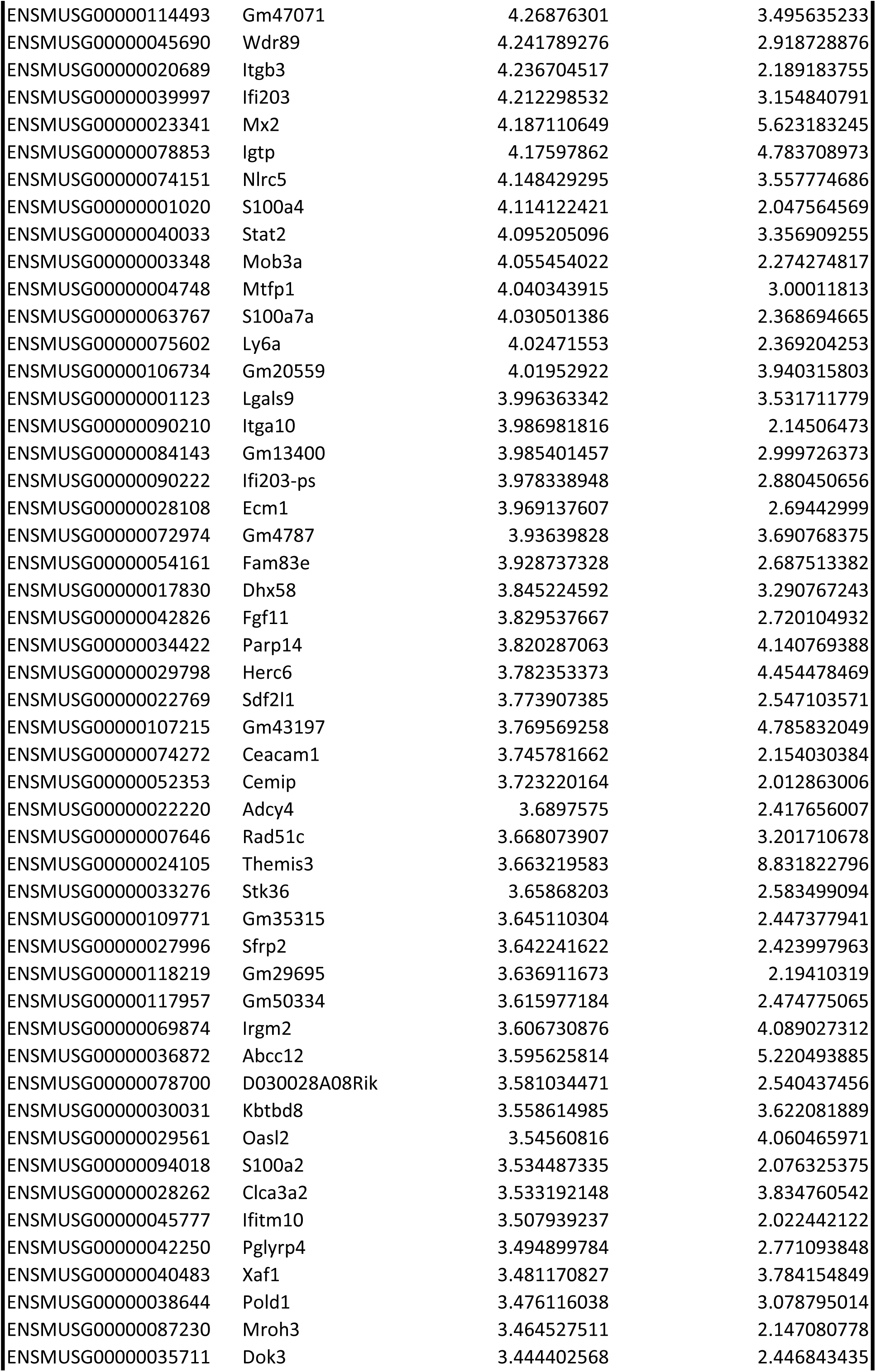
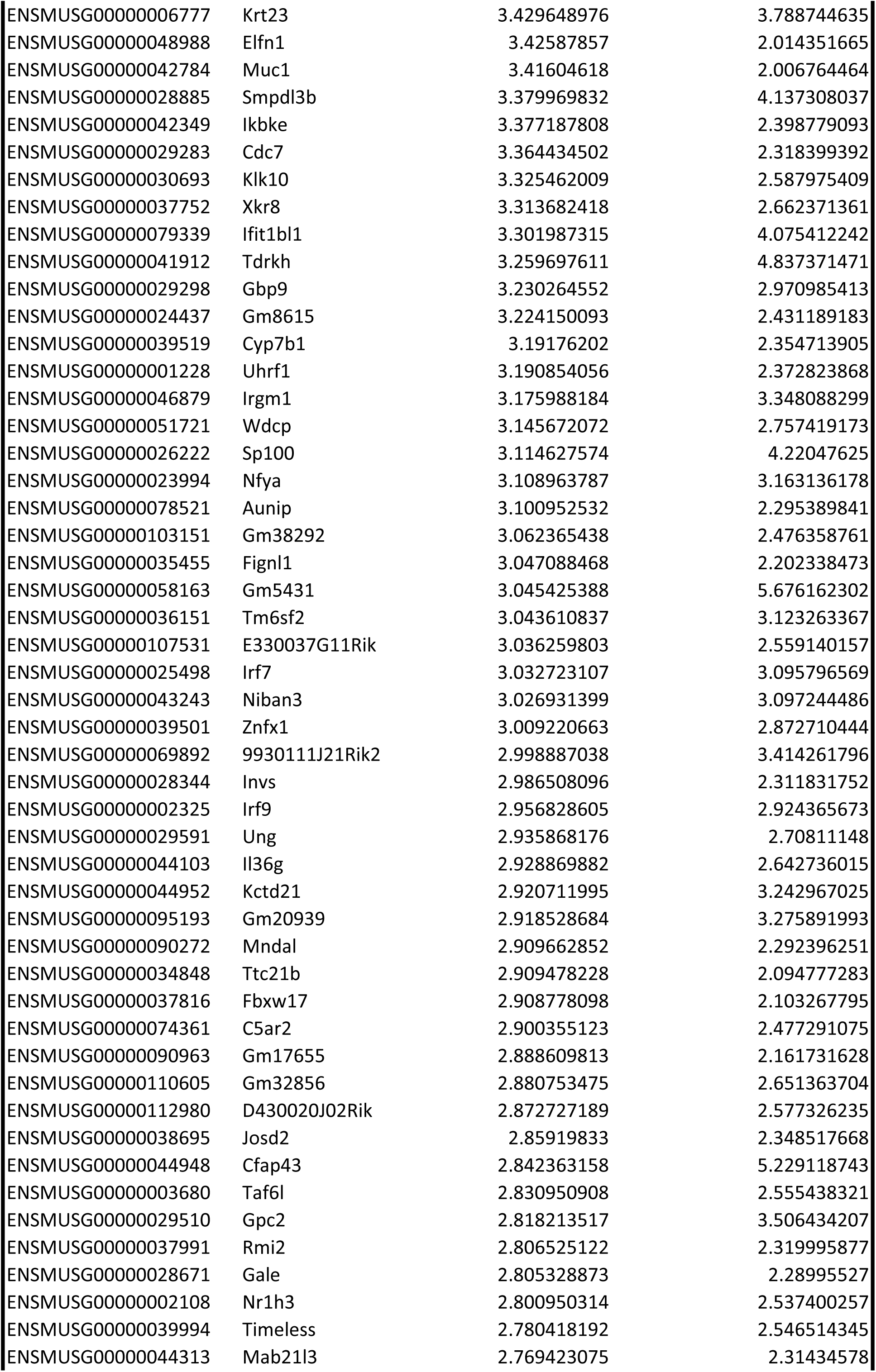
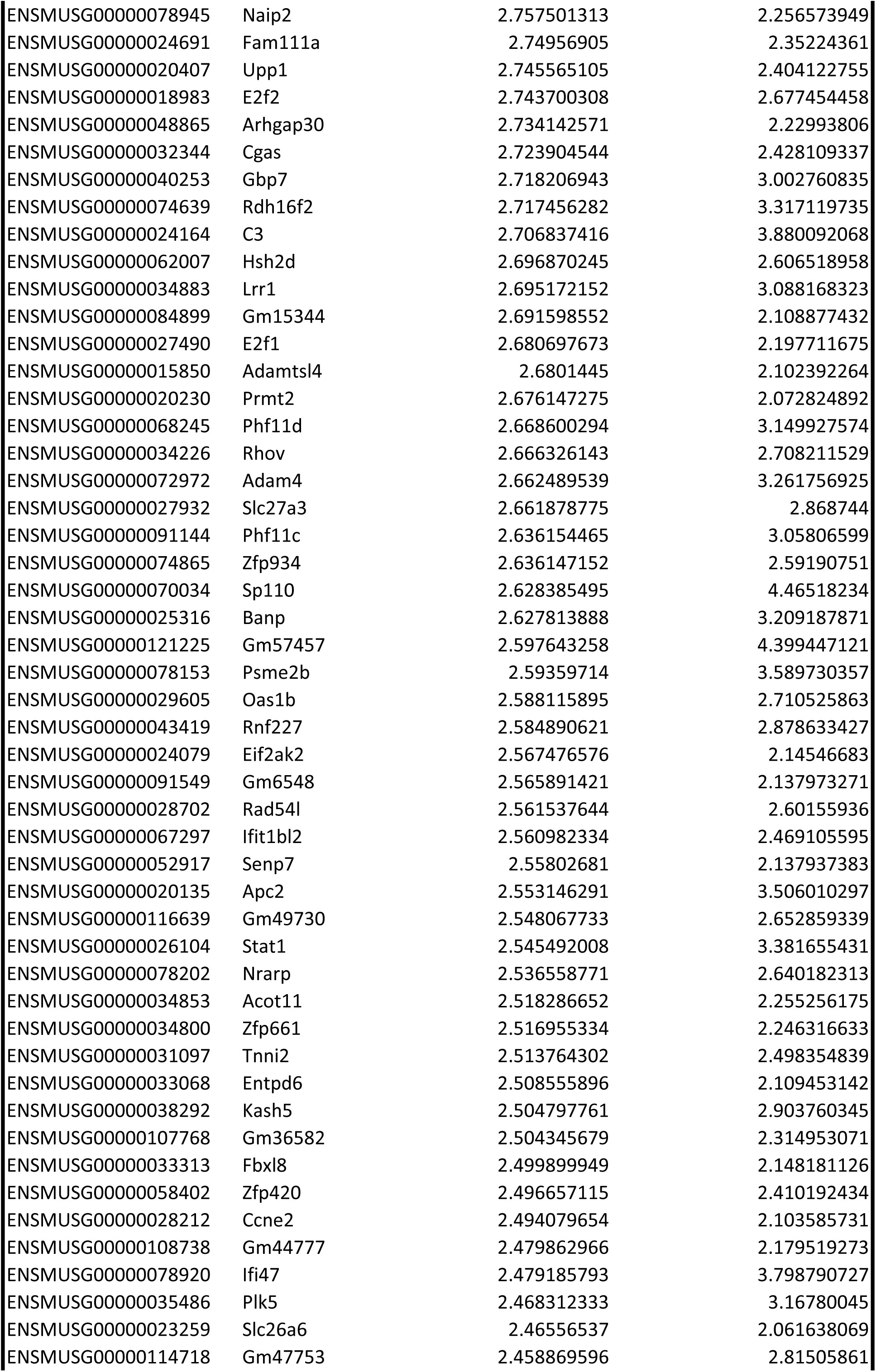
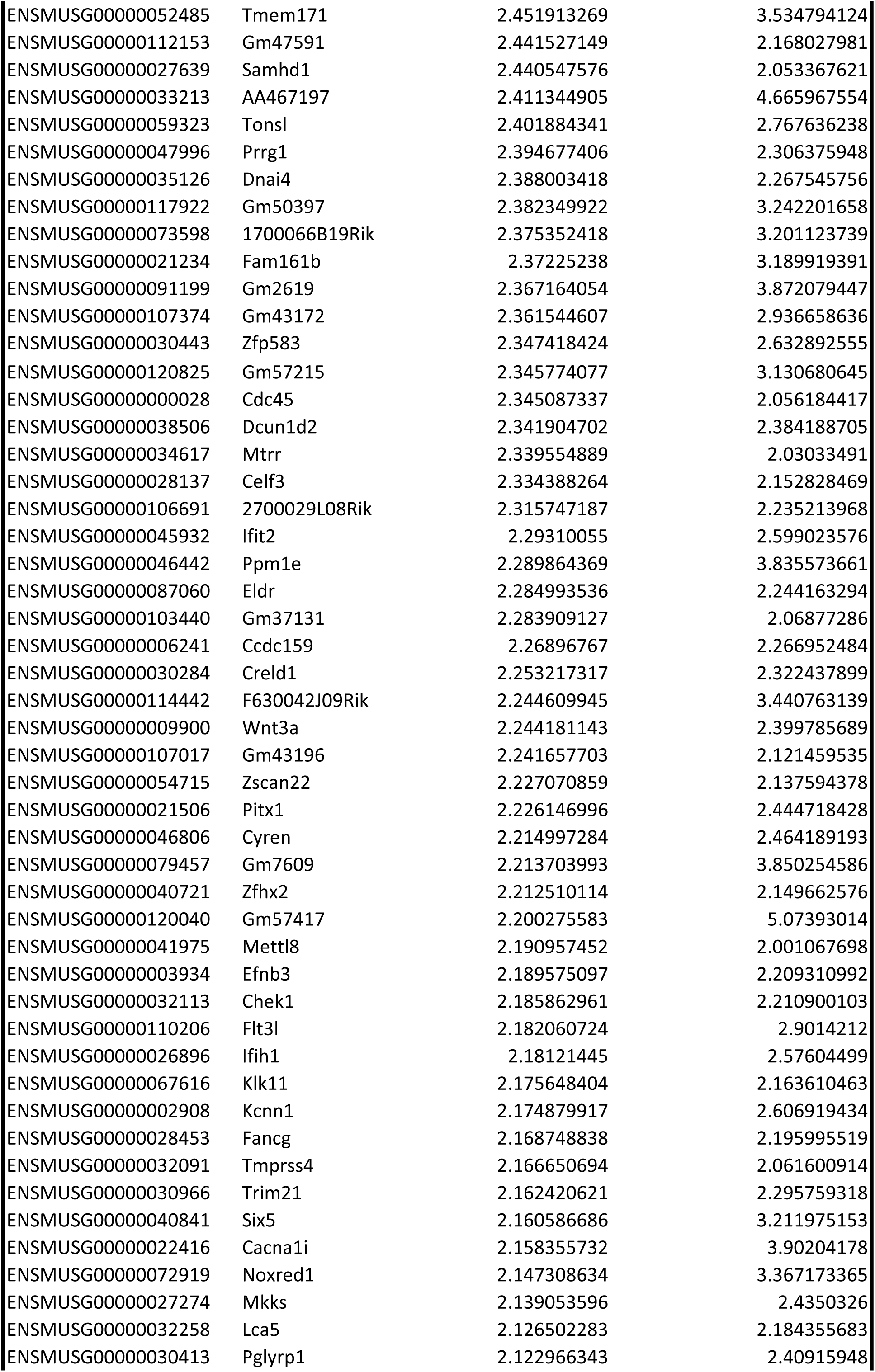
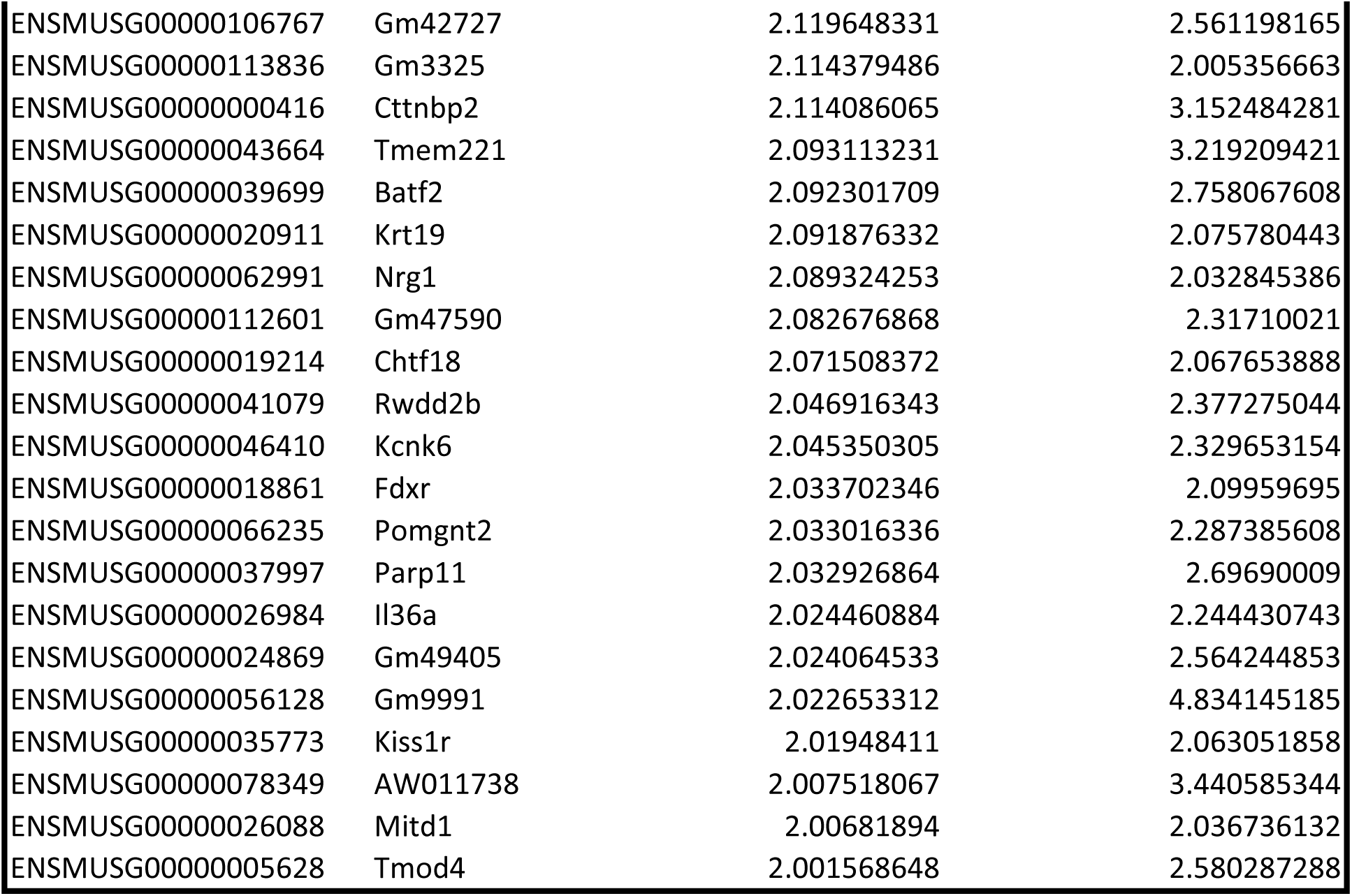
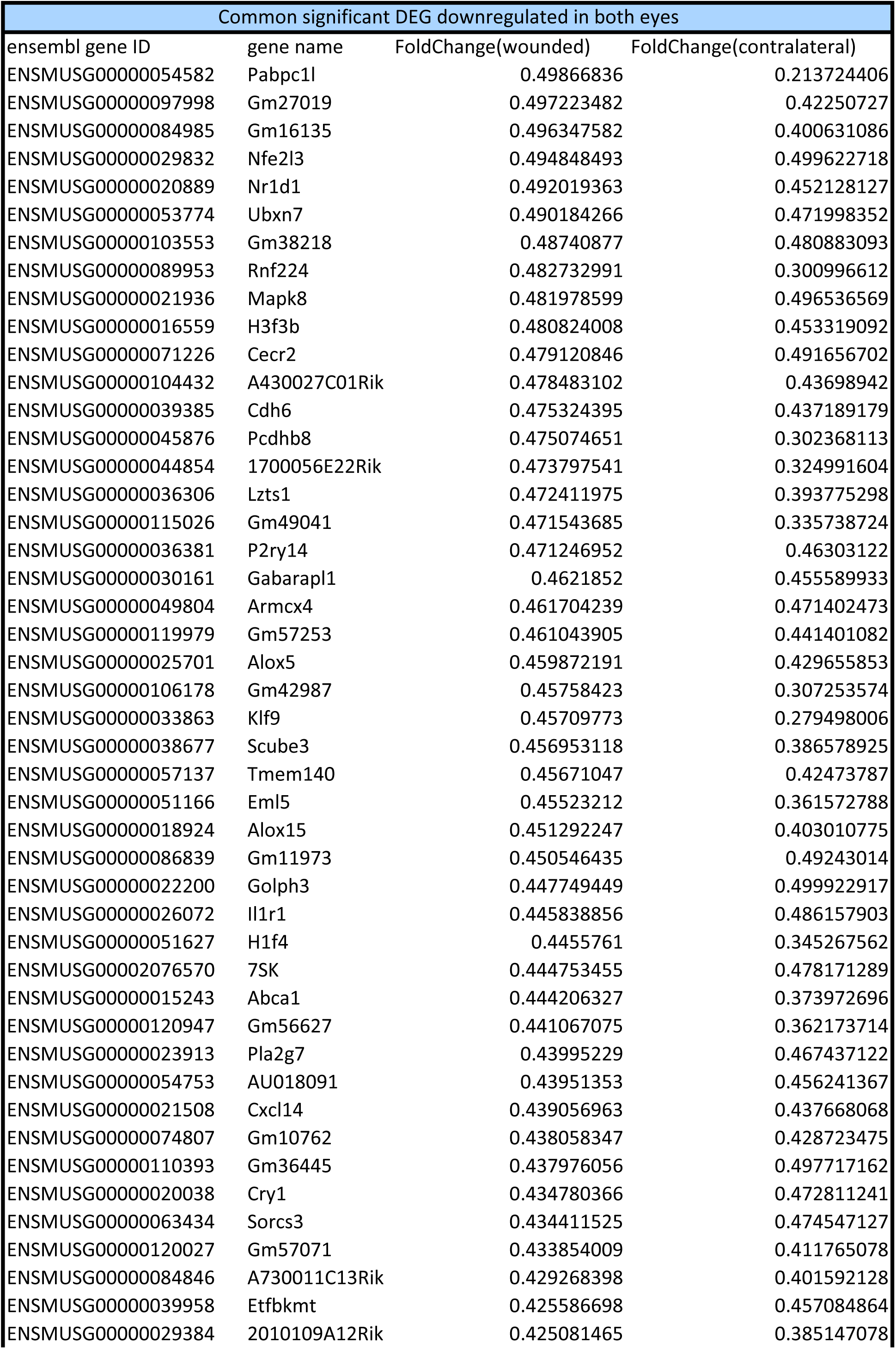
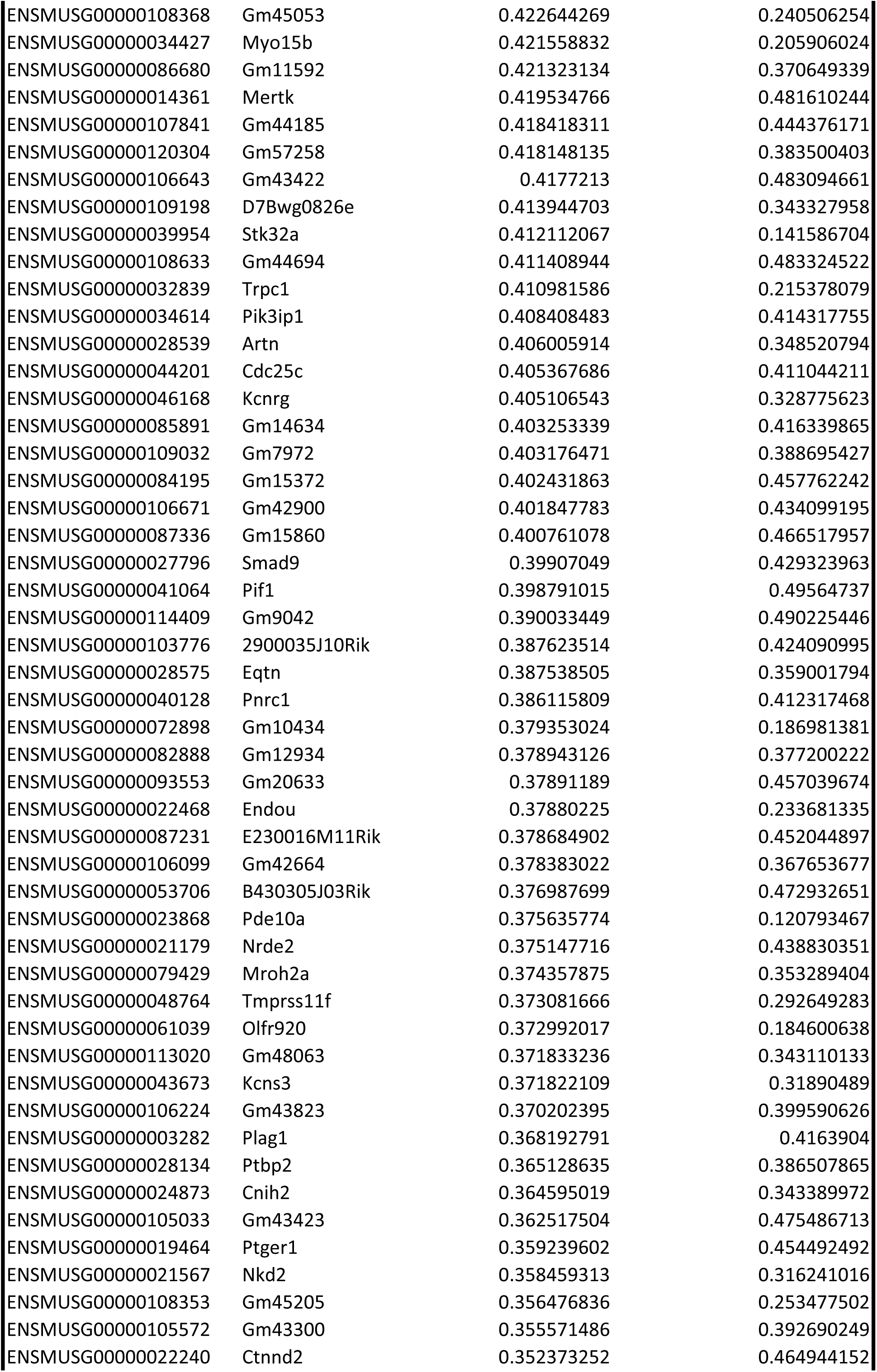
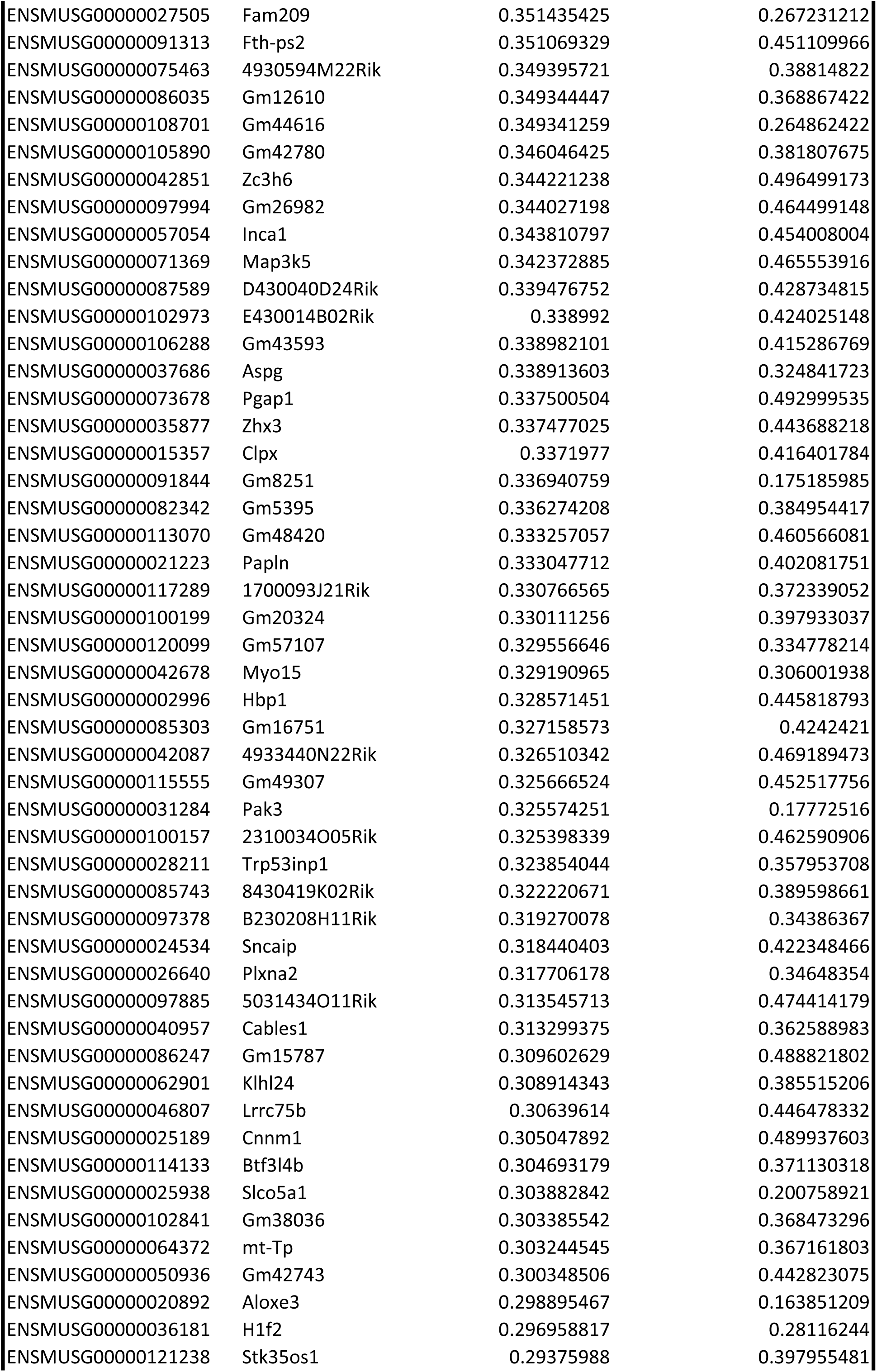
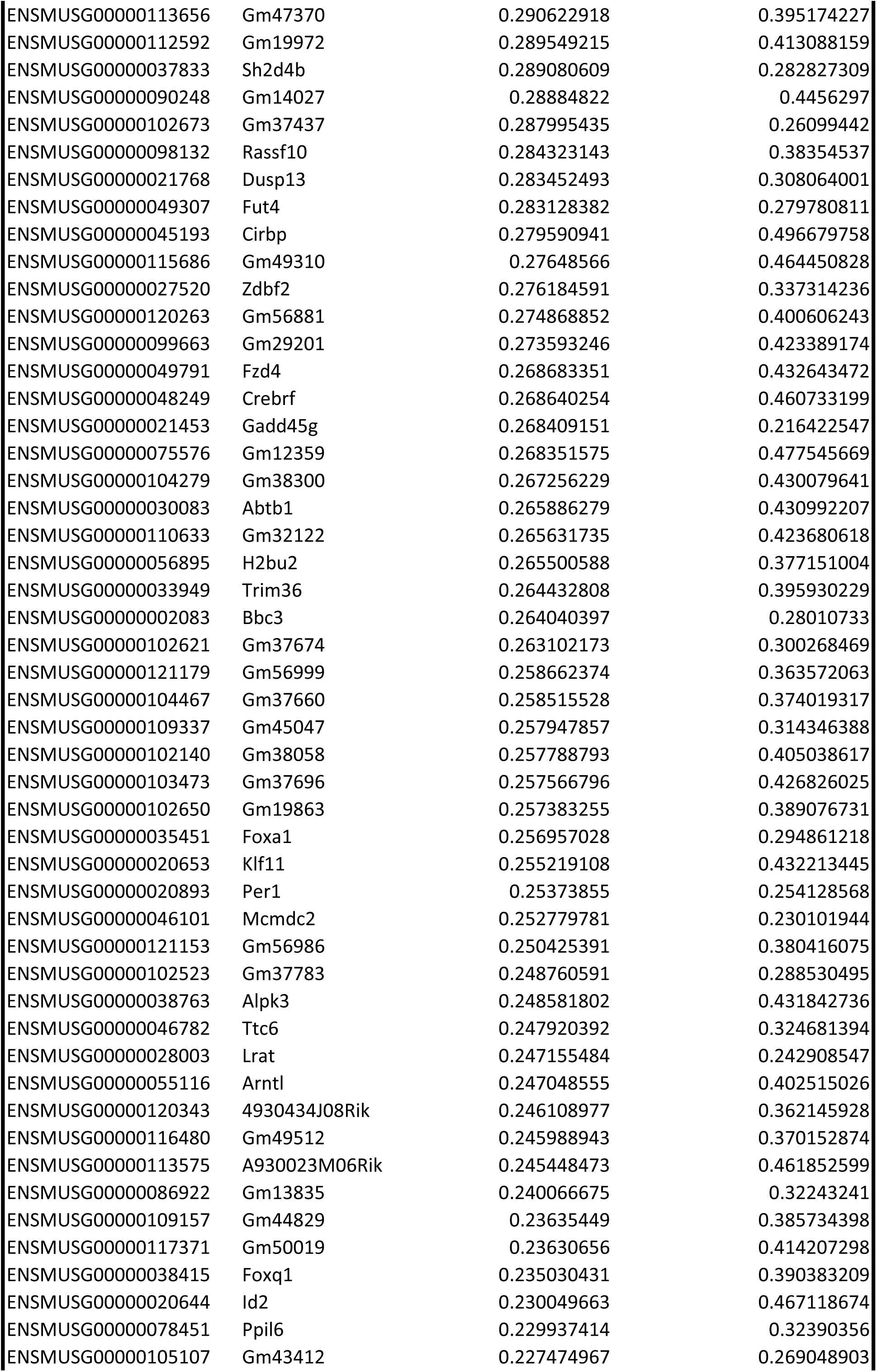
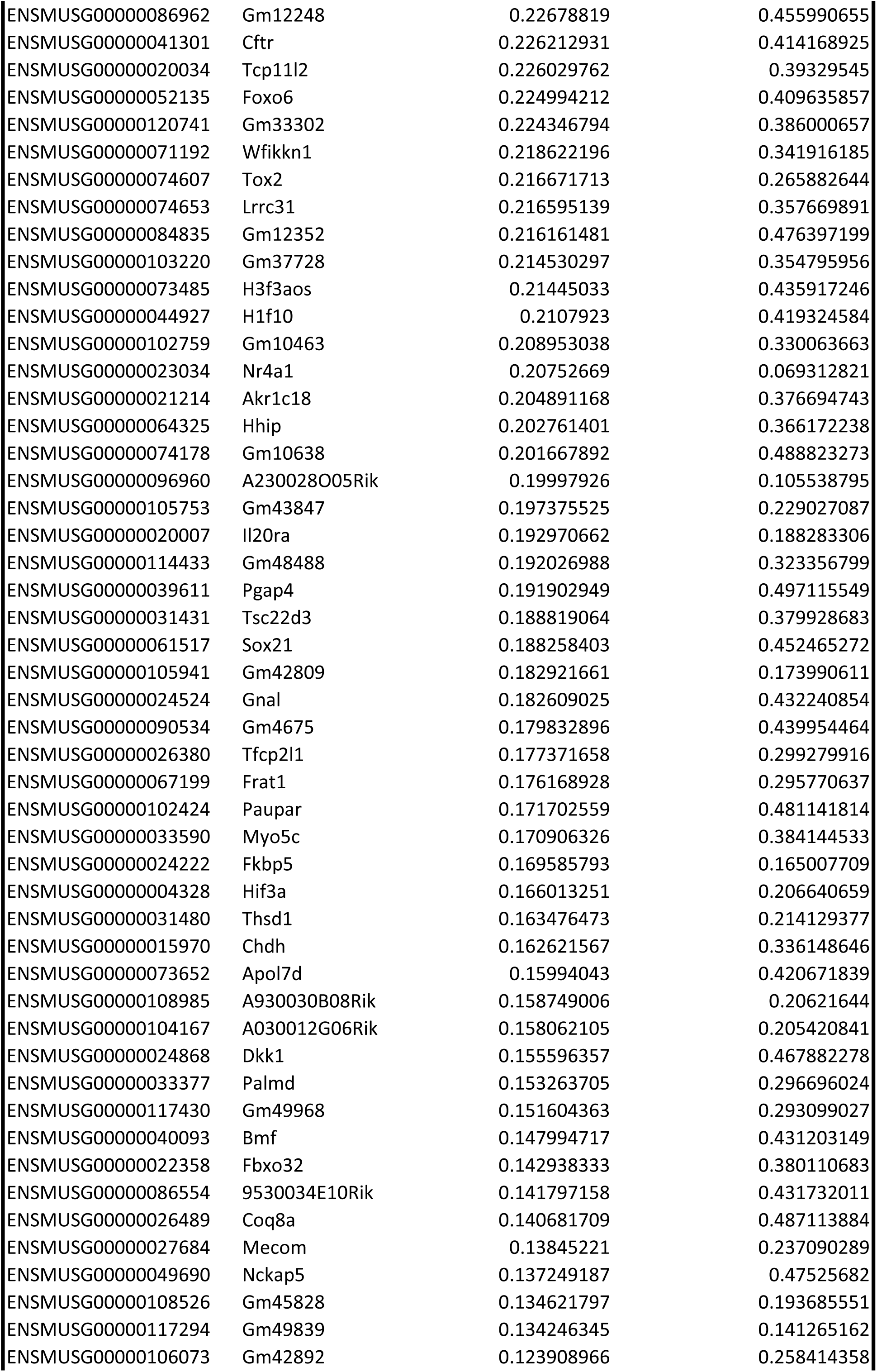
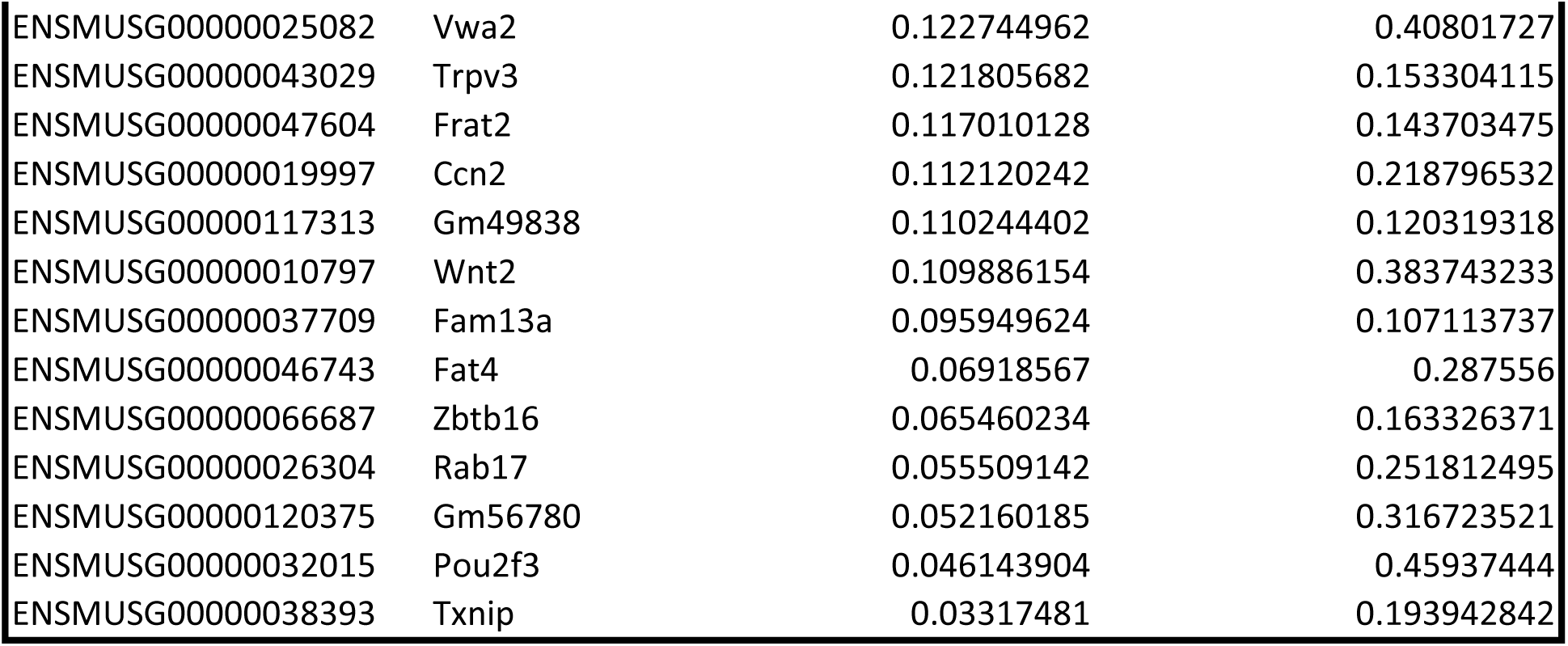
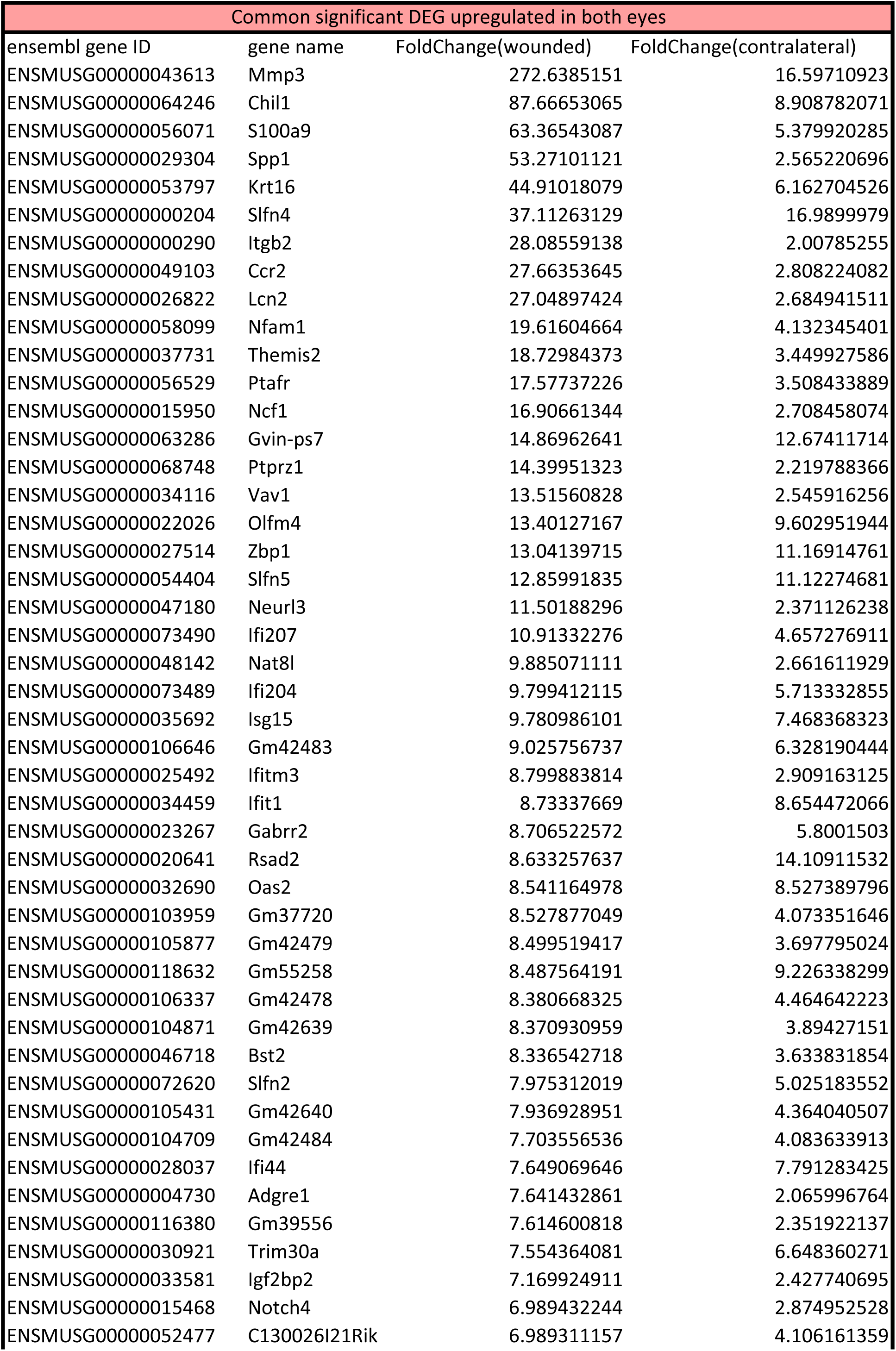
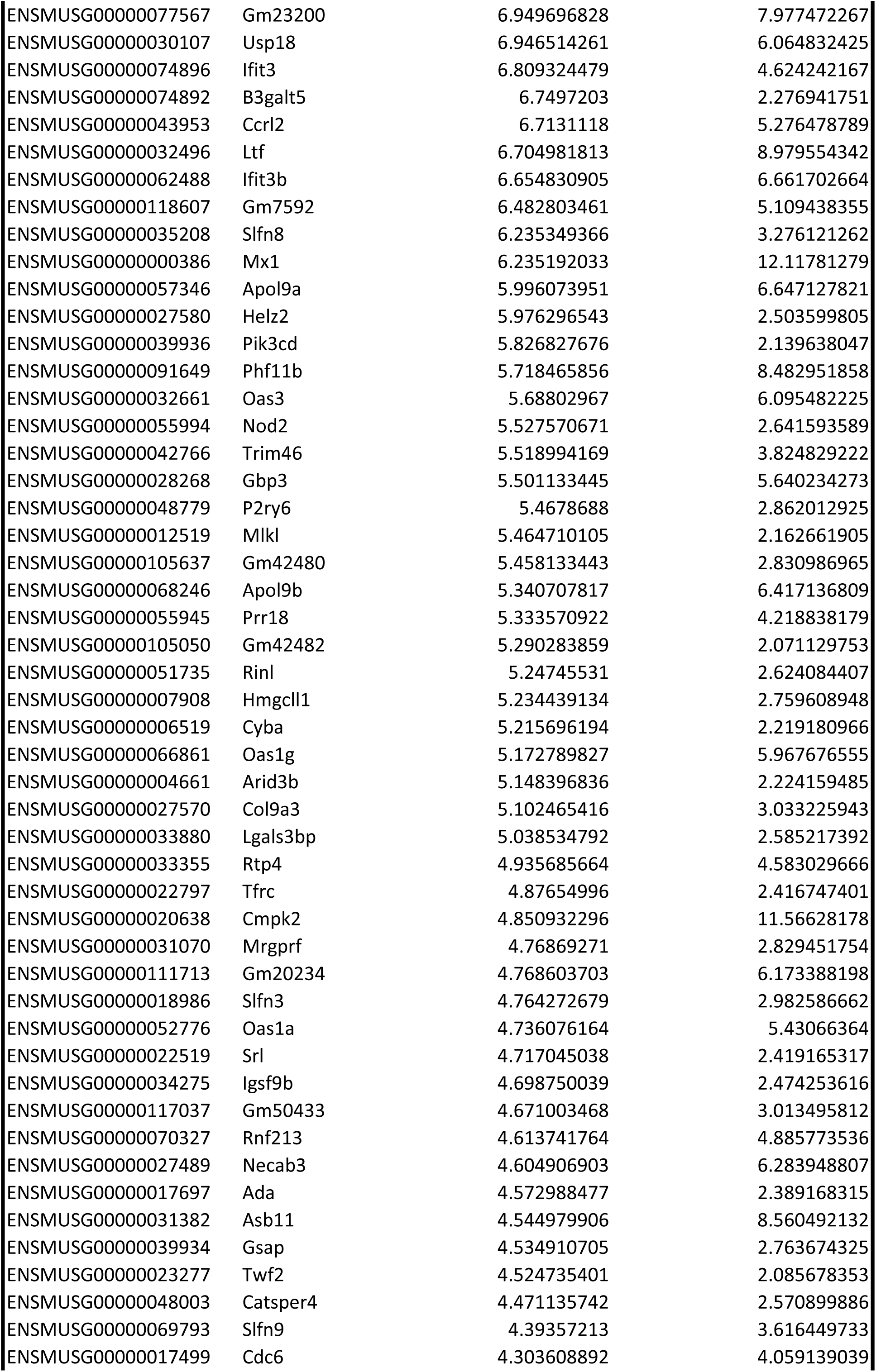
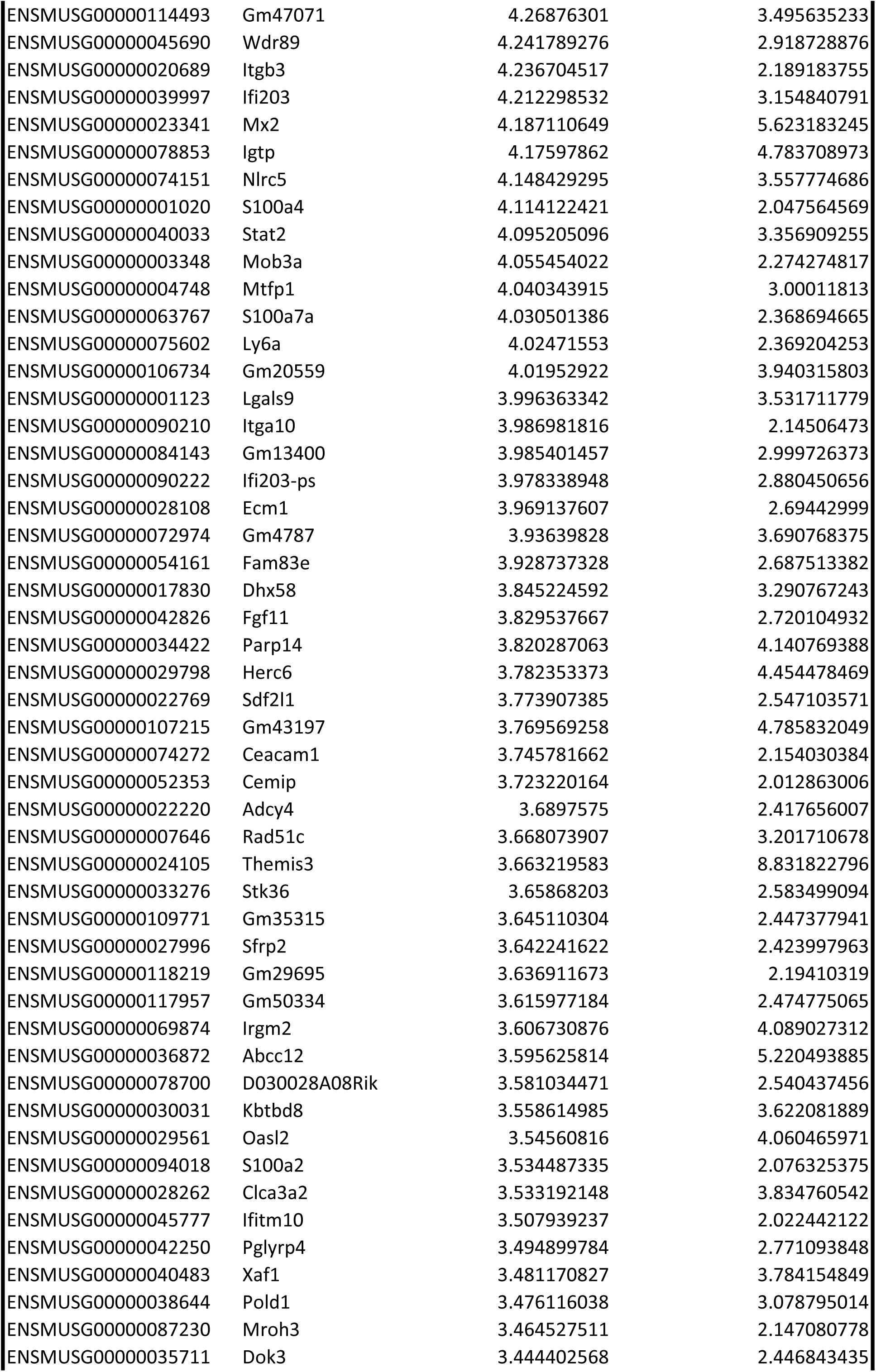
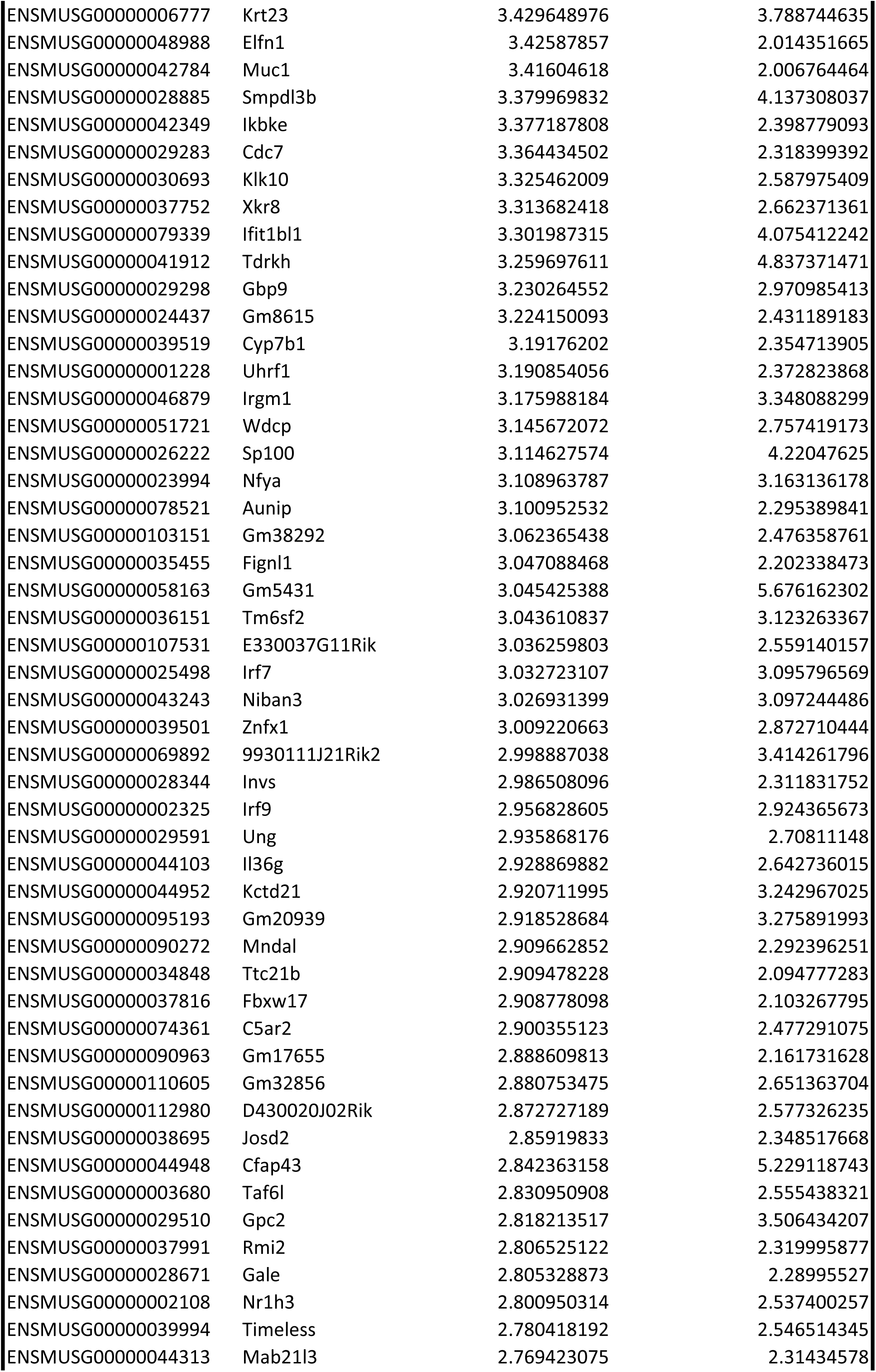
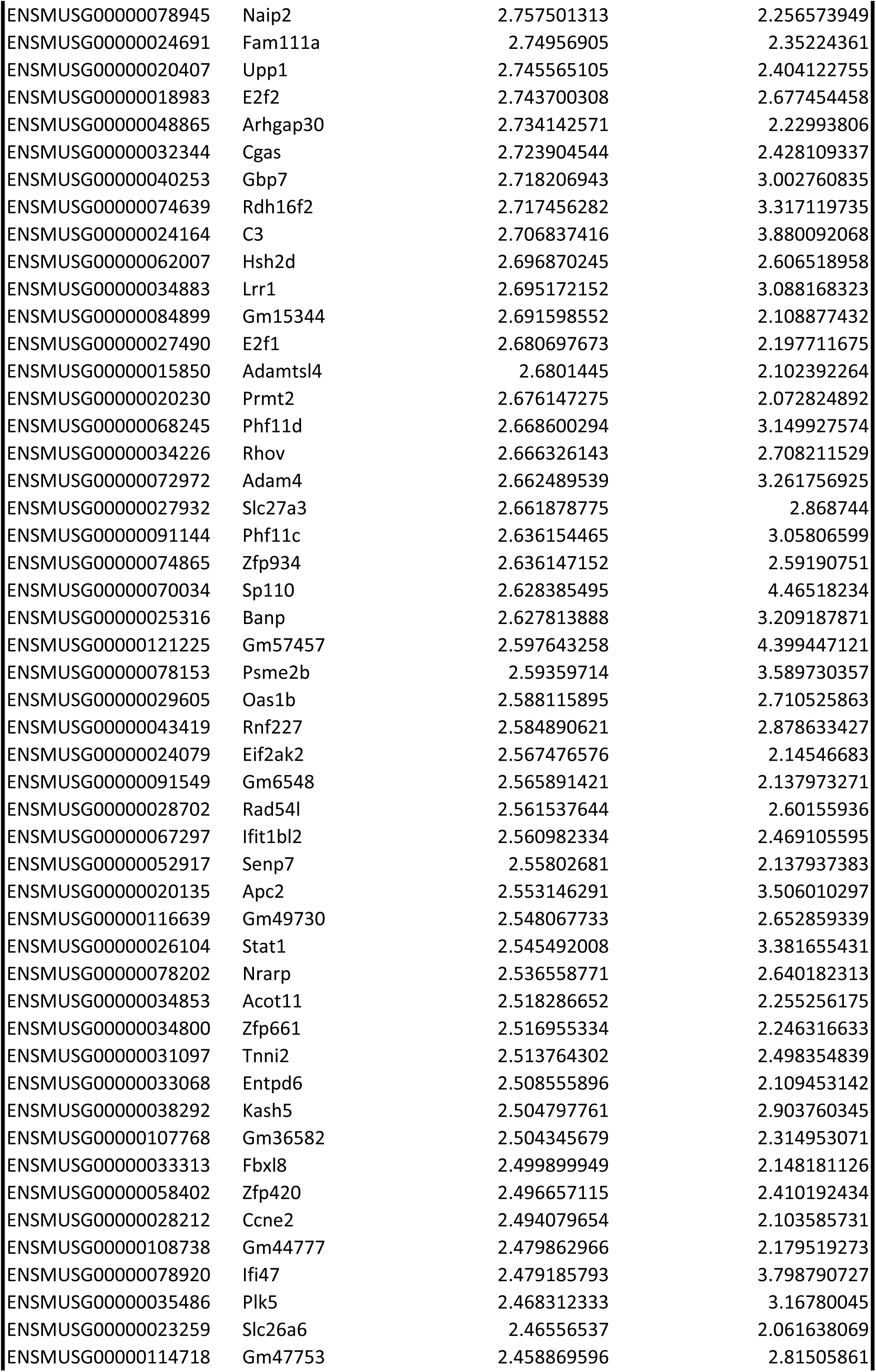
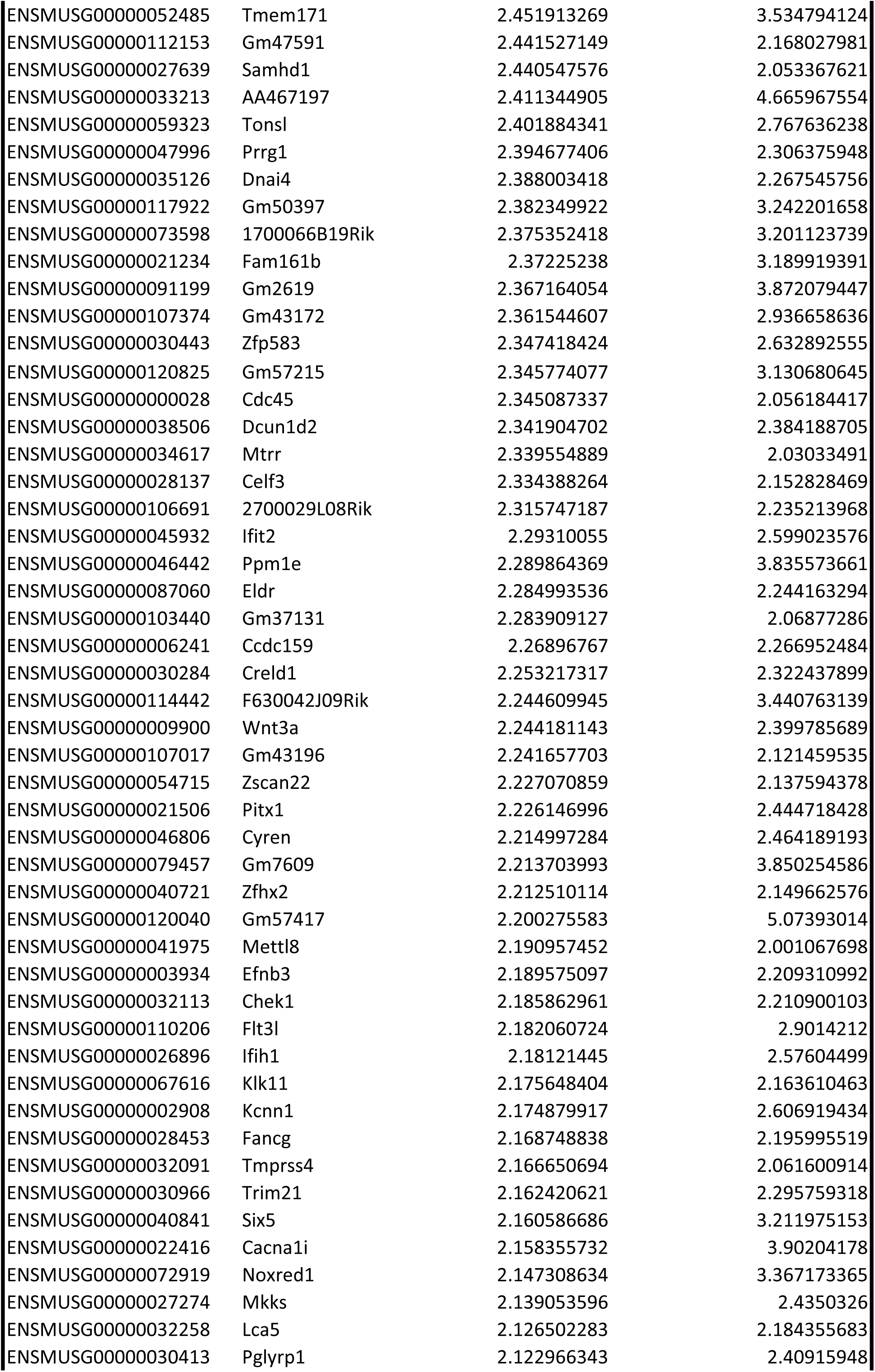
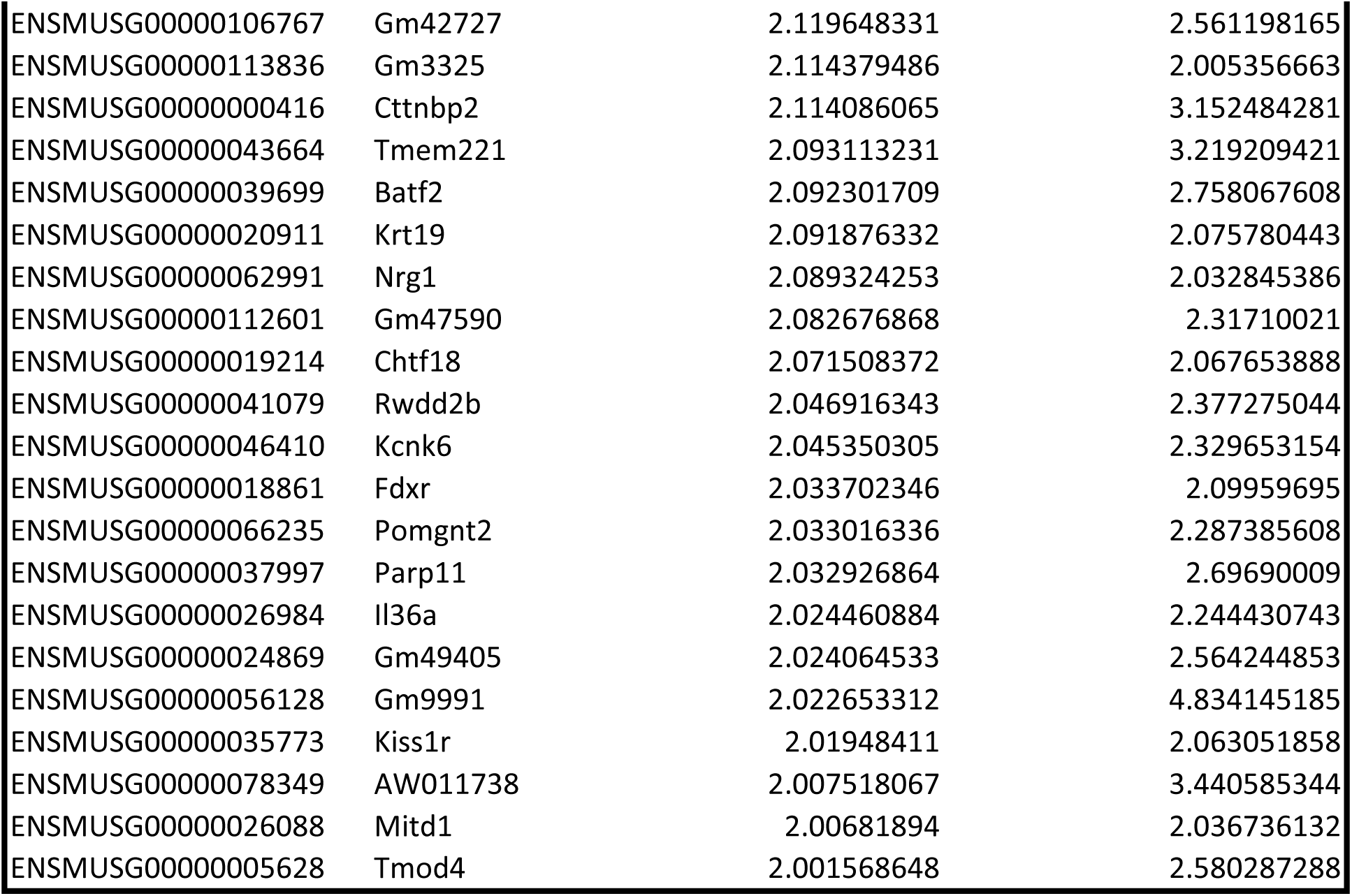
Overview of the common significant DEG found in both eyes. Significant genes with a p-adj <0.05 and a fold change lower than 0.5 (downregulated) or higher than 2 (upregulated) were isolated from RNA-Seq data at 18H post-abrasion. For each gene, the ensembl gene ID, the associated gene name and the fold changes in both eyes are given.

**Table S6.**
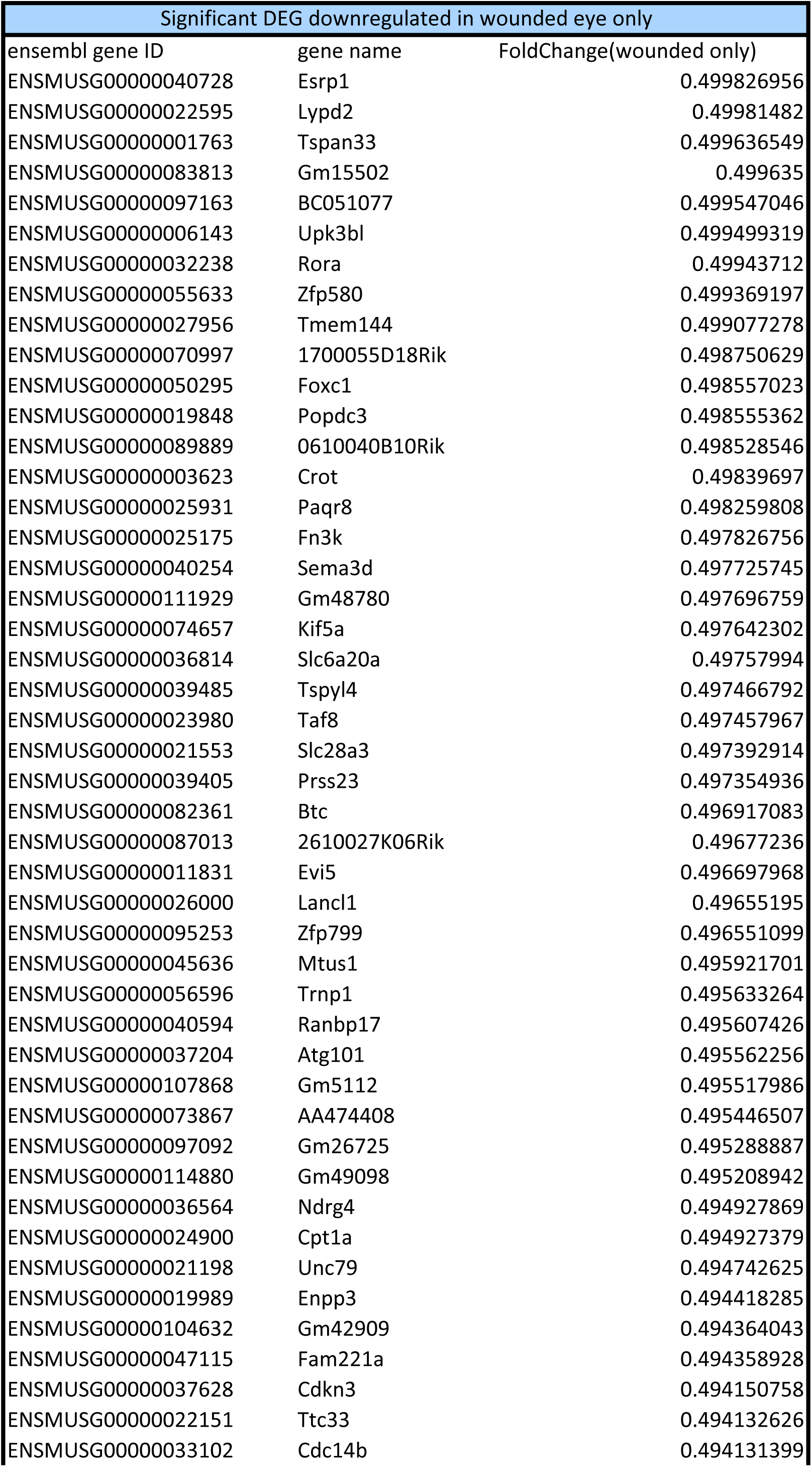
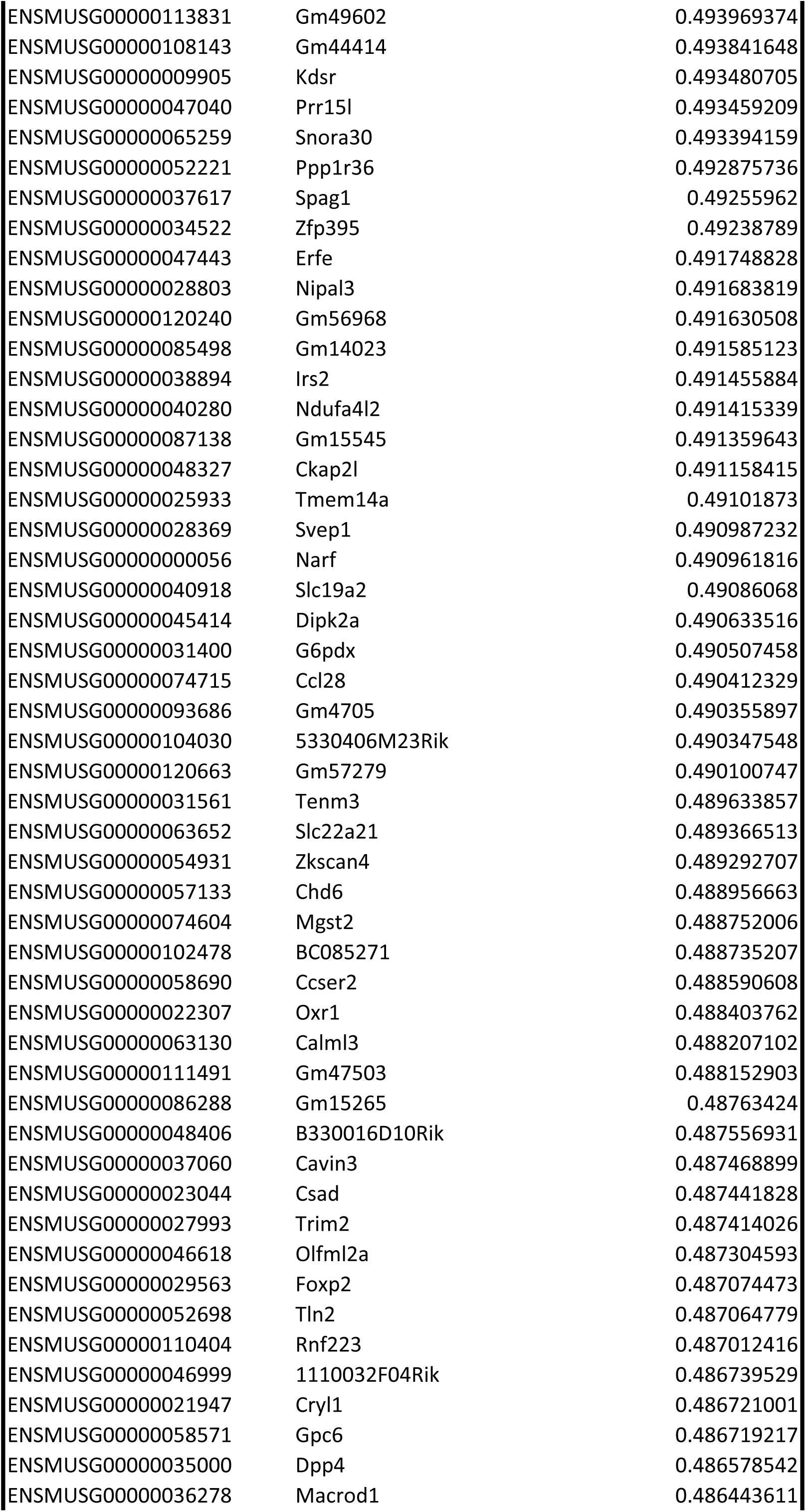
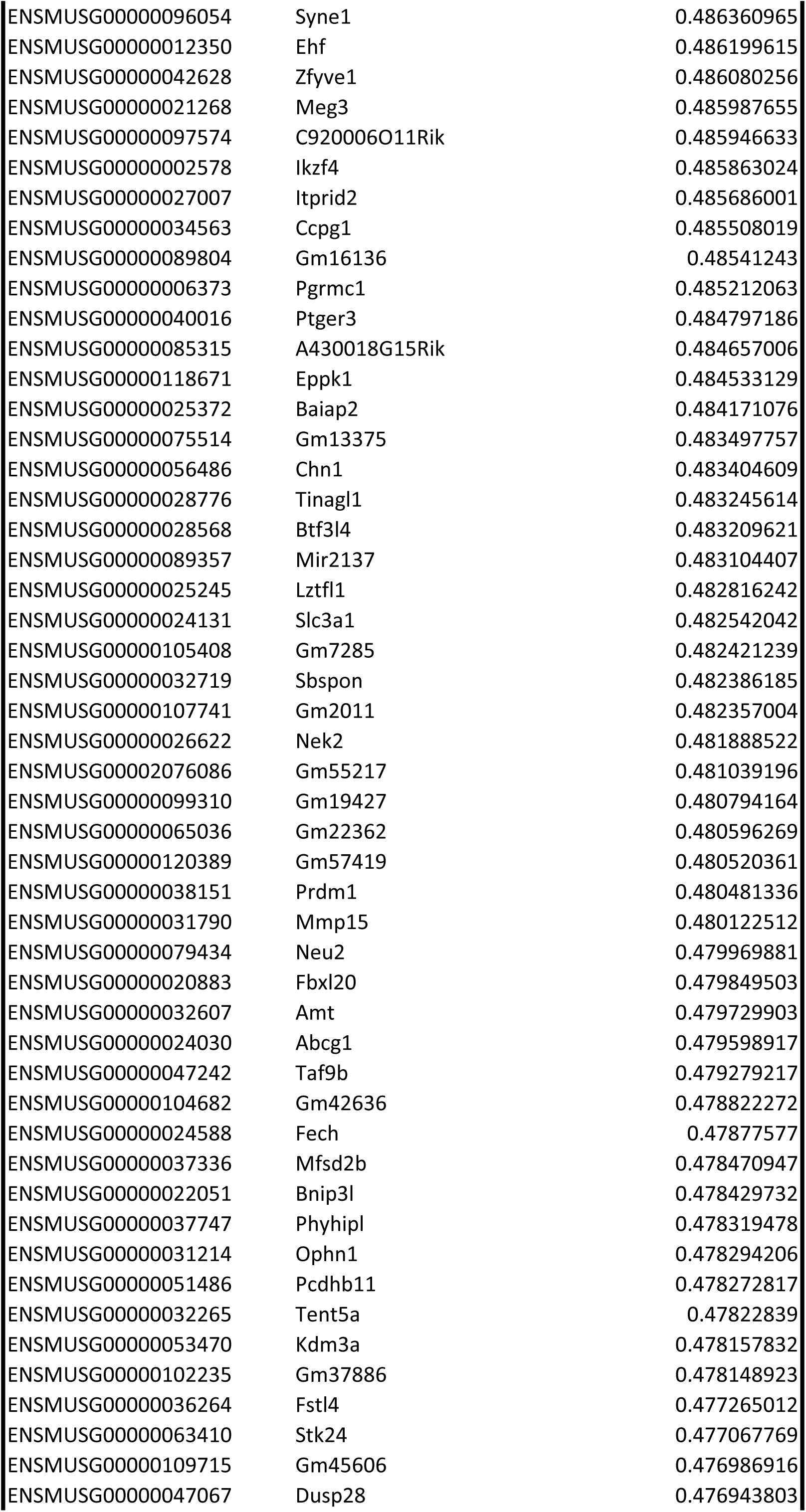
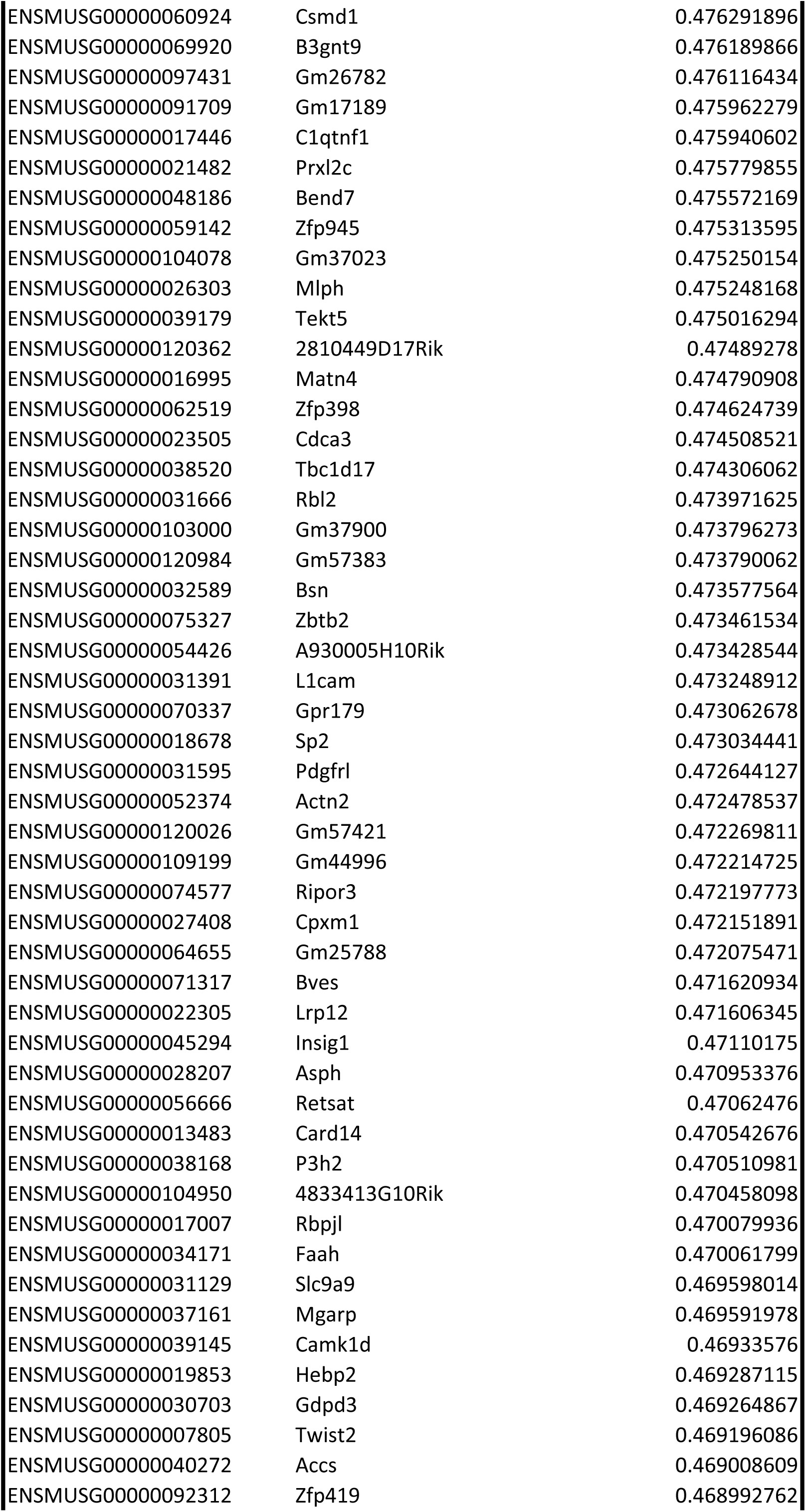
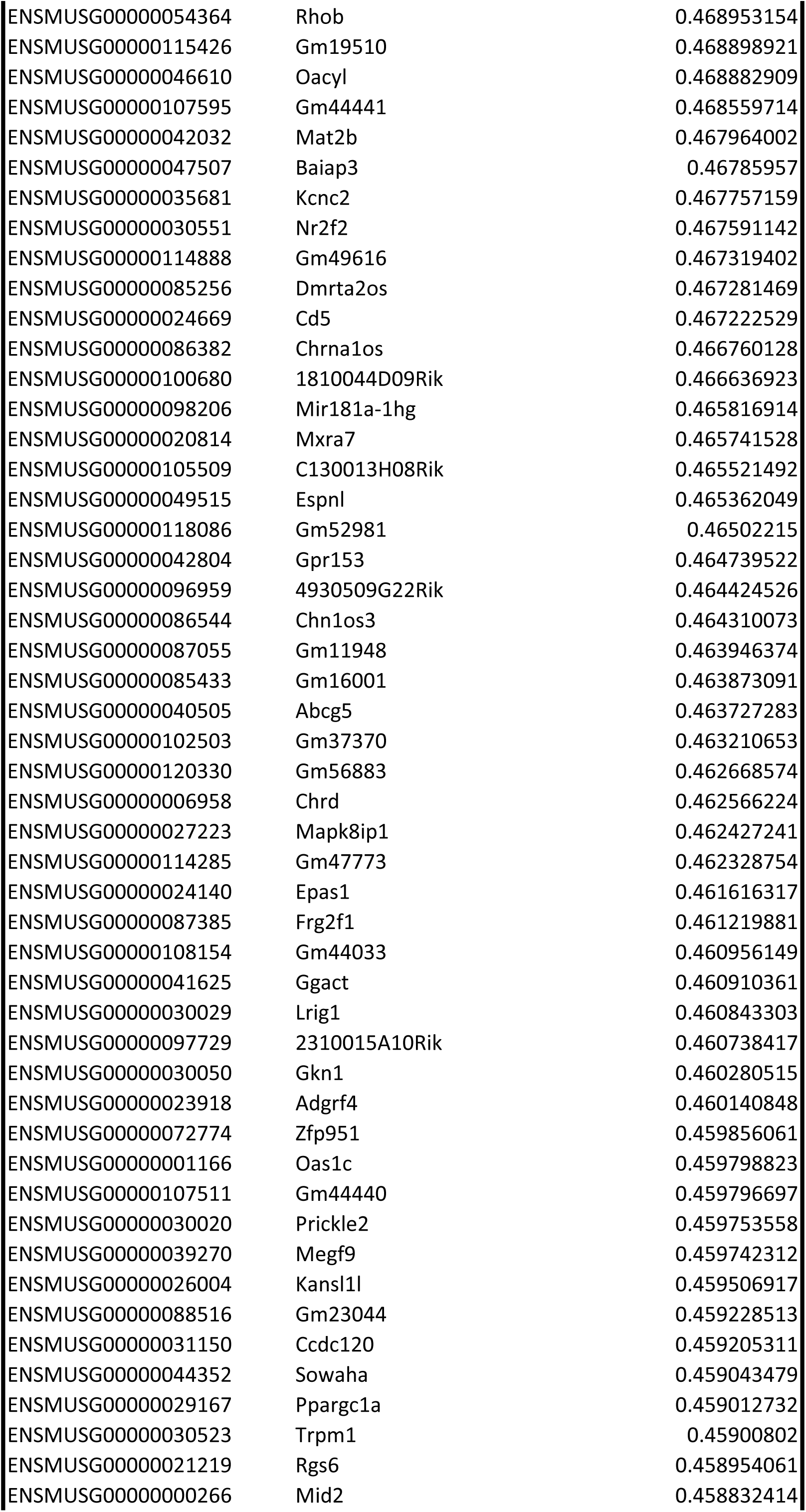
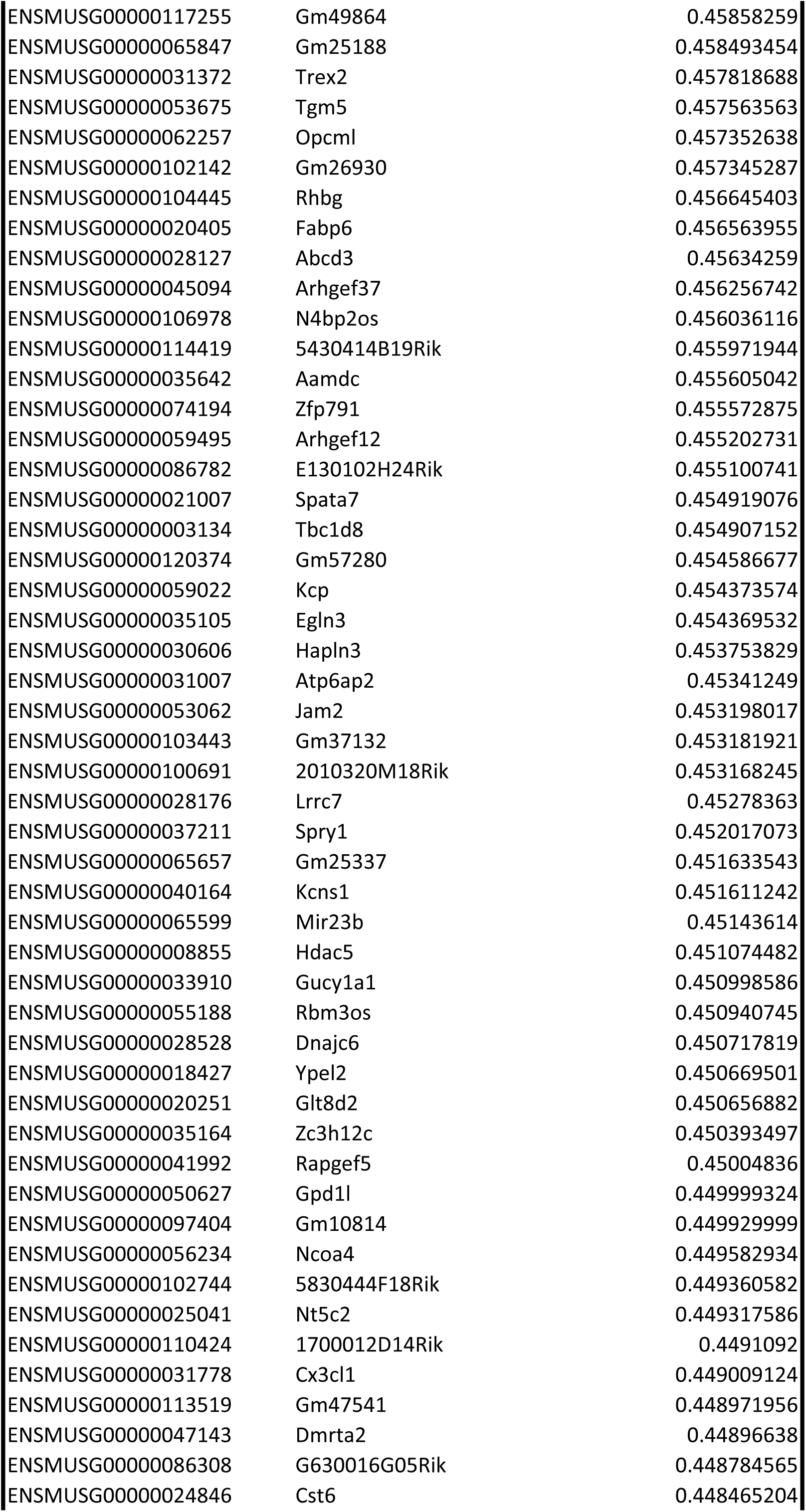
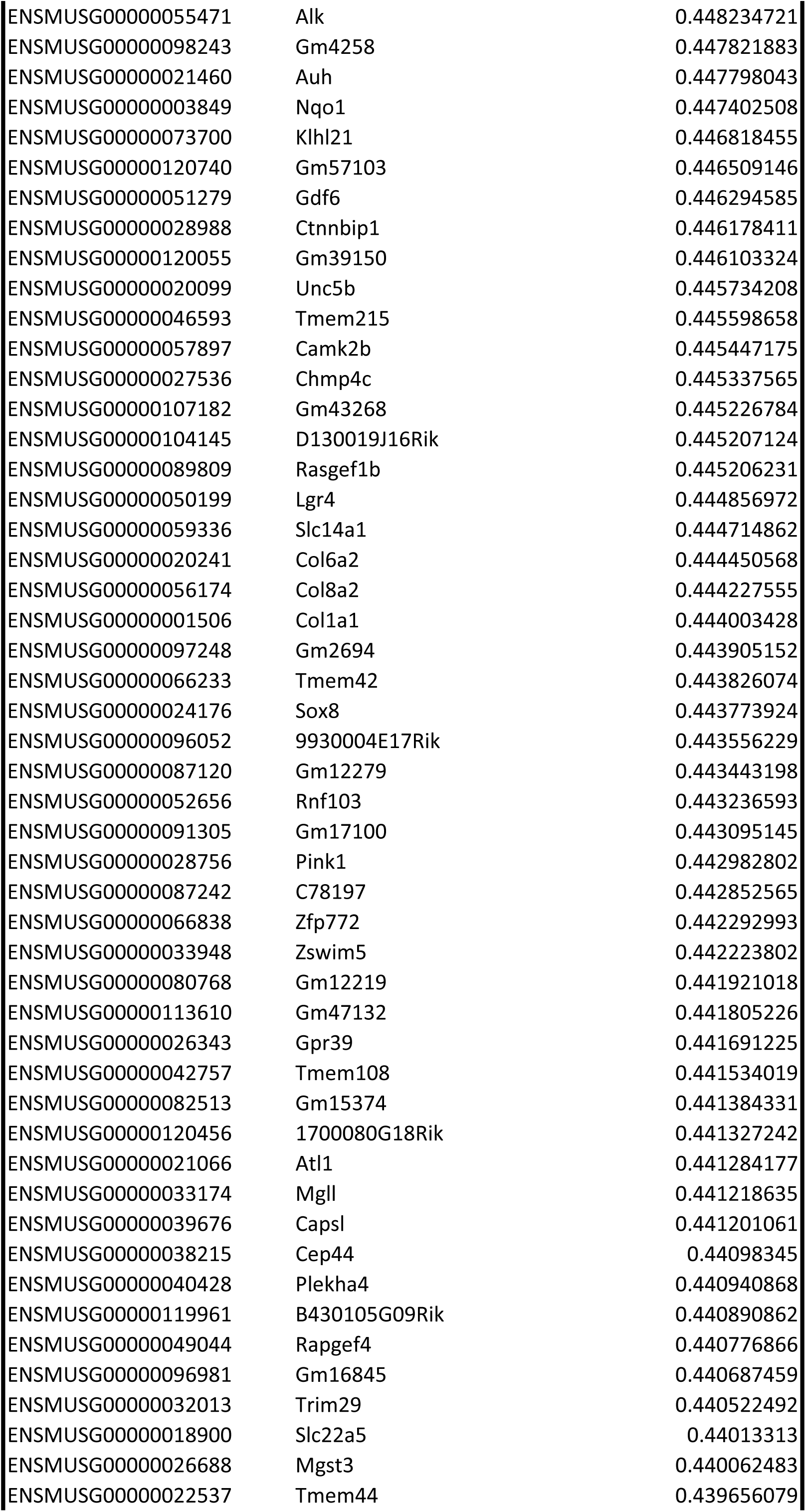
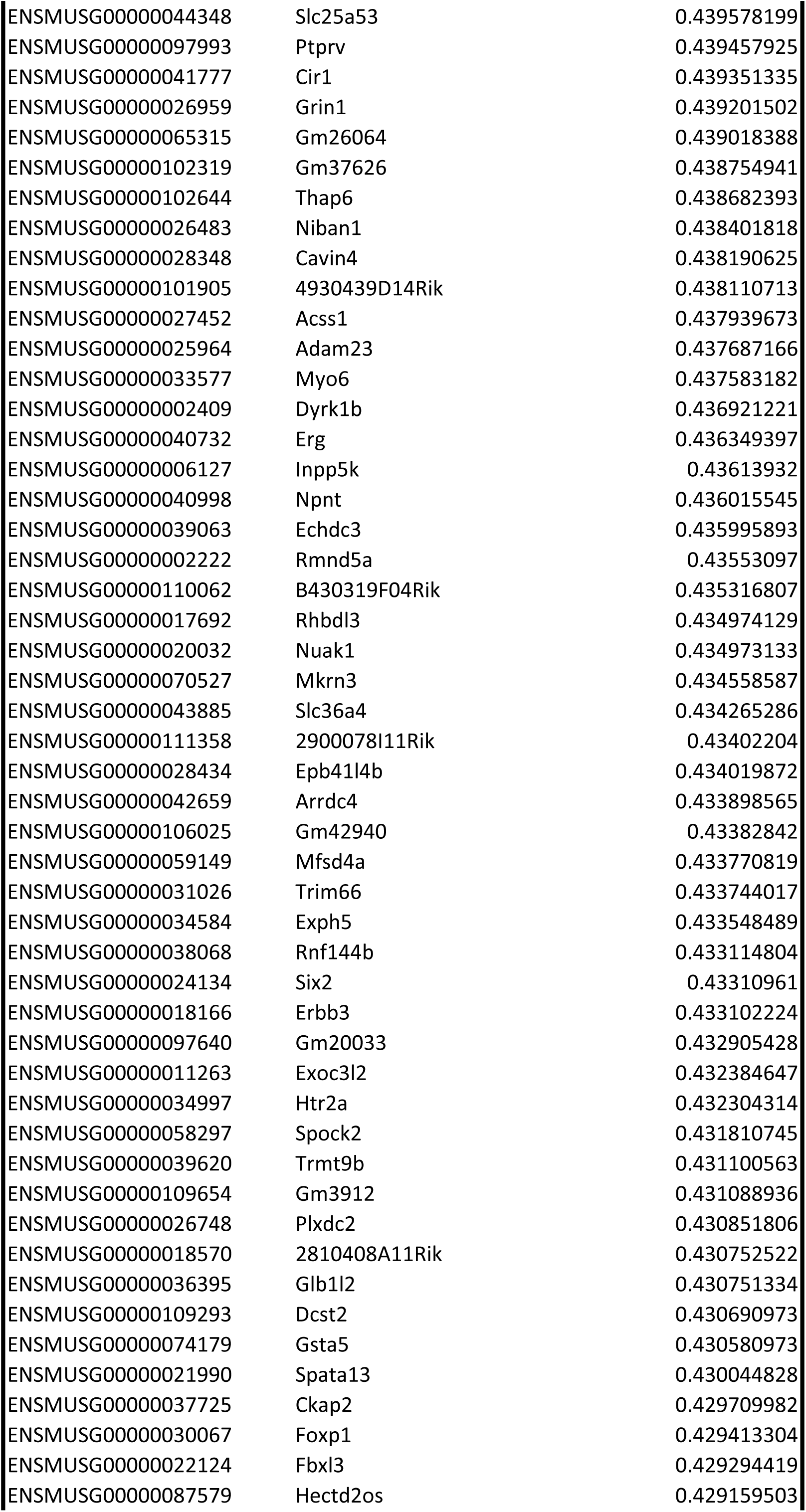
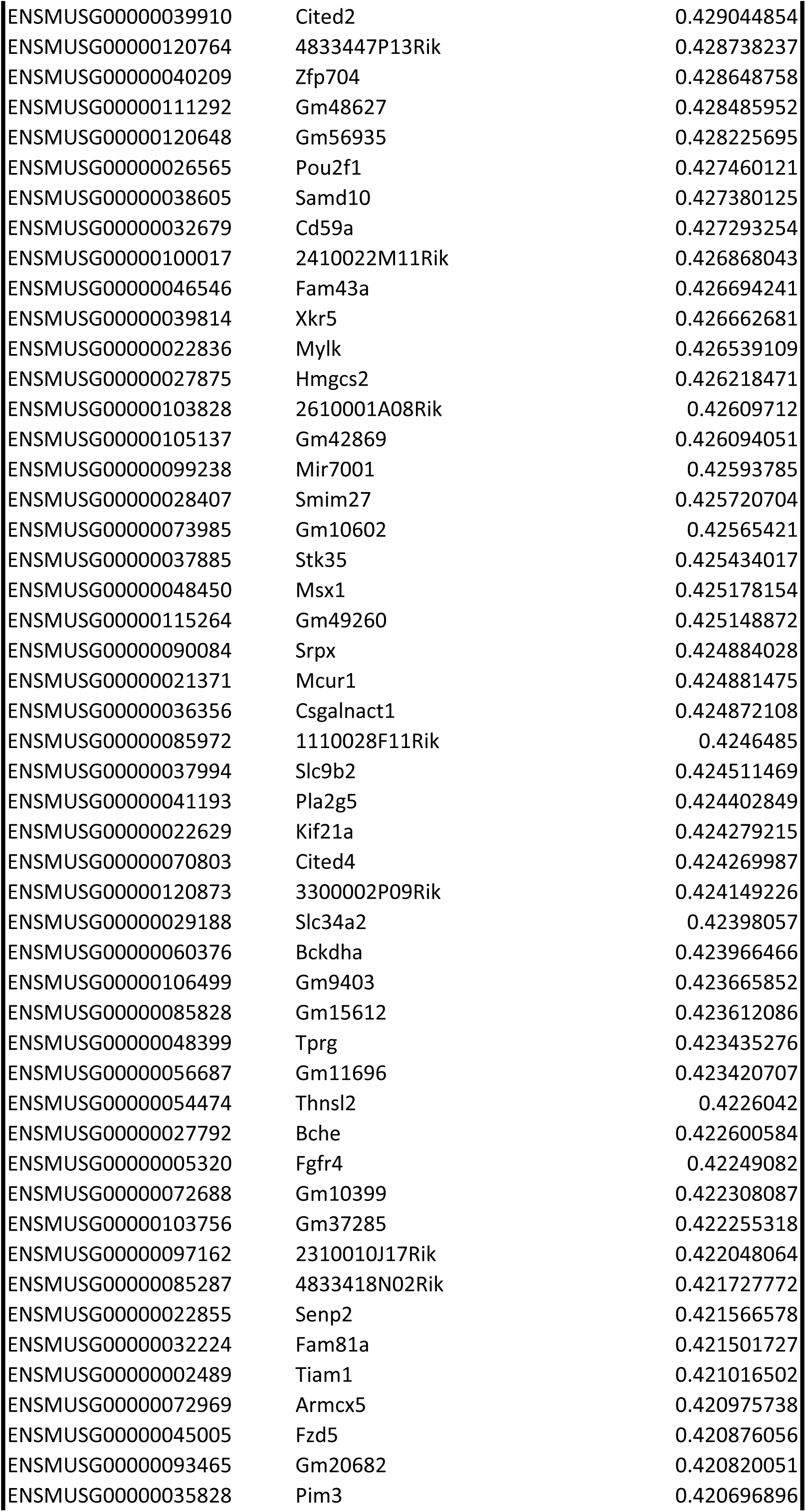
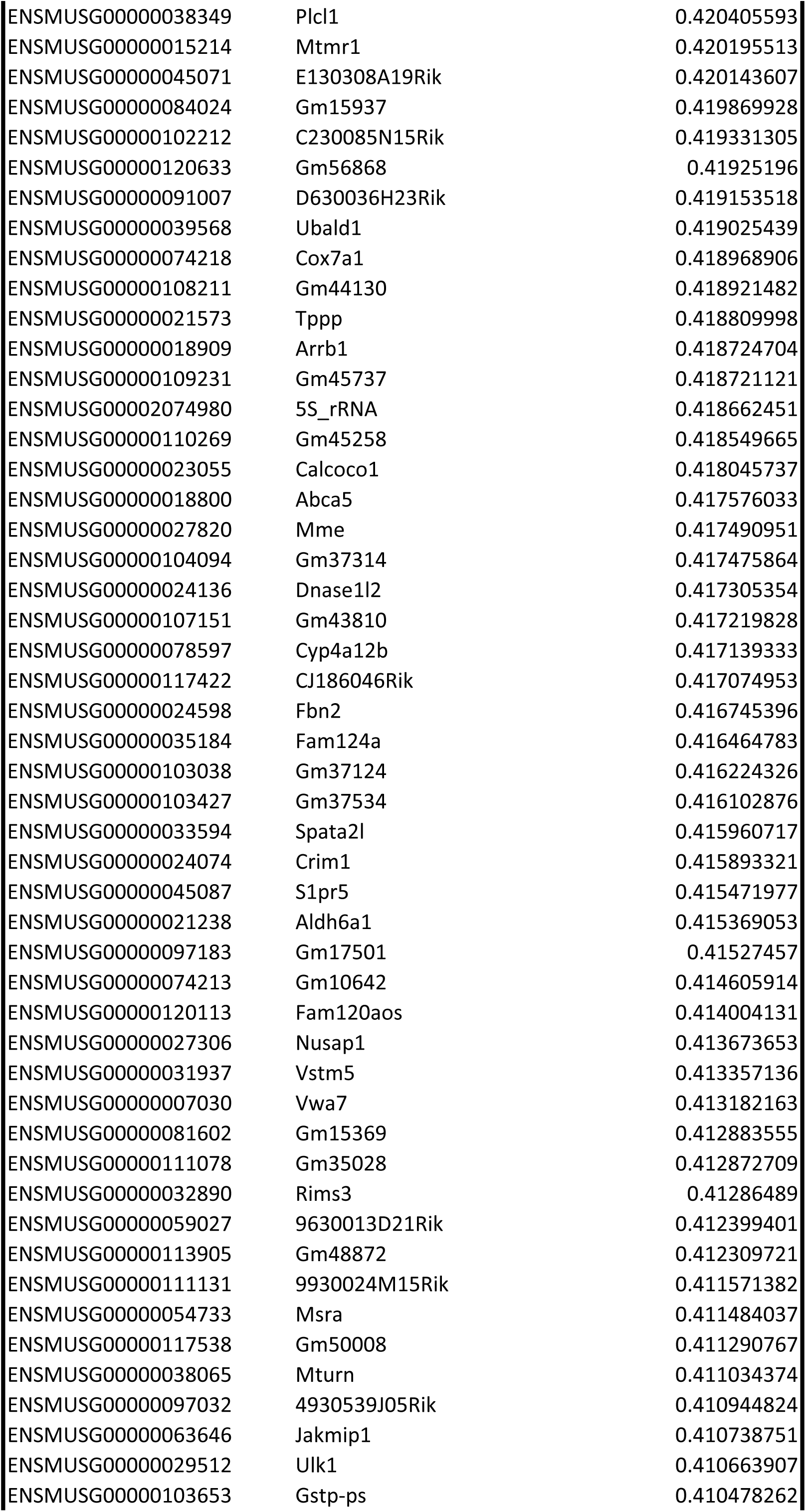
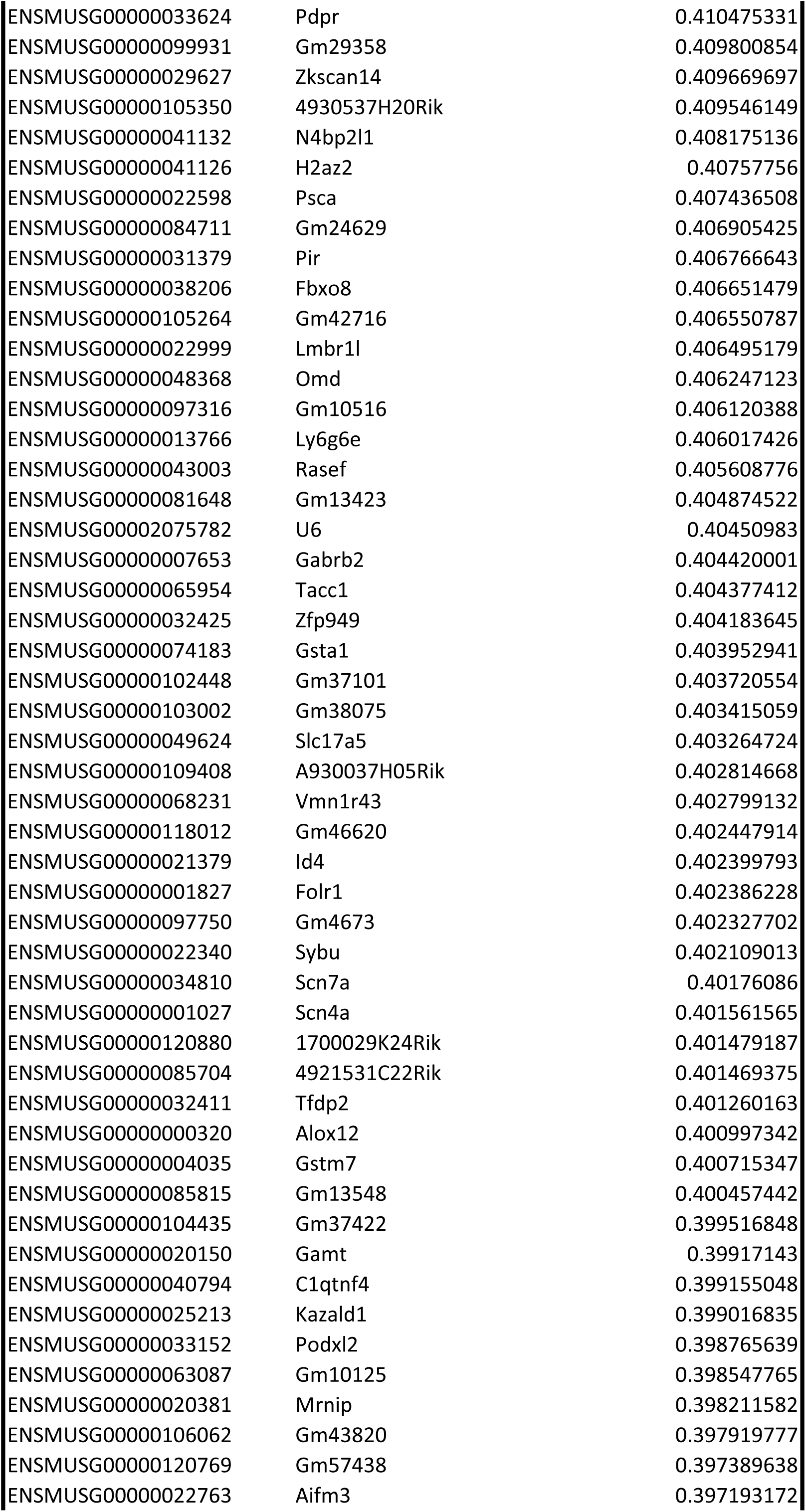
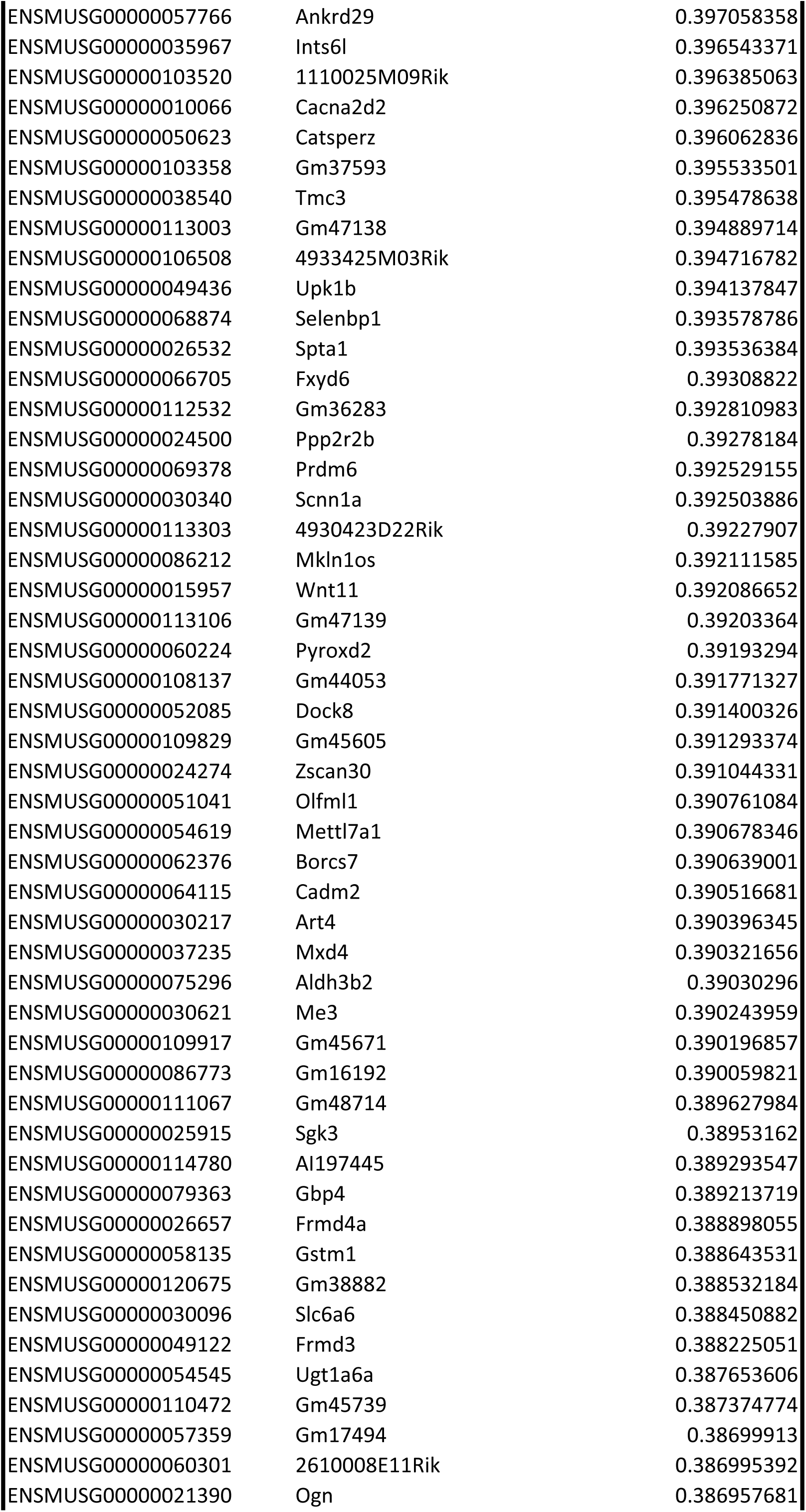
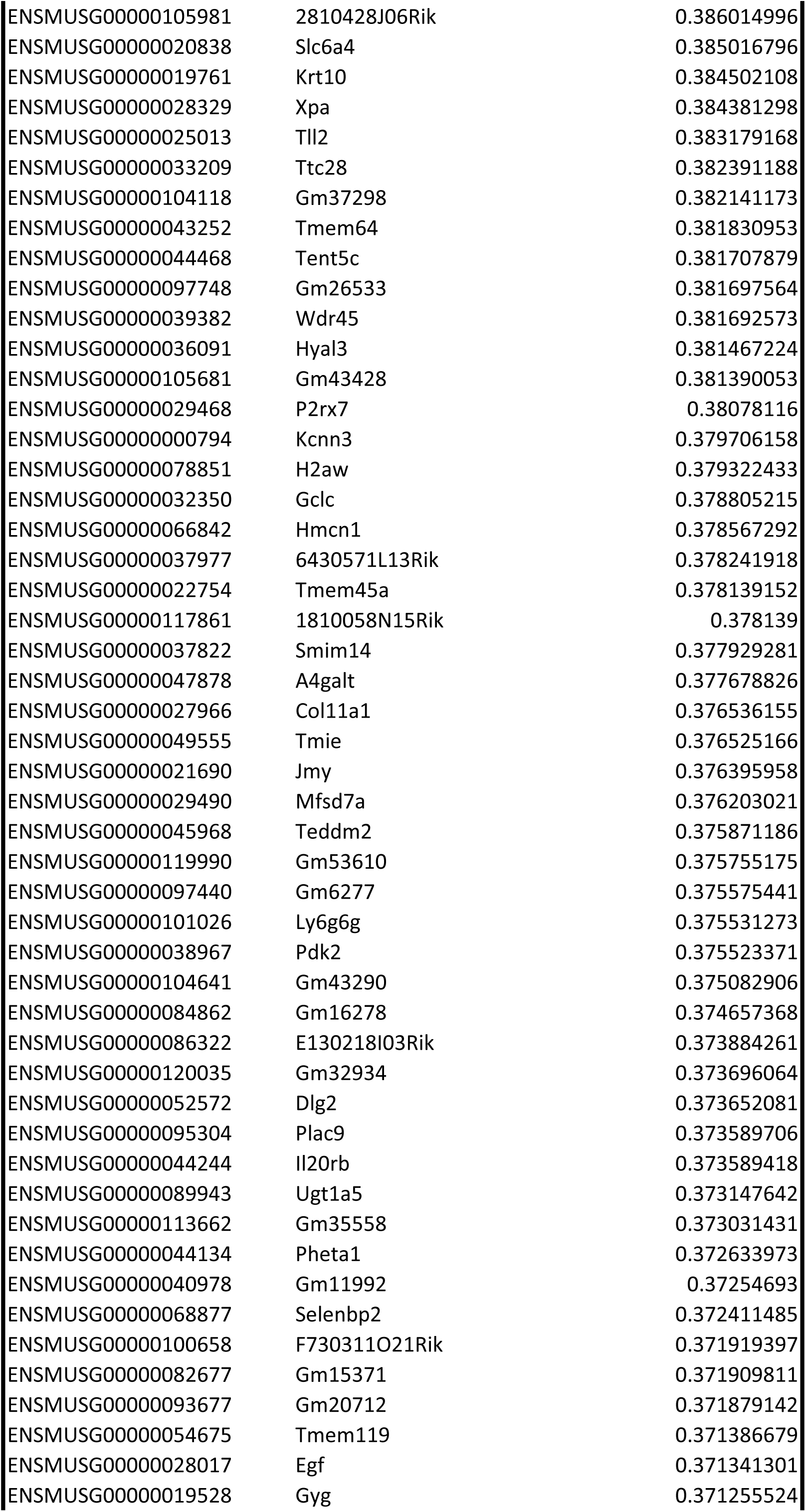
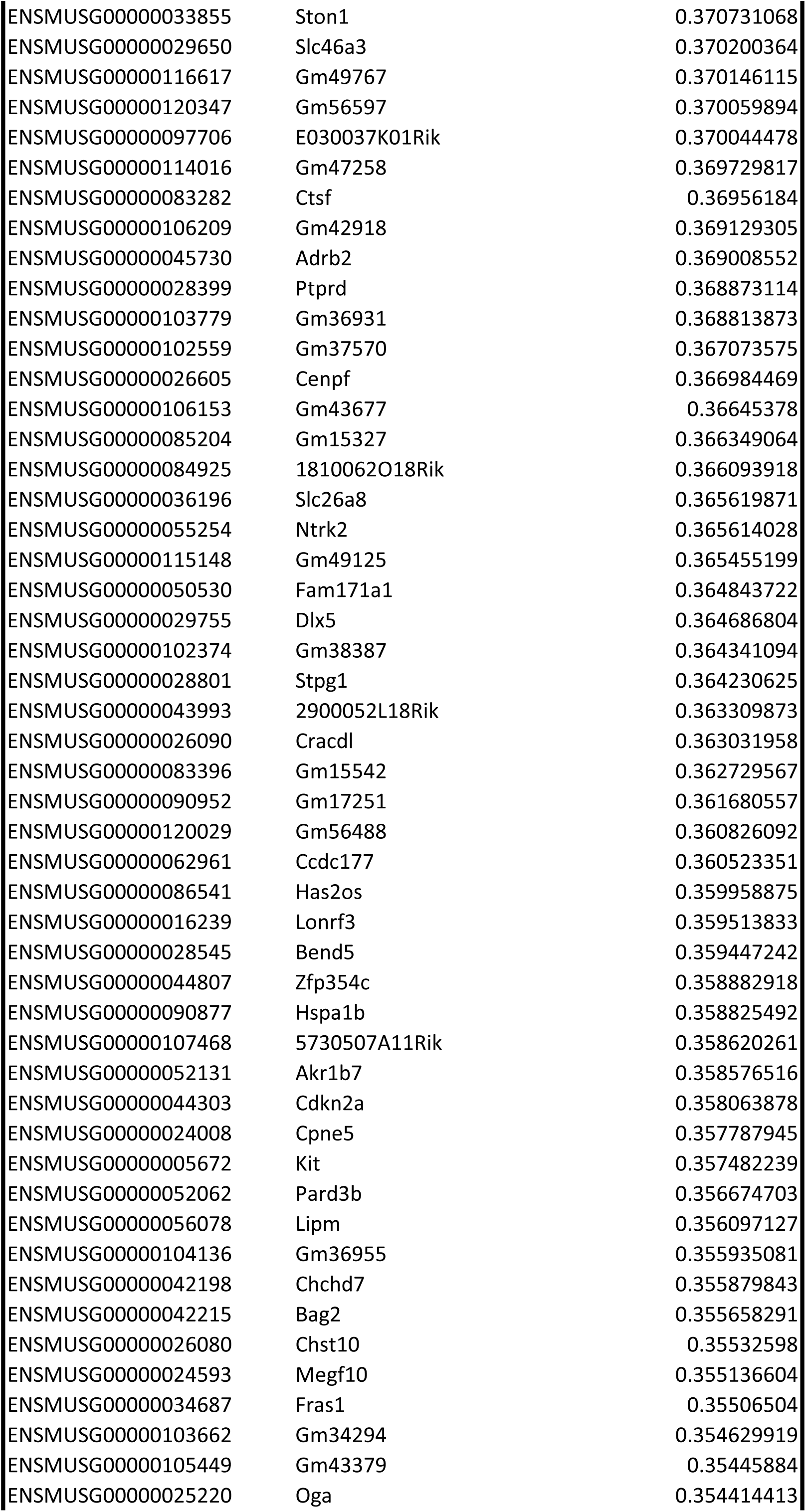
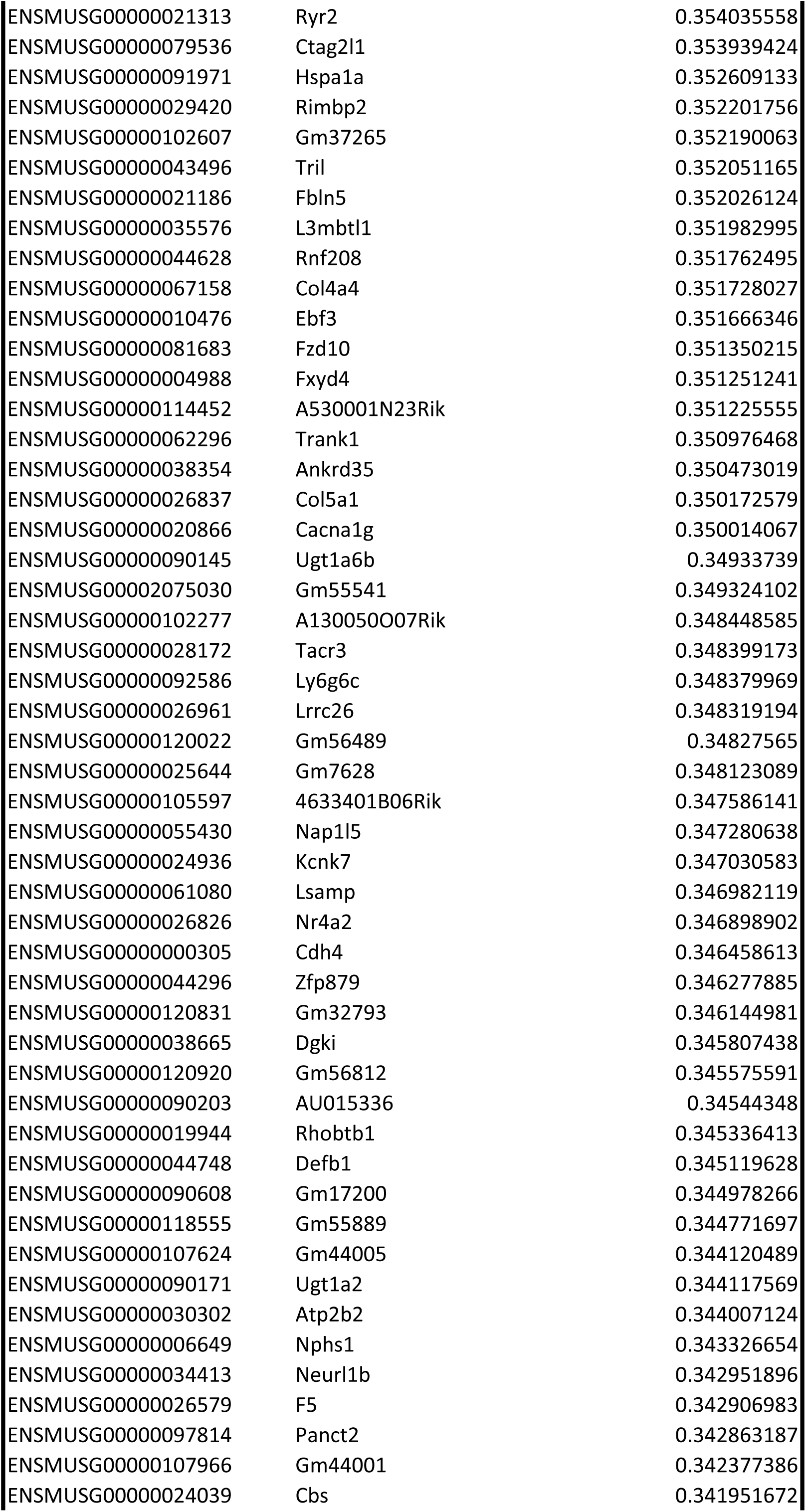
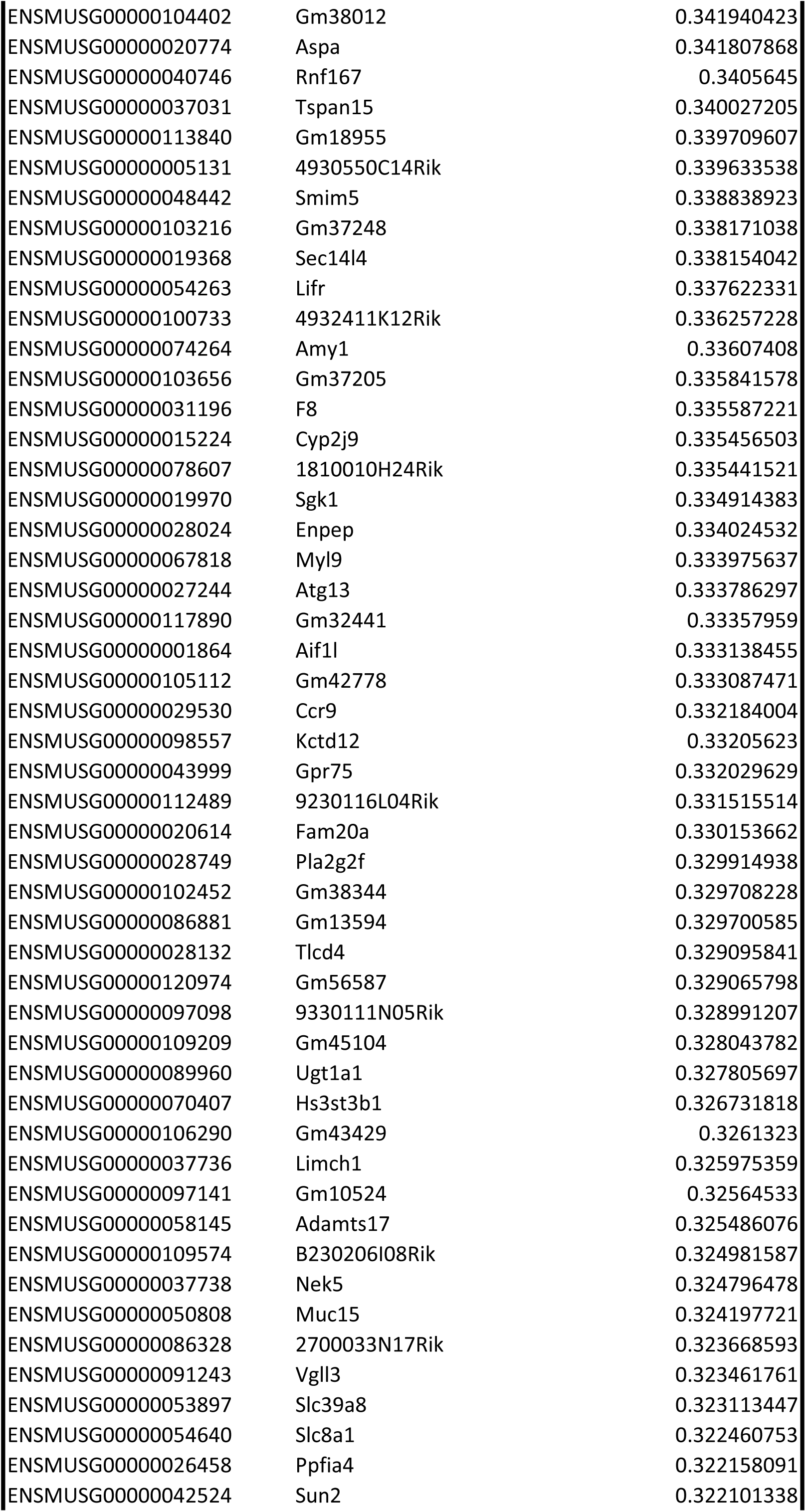
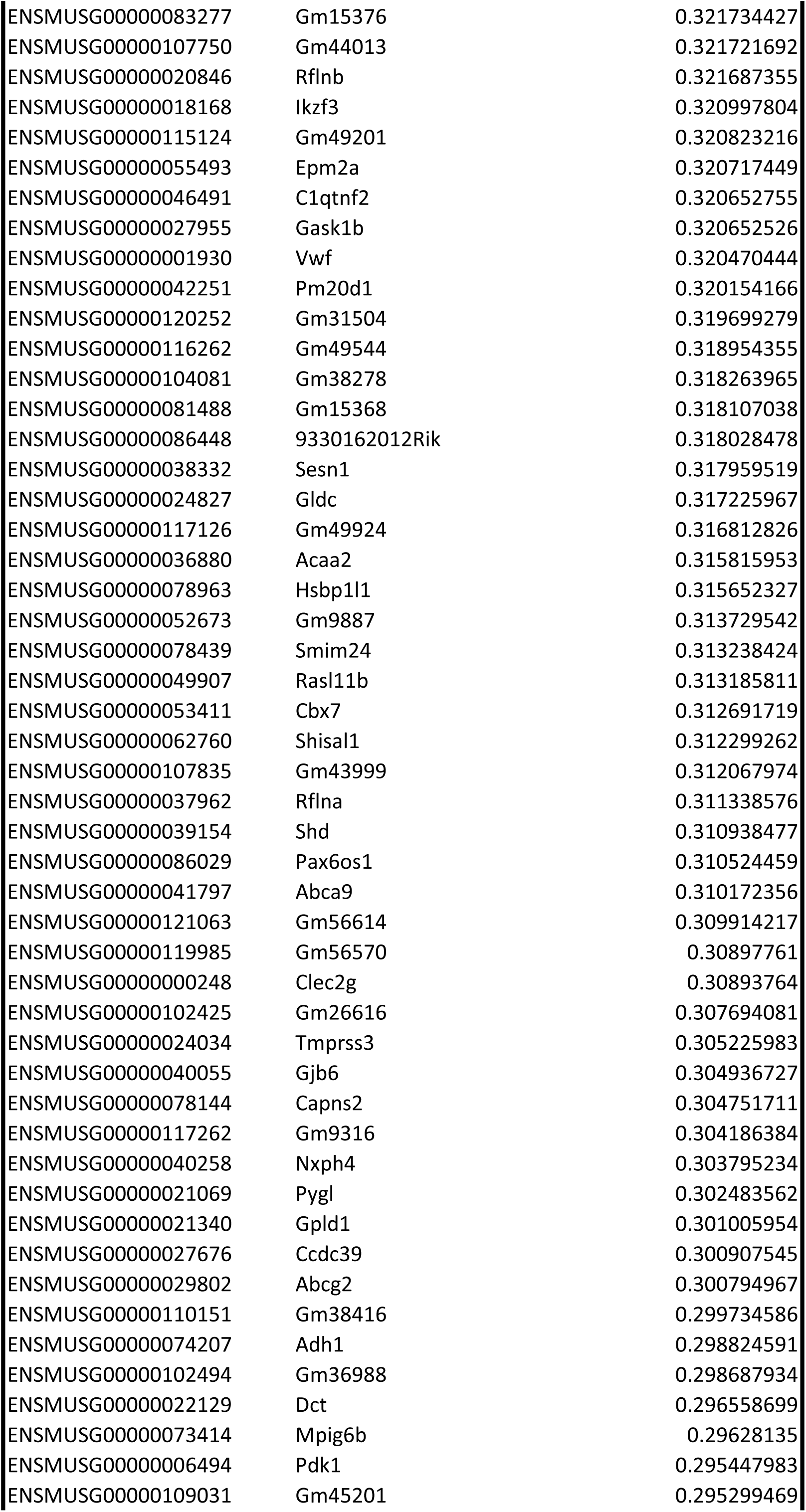
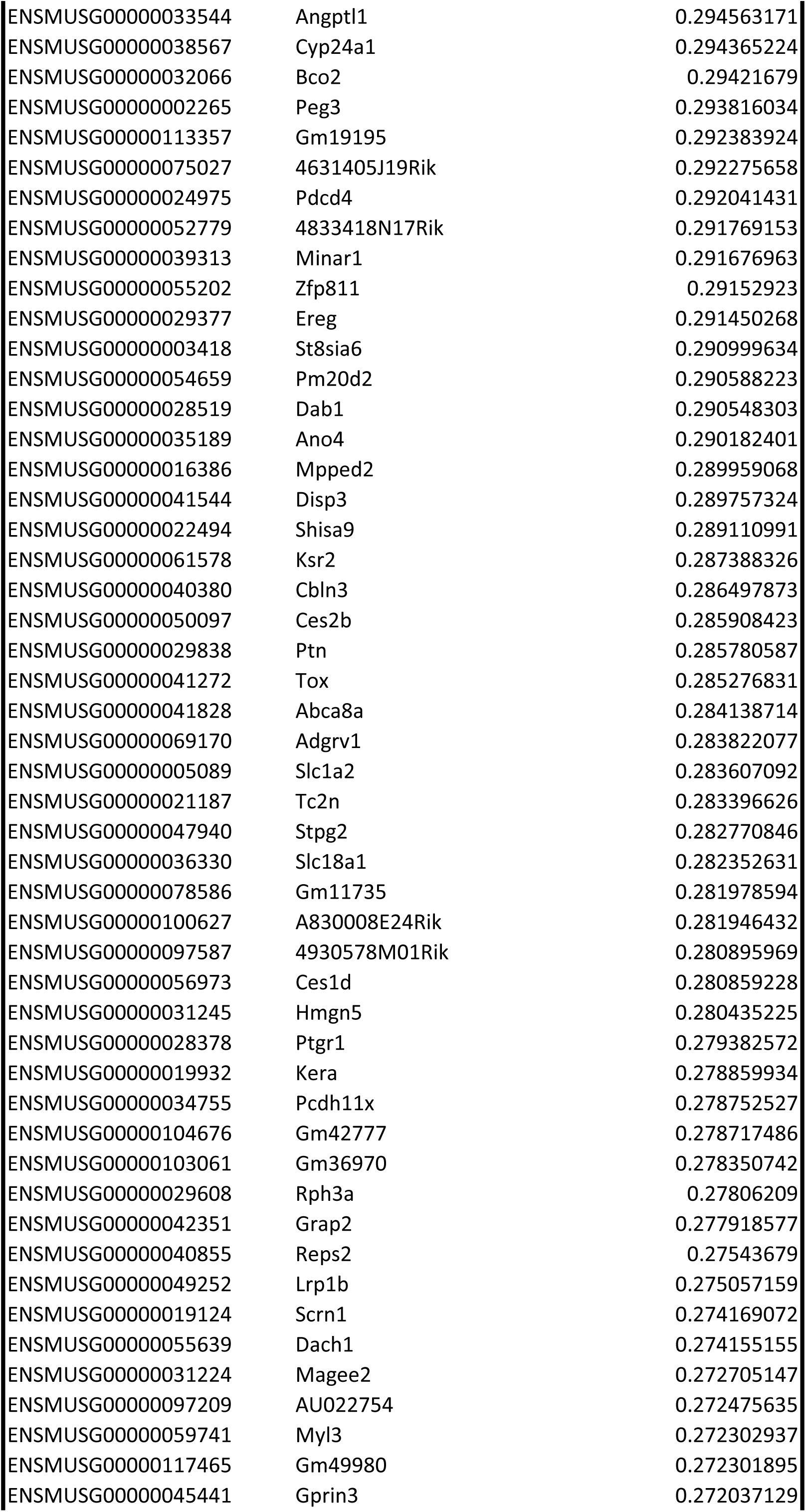
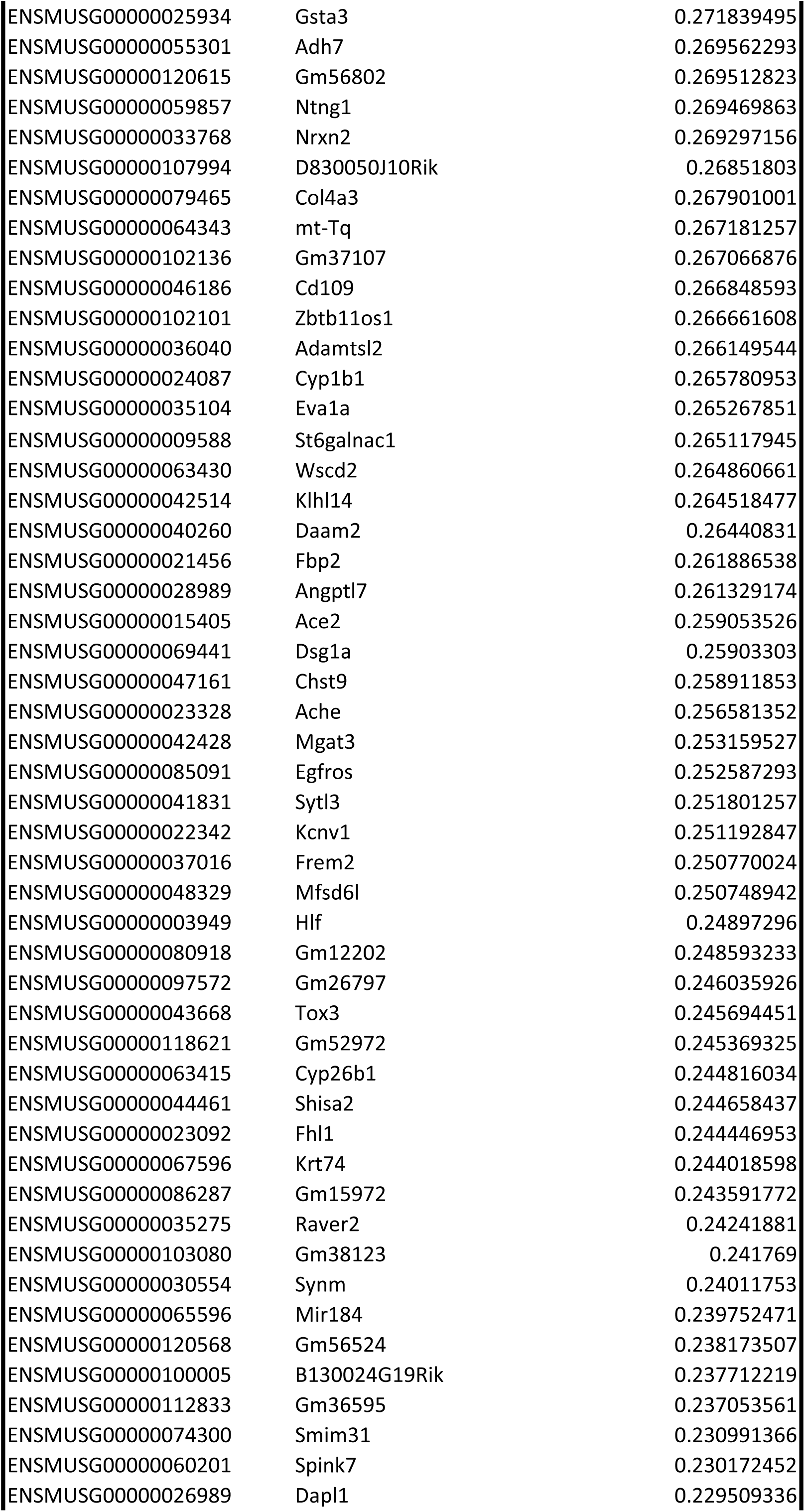
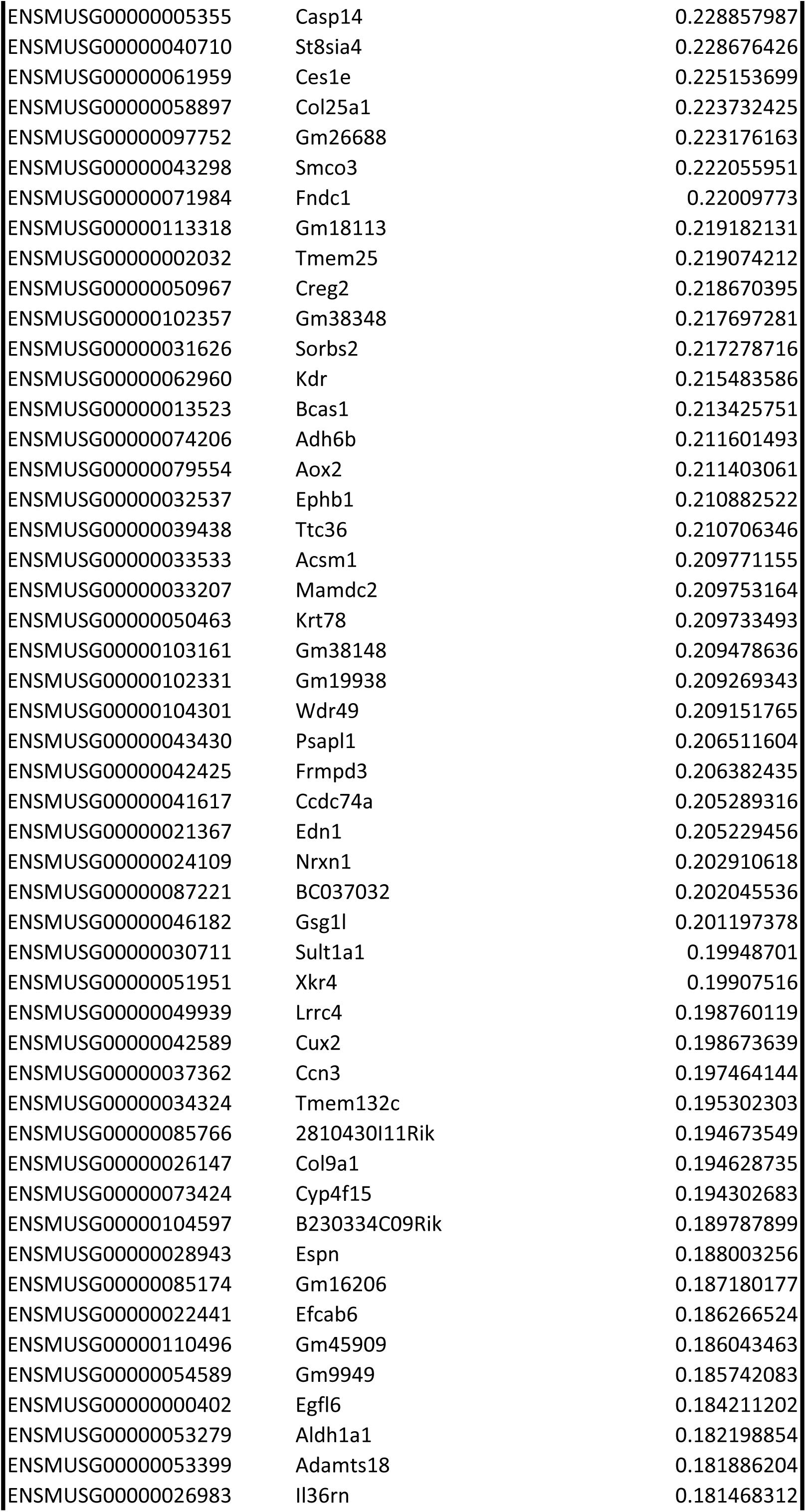
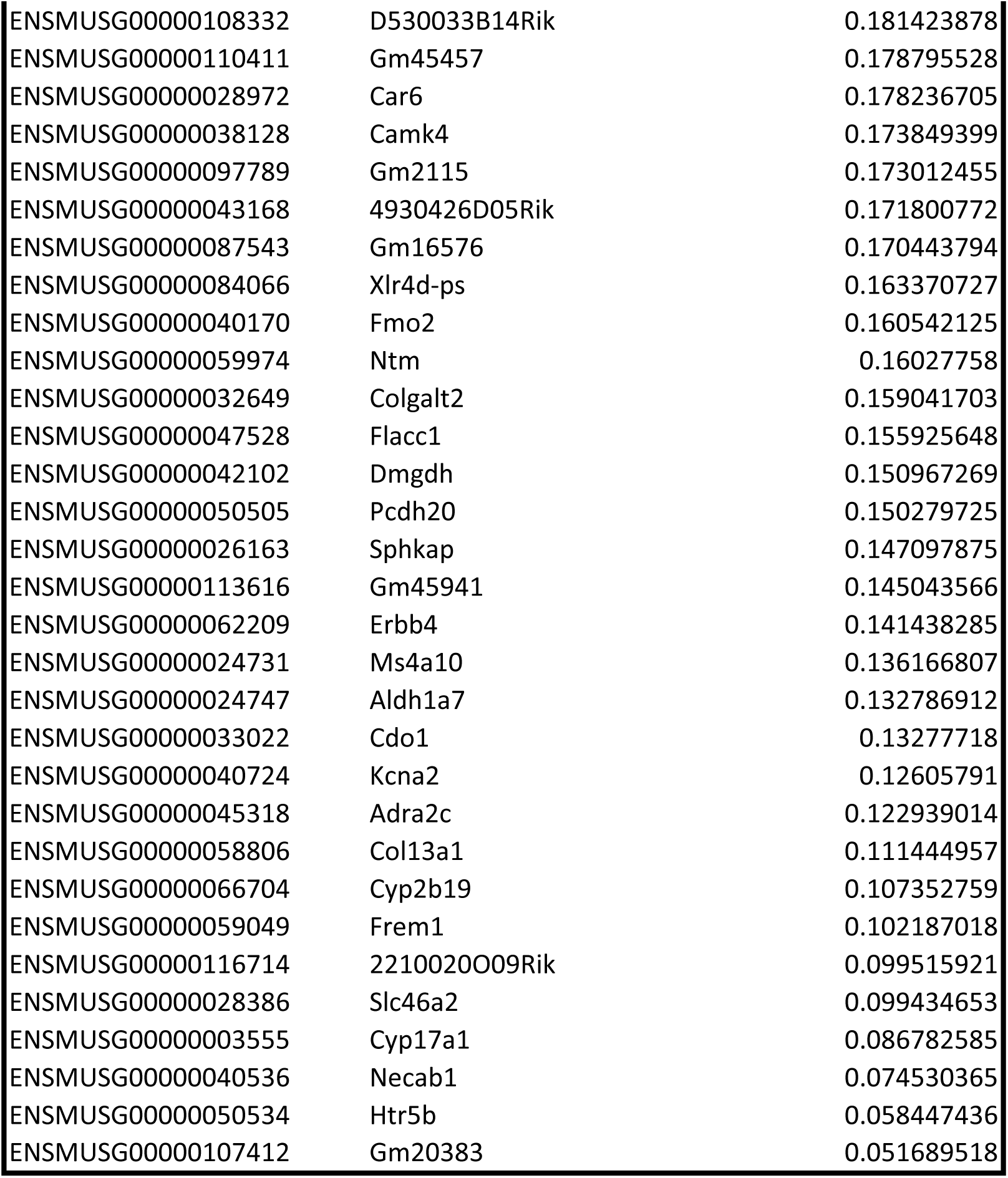
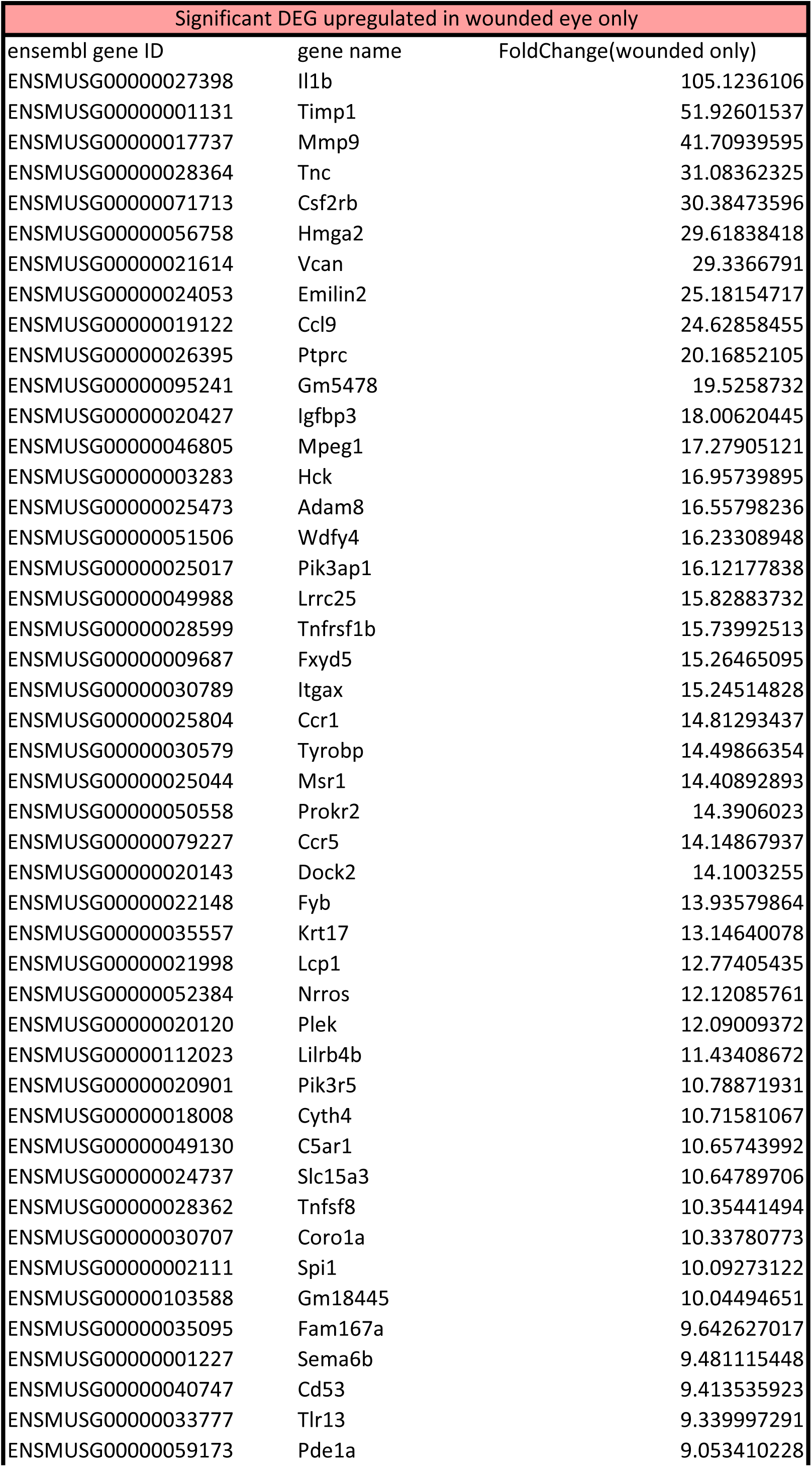
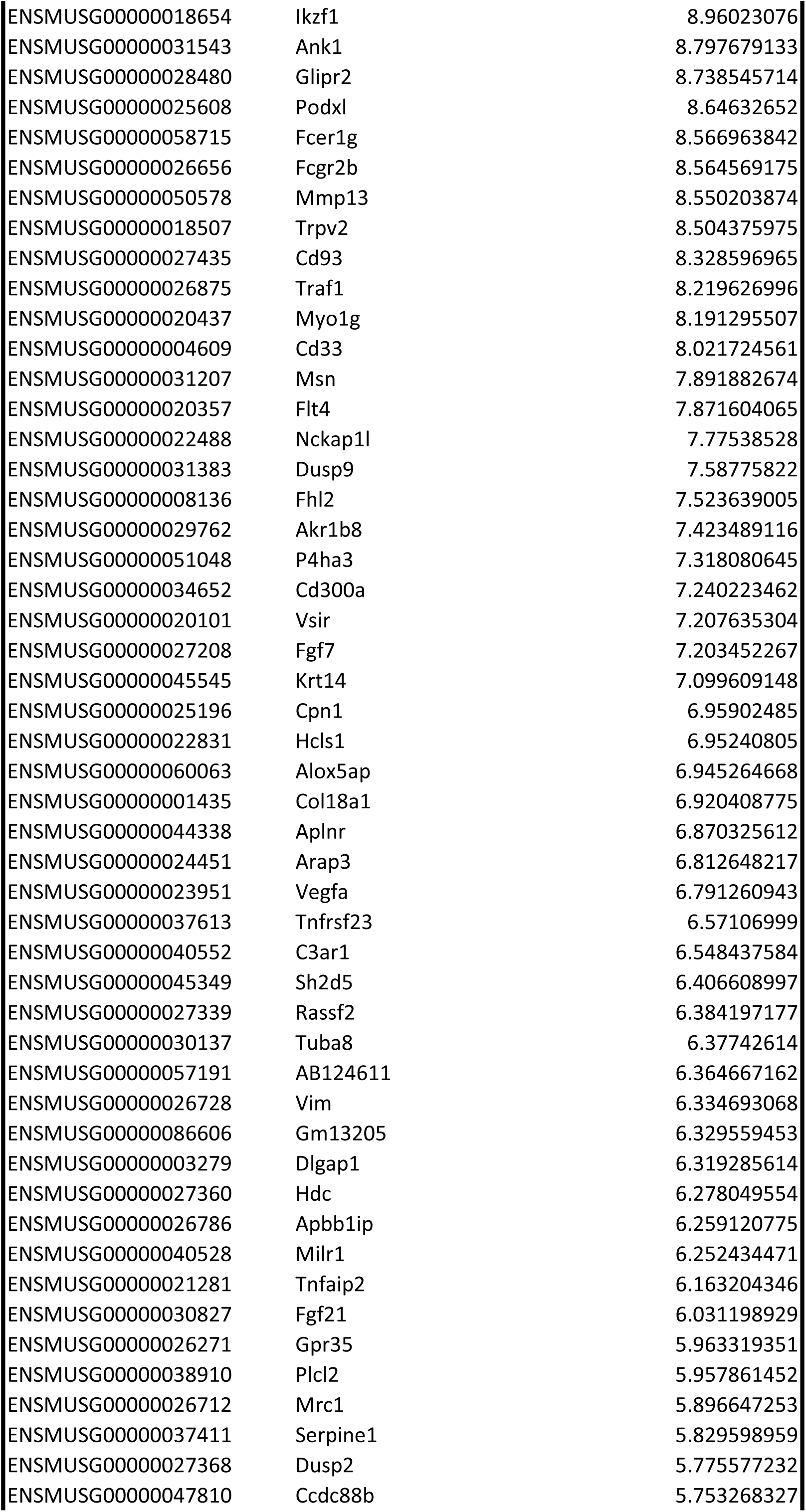
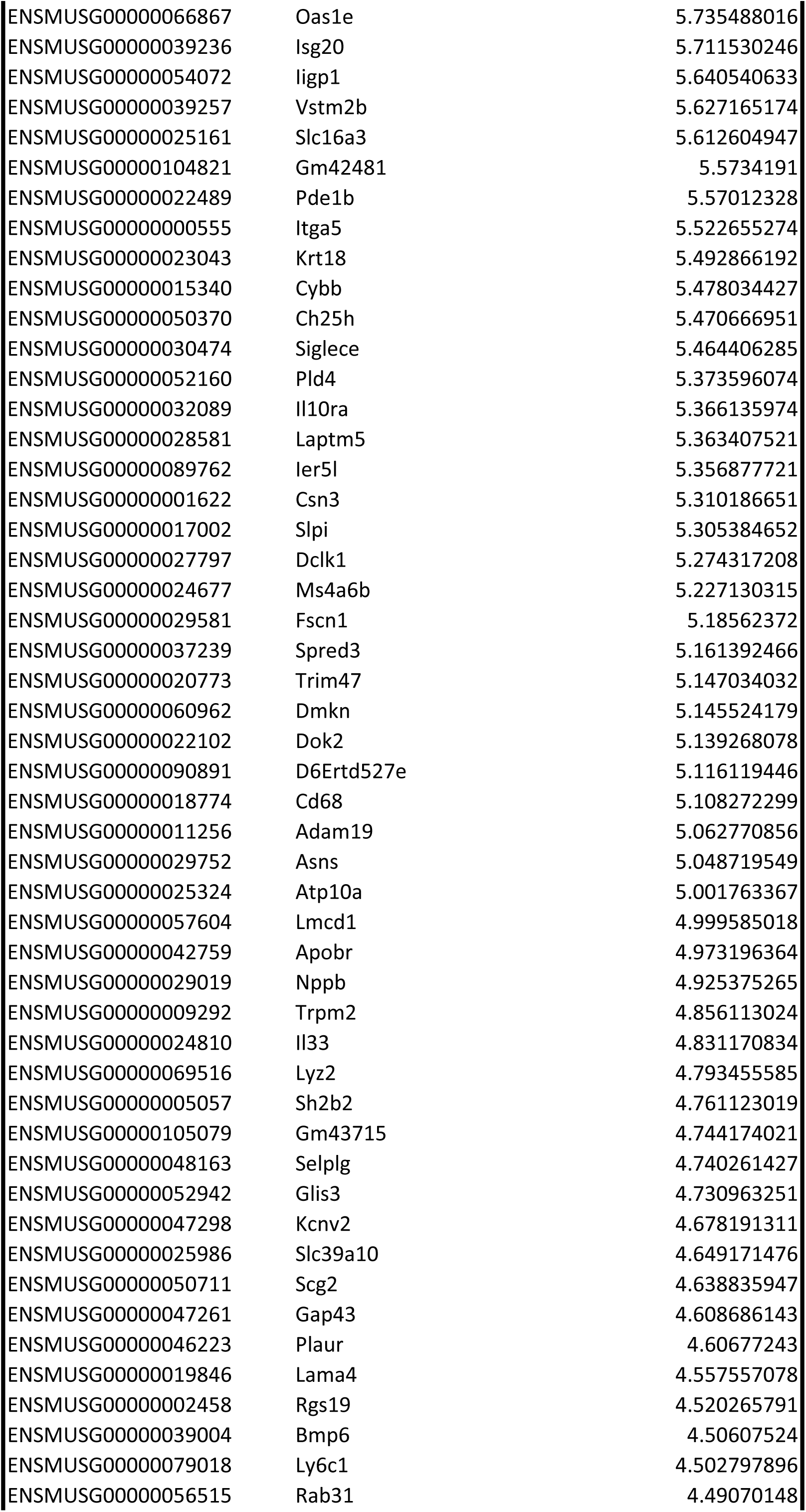
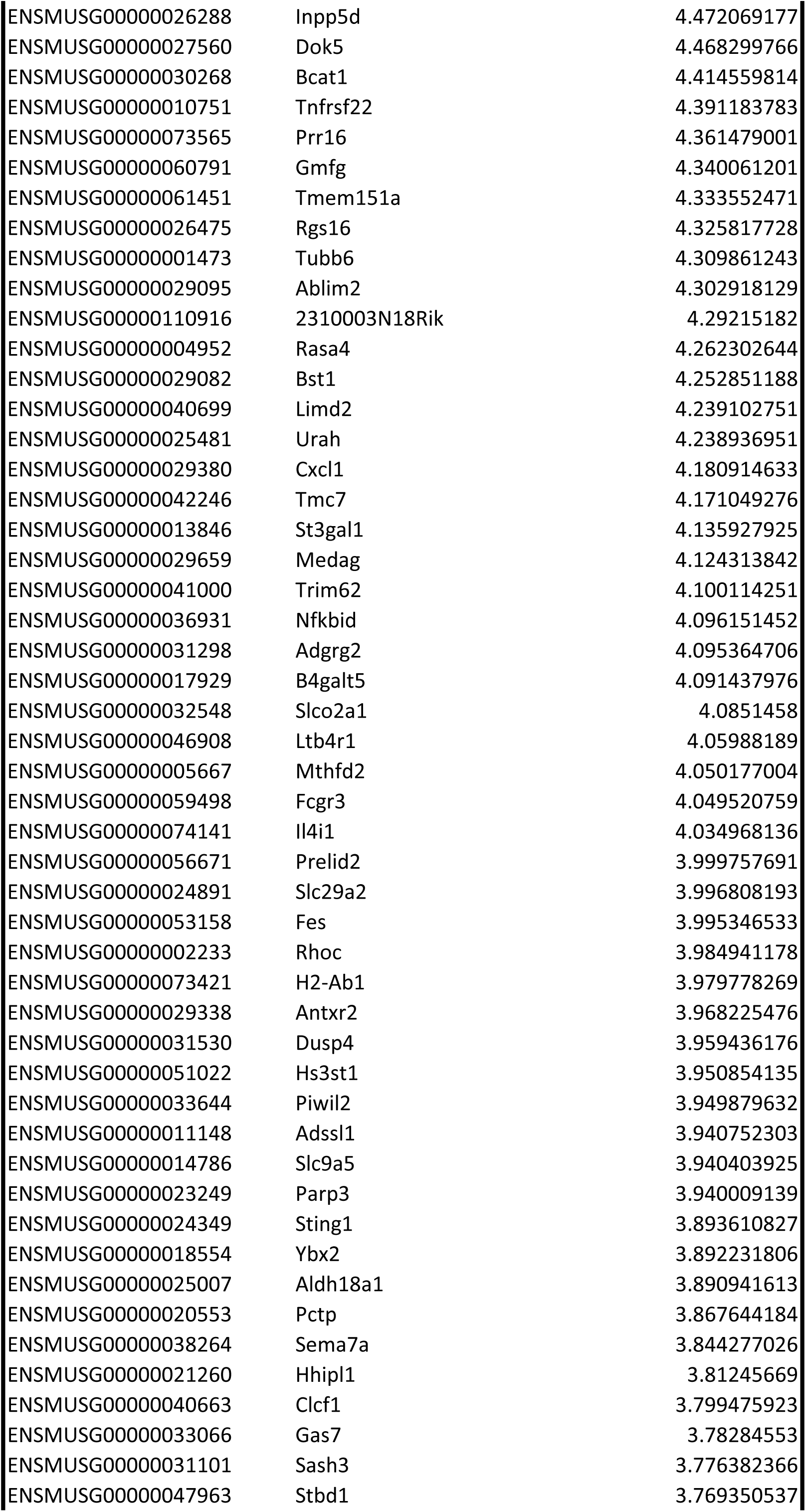
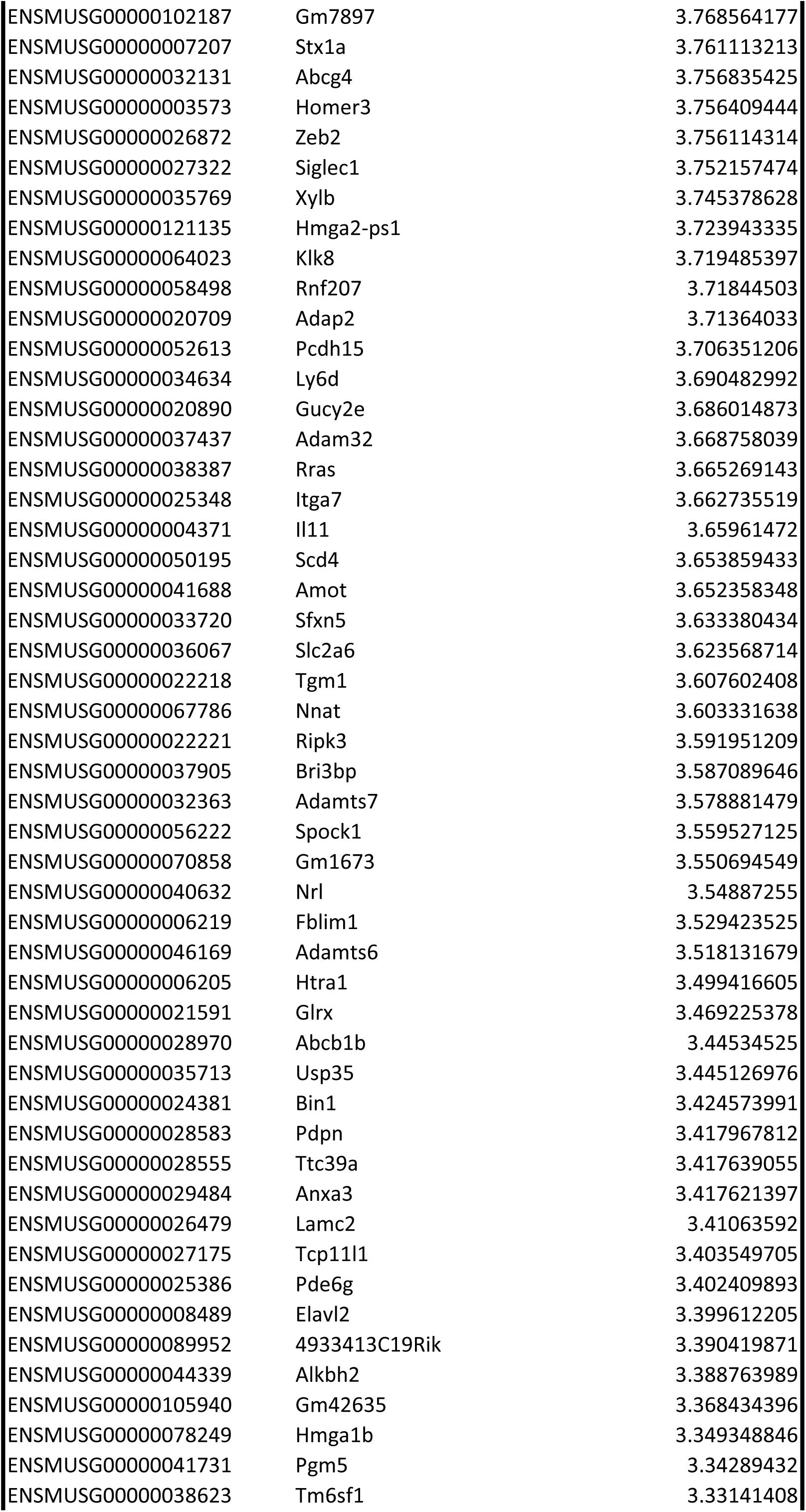
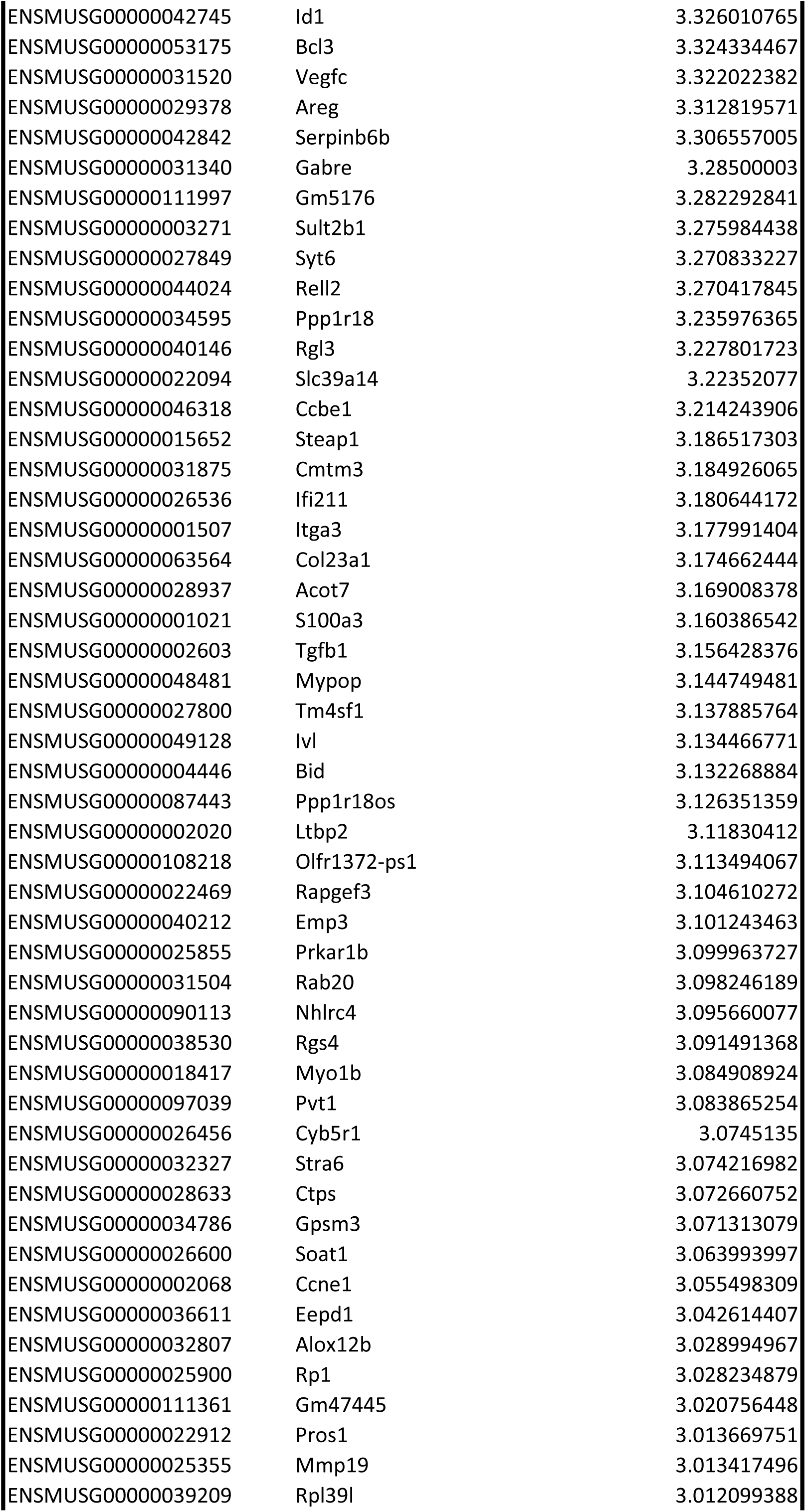
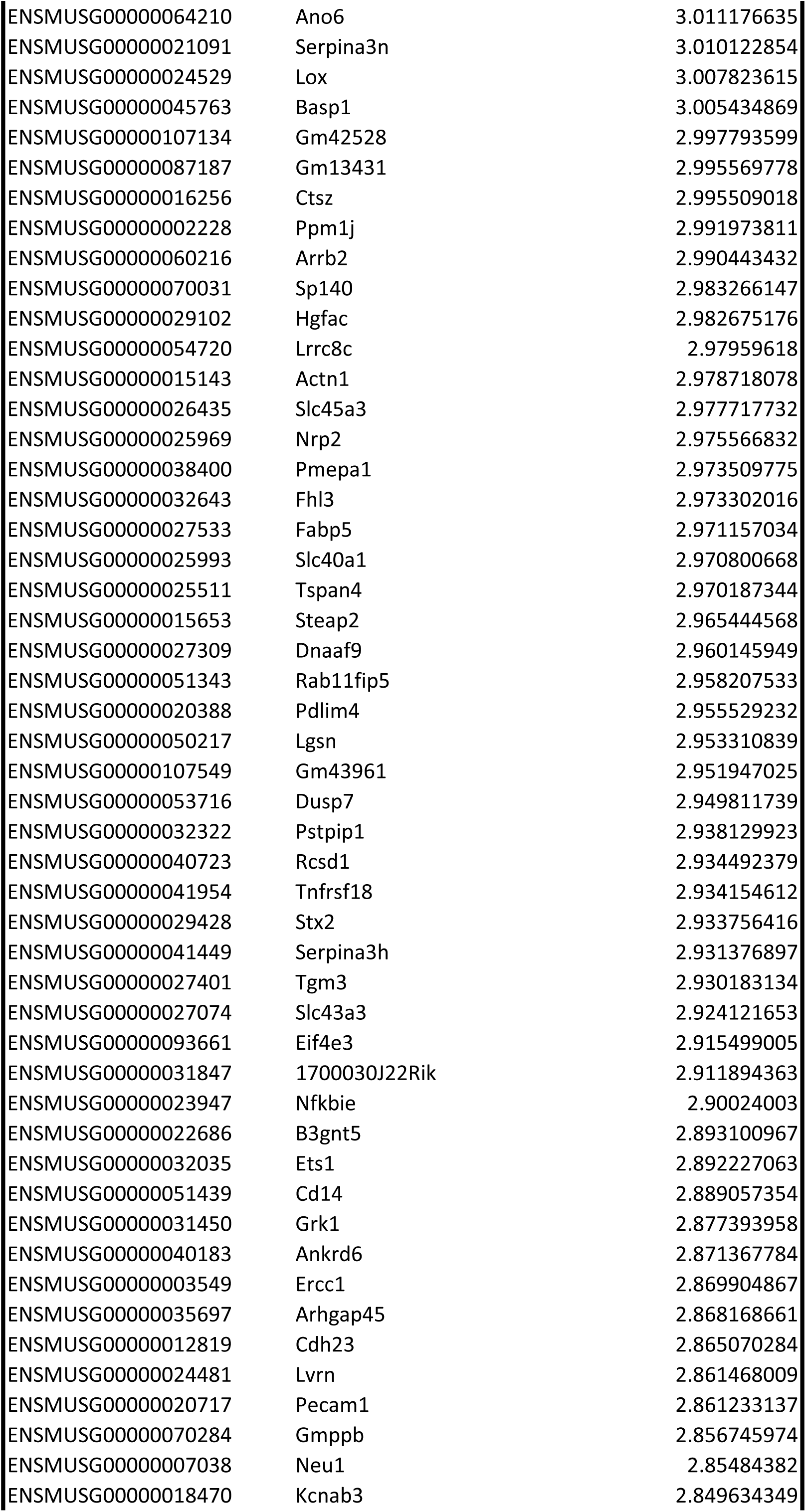
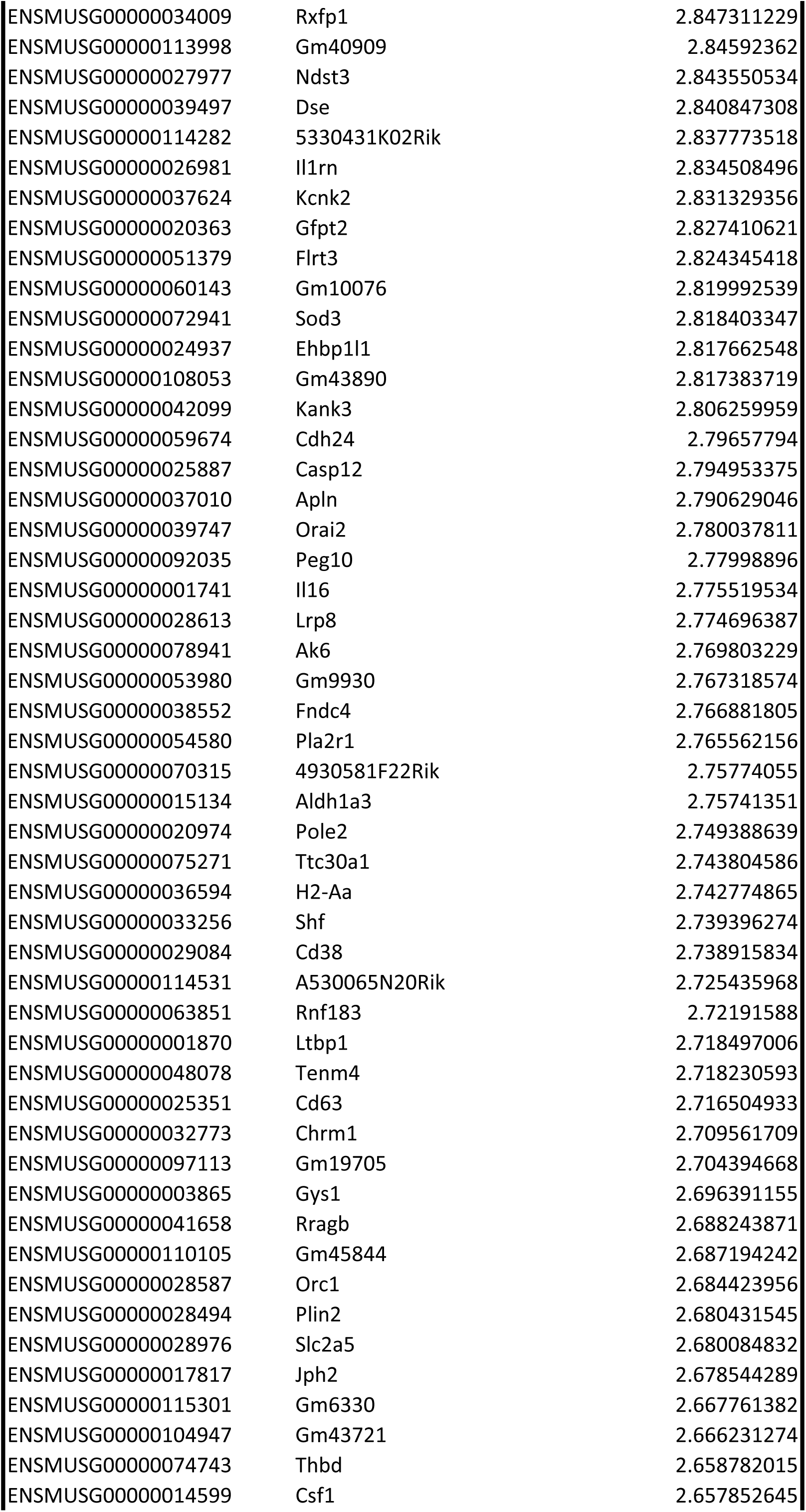
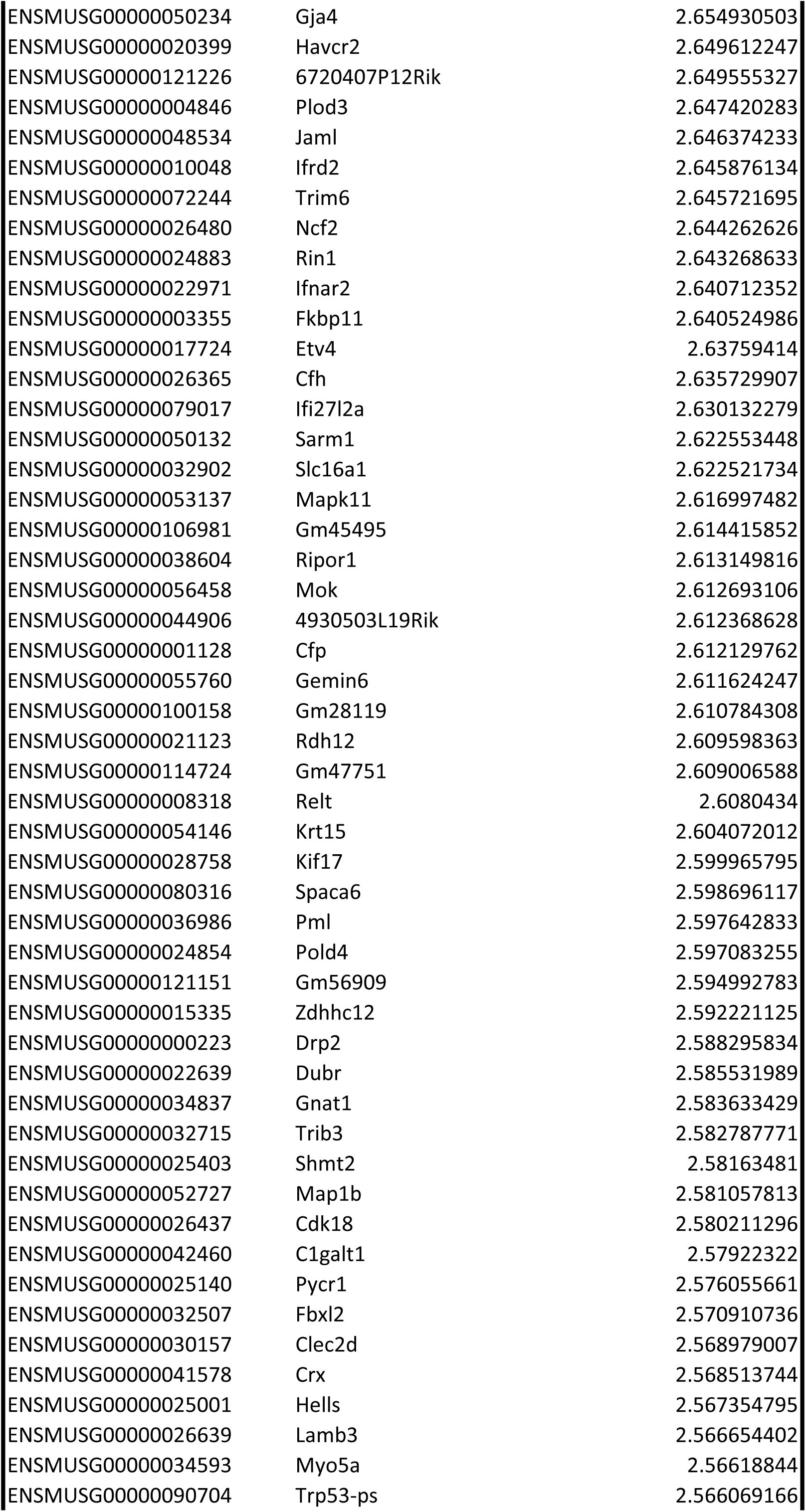
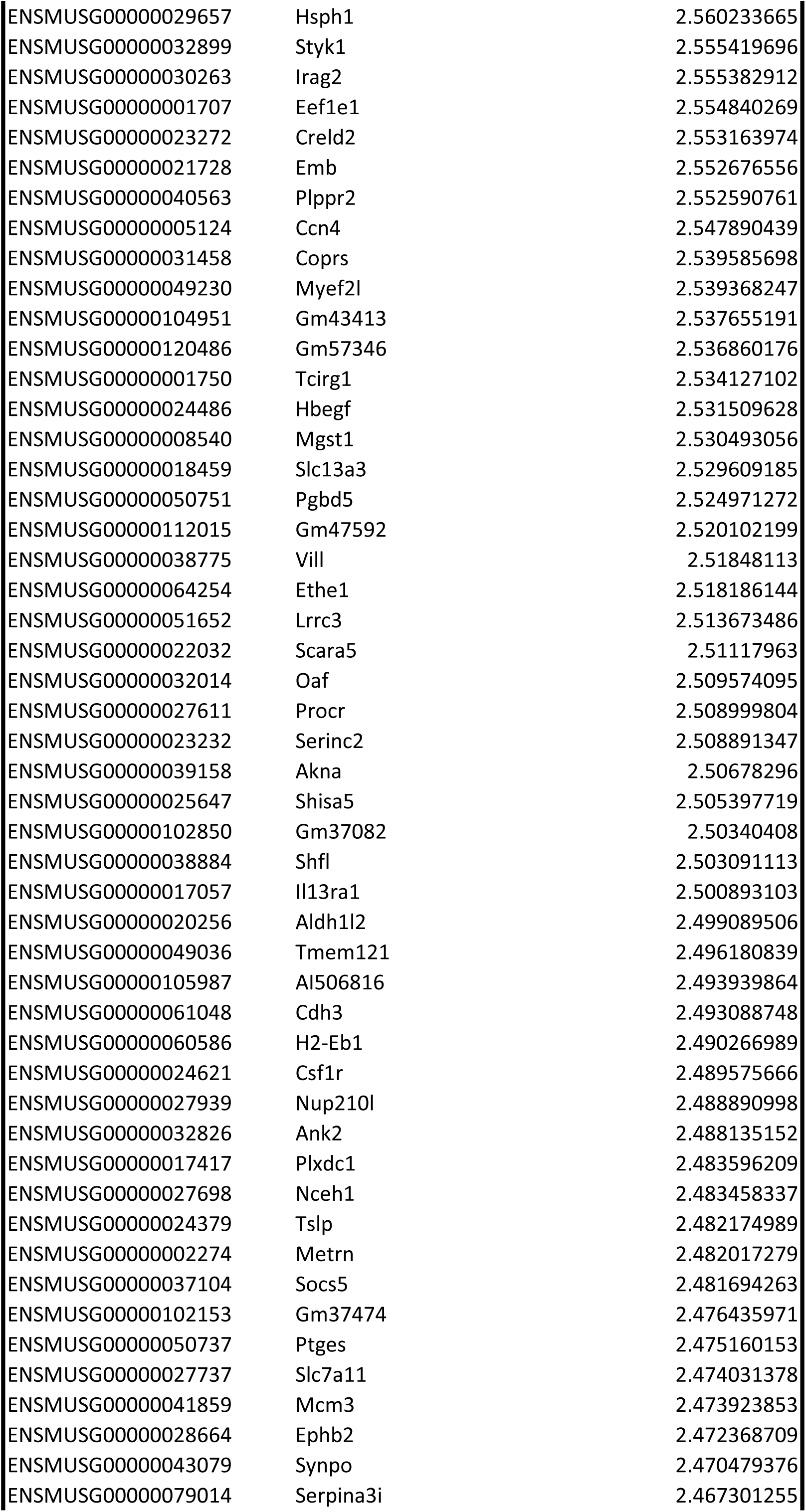
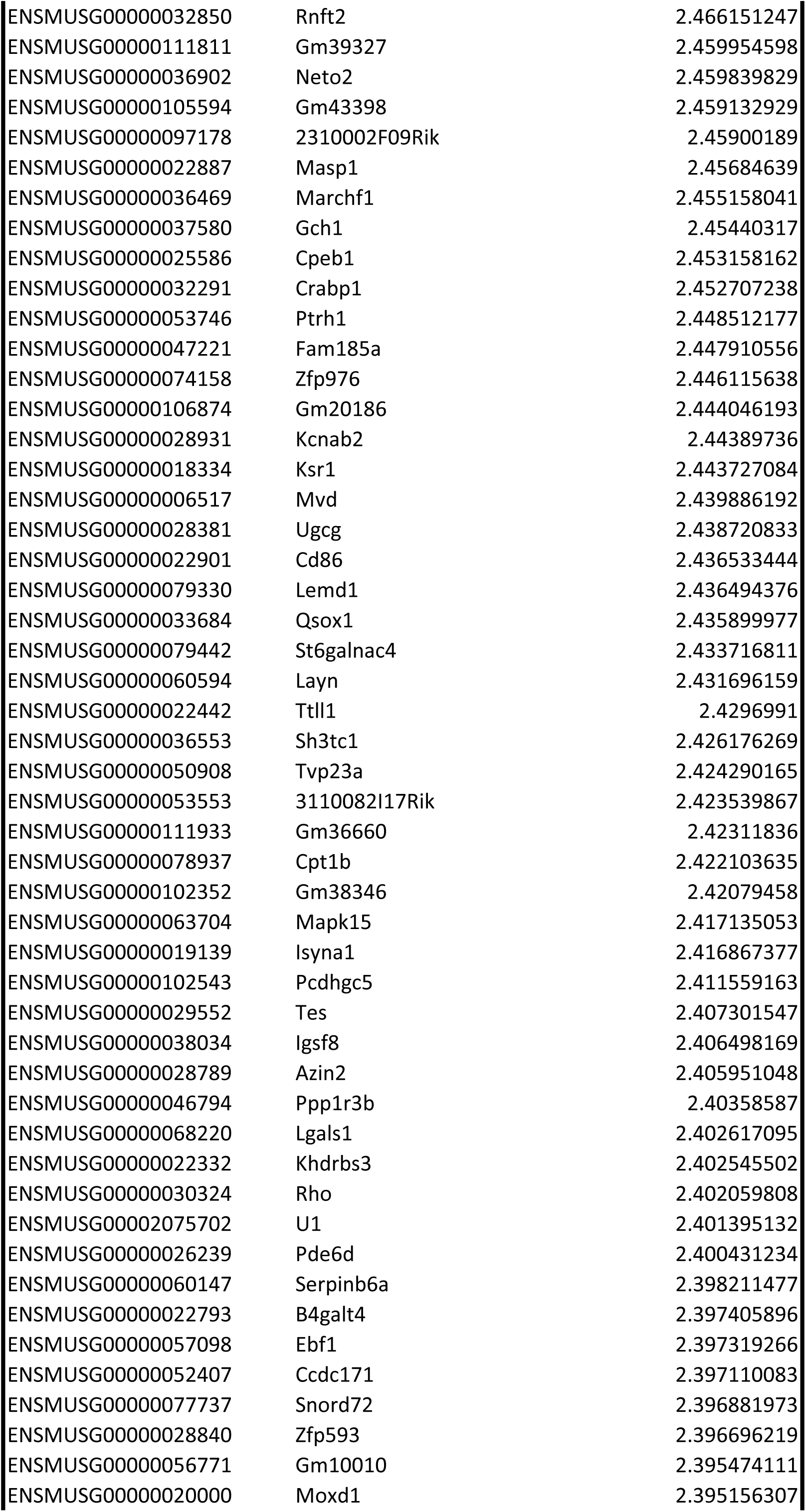
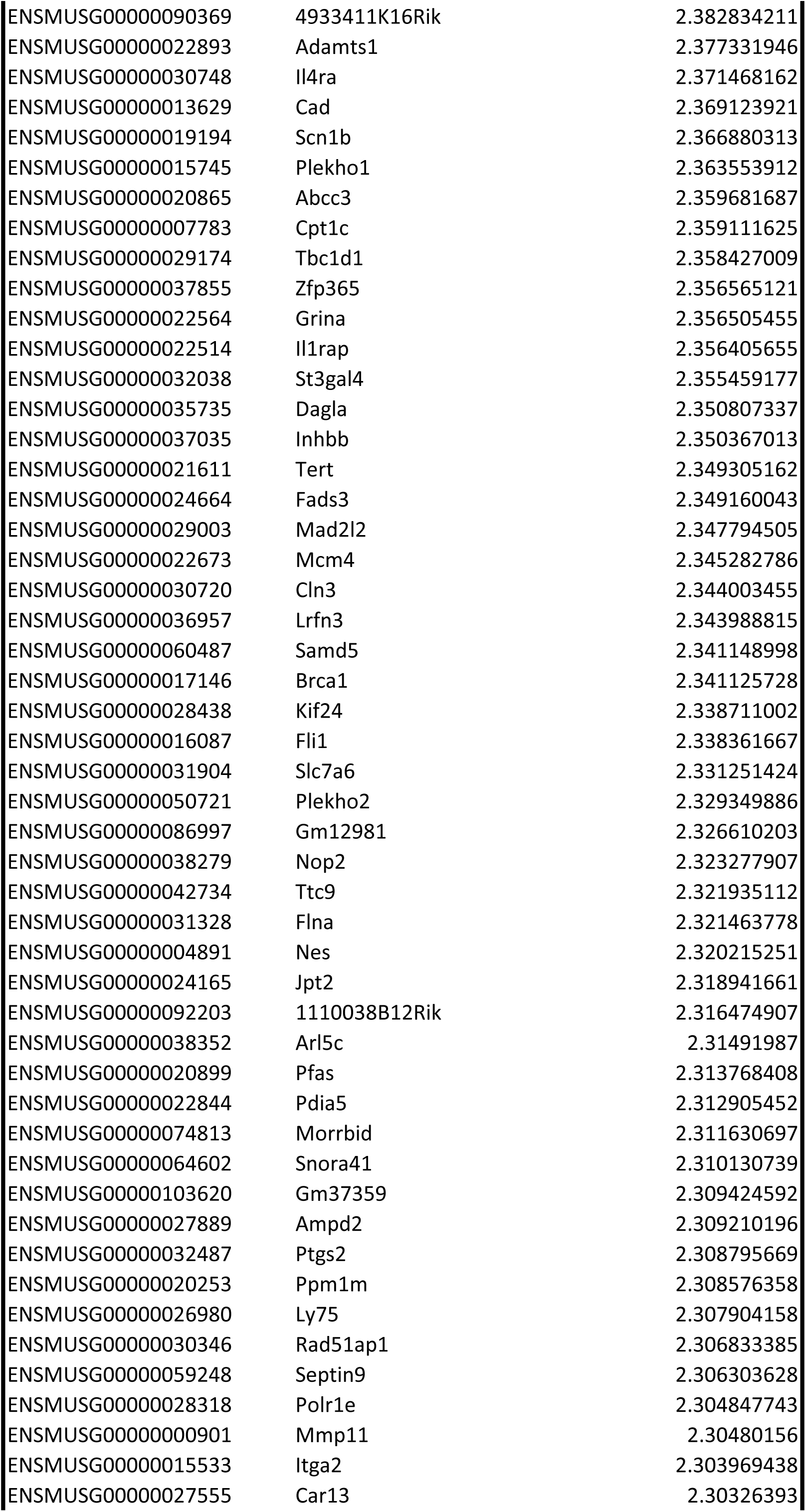
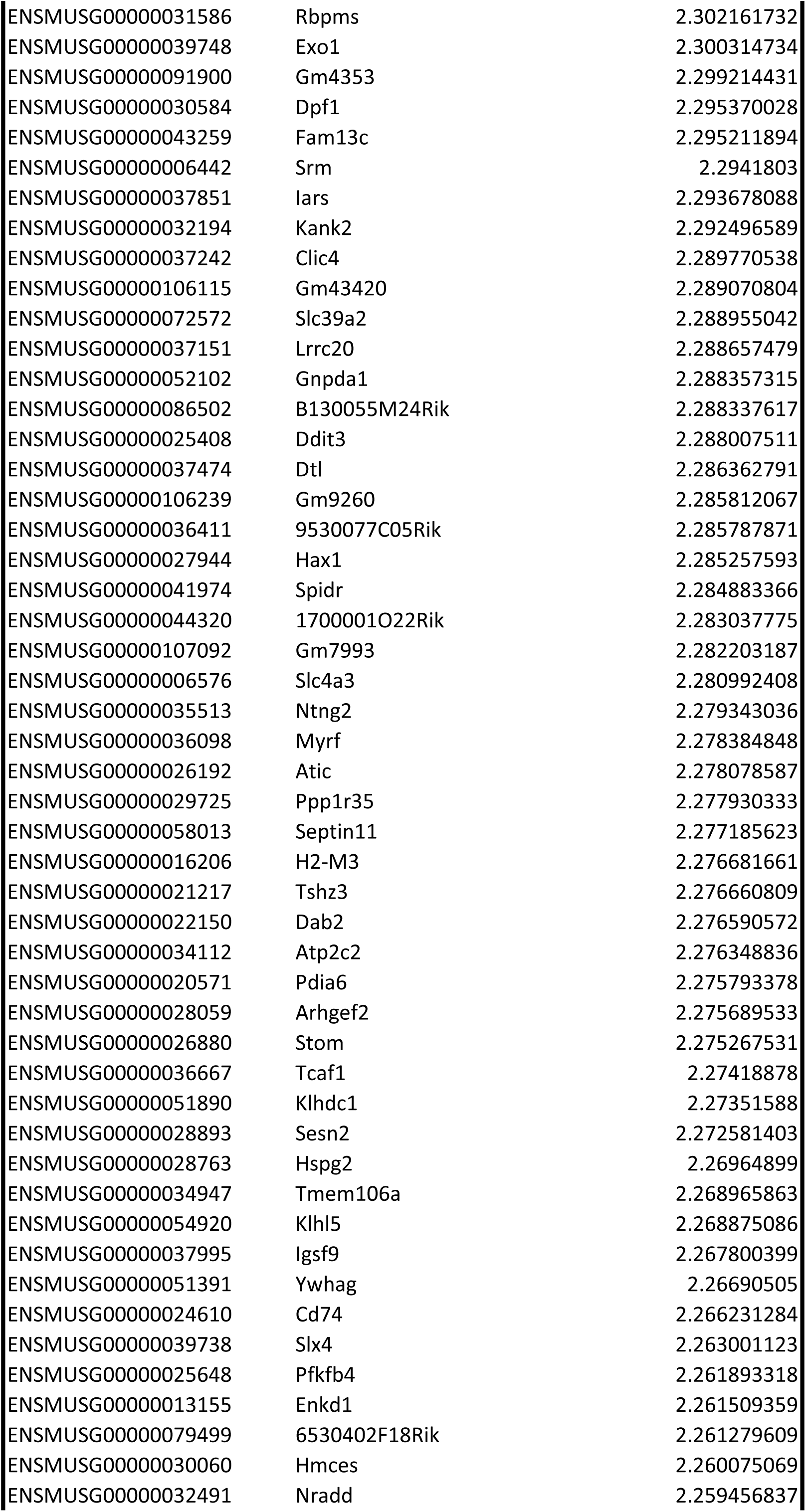
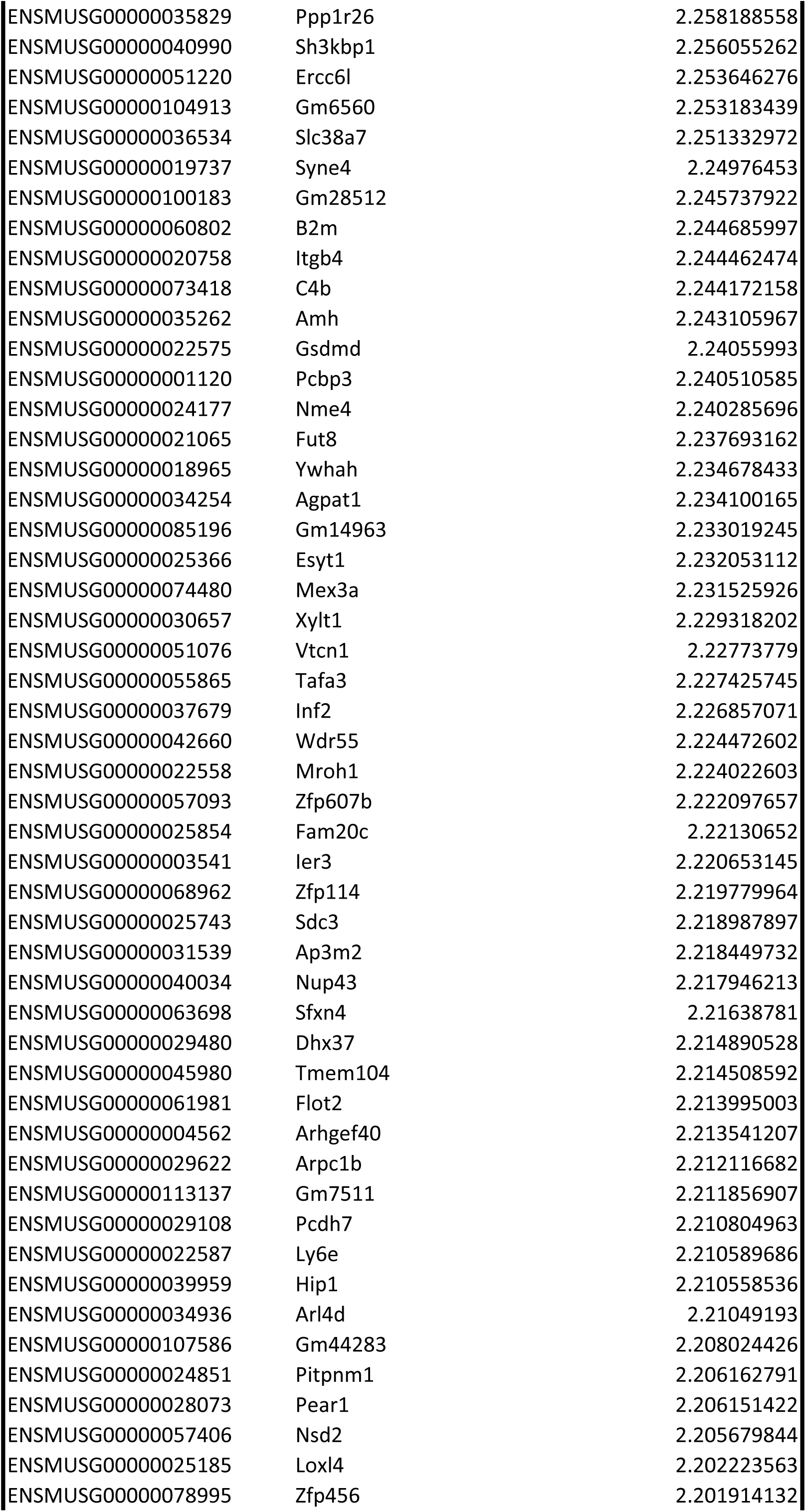
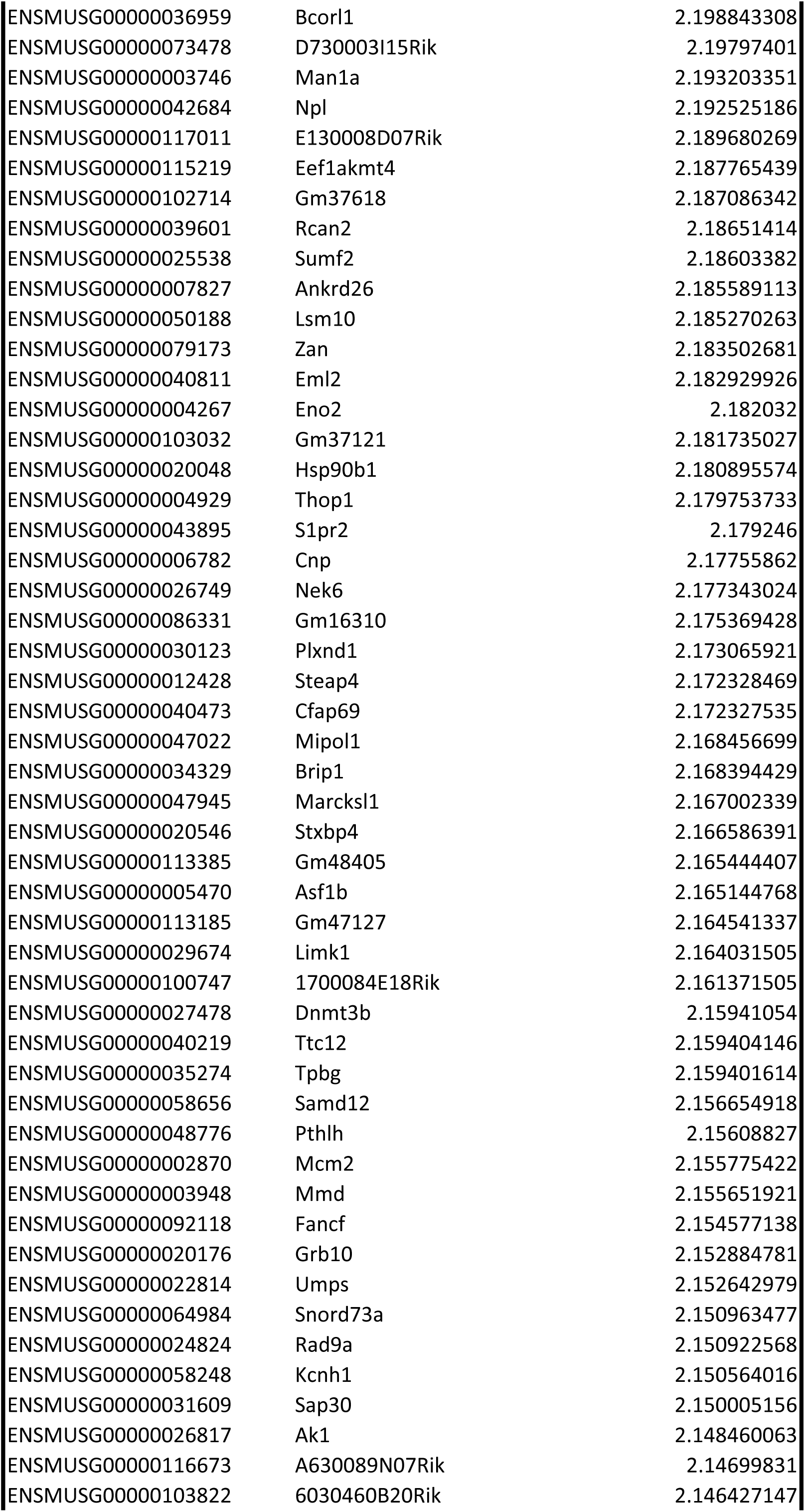
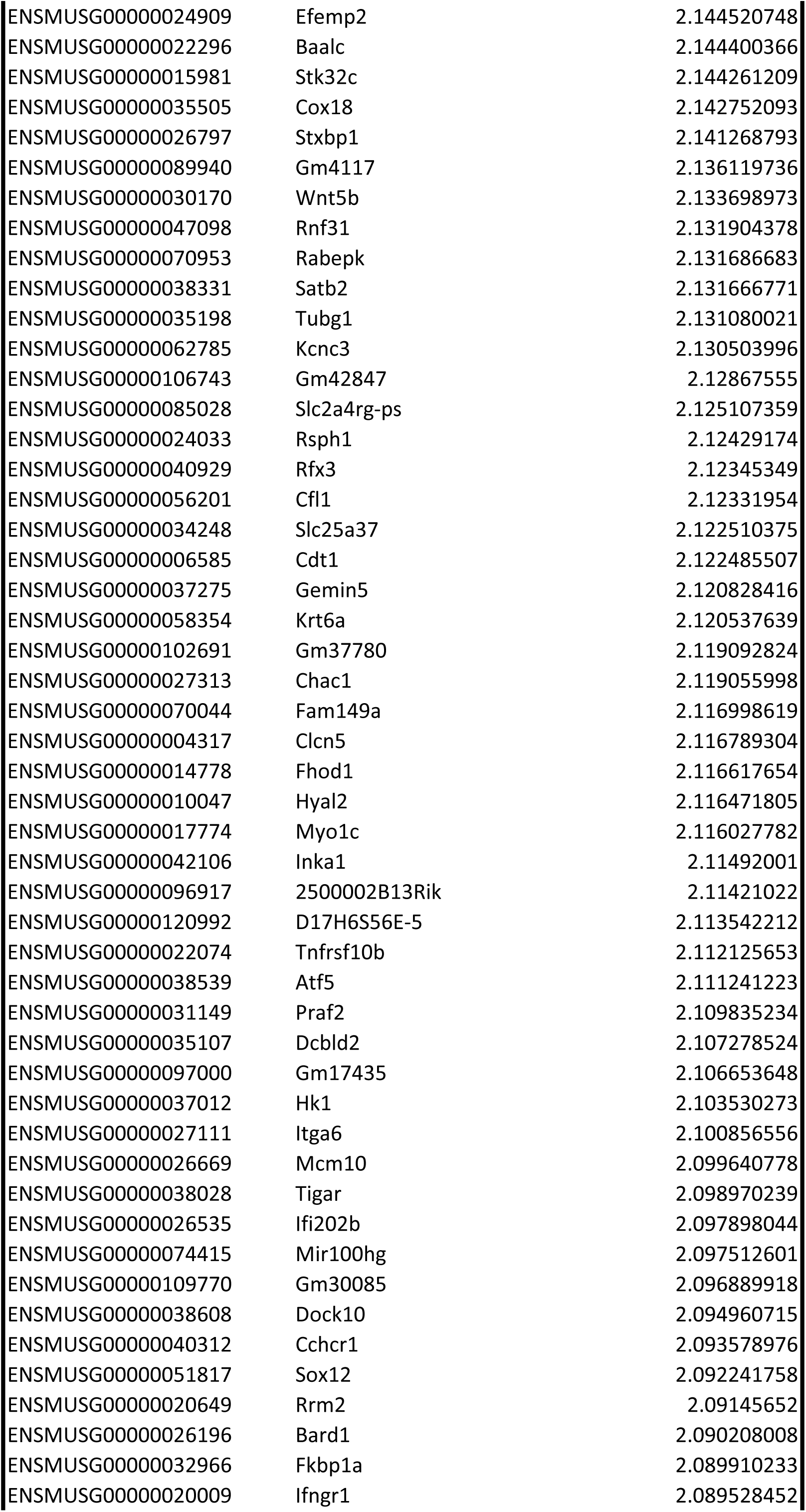
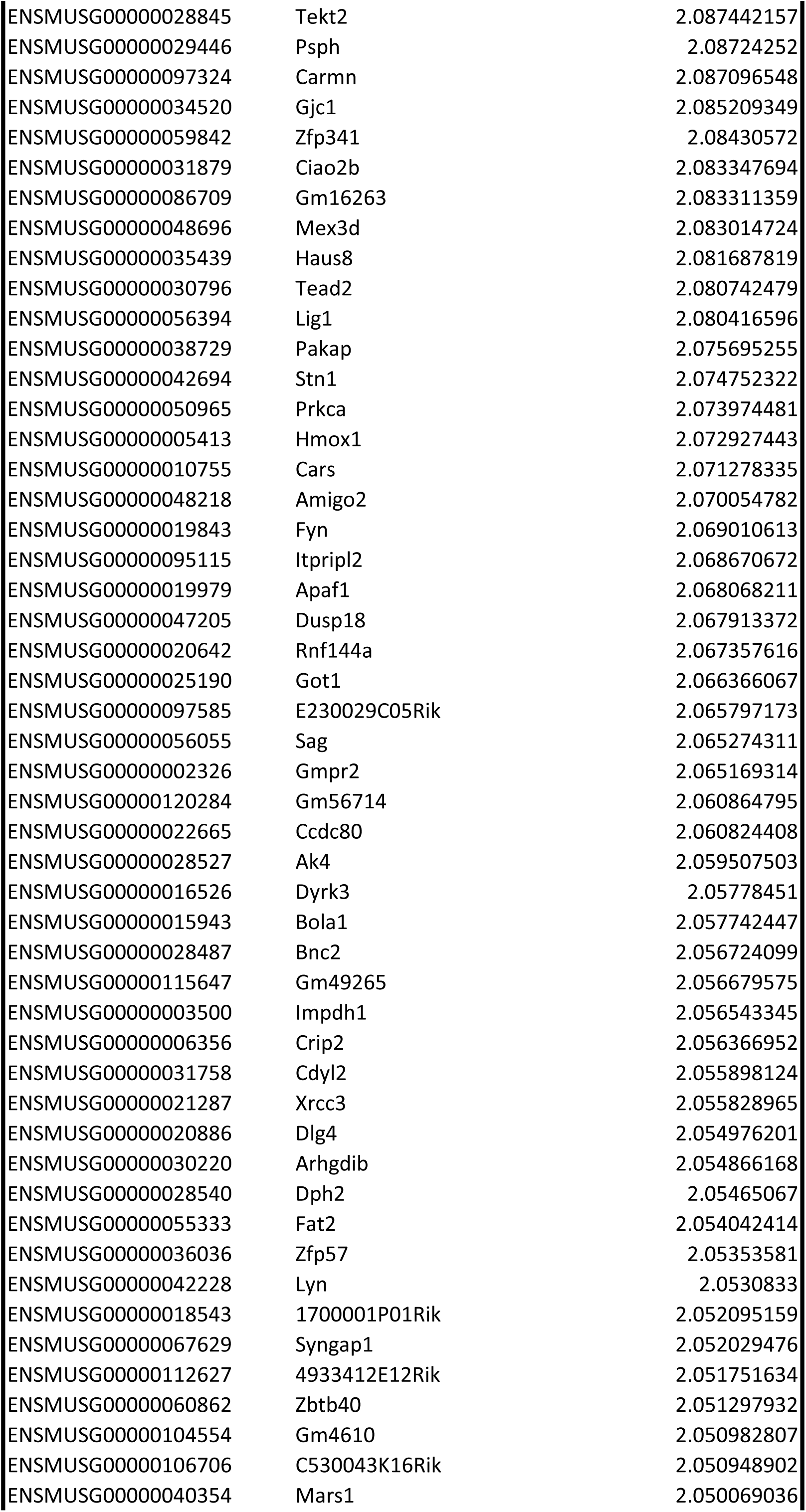
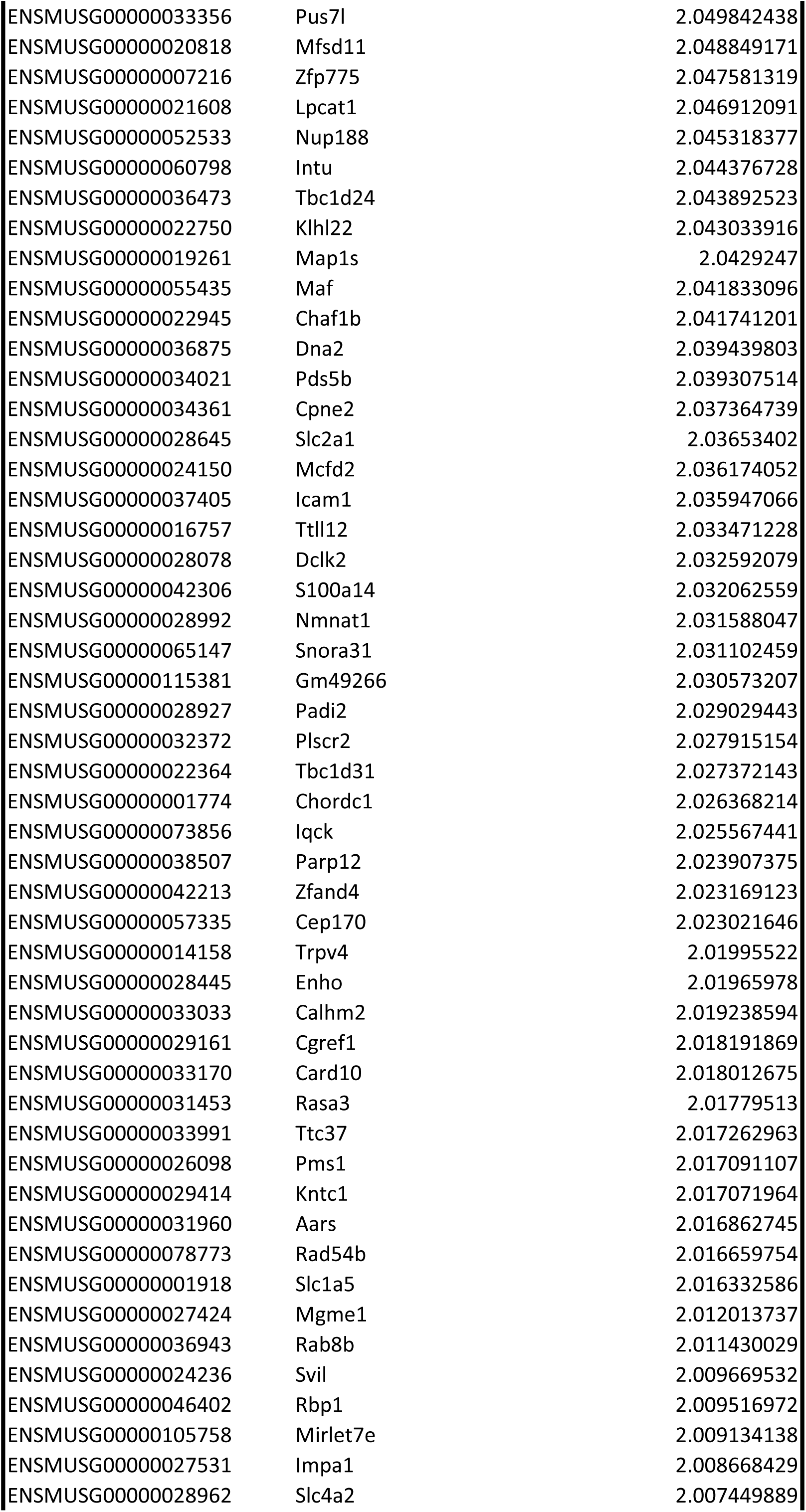
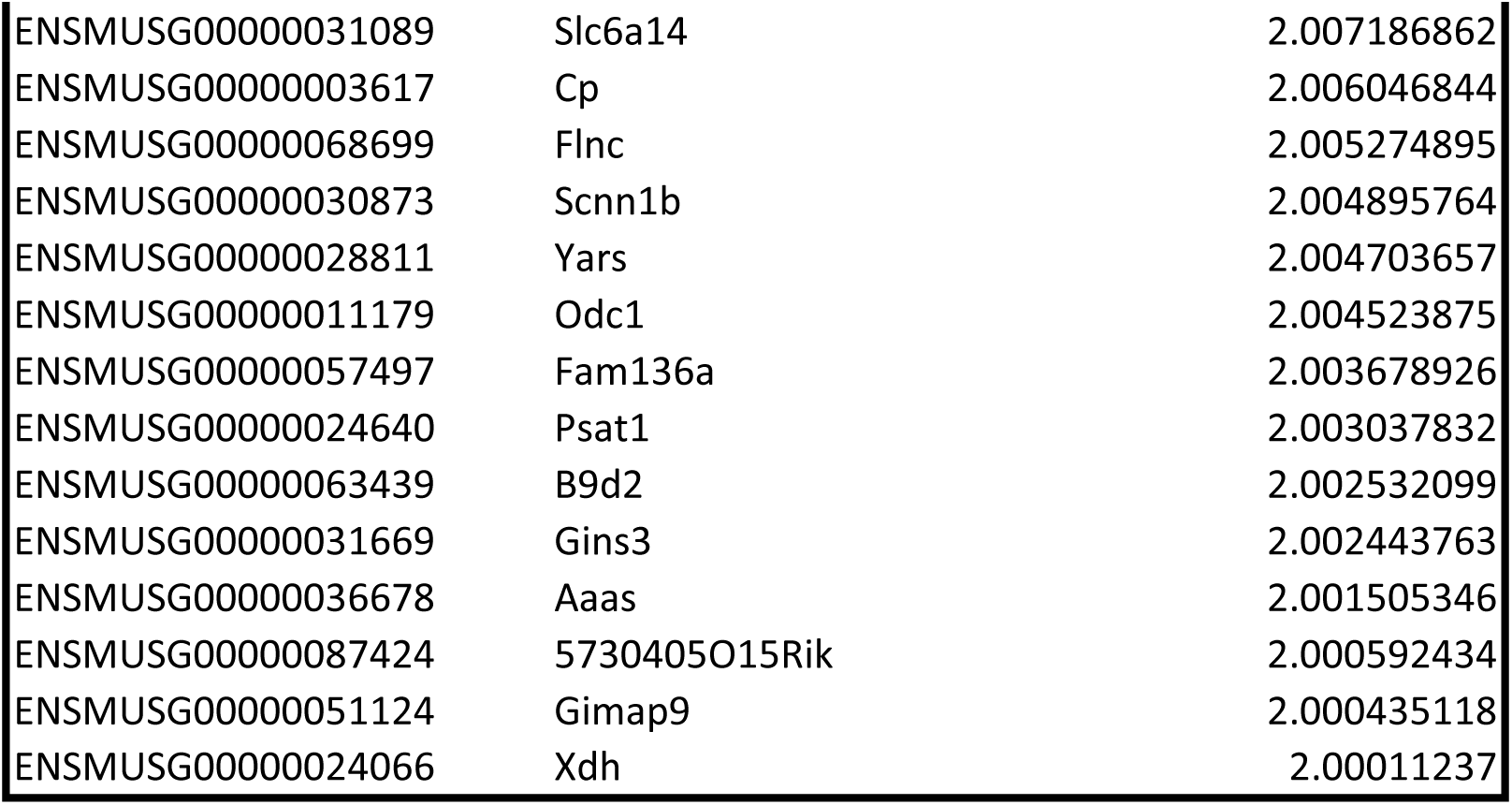
Overview of the significant DEG found in the wounded eye only. Significant genes with a p-adj <0.05 and a fold change lower than 0.5 (downregulated) or higher than 2 (upregulated) were isolated from RNA-Seq data at 18H post-abrasion. For each gene, the ensembl gene ID, the associated gene name and the fold change in the wounded eye are given.

**Table S7.**
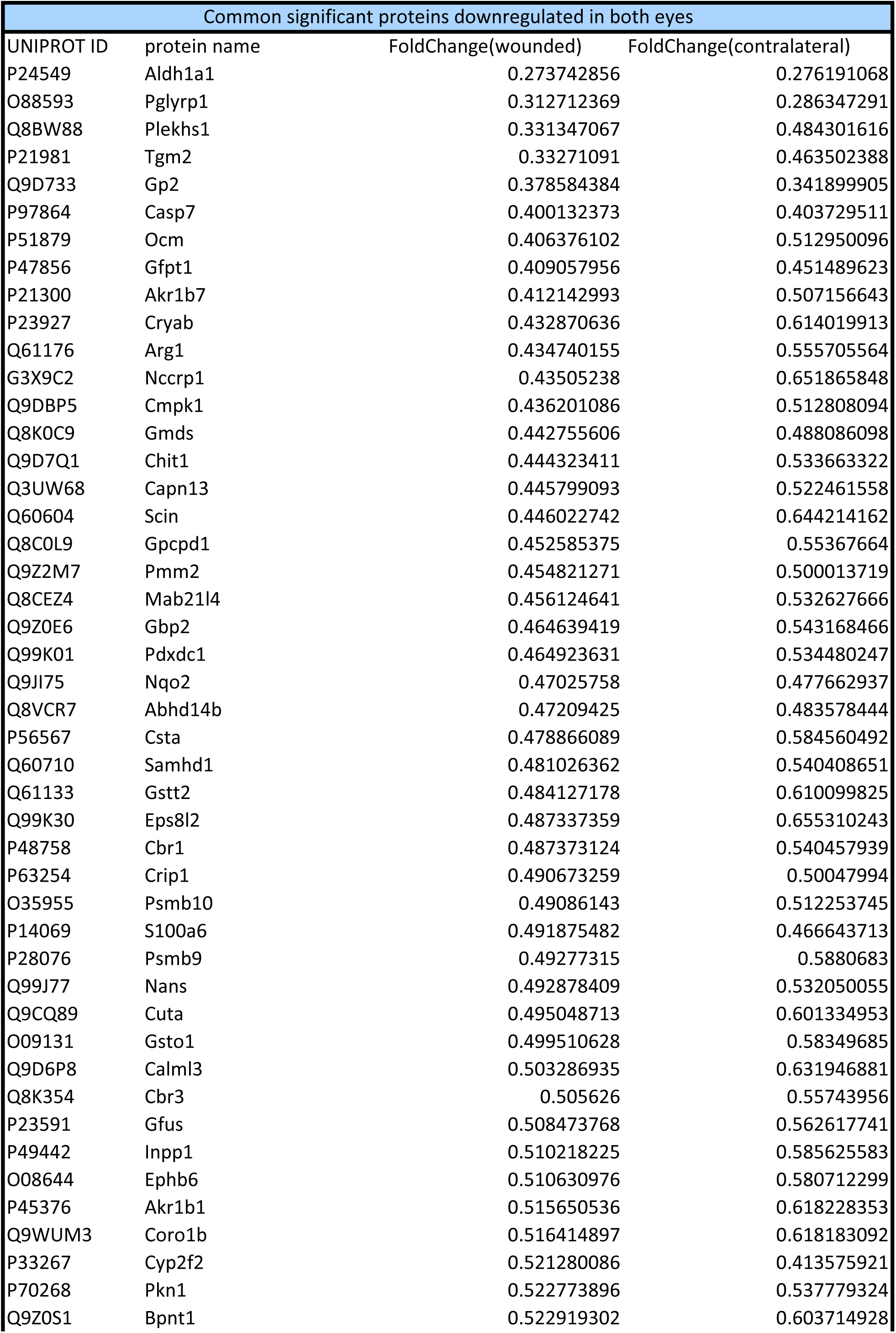
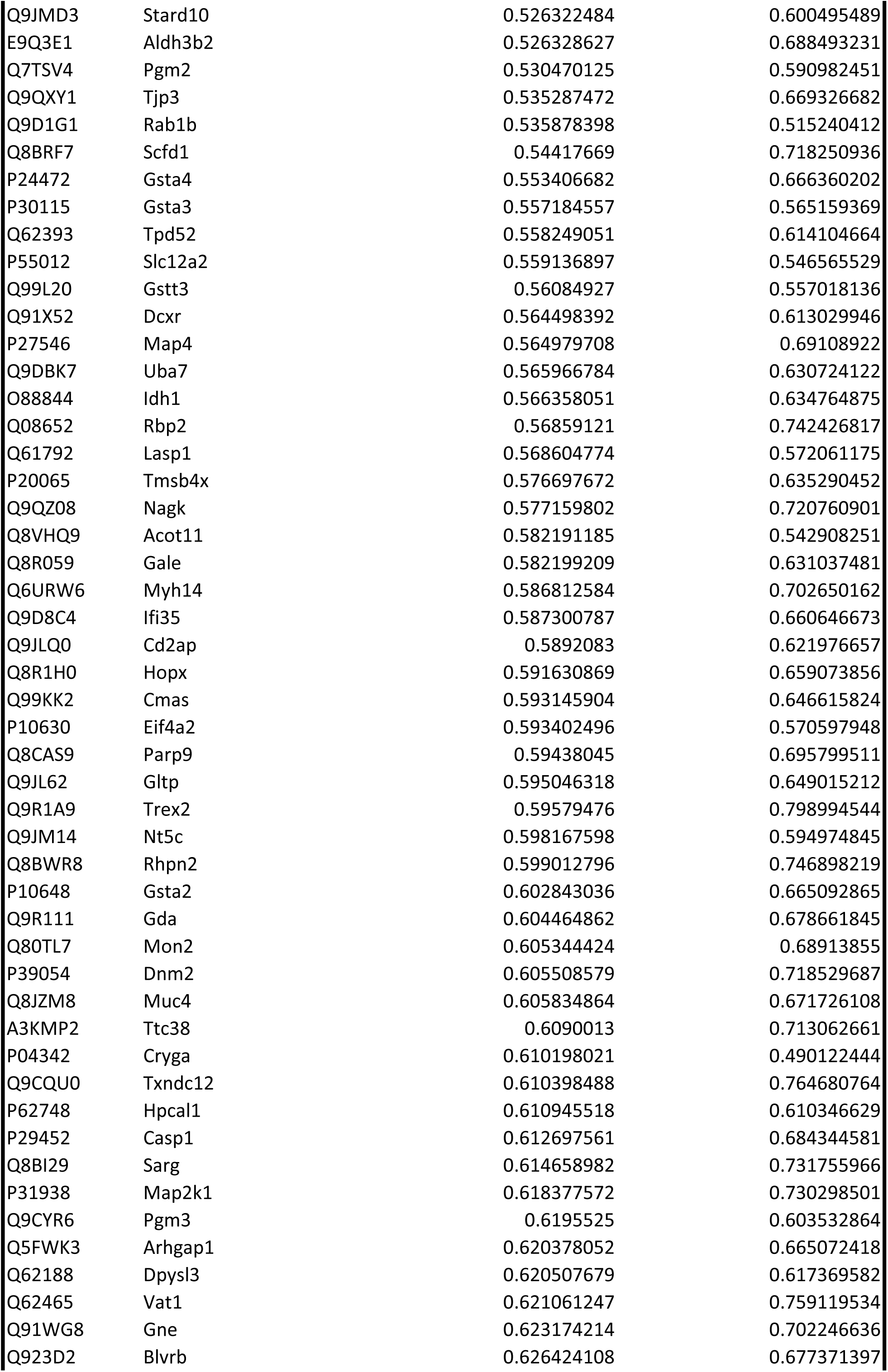
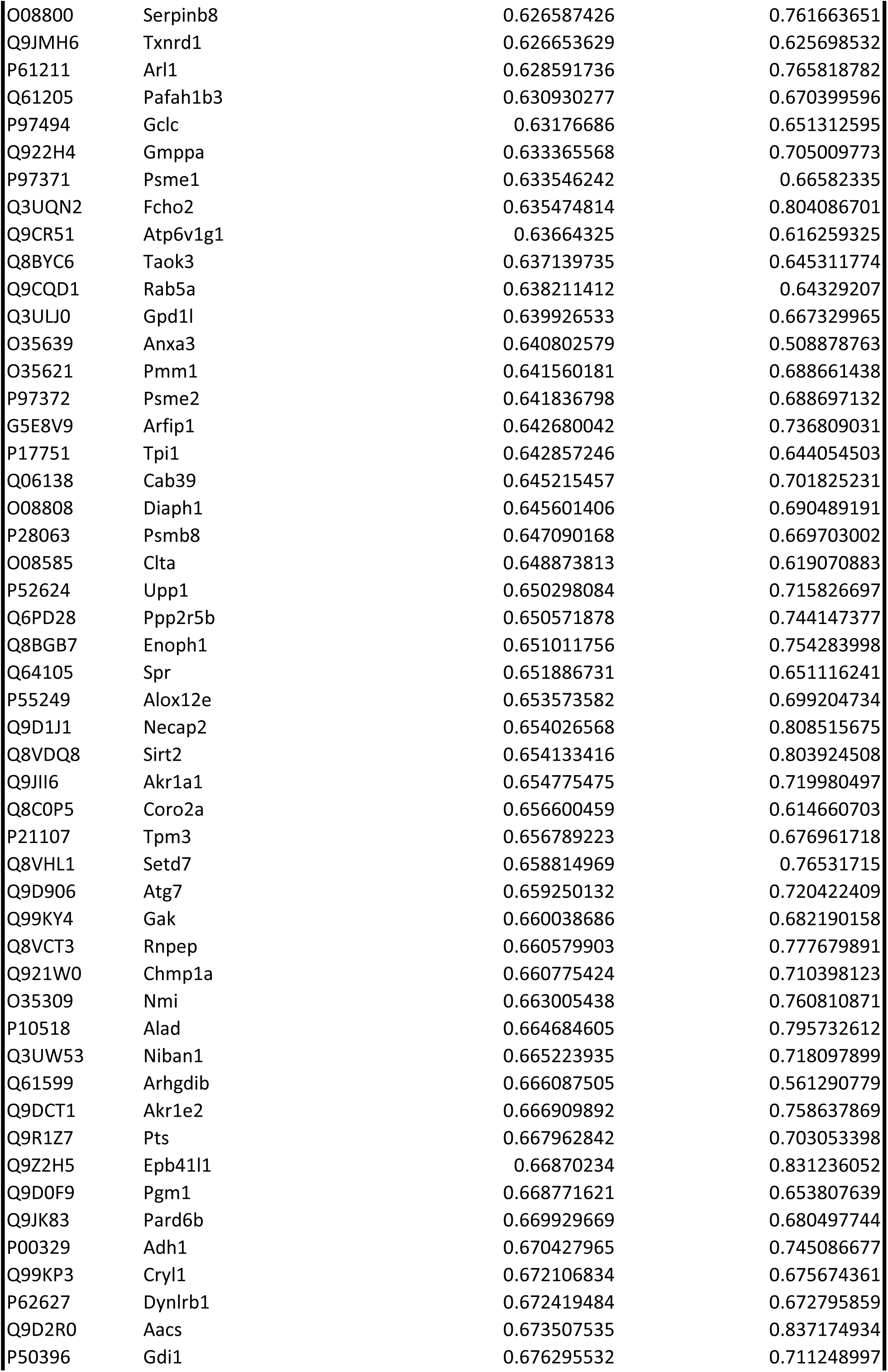
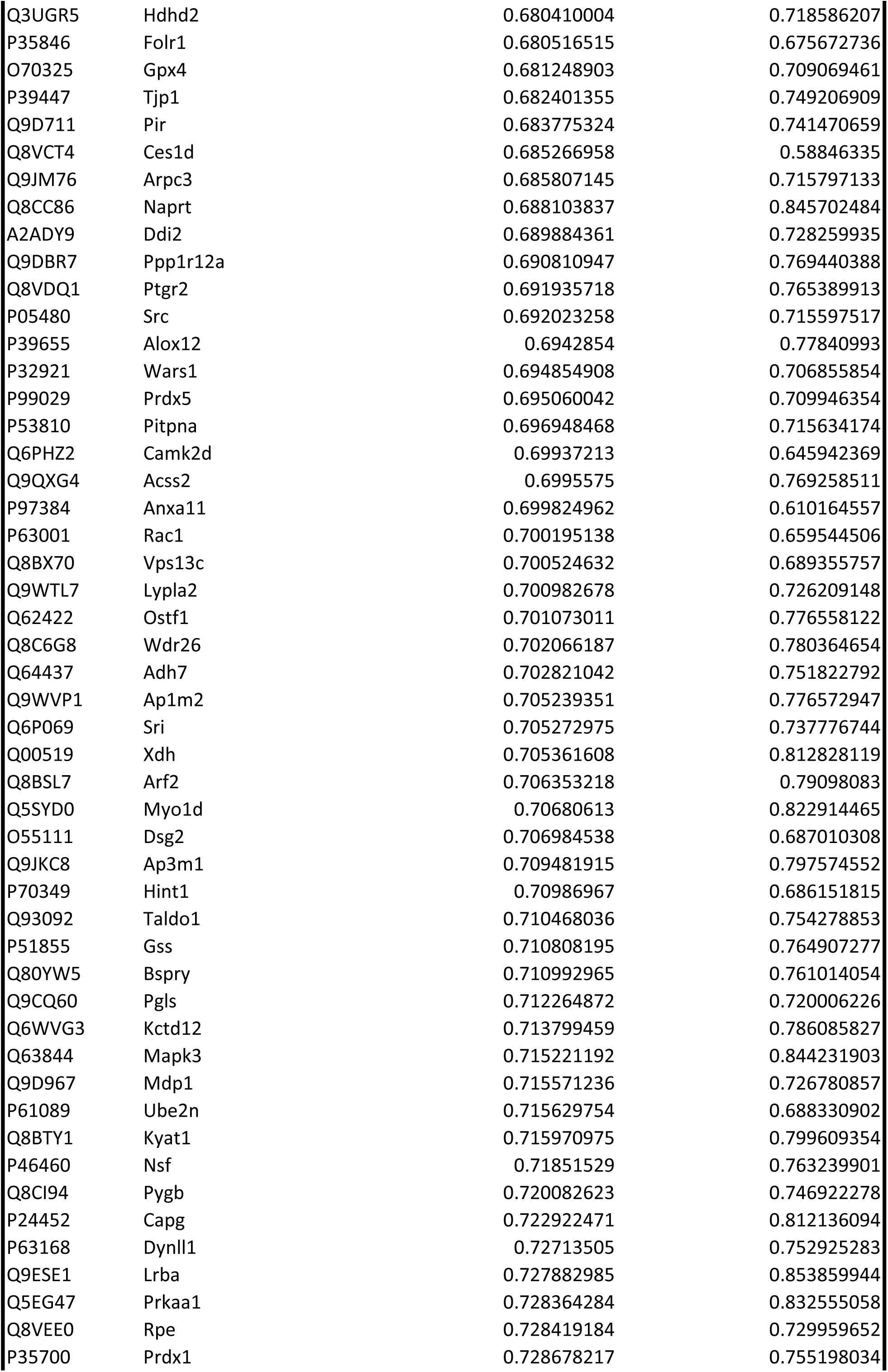
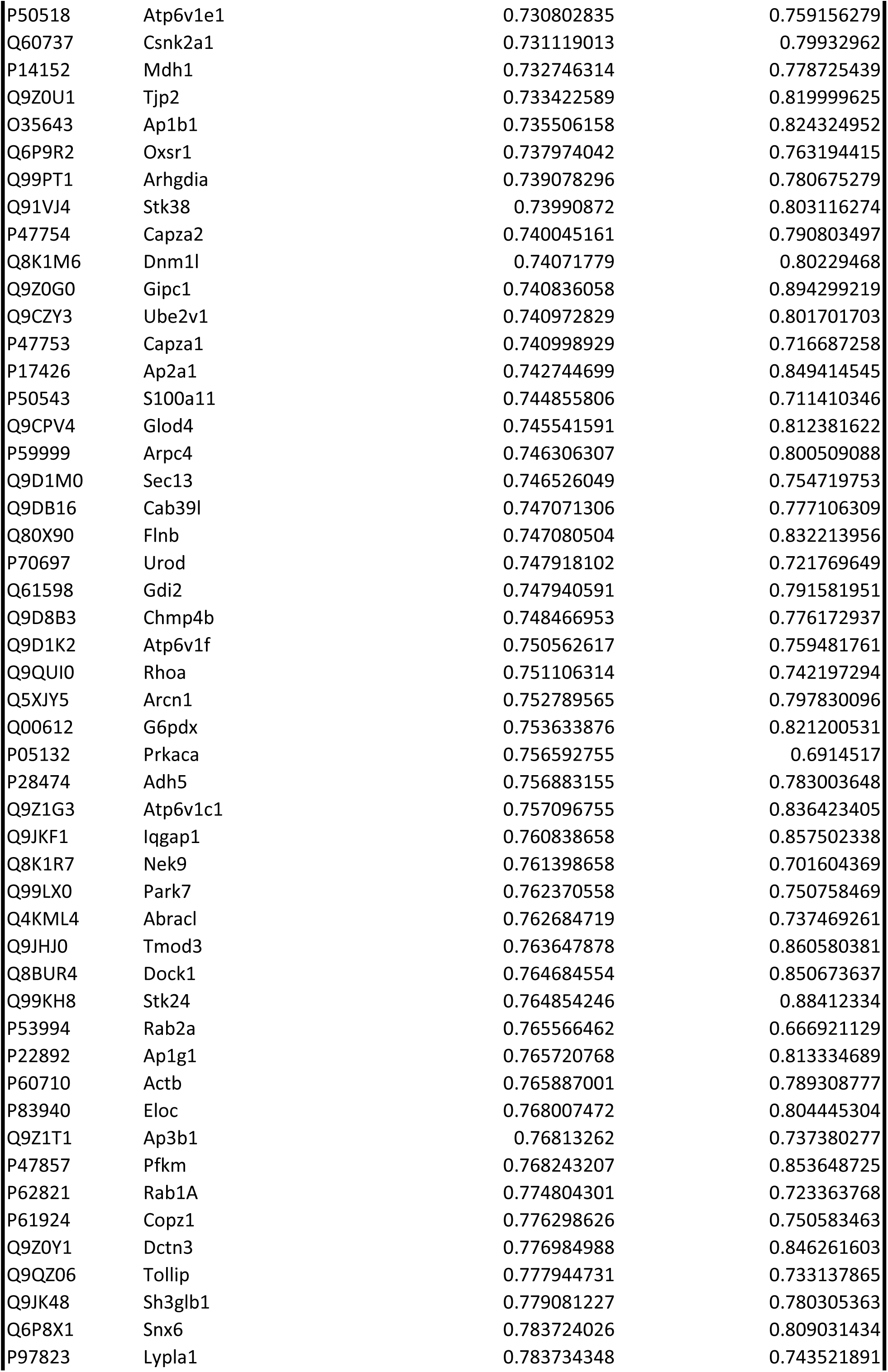
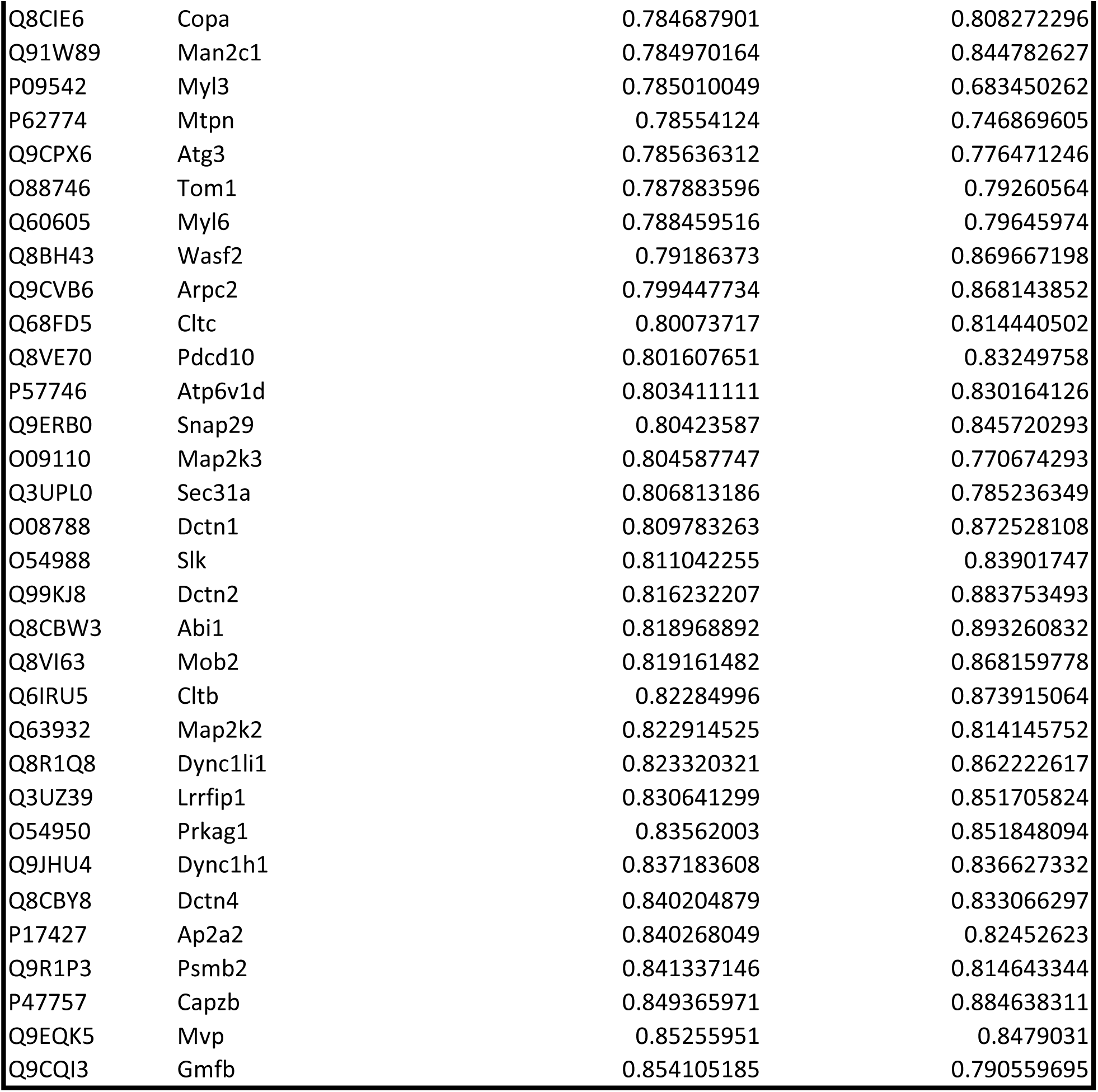
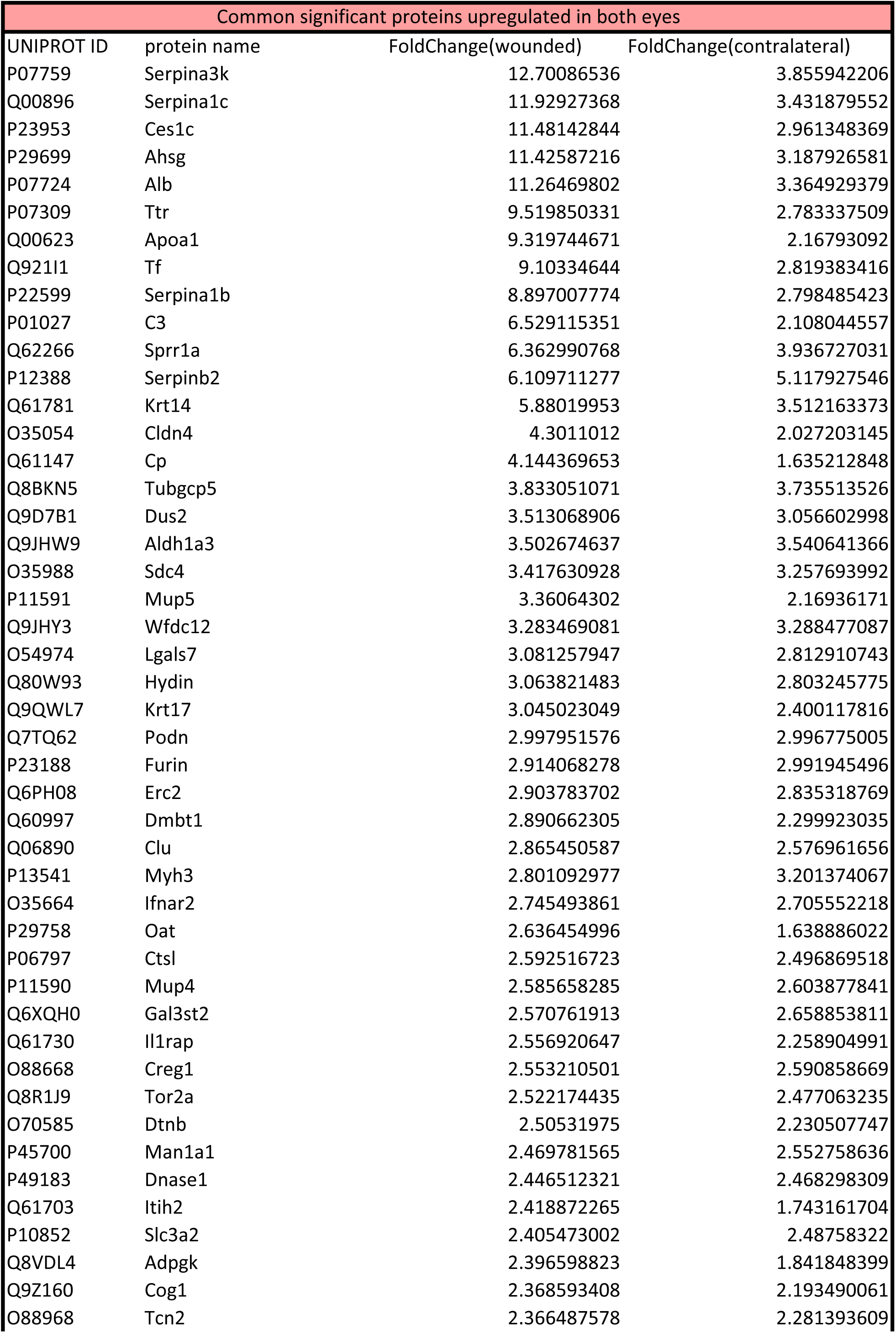
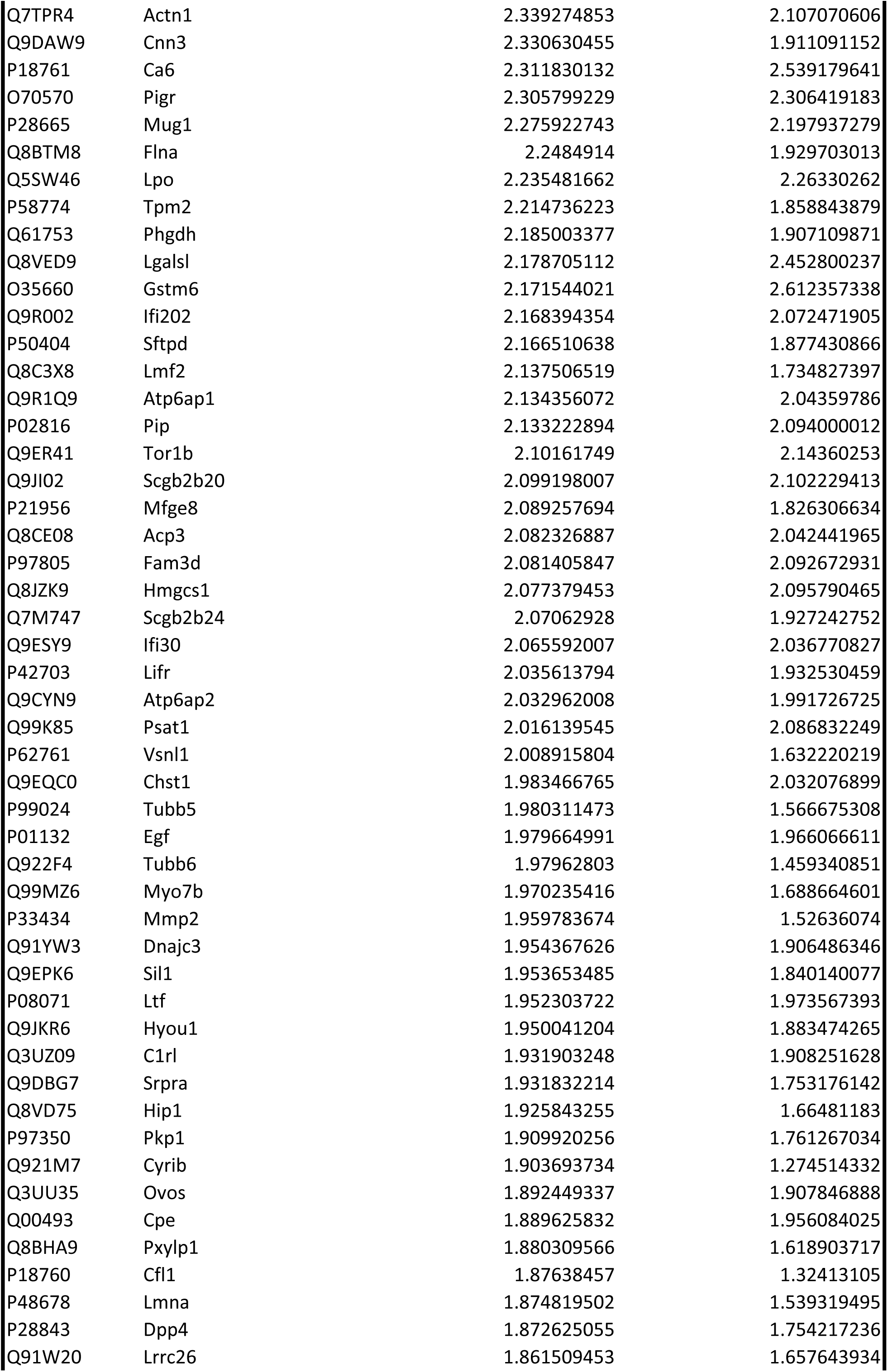
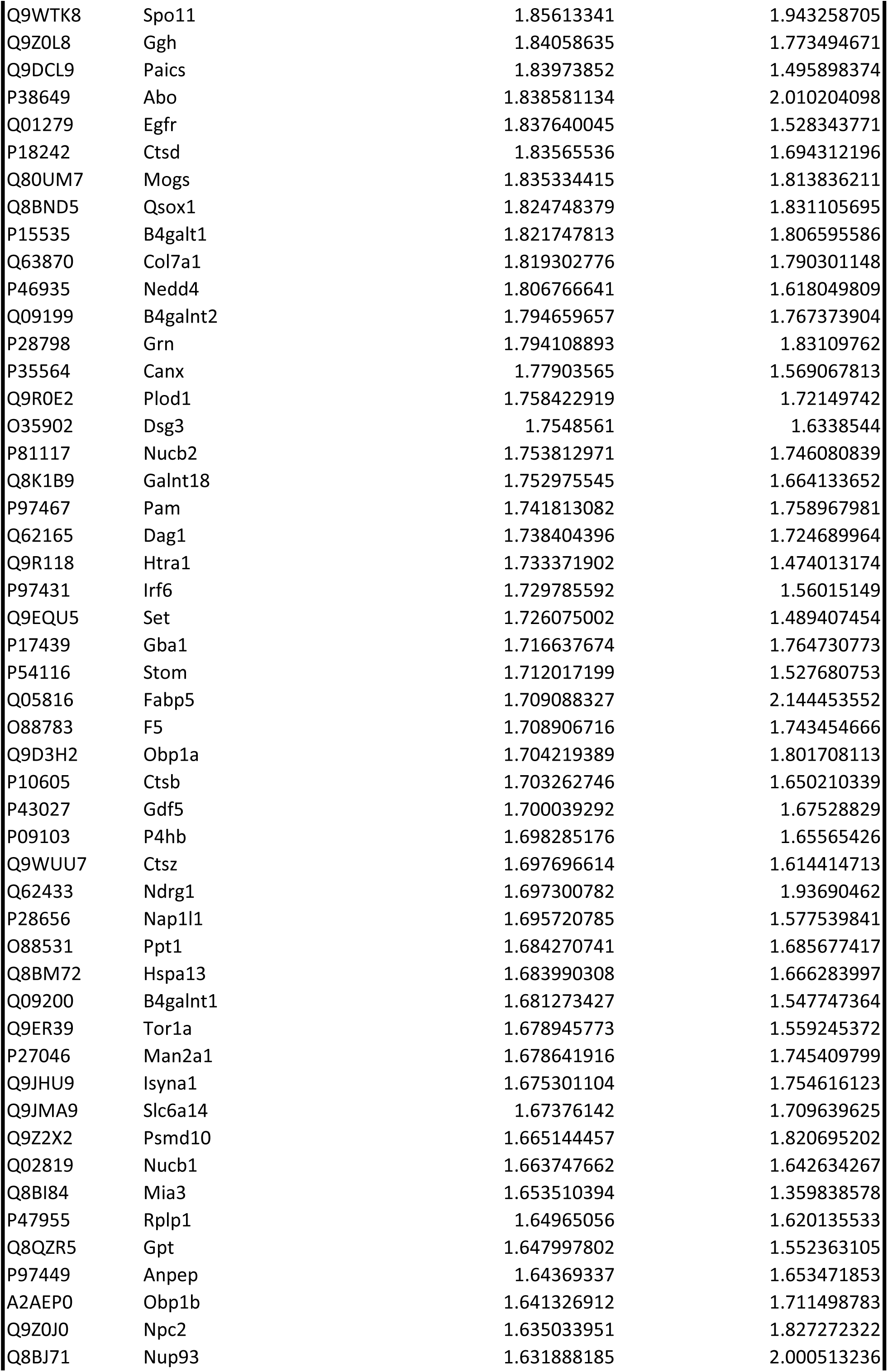
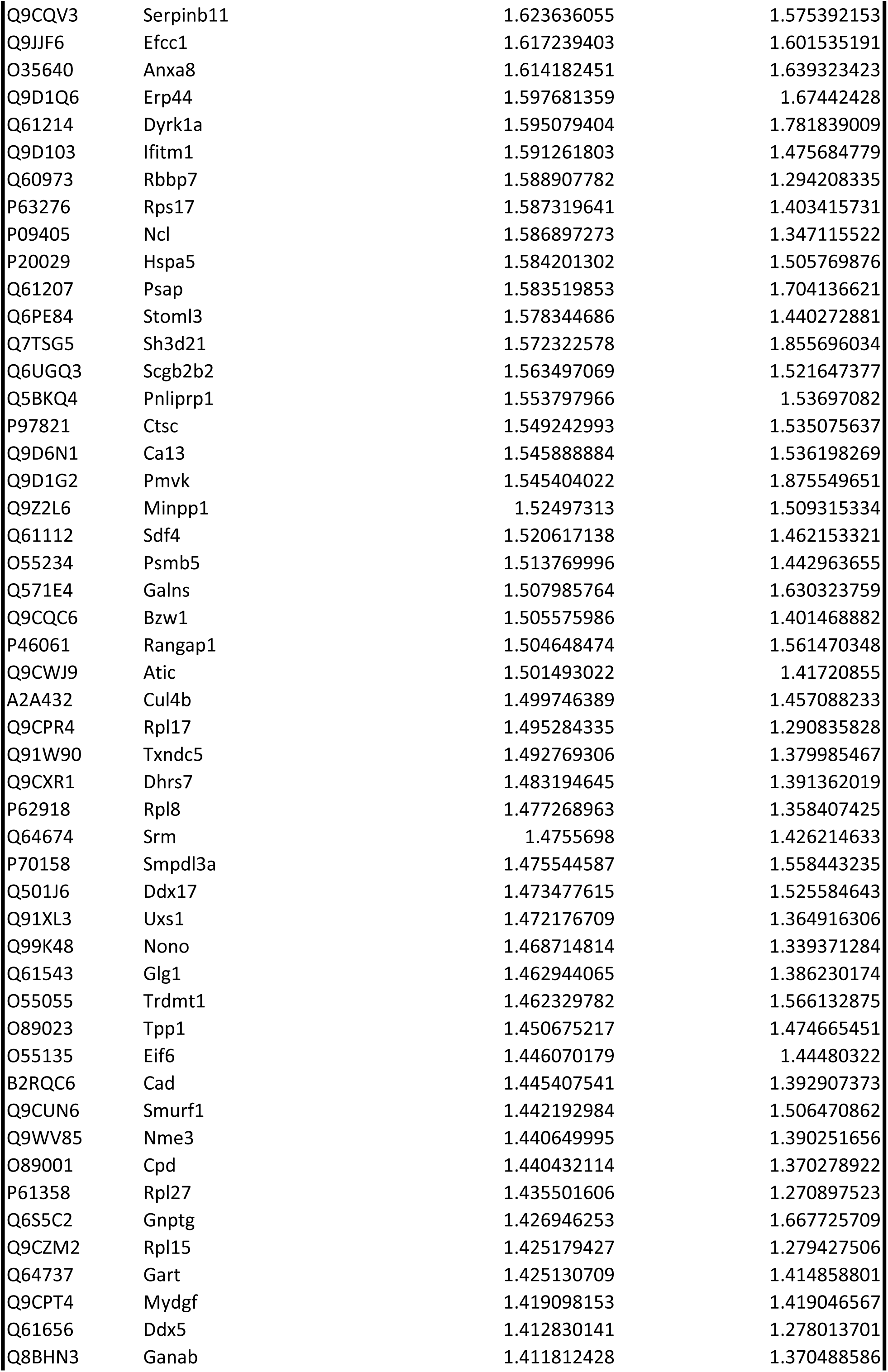
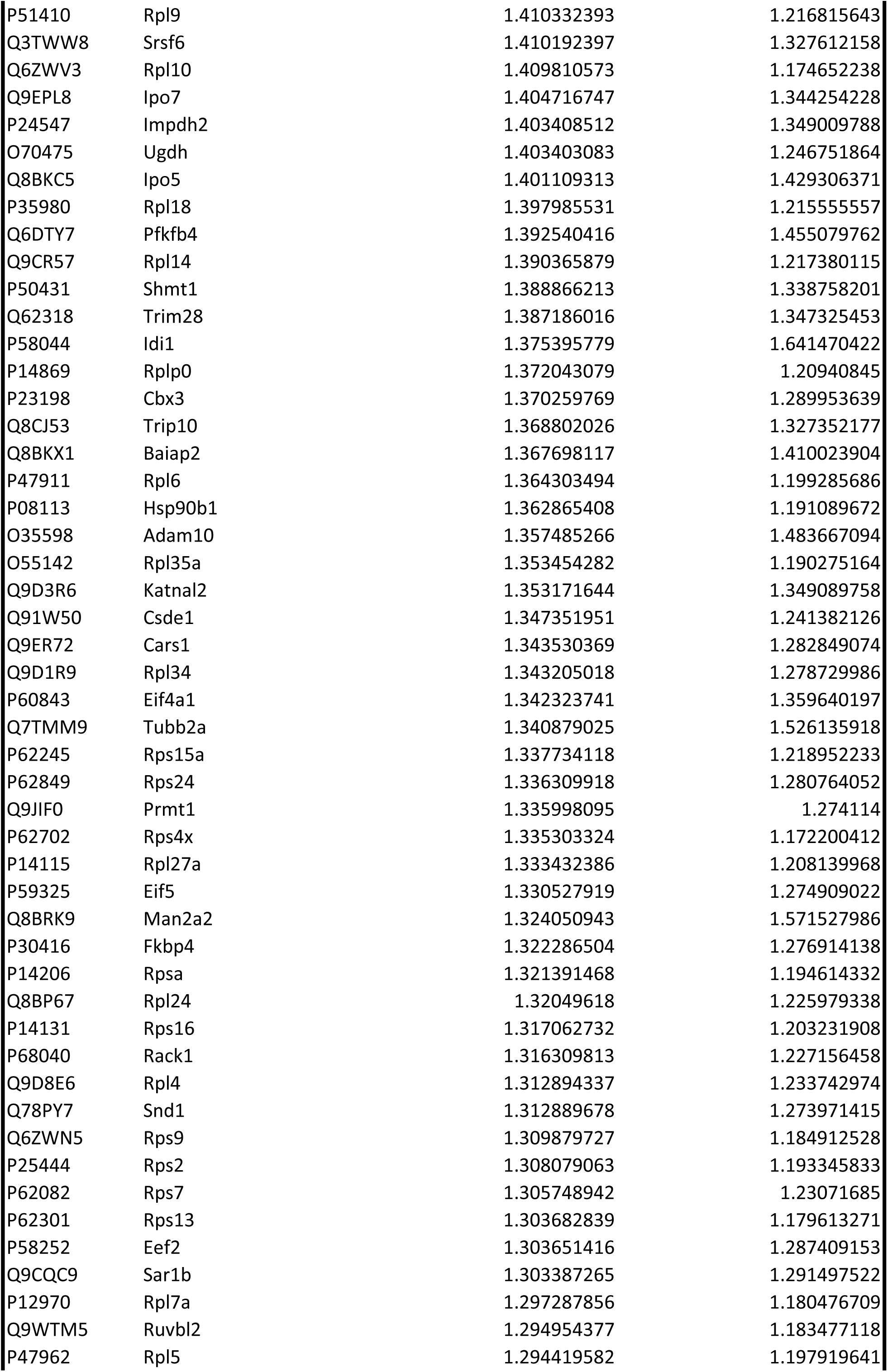
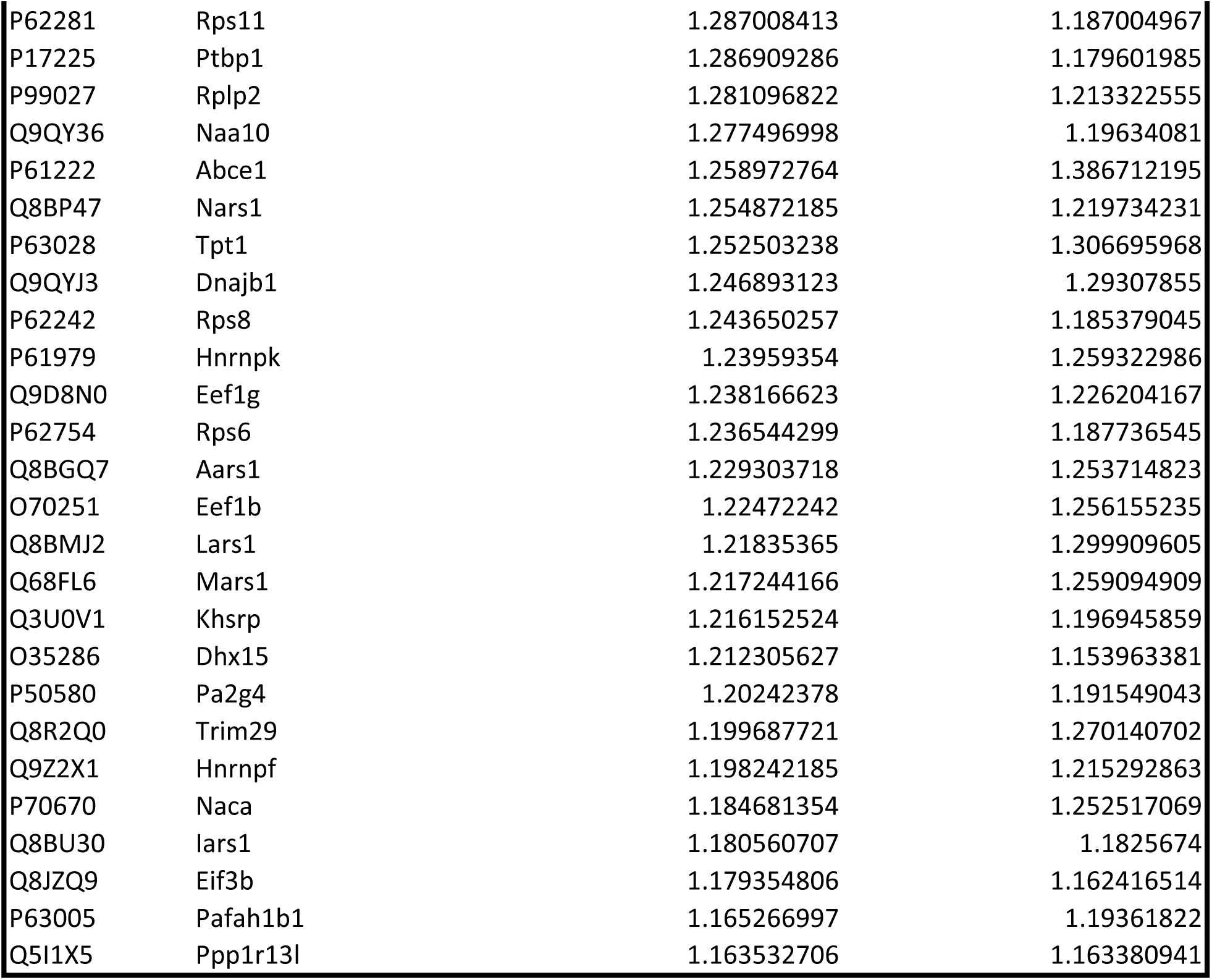
Overview of the common proteins found in both eyes. Significant proteins with a p-adj <0.05 were isolated from proteomics data, regardless of the fold change, at 18H post-abrasion. For each protein, the UniProt ID, the associated protein name and the fold changes in both eyes are given.

**Table S8.**
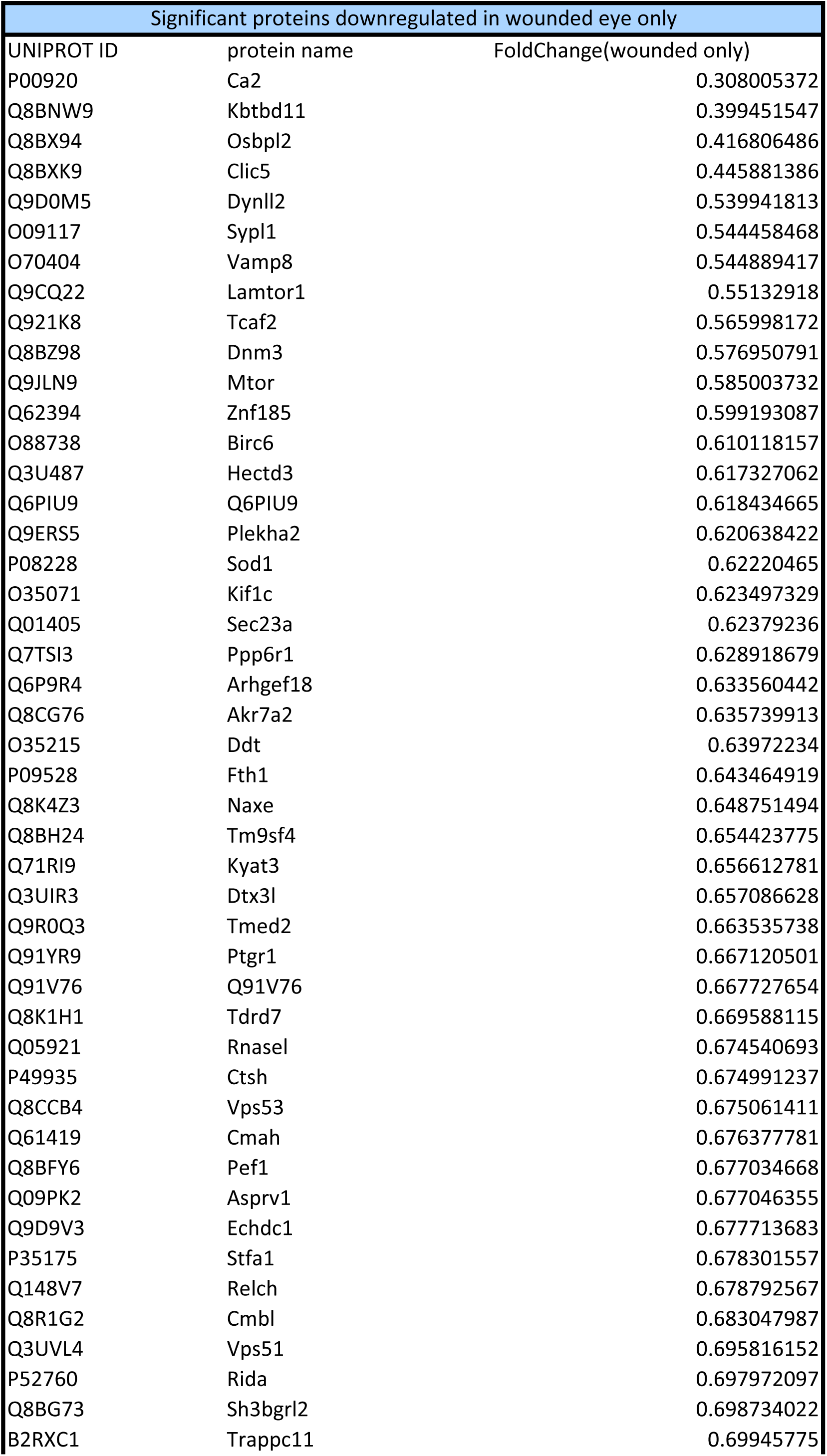
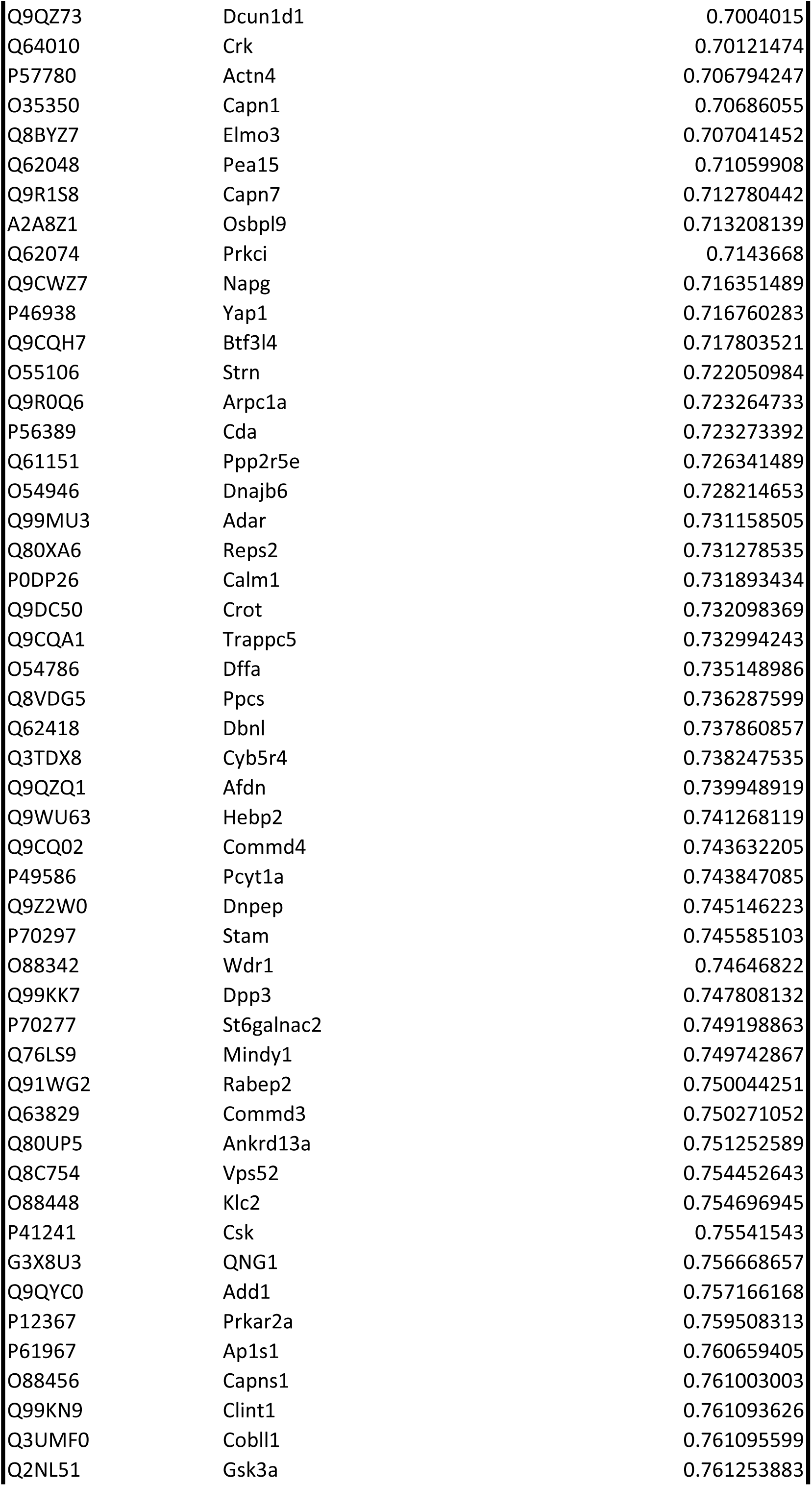
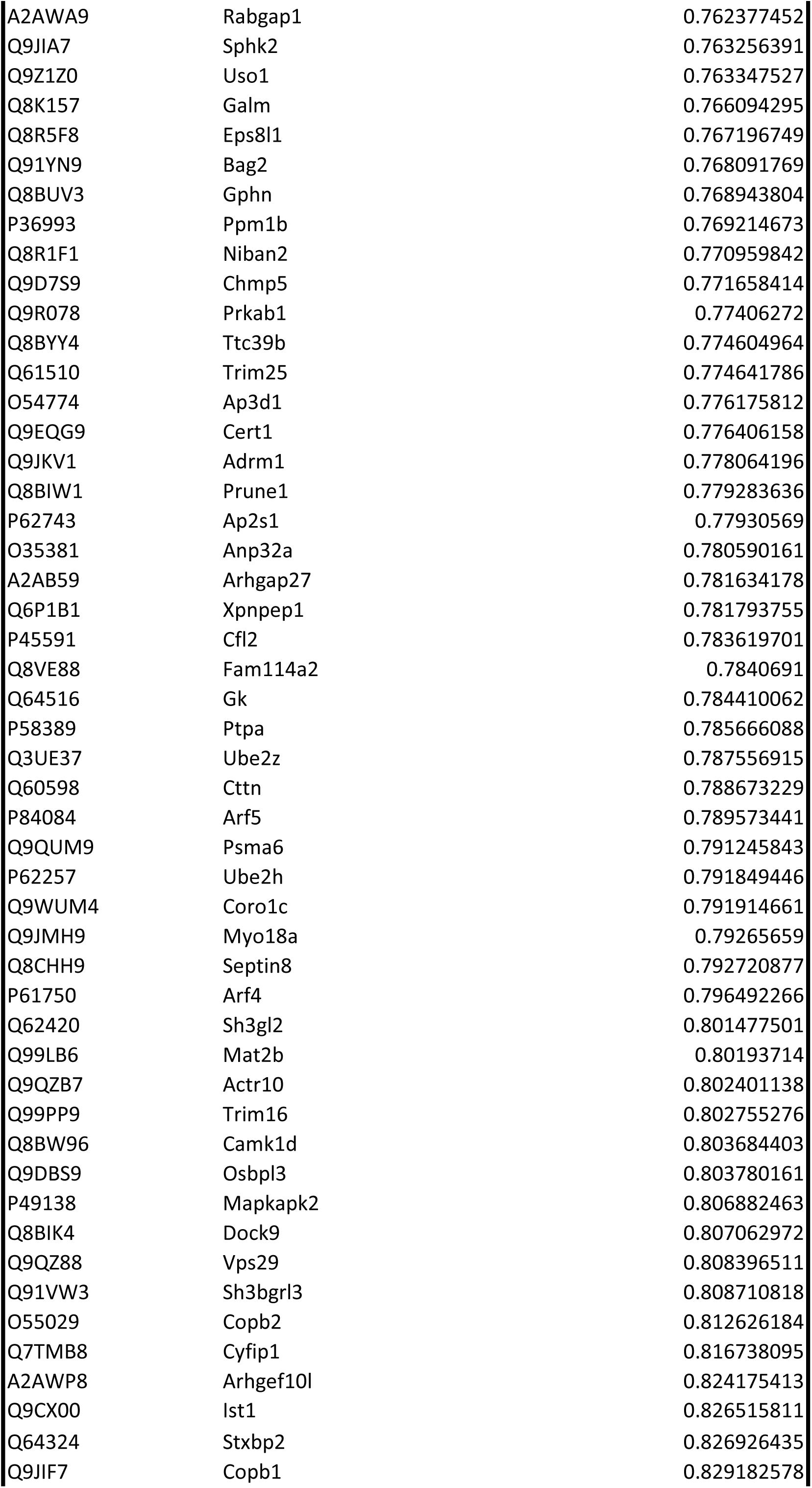
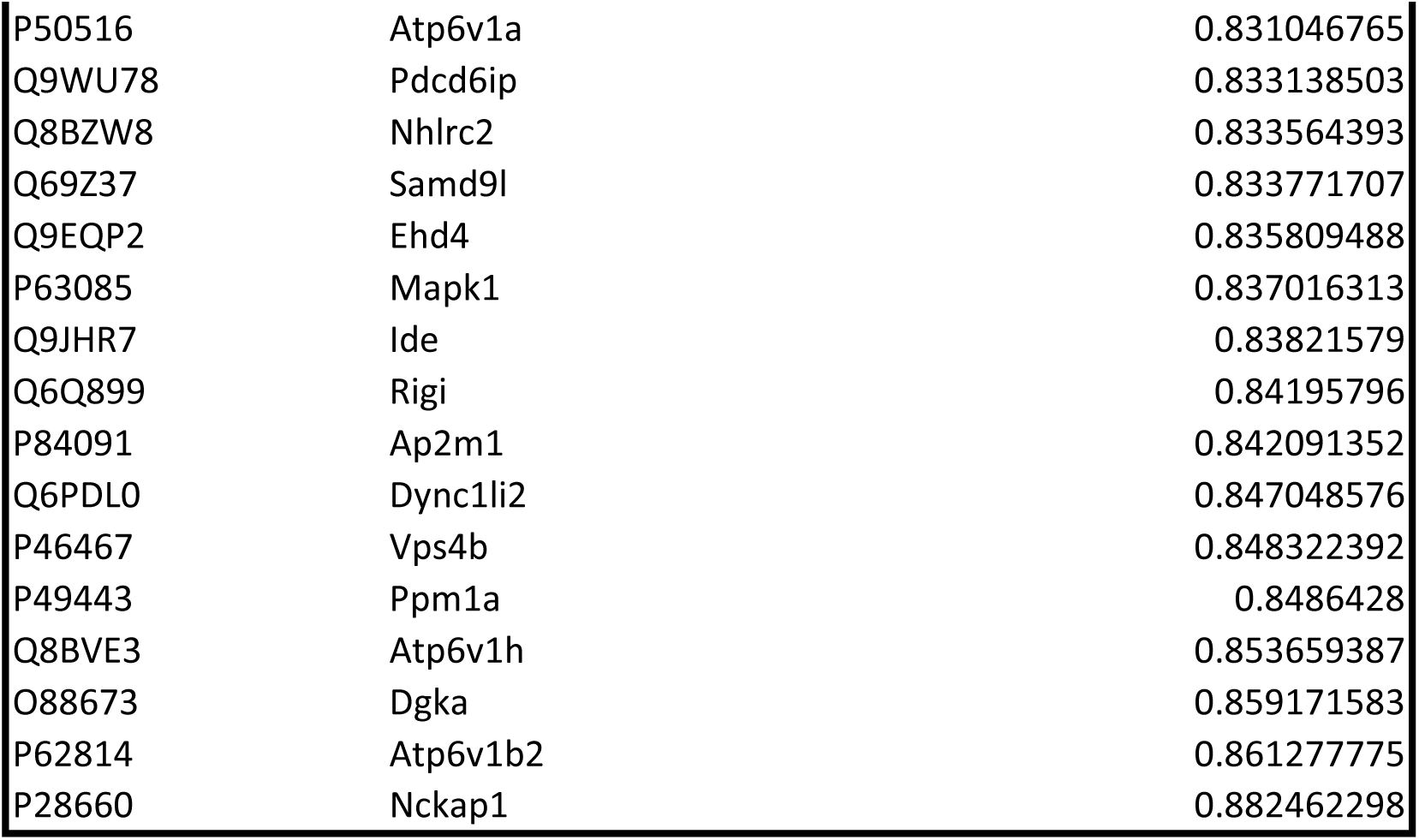
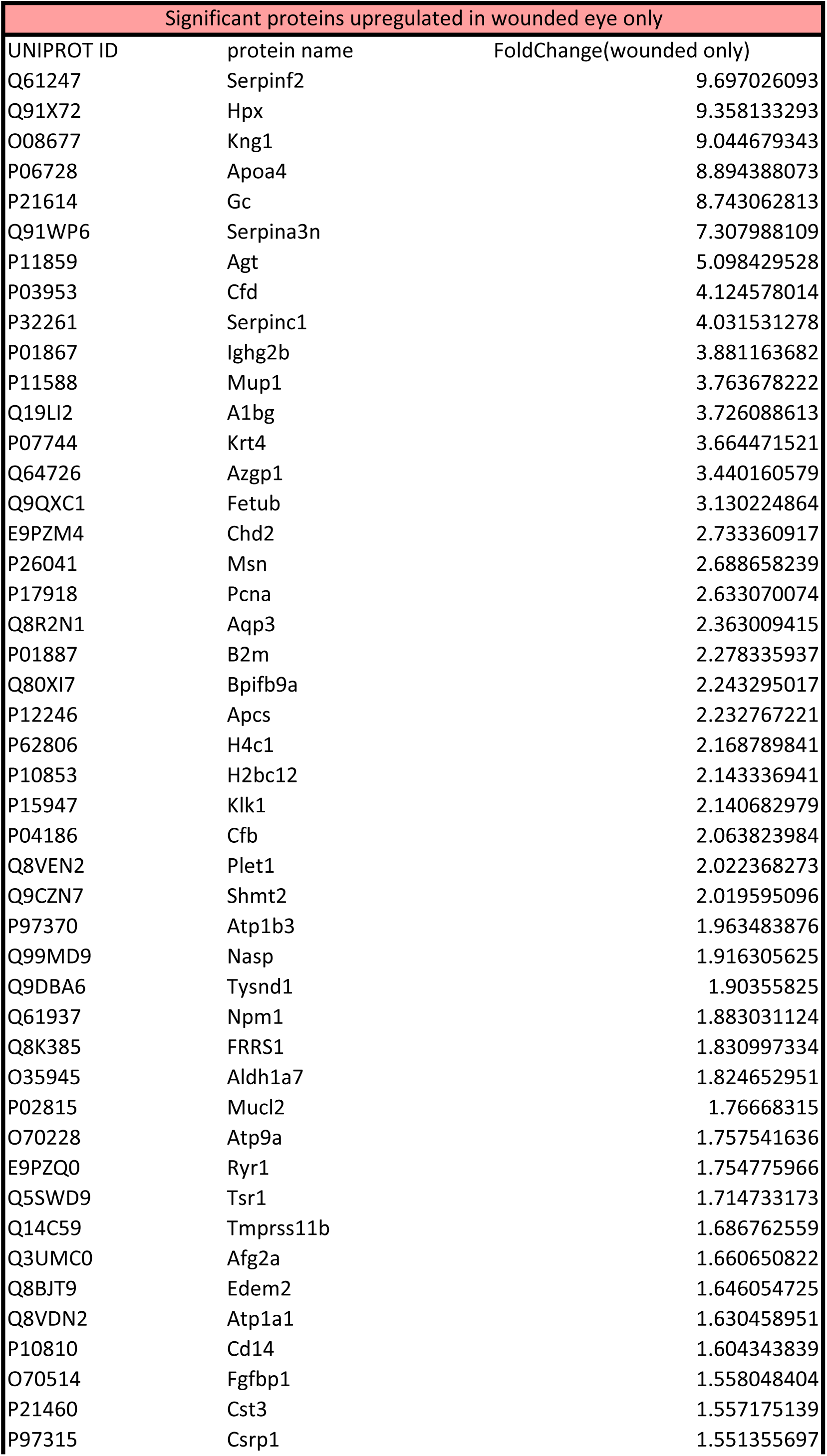
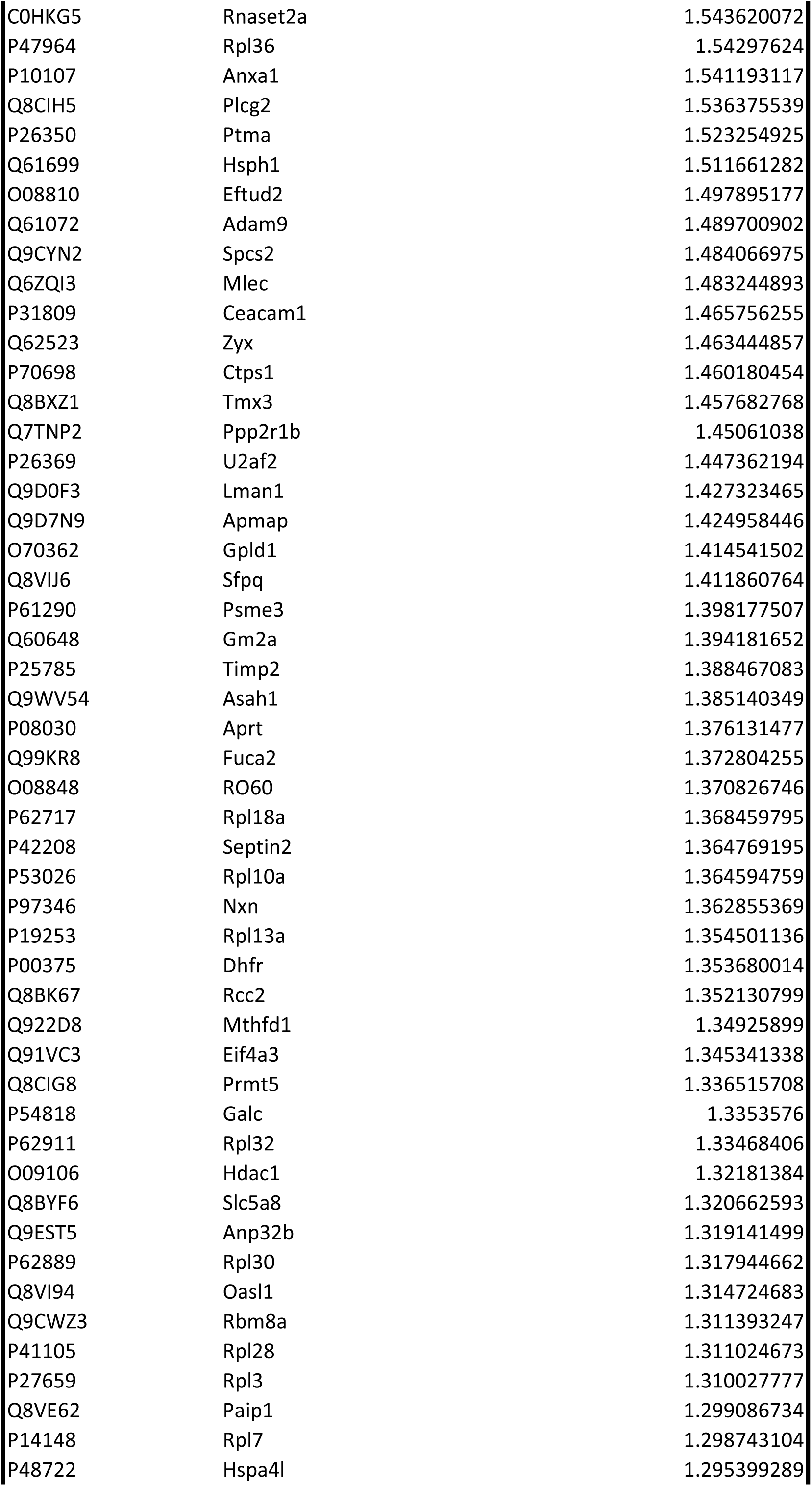
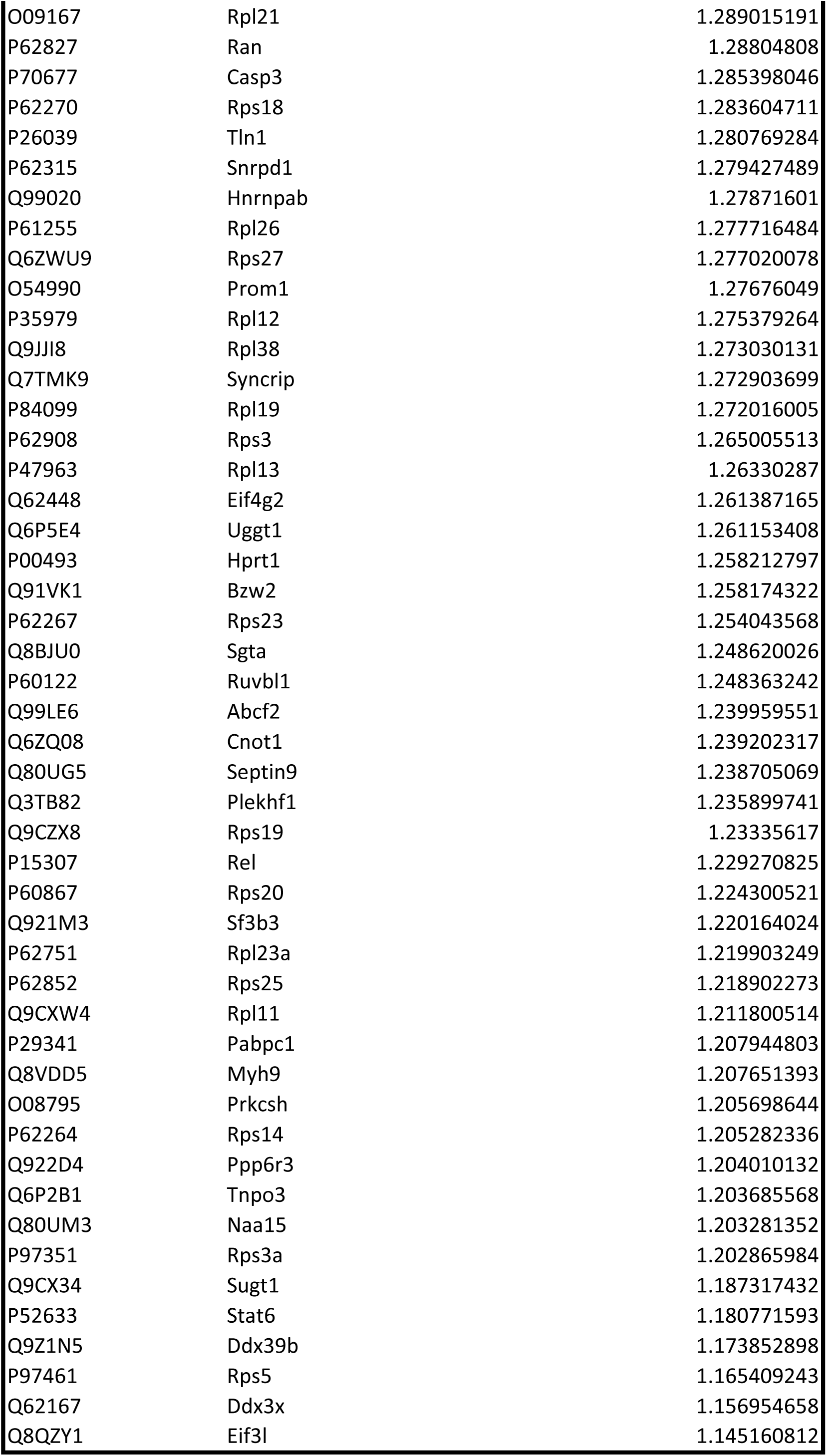
Overview of the common proteins found in the wounded eye only. Significant proteins with a p-adj <0.05 were isolated from proteomics data, regardless of the fold change, at 18H post-abrasion. For each protein, the UniProt ID, the associated protein name and the fold changes in the wounded eye are given.

**Table S9.** ESI source parameters for the detection of nucleosides.

| Parameter | Values |
| --- | --- |
| Nebulizer Gas Flow | 3 L/min |
| Heating Gas Flow | 10 L/min |
| Interface Temperature | 350°C |
| Desolvation Temperature | 602°C |
| DL Temperature | 225°C |
| Heat Block Temperature | 400°C |
| Drying Gas Flow | 3 L/min |
| Interface Voltage | 0.5 kV |
| Focus Voltage | 2 kV |

**Table S10.**
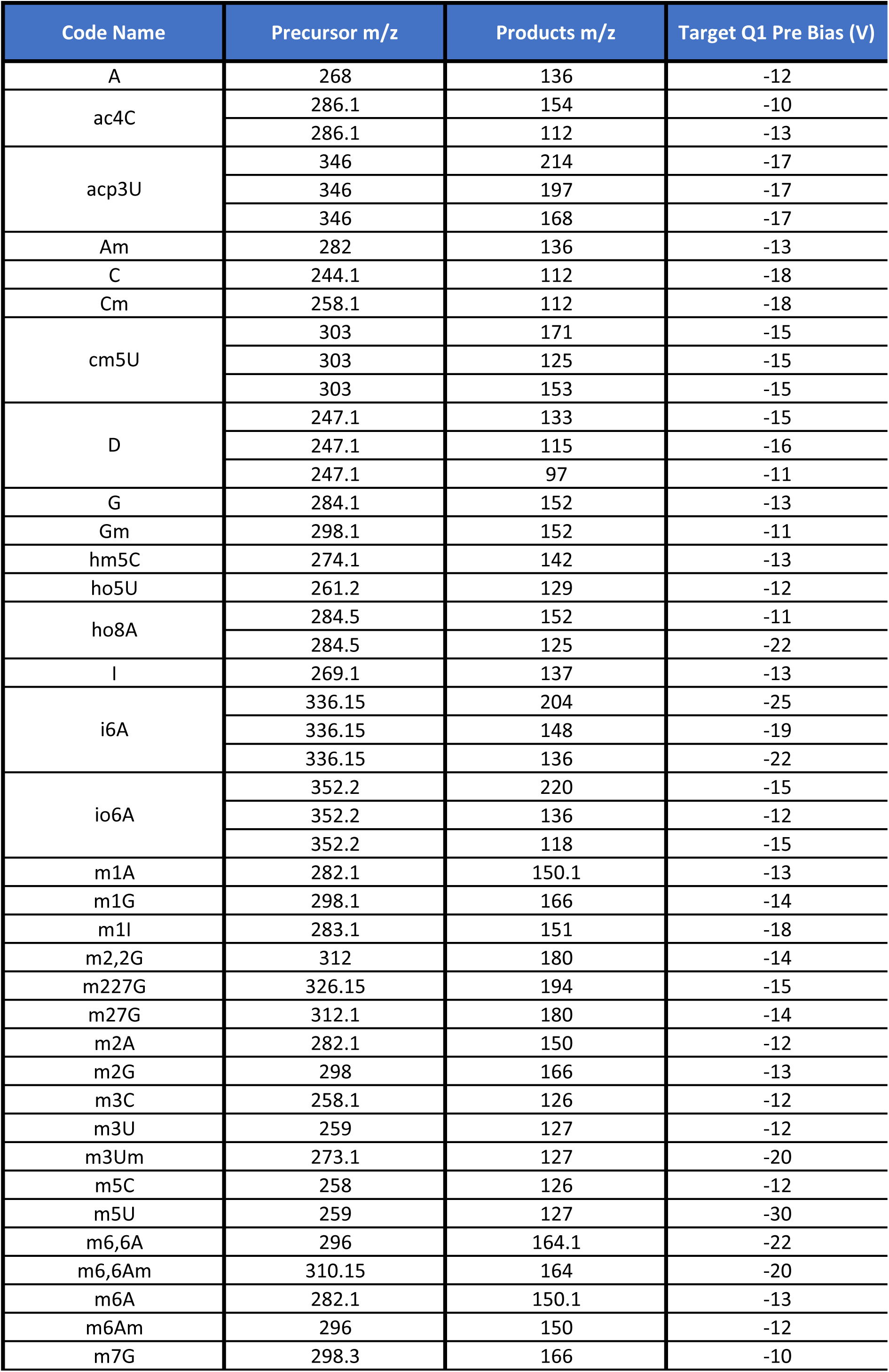
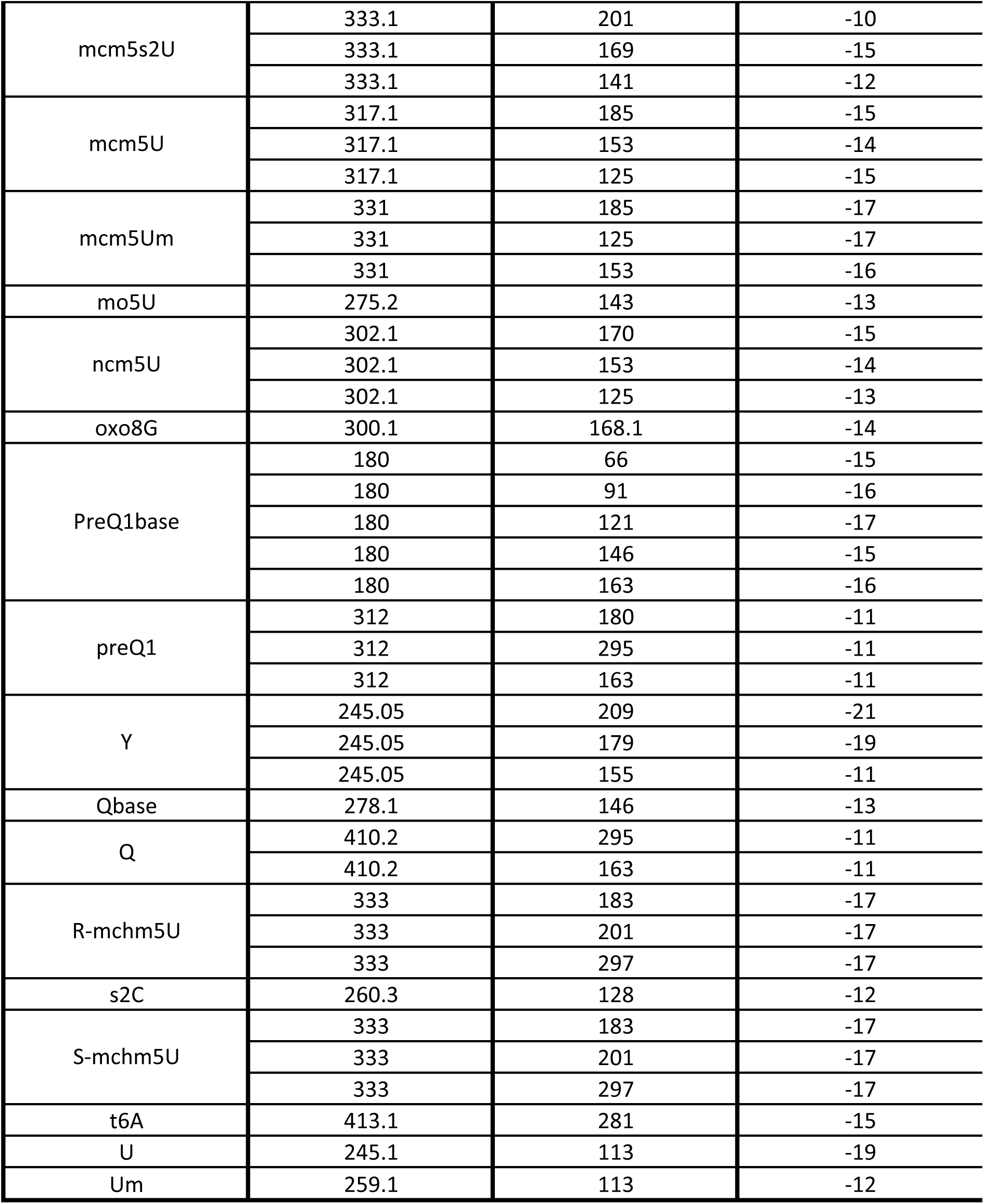
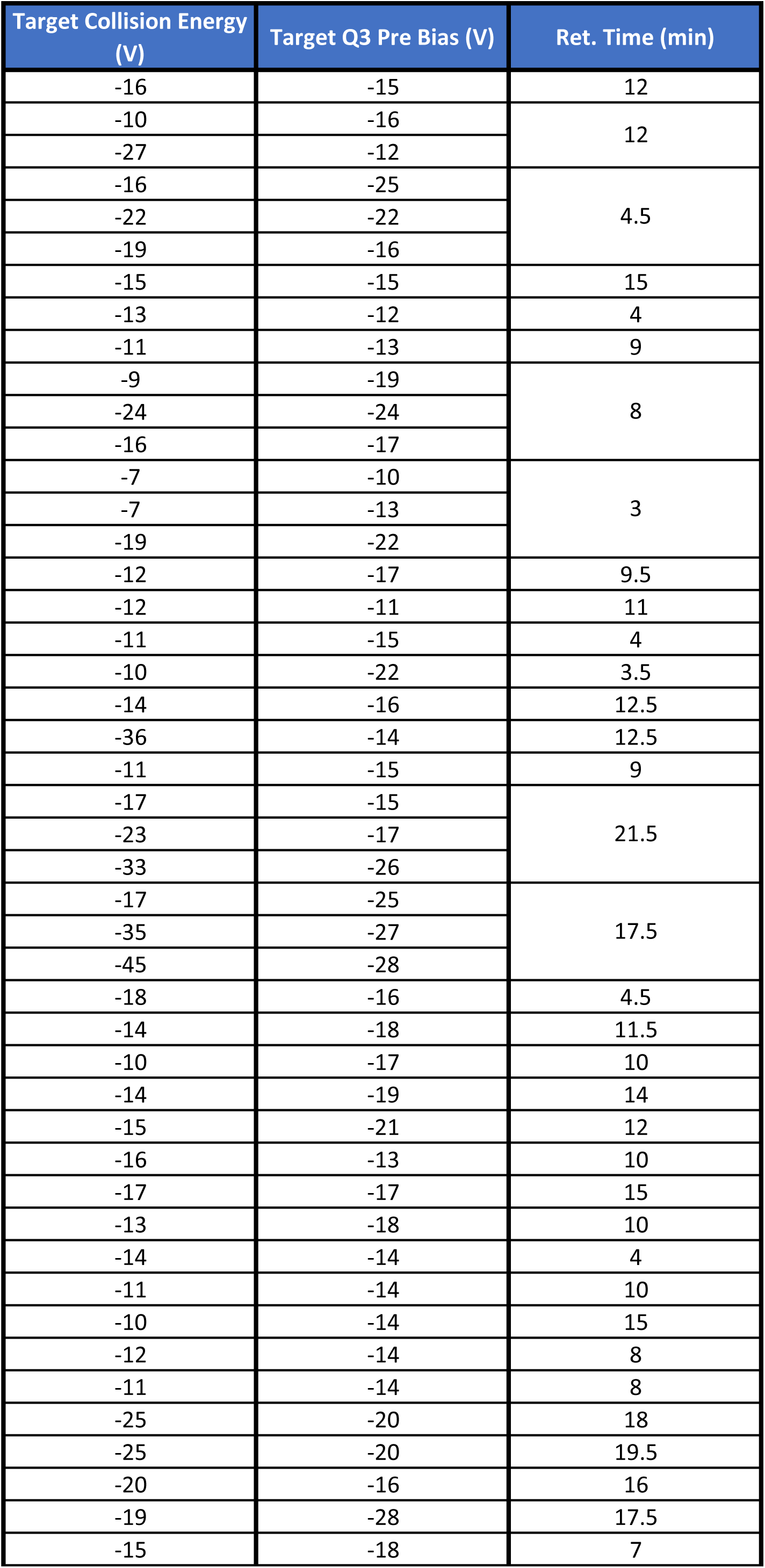
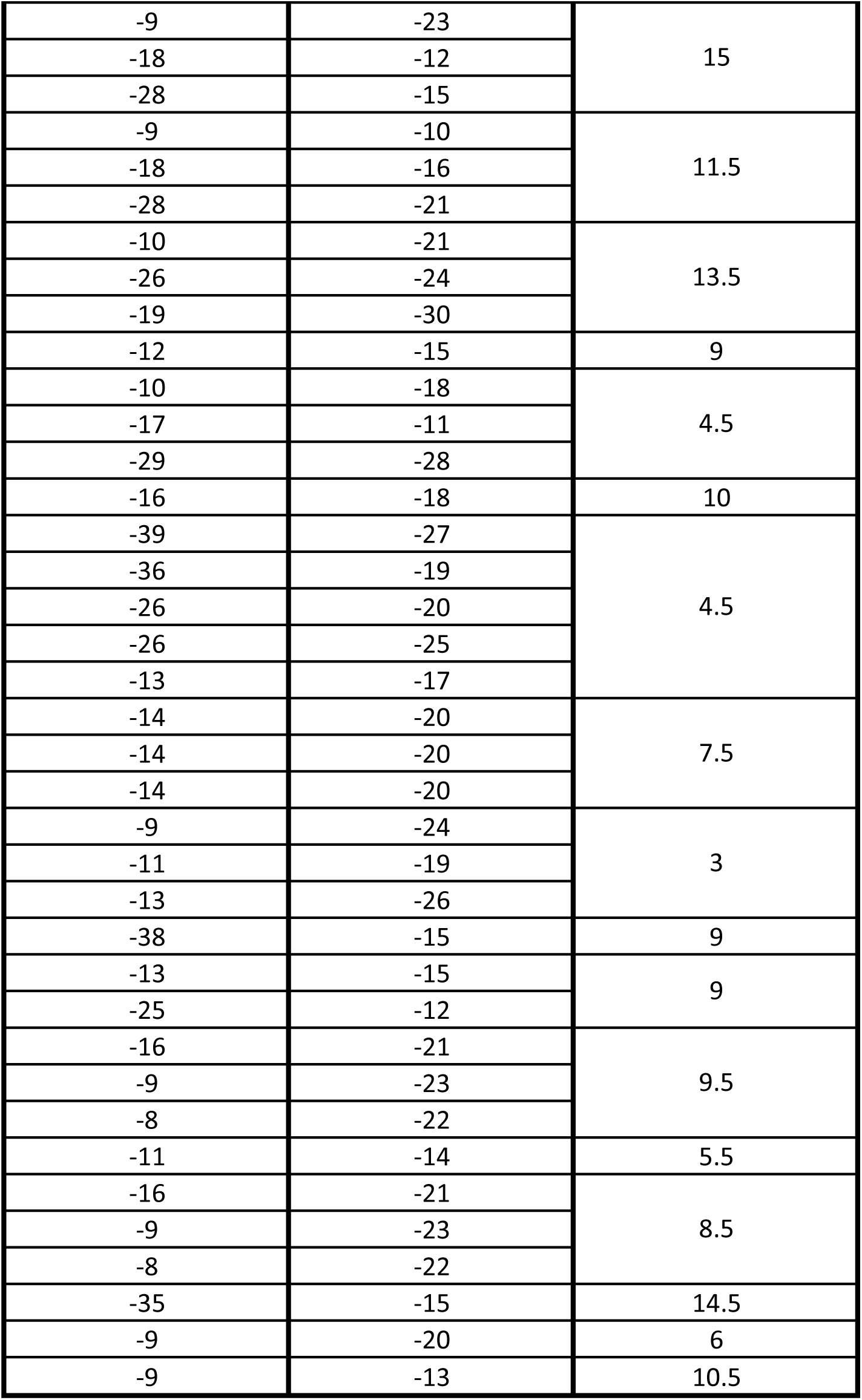
MS/MS parameters for MRM analysis of nucleosides.

| Code Name | Precursor m/z | Products m/z | Target Q1 Pre Bias (V) |
| --- | --- | --- | --- |
| A | 268 | 136 | -12 |
| ac4C | 286.1 | 154 | -10 |
|  | 286.1 | 112 | -13 |
| acp3U | 346 | 214 | -17 |
|  | 346 | 197 | -17 |
|  | 346 | 168 | -17 |
| Am | 282 | 136 | -13 |
| C | 244.1 | 112 | -18 |
| Cm | 258.1 | 112 | -18 |
| cm5U | 303 | 171 | -15 |
|  | 303 | 125 | -15 |
|  | 303 | 153 | -15 |
| D | 247.1 | 133 | -15 |
|  | 247.1 | 115 | -16 |
|  | 247.1 | 97 | -11 |
| G | 284.1 | 152 | -13 |
| Gm | 298.1 | 152 | -11 |
| hm5C | 274.1 | 142 | -13 |
| ho5U | 261.2 | 129 | -12 |
| ho8A | 284.5 | 152 | -11 |
|  | 284.5 | 125 | -22 |
| I | 269.1 | 137 | -13 |
| i6A | 336.15 | 204 | -25 |
|  | 336.15 | 148 | -19 |
|  | 336.15 | 136 | -22 |
| io6A | 352.2 | 220 | -15 |
|  | 352.2 | 136 | -12 |
|  | 352.2 | 118 | -15 |
| m1A | 282.1 | 150.1 | -13 |
| m1G | 298.1 | 166 | -14 |
| m1I | 283.1 | 151 | -18 |
| m2,2G | 312 | 180 | -14 |
| m227G | 326.15 | 194 | -15 |
| m27G | 312.1 | 180 | -14 |
| m2A | 282.1 | 150 | -12 |
| m2G | 298 | 166 | -13 |
| m3C | 258.1 | 126 | -12 |
| m3U | 259 | 127 | -12 |
| m3Um | 273.1 | 127 | -20 |
| m5C | 258 | 126 | -12 |
| m5U | 259 | 127 | -30 |
| m6,6A | 296 | 164.1 | -22 |
| m6,6Am | 310.15 | 164 | -20 |
| m6A | 282.1 | 150.1 | -13 |
| m6Am | 296 | 150 | -12 |
| m7G | 298.3 | 166 | -10 |
| mcm5s2U | 333.1 | 201 | -10 |
|  | 333.1 | 169 | -15 |
|  | 333.1 | 141 | -12 |
| mcm5U | 317.1 | 185 | -15 |
|  | 317.1 | 153 | -14 |
|  | 317.1 | 125 | -15 |
| mcm5Um | 331 | 185 | -17 |
|  | 331 | 125 | -17 |
|  | 331 | 153 | -16 |
| mo5U | 275.2 | 143 | -13 |
| ncm5U | 302.1 | 170 | -15 |
|  | 302.1 | 153 | -14 |
|  | 302.1 | 125 | -13 |
| oxo8G | 300.1 | 168.1 | -14 |
| PreQ1base | 180 | 66 | -15 |
|  | 180 | 91 | -16 |
|  | 180 | 121 | -17 |
|  | 180 | 146 | -15 |
|  | 180 | 163 | -16 |
| preQ1 | 312 | 180 | -11 |
|  | 312 | 295 | -11 |
|  | 312 | 163 | -11 |
| Y | 245.05 | 209 | -21 |
|  | 245.05 | 179 | -19 |
|  | 245.05 | 155 | -11 |
| Qbase | 278.1 | 146 | -13 |
| Q | 410.2 | 295 | -11 |
|  | 410.2 | 163 | -11 |
| R-mchm5U | 333 | 183 | -17 |
|  | 333 | 201 | -17 |
|  | 333 | 297 | -17 |
| s2C | 260.3 | 128 | -12 |
| S-mchm5U | 333 | 183 | -17 |
|  | 333 | 201 | -17 |
|  | 333 | 297 | -17 |
| t6A | 413.1 | 281 | -15 |
| U | 245.1 | 113 | -19 |
| Um | 259.1 | 113 | -12 |

| Target Collision Energy (V) | Target Q3 Pre Bias (V) | Ret. Time (min) |
| --- | --- | --- |
| -16 | -15 | 12 |
| -10 | -16 | 12 |
| -27 | -12 |  |
| -16 | -25 | 4.5 |
| -22 | -22 |  |
| -19 | -16 |  |
| -15 | -15 | 15 |
| -13 | -12 | 4 |
| -11 | -13 | 9 |
| -9 | -19 | 8 |
| -24 | -24 |  |
| -16 | -17 |  |
| -7 | -10 | 3 |
| -7 | -13 |  |
| -19 | -22 |  |
| -12 | -17 | 9.5 |
| -12 | -11 | 11 |
| -11 | -15 | 4 |
| -10 | -22 | 3.5 |
| -14 | -16 | 12.5 |
| -36 | -14 | 12.5 |
| -11 | -15 | 9 |
| -17 | -15 | 21.5 |
| -23 | -17 |  |
| -33 | -26 |  |
| -17 | -25 | 17.5 |
| -35 | -27 |  |
| -45 | -28 |  |
| -18 | -16 | 4.5 |
| -14 | -18 | 11.5 |
| -10 | -17 | 10 |
| -14 | -19 | 14 |
| -15 | -21 | 12 |
| -16 | -13 | 10 |
| -17 | -17 | 15 |
| -13 | -18 | 10 |
| -14 | -14 | 4 |
| -11 | -14 | 10 |
| -10 | -14 | 15 |
| -12 | -14 | 8 |
| -11 | -14 | 8 |
| -25 | -20 | 18 |
| -25 | -20 | 19.5 |
| -20 | -16 | 16 |
| -19 | -28 | 17.5 |
| -15 | -18 | 7 |
| -9 | -23 | 15 |
| -18 | -12 |  |
| -28 | -15 |  |
| -9 | -10 | 11.5 |
| -18 | -16 |  |
| -28 | -21 |  |
| -10 | -21 | 13.5 |
| -26 | -24 |  |
| -19 | -30 |  |
| -12 | -15 | 9 |
| -10 | -18 | 4.5 |
| -17 | -11 |  |
| -29 | -28 |  |
| -16 | -18 | 10 |
| -39 | -27 | 4.5 |
| -36 | -19 |  |
| -26 | -20 |  |
| -26 | -25 |  |
| -13 | -17 |  |
| -14 | -20 | 7.5 |
| -14 | -20 |  |
| -14 | -20 |  |
| -9 | -24 | 3 |
| -11 | -19 |  |
| -13 | -26 |  |
| -38 | -15 | 9 |
| -13 | -15 | 9 |
| -25 | -12 |  |
| -16 | -21 | 9.5 |
| -9 | -23 |  |
| -8 | -22 |  |
| -11 | -14 | 5.5 |
| -16 | -21 | 8.5 |
| -9 | -23 |  |
| -8 | -22 |  |
| -35 | -15 | 14.5 |
| -9 | -20 | 6 |
| -9 | -13 | 10.5 |

**Table S11.**
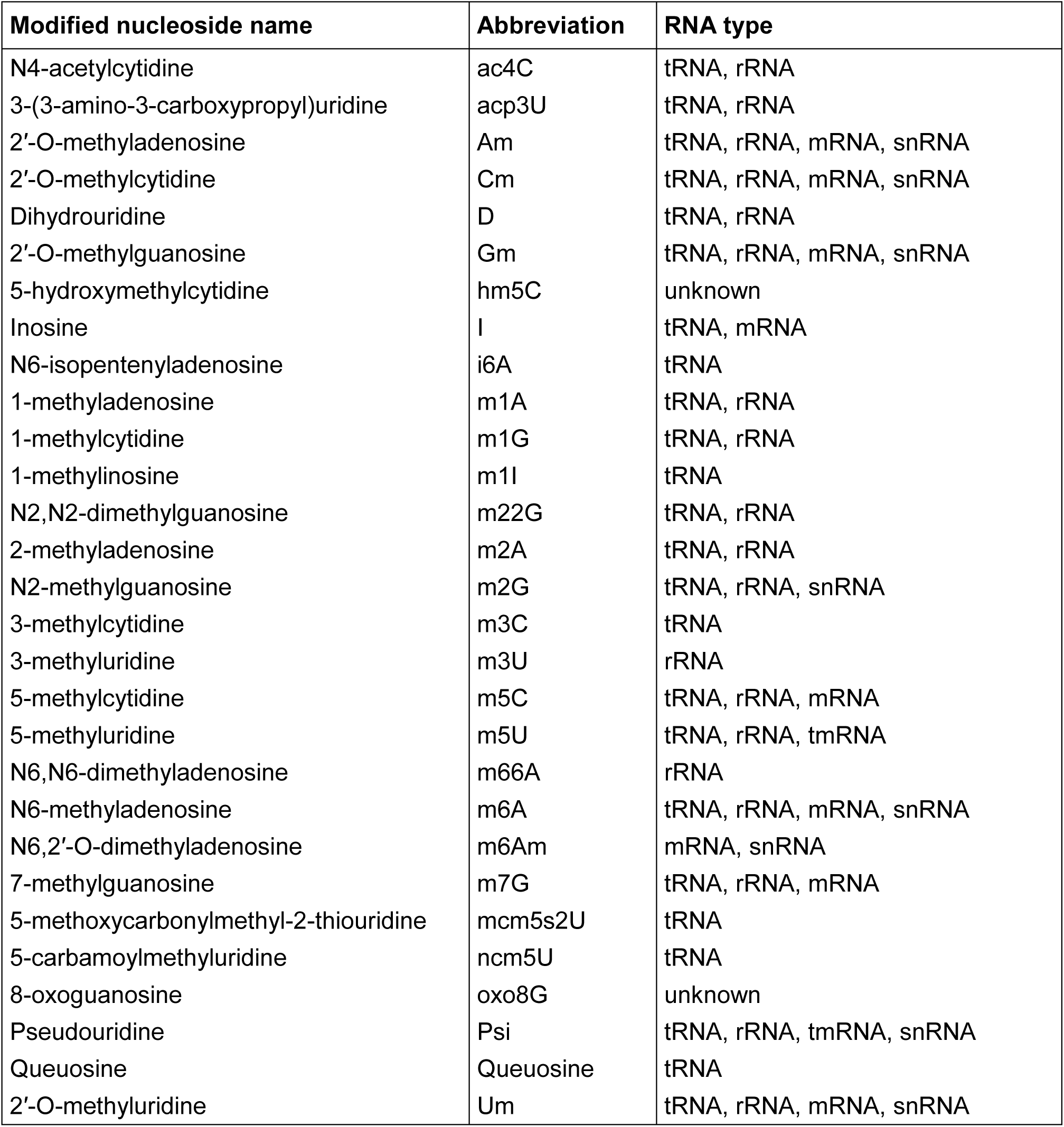
List of all modified nucleosides detected by LCMS.

